# Spatial Proteomics of the Normal Breast Collagen Stroma: Links to BI-RADS Categories and Body Mass Index

**DOI:** 10.1101/2025.07.01.660922

**Authors:** Jaclyn B Dunne, Denys Rujchanarong, Yeonhee Park, Jade K Macdonald, Taylor S Hulahan, Harrison B. Taylor, Laura Spruill, Heather Jensen-Smith, Michael Anthony Hollingsworth, George E Sandusky, Anand S Mehta, Richard R Drake, Marvella E Ford, Harikrishna Nakshatri, Peggi M Angel

## Abstract

Collagen breast stroma is the basis of increased breast density and a well-established breast cancer risk factor, yet proteomic regulation of normal breast stroma remains poorly defined. This study reports spatial regulation of the collagen proteome in normal breast tissue sections annotated by clinical characteristics. Normal breast samples from the Susan G. Komen tissue bank included data on genetic ancestry (n=40 total; n=20 African ancestry; n=20 European ancestry), body-mass-index (BMI), age, and mammogram density by the Breast Imaging Reporting and Data System (BI-RADS). Multiplexed cell marker staining showed CD44 and COL1A1 markers modulated with BMI. Collagen fiber widths by second harmonic generation (SHG) showed potential contrasts in BMI categories by genetic ancestry. Targeted extracellular matrix proteomics mass spectrometry imaging showed collagen alpha-1(I) chain domain proteome was spatially heterogenous across the normal breast microenvironment with site specific post-translational modification of proline hydroxylation. Signatures computationally extracted from breast stroma reported that 47 collagen peptides distinguished BI-RADS categories (area under the receiver operating curve>0.7; p-value>0.05). Proteomic alterations were found between overweight to obese categories with strong positive associations to BMI by multivariate analysis. This study provides the first spatial analysis of the collagen proteome in normal breast within contexts of cellular markers and clinical characteristics.

## Background

Breast stroma is evaluated throughout the course of clinical care, where an increased risk of breast cancer is related to increased stromal density(1–5). Mammographic imaging shows that breast stroma emerges as filamentous strands that in high cancer risk(6) to become dense fields of connective tissue throughout the breast microenvironment. Lifestyle and hereditary factors such as obesity, or having body mass index above 30(7), further increase the risk of cancer through diverse multiomic changes(8). At a molecular level, stroma forms the cell-matrix interface for cell recruitment, movement, and localization within the breast microenvironment(9–11). Recent studies have shown that the cellular microenvironment within the normal breast is a heterogenous mixture of perivascular, endothelial, numerous luminal epithelial cell types, immune cells, and fibroblast-like cells, all varying over age and ethnicity(12–15). These detailed studies provide a wealth of information on genetic and transcriptomic mechanisms of the breast microenvironment leading to emergent tumors. A significant challenge remains in that while stroma is used to assess for clinical risk, this remains uncharacterized at the molecular level as a largely proteinaceous composition, rich in post-translational modifications.

Breast stroma is composed of collagen type proteins that form collagen fiber superstructures shaped by extracellular glycoprotein modulation(16). The organization of collagen fibers is predictive of breast cancer outcomes and is altered by therapies(17–19), implicating collagen protein regulation as a fundamental breast biology that can lead to a tumor permissive or therapeutically resistant microenvironment. Fibrillar collagens of type I-III, V and XI, are the primary collagen type proteins of breast, presenting complex network throughout the breast microenvironment under reproductive control(20) with multiple and diverse domains that directly interface with cells to alter cell function (21, 22). Cell interactive domains are dispersed within the tissue microenvironment as bundles of triple helical constructs, composed of three intertwined polypeptide domains from multiple fibrillar collagen proteins (23). Post-translational modification of triple helical domains through hydroxylation of proline (HYP) serves to control exposure or concealing of cell interactive domains(21, 22, 24) and presents a heterogenous network in breast stroma changing with invasive breast cancer(25). Proline hydroxylases, enzymes essential to forming HYP sites(26), regulate collagen synthesis (27), increasing with breast cancer progression and forming components of therapeutic resistance(28–30) that are reduced with inhibition(31, 32). Fibrillar collagen protein types contain several hundred prolines that may be modified or unmodified. For instance, collagen alpha-1(I) chain (COL1A1) has 283 prolines that may be variably modified; collagen alpha-2(I) chain (COL1A2) has 237 proline sites that may be variably modified. Critically, both collagen protein composition and HYP site modifications present significant cell-interactive mechanisms of the breast microenvironment that remain largely undefined.

Obesity increases breast connective tissue, creating a more favorable microenvironment for proliferation and emergence of breast cancer(33, 34). Evidence of increased BMI has been associated with an increase in postmenopausal breast cancer risk(35). Additionally, longitudinal studies show that longer duration, greater severity, and earlier onset of obesity are strongly associated with postmenopausal breast cancer risk(36). There is sufficient evidence for the contribution of obesity to breast cancer in postmenopausal women that nearly a decade ago, physician working groups recommended the avoidance of weight gain as breast cancer risk control(35). Obesity intersects with genetic ancestry within cancer risk and breast cancer outcomes. Obese women self-identified as Black (with presumed African ancestry) are more frequently diagnosed with aggressive triple-negative breast cancer (TNBC) and at a younger age compared to self-identified White counterparts (with presumed European ancestry)(37, 38). The effect of obesity is well-documented in the TNBC breast microenvironment, including impairment of immune response(39–41), meta-inflammation increases(34, 42), and altered regulation of cytokines and adipokines(43, 44) that propel stroma modulation in tumor invasion and metastasis(39, 45). As of yet, few studies exist to understand how BMI influences the normal breast microenvironment from the standpoint of genetic ancestry and collagen stroma modulation.

In this study, we investigate the hypothesis that the normal breast microenvironment contains a gradient composition of fibrillar collagens altered by risk factors, particularly BMI, that also associate with breast cancer. The study leverages a unique cohort from the Susan G. Komen Tissue Bank derived from healthy female donors (n=40) and clinically characterized for risk using the Breast Imaging System as well as risk algorithms Gail and Tyrer-Cuzick. Key cellular expression patterns, collagen fiber measurements, and collagen proteomic regulation including gradients of HYP site modifications are used with clinical characteristics age, ancestry and BMI to understand dynamic collagen proteome regulation in the normal breast microenvironment. This study provides a foundation for understanding the molecular characteristics of normal breast stroma that may increase breast cancer risk.

## Methods & Materials

### Materials

100% ethanol (pharmaceutical grade), ammonium bicarbonate for LC-MS (40867-50G-F), acetic acid glacial, bovine serum albumin, Advanced PAP Pen (5-mm tip width), 0.5 mL Ultrafree-MC centrifugal 0.45-µm filter devices, and octyl ß-d-glucopyranoside (OBG) (≥95% (HPLC), 50% (w/v) in H_2_O) were from Millipore Sigma (Burlington, MA/St. Louis, MO); xylenes (semiconductor grade) and water (LC-MS grade) were from Thermo Scientific (Waltham, MA); chloroform (HPLC Grade) was from Alfa Aesar (Haverhill, MA); isopropanol (Optima™ LC-MS Grade) was from Fisher Chemical™ (Hampton, NH); tris base (molecular biology grade) and tris-HCl (molecular biology grade) were from Promega (Madison, WI); and antigen retrieval buffer (100× Tris-EDTA buffer, pH 9.0) was from ABCAM (Cambridge, MA). Trifluoroacetic acid, acetonitrile, and α-cyano-4-hydroxycinnamic acid were purchased from Sigma Aldrich (St. Louis, MO, USA). Collagenase type III (COLase3) was from Worthington Biochemical (Lakewood, NJ, USA) and PNGase F PRIME™ was from N-Zyme Scientifics (Doylestown, PA, USA).

### Human Tissue

Formalin-fixed, paraffin-embedded (FFPE) normal breast tissues were obtained through collaboration with the Susan G. Komen Tissue Bank at Indiana University Simon Cancer Center (46). Analysis was done on the tissues with MUSC IRB permissions. Samples from 20 women of African Ancestry (AA) and 20 of European Ancestry (EA) were annotated with clinical characteristics including genetic ancestry, age, body mass index (BMI), and breast density based on Breast Imaging Reporting and Database System (BI-RADS) scoring. Although Body Mass Index is an imperfect measure of obesity, we used this clinical characteristic to group the women into the following categories based on the National Institute of Health (NIH) definitions: Normal (18.5 to 24.9 kg/m^2^), Overweight (25 to 29.9 kg/m^2^), or Obese (> 30 kg/m^2^). Genetic ancestry was defined for each donor (**Figure 1**) using a panel of 41 genetic polymorphisms, called Ancestry Informative Markers (AIMs), to determine ancestry percentage by continent(47). Breast cancer risk was defined by Gail Score and Tyrer-Cuzick models, which consider several risk factors for breast cancer(48–50).

**Figure 1.**
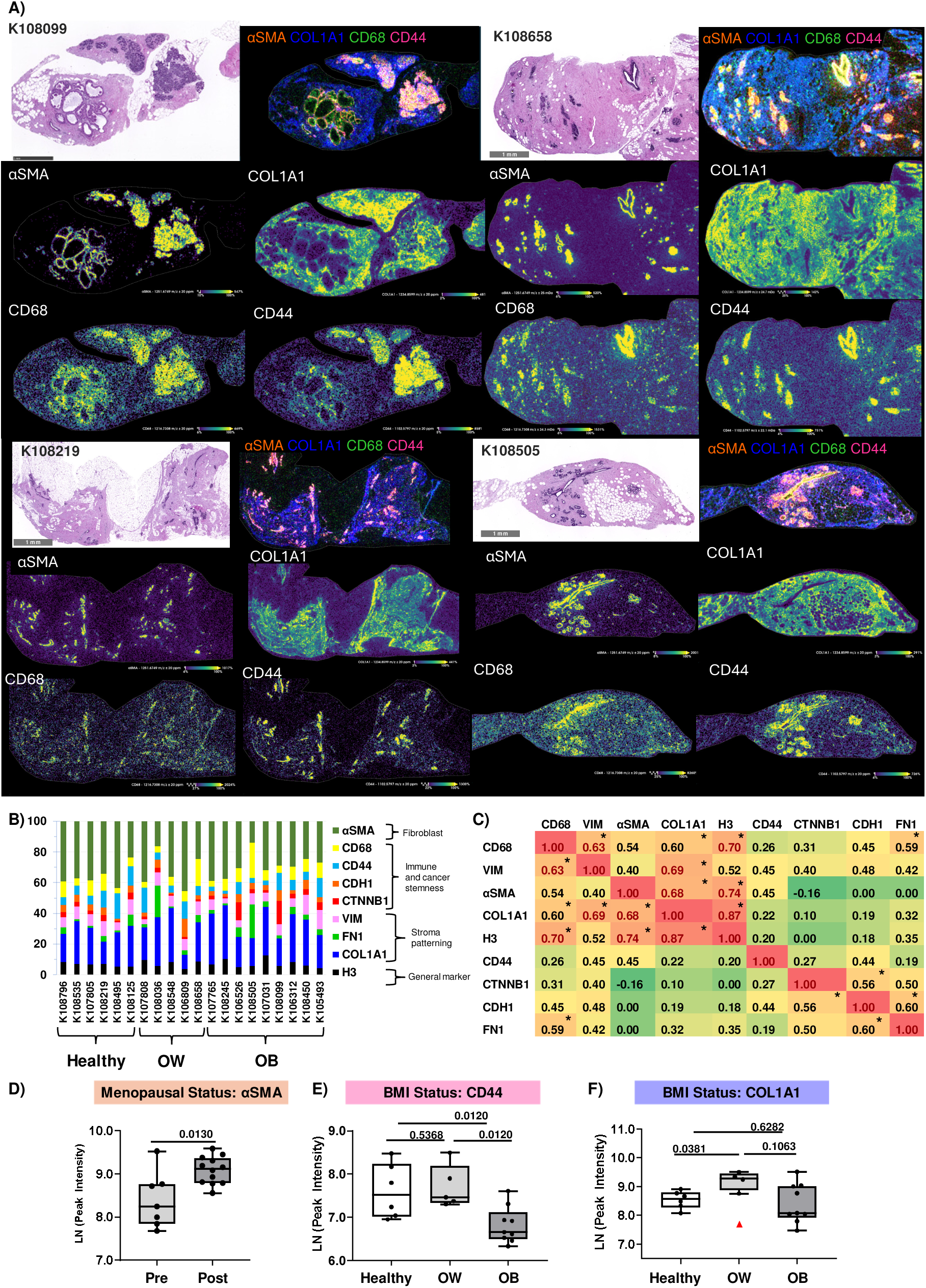
The normal breast microenvironment shows a complex cellular and extracellular expression. Example images of αSMA, COL1A1, CD68 and CD44 localization on normal breast. CD44 and CD68 were detected around ductal glands. B) COL1A1 and alpha-SMA represented high proportions of staining within stroma regions. C) Correlation by intensity demonstrated significant correlations of COL1A1 with CD68, VIM, and aSMA. Spearman’s correlation r is shown; * = p-value <0.01. D) Menopausal status showed a significant increase in postmenopausal stroma-rich breast. E) CD44 decreased with high BMI status. F) COL1A1 showed an increase in detection in samples connected with a midrange BMI. Outlier is demarked by a red triangle; Outlier method is ROUT (Q=10%). E,F: Mann-Whitney U p-value is shown. BMI: <25 conventionally classified as Healthy; midrange BMI 25-30, conventionally classified as overweight; High BMI, >30, conventionally classified as obese.

### MALDI-IHC Workflow

A subset of 20 normal breast tissues (10 AA & 10 EA) was selected based on the highest stromal content for staining with photocleavable-mass tags (PC-MTs; MALDI-IHC). The protocol for MALDI-IHC by AmberGen, Inc. was followed (51). In brief, FFPE tissues were melted at 60 °C for 2 hours followed by deparaffinization (xylenes 3x 5 minutes, 50% xylenes/50% ethanol 3 minutes) and rehydration (100% ethanol 2x 2 minutes, 95% ethanol 3 minutes, 70% ethanol 3 minutes, 50% ethanol 3 minutes, and tris-buffered saline 10 minute) washes. Tissues were then antigen retrieved in 1x tris-EDTA, pH 9.0 for 30 minutes at 95 °C. Blocking buffer (2% normal serum from mouse and rabbit and 5% BSA in TBS-OBG [0.05% OBG]) was applied on tissues for 1 hour. A PAP pen was used to encircle the tissue before antibody application. A panel of 10 PC-MT antibodies (AmberGen, Inc.) was combined into a master mix with 3.75 ug/mL of each antibody (diluted in blocking buffer). This PC-MT antibody master mix was applied to tissues and incubated for 1 hour at 38 °C, protected from any light. Slides were then washed in TBS (3x 5 minutes) and 50 mM LC-MS grade ammonium bicarbonate (3x 2 minutes) in a petri dish with gentle shaking. Tissues were dried for 1 hour in a vacuum desiccator and stored at 4 °C until ready for mass spectrometry imaging. Before MALDI-MSI PC-MTs must be photocleaved for 10 minutes with 3 mW/cm^2^ of 365-nm light in a UV Illumination Box (AmberGen, Inc). CHCA matrix (10 mg/mL in 70% acetonitrile, 0.1% TFA, 10 mM ammonium phosphate) was sprayed on tissue using an HTX M5 automated sprayer with the following parameters: 60 °C, 0.1 mL/min flow rate, 1350 mm/min velocity, 8 passes, 3 mm track spacing, 40 mm nozzle height, 10 PSI, 2 L/min gas flow, 10 s drying time. Matrix was recrystallized with 1.0 mL 5% isopropyl alcohol for 1 minute in a petri dish with filter paper (Whatman™ Qualitative Grade Plain Circles, Thermo Fisher Scientific).

### MALDI-TOF Mass Spectrometry Imaging of Photocleavable-Mass Tags

MALDI-IHC PC-MTs were detected using a MALDI-QTOF (timsTOF flex, Bruker) in positive ion mode from 700-2000 m/z. Tissues were scanned using a 20 µm step size with a laser spot size of 14 µm (resulting field size of 18 µm), 80 µs transfer time, and 20 µs pre-pulse storage. Data were uploaded to SCiLS Lab 2024 for evaluation of mass tags’ spatial localization.

### Second Harmonic Generation

High contrast images of endogenous collagen second harmonic generation (SHG) were collected at the Multiphoton Intravital and Tissue Imaging (MITI) core at University of Nebraska Medical Center using an upright Olympus FVMPE-RS microscope equipped with a 25x (1.05 NA) objective. A Spectra-Physics InSight X3 laser tuned 860 nm with a pulse width of ∼120 fs and 80 MHz repetition rate was used to excite collagen SHG. SHG emission (backscatter) was collected from individual images taken at 2 μm intervals throughout each 5 μm section using a 432 nm (45 nm bandpass) emission filter and zoom of 1.2. To optimize the visualization of collagen fiber lengths, 6 mm image stacks were compressed (max intensity projection) using NIH ImageJ (52). Images were acquired from 2-6 discrete locations of uniform sizes (509.12 mm x 509.12 mm x 5 mm) depending on what was present in each sample. Measurements were classified as either ductal (presence of duct) or fibrotic (absence of ductal features). Stromal measurements from a single patient were averaged together for a representative fiber characteristic for either ductal or fibrotic categories. Collagen fiber counts and organization (fiber length, width, curvature, alignment, orientation) were quantified in individual SHG images using CT-FIRE and CurveAlign for Fibrillar Collagen Quantification (53–55) in areas annotated by a surgical pathologist.

### Tissue Preparation for Collagen-Targeted MALDI-MSI

FFPE tissues were processed to remove N-glycans by previously published protocols (56, 57). In brief, these tissues were deparaffinized, rehydrated, and antigen retrieved using 10 mM citraconic anhydride (pH 3) before PNGase F PRIME™ (N-zyme Scientifics) application (M5 HTX Technologies Sprayer) using the following parameters: 15 passes, 45 °C, 10 psi, 25 mL/min, 1200 velocity, 40 mm nozzle height, and 2.5 mm offset. Tissues were placed in individual, preheated humidity chambers (37 °C) for a 2-hour enzymatic digestion. The cleaved N-glycans were then washed off the tissue by a series of ethanol, water, high pH, and low pH rinses. A second antigen retrieval was then performed using 10 mM tris buffer (pH 9) for 25 minutes at 95 °C (vegetable steamer). Collagenase type III (Worthington) was prepared at 0.01 mg/mL in 10 mM ammonium bicarbonate with 1 mM CaCl_2_ and sprayed by HTX Technologies M5 automated sprayer (15 passes, 45 °C, 10 psi, 25 mL/min, 1200 velocity, 40 mm nozzle height, and 2.5 mm offset) followed by digestion (5 hours at 37 °C). CHCA matrix was prepared (7 mg/mL in 50% acetonitrile and 1% TFA with Glu-1-Fibrinopeptide spiked in at 0.15 picomole and sprayed for 10 passes, 79 °C, 10 psi, 70 mL/min, 1300 velocity, 40 mm nozzle height, 3.0 mm offset. The slides were then dipped once in cold 5 mM ammonium phosphate and dried in a desiccator. Detailed tissue preparation specifications can be found in previously published protocols (58).

### Mass Spectrometry Imaging

Prepared normal breast tissues was analyzed by Matrix-Assisted Laser Desorption/Ionization – Mass Spectrometry Imaging (MALDI-MSI) using a trapped ion mass spectrometer quadrupole time of flight (Bruker timsTOF flex) in positive ion mode. Tissues were sampled using a 60 µm step size with 300 laser shots per pixel and a 18 µm laser size (22 µm resulting field size). The m/z range was set to 600-2500 with a transfer time of 75 µs and a pre-pulse storages of 20 µs, respectively. MALDI-MSI data was imported into SCiLS Lab 2025a and normalized to the internal standard. Stromal regions were identified by pixel classification using QuPath digital pathology software(59). QuPath annotations were applied into SCiLS Lab via QuPath to SCiLS Lab plug-in. Maximum peak intensities from stromal regions classified in QuPath were exported for analyses. Peptides identified by LC-MS/MS were used to perform image segmentation based on K-means algorithm using Manhattan distance. Plots displayed were made in GraphPad software v10.4.1 and analyses were done in MetaboAnalyst 6.0(60).

### Tissue Preparation for Extracellular Matrix-Targeted LC-MS/MS Proteomics

A serial section of normal breast tissue was used for extracellular matrix-targeted LC-MS/MS proteomics targeting stroma. Tissues were heated at 60 °C for one hour and deparaffinized before being scraped from microscope slides into individual centrifuge tubes. A 200 µL volume of 25 mM ammonium bicarbonate, 3mM calcium chloride, pH 7.4 was added to the tubes, and the tissues were spun down using a benchtop centrifuge at 3,000 rpm for 1 min. Ultrasonication was performed in a cool water bath on a closed tube capped with parafilm with the following parameters: 35% amplitude, 5 minutes with 20 seconds pulse time on and 5 seconds off until fragmentation of the tissue was observed. Collagenase type III (5 µg) was added to the samples and digested overnight at 38 °C with 450 rpm shaking. Following overnight digestion, tissues were sonicated for 30 minutes, and an additional aliquot of collagenase type III (2 µg) was added for a 5-hour digestion at 38 °C with 450 rpm shaking. The tissues were then pelleted for 10 mins at 14000 rpm. The supernatant was collected and vacuum-concentrated until dry. Samples were cleaned using a 10-µg capacity C18 StageTip (Pierce, Thermo Fisher Scientific). After the eluent was dried, samples were further processed using a 2-ug capacity C18 ZipTip (Millipore), a step we have found helpful when dealing with FFPE tissues.

### LC-MS/MS Extracellular Matrix Peptide Sequencing

Pooled samples were injected in sextuplicate (200 ng) through a NanoElute coupled to a timsTOF FleX (Bruker Daltonics, Bremen Germany). Peptides were separated with a 40-minute gradient from 2% acetonitrile, 0.1% formic acid (mobile phase A) to 30% acetonitrile, 0.1% formic acid (mobile phase B) through an Aurora Ultimate Ionopticks C18 column (75 µm inner diameter, 25 cm length, 1.7 µm particle size, 120 Å pore size). Data dependent acquisition was performed in positive ion mode over a 150-1700 m/z mass range, 0.5-1.85 1/K0 trapped ion mobility range, 0.96 s cycle time with 8 PASEF ramps per cycle, 60 µs transfer time, and a pre pulse storage of 12 µs. Ion mobility and m/z were calibrated using a m/z = 622, 922, 1221 calibrant filter standard (Low concentration ESI-L LCMS Tuning Solution, Agilent). System quality was assessed using HeLa Protein Digest Standard (Pierce, thermos Fisher Scientific) as a control at the beginning and end of the run.

Data was searched using Fragpipe v20.0 and MSFragger v3.8 using a nonspecific digest, fragment tolerance of 40 ppm and parent tolerance of 25 ppm, variable methionine oxidation, proline hydroxylation, and asparagine or glutamine deamidation with a protein and peptide false discovery rate set to 0.01 (61–71). Data was searched against a curated SwissProt-reviewed database downloaded from UniProt on January 24, 2024, containing 1,020 entries and using search terms “homo sapiens” AND subcellular location search terms extracellular matrix, SL-0111. Reverse decoys and contaminants were added to the database using FragPipe v20.0. PSM files were used to report peptide domains using a hyperscore cutoff of >18, peptide probability >0.95 and single-site proline modification >0.8 (**Supplemental Table 1**). Image data was matched to peptide domain sequence within 10 ppm. Data were uploaded into Scaffold (v5.3.1) and protein quantitative values were evaluated using normalized average precursor intensities (**Supplemental Figure 1**).

## Results

### The normal breast microenvironment is defined by pathological markers within collagen-rich stroma

To understand the mechanisms of the normal breast microenvironment, we first sought to define cellular expression with extracellular morphological patterns within the stroma. A total cohort of samples from 40 women with genetic ancestry classified as African ancestry >70% (AA = 20) or European ancestry >50% (EA = 20) was curated from the Susan G. Komen Tissue Bank based on intermediate risk for breast cancer defined by the Gail score and Tyrer-Cuzick score (**Table 1**). On a subset of 20 tissues with greater than 20% stroma present by area per H&E, multiplexed immunohistochemistry using cell and extracellular markers associated with breast cancer outcomes was used to assess the status of each tissue. By antibody staining, stroma rich regions showed intense staining for COL1A1 with co-localized CD44+ and αSMA+ myoepithelial cells amid a milieu of CD68+ macrophages surrounding ductal regions (**Fig. 1A**). Surprisingly, markers of poor prognosis in breast cancer, including vimentin (VIM; 7.4 ± 2.5%), cadherin (CDH1; 4.2 ± 2.4%), and beta catenin (CTNNB1; 3.6 ± 3.6%) showed expression within the normal breast microenvironment with heterogeneity across samples (**Fig. 1B, Supplemental Table 2, Supplemental Figure 2, 3**). Measurement of intensity per stroma-rich area showed positive correlation of COL1A1 with αSMA, VIM, and CD68 (**Fig. 1C**). Genetic ancestry-dependent evaluation of markers showed no significant alterations per clinical characteristics. However, αSMA was increased in postmenopausal women (**Fig. 1D**). Body mass index (BMI) showed a decrease in CD44+ myoepithelial cells associated with high BMI (**Fig. 1E**). COL1A1 demonstrated an increase in overweight BMI (BMI 25-30 kg/m^2^) (**Fig. 1F**). These results show that the normal breast stroma carries complex cellular signatures previously associated with pathological processes of breast cancer.

**Table 1.** Cohort characteristics are statistically comparable when investigating differences in clinical categories based on genetic ancestry. The cohort is split by genetic ancestry to show similarities between the two groups. The last column displays p-values when comparing women of African vs. European ancestry for each category (t-test for first 5 rows, asterisk * indicates p-value calculated by chi-square test). The characteristics for age, BMI, Gail Score, and Tyrer-Cuzick scores are shown as the average value with 95% confidence interval.

| Parameters | African Ancestry<br>[± 95% CI] | European Ancestry<br>[± 95% CI] | P-value |
| --- | --- | --- | --- |
| Age | 59.4 [55.9, 62.9] | 55.3 [51.9, 58.6] | 0.10 |
| Body Mass Index | 28.1 [25.5, 30.7] | 29.4 [27.5, 31.4] | 0.43 |
| Gail Score | 7.3 [5.5, 9.0] | 7.1 [5.2, 8.9] | 0.88 |
| Tyrer-Cuzick Score<br>(10 year) | 4.2 [3.0, 5.5] | 3.6 [2.5, 4.7] | 0.45 |
| Tyrer-Cuzick Score (Lifetime) | 9.9 [7.0, 12.8] | 10.7 [7.5, 13.9] | 0.71 |
| BI-RADS<br>(Breast Imaging Reporting and<br>Data System) | (1) Negative = 12<br>(2) Benign = 7<br>(3) Probably benign = 1<br>(4) Suspicious abnormality = 0 | (1) Negative = 7<br>(2) Benign = 12<br>(3) Probably benign = 0<br>(4) Suspicious abnormality= 1 | 0.20† |
| Breast Density | Scattered fibroglandular = 7<br>Mixed fatty & fibroglandular = 0<br>Mild Density = 2<br>Heterogeneously dense= 10<br>Dense Fibroglandular = 0<br>Extremely dense = 2 | Scattered fibroglandular = 5<br>Mixed fatty & fibroglandular = 1<br>Mild Density = 0<br>Heterogeneously dense = 12<br>Dense Fibroglandular = 1<br>Extremely dense = 0 | 0.27† |
| Menopausal status | Premenopausal = 4<br>Postmenopausal = 16 | Premenopausal = 7<br>Postmenopausal = 12 | 0.24† |

### Collagen fiber width in the normal breast microenvironment is altered in association with clinical characteristics

Previous studies have shown a systematic alteration in collagen fiber organization in breast cancer predictive of poor prognoses(18). To understand collagen fiber regulation within the normal breast microenvironment, collagen fiber measurements were investigated across the cohort of normal breast tissue sections (n=40). Hematoxylin & Eosin (H&E) stained tissue sections were imaged by second harmonic generation (SHG) microscopy to visualize and measure collagen fiber characteristics within the normal breast (**Fig. 2A**). The measures were assessed by different clinical characteristics including ancestry, age, Body Mass Index (BMI), menopausal status, BI-RADS category, breast density categories (**Supplemental Table 3**). There were no significant differences in fiber length or angle of orientation (from ductal regions) in any category. Collagen fiber width appeared to decrease with age when women were categorized into groups below (n = 17) and above (n = 19) 55 years old (**Fig. 2B**). When comparing collagen widths by BMI status independent of ancestry, there were no significant differences in ductal nor fibrotic regions (**Fig. 2C**). However, when genetic ancestry was considered, collagen fiber width altered significantly in comparisons of overweight and obese women for both ductal and fibrotic regions (**Table 2**). Overweight EA women showed significantly wider collagen fibers compared to overweight AA women in both ductal and fibrotic regions (**Fig.2D**). Interestingly, this trend switched in samples associated with obese BMI, showing AA women have significantly wider collagen fibers than EA women (**Fig. 2E; Supplemental Figure 4**). Collagen fiber widths varied by BMI and genetic ancestry, with obese AA donors and overweight EA donors showing wider fibers in ductal and fibrotic regions than their overweight and obese counterparts (**Fig. 2F, 2G**, respectively; **Supplemental Figure 4**). These findings indicate that the normal breast microenvironment exhibits dynamic changes in collagen fiber structure associated with obesity, potentially influenced by genetic ancestry.

**Figure 2.**
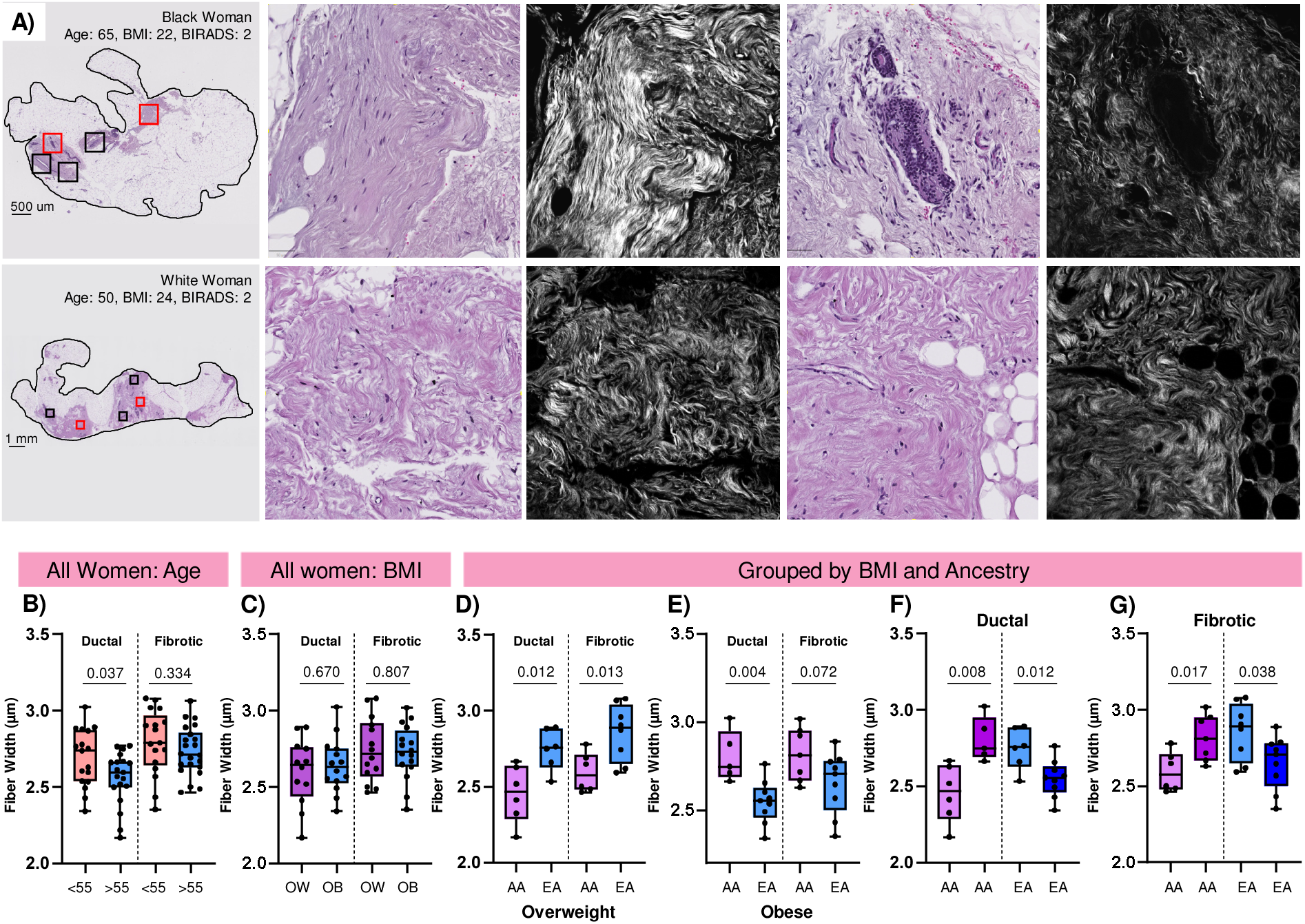
Collagen fiber measurements alter by clinical characteristic in the normal breast microenvironment. Second harmonic generation microscopy (SHG) measurements were recorded using CT-FIRE and CurveAlign for 2-6 different locations of uniform sizes per tissue sample (n = 20 AA; n = 20 EA). All stromal measurements from a single patient were averaged together for a representative fiber characteristic in either ductal or fibrotic regions. A) Example whole tissue H&E stain of a normal breast tissue section from a donor of African Ancestry and a donor of European Ancestry. Black boxes indicate all areas that were imaged by SHG. Fiber measurements compared in (B-D) were compared by t-test with 95% confidence. B) Ductal collagen fiber widths tend to decrease with age in the normal breast microenvironment. C) All samples in different BMI categories show similar collagen fiber widths. When samples are categorized by BMI and ancestry status, differences in collagen widths are present. D) In samples with midrange BMI (BMI 25-30, conventionally overweight status), EA showed an increase in collagen fibers while E) Black women with obese BMI have wider fibers F-G) Comparisons between samples associated with overweight and obese status show an increase in BW collagen fiber width for both F) ductal and G) fibrotic measurements.

**Table 2.**
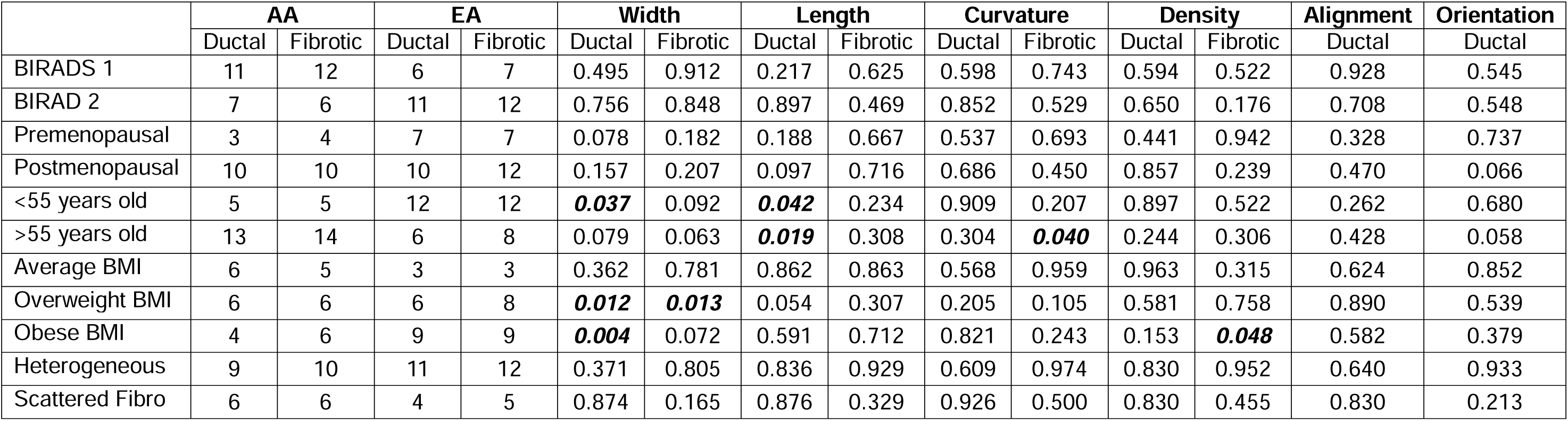
Genetic ancestry-based comparisons of collagen measurements show the differences in ductal width category by second harmonic generation with respect to different clinical characteristics. Significant differences in collagen width measurements appear when comparing overweight or obese by genetic African ancestry (AA) or European Ancestry (EA). Ancestry-based comparisons of BIRADS, menopausal status, and breast density do not show significant differences.

|  | AA |  | EA |  | Width |  | Length |  | Curvature |  | Density |  | Alignment | Orientation |
| --- | --- | --- | --- | --- | --- | --- | --- | --- | --- | --- | --- | --- | --- | --- |
|  | Ductal | Fibrotic | Ductal | Fibrotic | Ductal | Fibrotic | Ductal | Fibrotic | Ductal | Fibrotic | Ductal | Fibrotic | Ductal | Ductal |
| BIRADS 1 | 11 | 12 | 6 | 7 | 0.495 | 0.912 | 0.217 | 0.625 | 0.598 | 0.743 | 0.594 | 0.522 | 0.928 | 0.545 |
| BIRAD 2 | 7 | 6 | 11 | 12 | 0.756 | 0.848 | 0.897 | 0.469 | 0.852 | 0.529 | 0.650 | 0.176 | 0.708 | 0.548 |
| Premenopausal | 3 | 4 | 7 | 7 | 0.078 | 0.182 | 0.188 | 0.667 | 0.537 | 0.693 | 0.441 | 0.942 | 0.328 | 0.737 |
| Postmenopausal | 10 | 10 | 10 | 12 | 0.157 | 0.207 | 0.097 | 0.716 | 0.686 | 0.450 | 0.857 | 0.239 | 0.470 | 0.066 |
| <55 years old | 5 | 5 | 12 | 12 | **0.037** | 0.092 | **0.042** | 0.234 | 0.909 | 0.207 | 0.897 | 0.522 | 0.262 | 0.680 |
| >55 years old | 13 | 14 | 6 | 8 | 0.079 | 0.063 | **0.019** | 0.308 | 0.304 | **0.040** | 0.244 | 0.306 | 0.428 | 0.058 |
| Average BMI | 6 | 5 | 3 | 3 | 0.362 | 0.781 | 0.862 | 0.863 | 0.568 | 0.959 | 0.963 | 0.315 | 0.624 | 0.852 |
| Overweight BMI | 6 | 6 | 6 | 8 | **0.012** | **0.013** | 0.054 | 0.307 | 0.205 | 0.105 | 0.581 | 0.758 | 0.890 | 0.539 |
| Obese BMI | 4 | 6 | 9 | 9 | **0.004** | 0.072 | 0.591 | 0.712 | 0.821 | 0.243 | 0.153 | **0.048** | 0.582 | 0.379 |
| Heterogeneous | 9 | 10 | 11 | 12 | 0.371 | 0.805 | 0.836 | 0.929 | 0.609 | 0.974 | 0.830 | 0.952 | 0.640 | 0.933 |
| Scattered Fibro | 6 | 6 | 4 | 5 | 0.874 | 0.165 | 0.876 | 0.329 | 0.926 | 0.500 | 0.830 | 0.455 | 0.830 | 0.213 |

### Collagen domains are spatially modulated within the normal breast microenvironment

COL1A1 transcriptional and protein expression has been linked to breast cancer progression and therapeutic response(72), yet the fundamental mechanism of COL1A1 domain regulation has not been spatially investigated in normal breast. In this study, peptide domains of collagen α-1(I), including the triple-helical domain and cell-interacting domain (73), were assessed for spatial distribution throughout stroma regions (**Figure 3A**). Spatial segmentation to cluster collagen domain peptides based on intensity and localization showed the normal breast proteome aligns with histology, but shows significant heterogeneity (**Figure 3B&C, Supplemental Figure 5**). Spatial heterogeneity was also observed at the single collagen peptide domain level when visualized as intensity maps (**Figure 3D).** Generally, high intensity regions were positively correlated with stroma depicted in histology stains. However, proteomic imaging of COL1A1 domains showed that in certain tissues, expression extended to the adipocyte regions (example, **Figure 3D i, iii, vi, ix**). This expression pattern varying from stroma to adipocyte included a known cell binding region (**Figure 3D, v**). COL1A1 peptide domain distribution varied independent of hydroxylation of proline status. COL1A1 domain peptides that were unmodified showed differential patterns when compared to each other (example, **Figure 3D, i, iv, ix**). COL1A1 domains that showed all prolines as fully hydroxylated also varied in distribution, with certain domains showing highly localized expression (**Figure 3D, vi, vii, viii**) that included adipocyte regions. COL1A1 domain peptides showed spatial variation per donor (example **Figure 3D iv, v, vi**). Altogether, the triple-helical collagen peptide domains in the normal breast are spatially regulated and localize to histologically relevant, stromal and adipose-rich regions.

**Figure 3:**
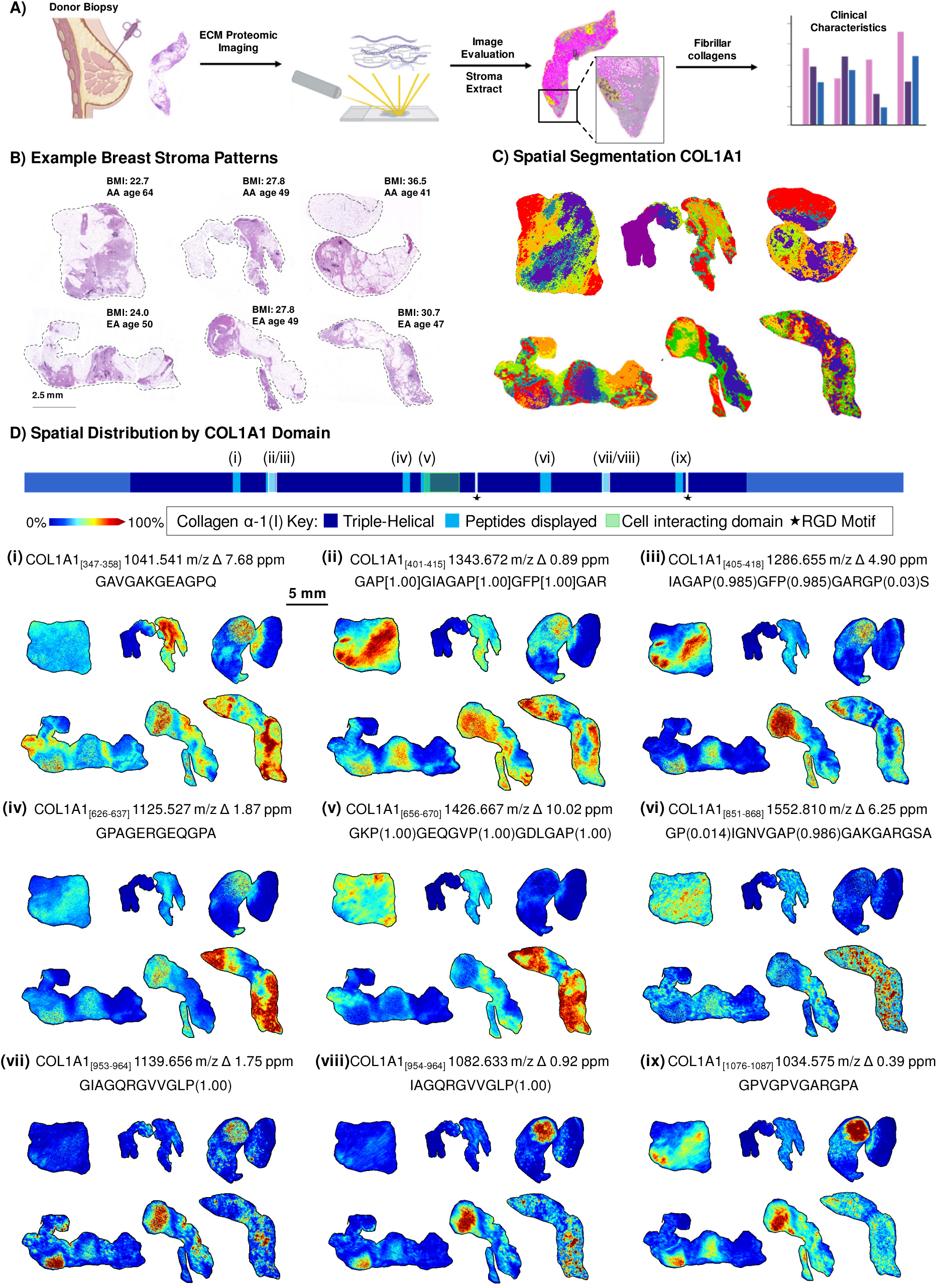
Spatial regulation of collagen domains in normal breast. A) Workflow isolating stromal signatures from normal breast histology. H&E-stained tissues were aligned with sections that were analyzed by ECM-targeted mass spectrometry imaging proteomics. QuPath was used to extract stroma specific regions from each section. Stroma extracted data were compared by clinical characteristics. B) Example breast stroma patterns shown for women with defined BMI and genetic ancestry. C) Spatial segmentation of the stromal proteome collected by mass spectrometry imaging. Colors represent different proteome signatures segmented by spatial organization and intensity. D) Annotated Collagen α-1(I) chain domain mapping. Roman numerals represent peptide domains that were mapped by mass spectrometry imaging. Domains reflect spatial regulation of the collagen network within breast stroma.

### Spatially distributed collagen domains distinguish BI-RADS breast density categories

Increased mammographic density is a significant risk factor for breast cancer, as dense breast tissue complicates tumor detection due to its radiological similarity to malignant lesions, both appearing as white regions on mammograms (1, 74). Currently, around 80% of women fall into two major breast density categories as defined by fibroglandular content, either scattered fibroglandular (SF; 25 to ≤50%) or heterogeneously dense (HD; 50 to ≤75%)(75). Using stroma pixel data that were extracted from image data by computational classification to compare stroma-only regions, scattered fibroglandular signatures were compared to heterogeneously dense signatures. Molecular characterization of the stromal content of SF (n = 12) and HD (n = 24) normal breast tissue revealed 47 peptides that significantly differentiated between these breast density categories (**Figure 4A**). Example peptide domains map to collagen α-1(I) including 981.431 m/z, 1034.575 m/z, and 1082.633 m/z and show significant Area Under the Receiver Operating Characteristic curve (AUROC; >0.73, p value < 0.02; **Figure 4B, Supplemental Figure 6**), highlighting ability to distinguish molecular features of breast density. ECM peptide intensities extracted from stroma regions were further compared among AA and EA women within either the SF (n = 7 AA, n = 5 EA) or HD (n = 10 AA, n = 14 EA). Scattered fibroglandular mammograms showed expected sparse fibroglandular mapping (**Figure 4C**) correlating with sparse stroma seen on histology stains by hematoxylin and eosin (**Figure 4D**). Two peptide domains were compared by ancestry within each category; image data reflected that the stroma signal altered within stroma regions that extended into the adipose regions. (**Figure 4E, Supplemental 7**). Mammograms from breasts categorized as heterogeneously dense showed expected similar compositions of dense glandular tissue with some areas of adipose tissue (**Figure 4F**). Histology staining reflected this increased stroma composition (**Figure 4G**). A single peptide domain demonstrated significant alterations when compared by ancestry with the heterogeneously dense category peptide P1142 (**Figure 4H**). To summarize, BI-RADS density showed distinguishing molecular signatures from stroma regions; subsequent genetic ancestry-associated analysis supports fundamental molecular differences in stroma from the normal breast microenvironment.

**Figure 4.**
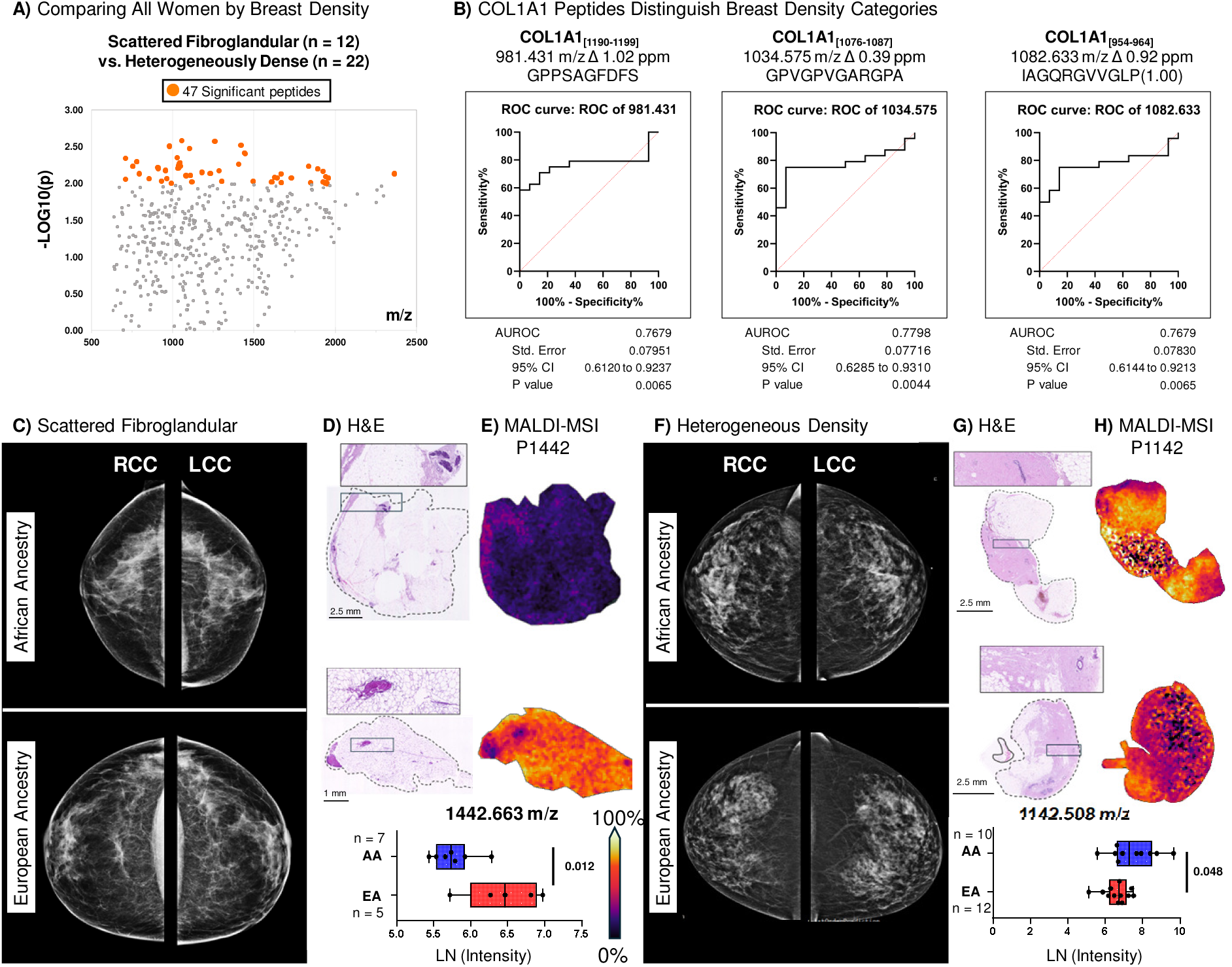
The normal breast collagen proteome distinguishes scattered fibroglandular and heterogeneously dense breast density categories. A) Stromal proteome comparisons reveal 47 significantly altered peptides between scattered fibroglandular (n = 12) and heterogeneously dense (n = 22) breast density categories. B) Specific identified collagen α-1(I) peptide domains differentiate between the two categories (AUROC > 0.75). C) Right and left craniocaudal (RCC/LCC) views of example mammograms from women of African or European ancestry with scattered fibroglandular breast density. D) H&E-stained tissue sections donated from the women whose mammograms are shown in (C). E) Spatial distribution and intensity of P1442 within scattered fibroglandular breast tissues alters by genetic ancestry. F) RCC and LCC views of mammograms from women of African or European ancestry with heterogeneously dense breasts. G) H&E-stained tissue sections donated from women whose mammograms are shown in (F). H) Peptide P1142 alters within heterogeneously dense breasts by genetic ancestry.

### The collagen proteome in normal breast is altered in association with body mass index

Obesity is a well-known risk factor for the development of breast cancer in both Black women and White women in the United States(76, 77). Black women are affected by obesity at higher prevalence compared to White women(78), yet the effects of obesity on the stromal-specific healthy breast proteome are relatively understudied. Overall stromal-specific profiles revealed complex proteomic signatures from healthy, overweight, and obese stroma with increased intensities in obese donors (**Figure 5A**). The top 50 differentially expressed peptides show variation in peptide domain profiles across conventional body mass index categories (**Figure 5B, Supplemental Table 4**). Clustering of samples based on BMI categories (healthy, overweight, and obese) revealed distinct clusters by collagen profiles within each BMI group (**Figure 5C**). The majority of ECM peptide domains extracted from stroma regions were significantly upregulated in obese donors compared to overweight donors, as anticipated based on spectral evaluation (**Figure 5D**). Identified domains within collagen α-1(I) (COL1A1) containing hydroxyproline residues (modified proline) show decreased intensities within all BMI classes compared to unmodified (**Figure 5E**). However, while healthy BMI showed a small significant change in HYP modified COL1A1 peptides, overweight and obese reported a more significant decrease in HYP modification. Single COL1A1 peptides were found to distinguish between categories of overweight and obese (AUROC > 0.8, p-value < 0.005, **Figure 5 F, G**). Triple helical regions are generally recognized to be in a 2:1 ratio COL1A1 and COL1A2; here COL1A2 did not show a significant change in unmodified versus modified status within BMI categories (**Figure 5H**). However, COL1A2 showed a decrease in both unmodified and modified peptides in overweight BMI compared to either healthy BMI or obese BMI. Peptide domain differences based on genetic ancestry were seen within overweight and obese BMI categories (OW: 3 peptides, OB: 26 peptides). Interestingly, all 26 peptides identified within the obese class are increased in EA compared to AA (**Supplemental Figure 8**). Example distribution patterns in stroma are shown for ancestry-based comparisons of overweight women (P1321, **Figure 5I**) and obese women (P1188, **Figure 5J**). Obesity significantly impacts the breast collagen proteome, leading to increased peptide domain expression in obese vs. overweight donors, with potential genetic ancestry-based differences among overweight and obese women.

**Figure 5.**
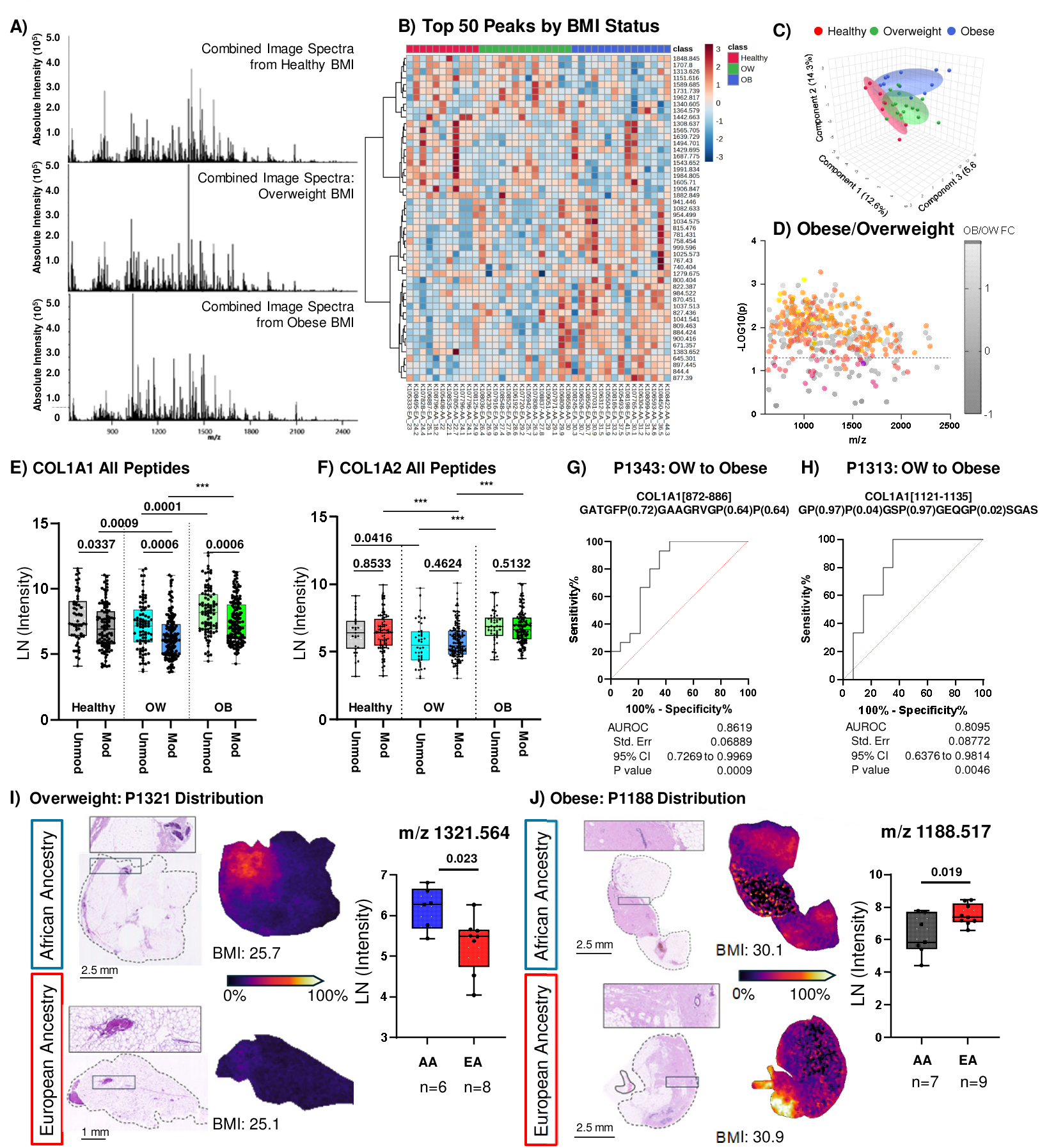
Normal breast collagen proteome alters with BMI. A) Proteomic imaging spectra of the normal breast shows complexity of the combined collagen proteome within healthy, overweight, or obese BMI. B) Top 50 peptides distribute based on BMI status. C) Principal components analysis shows clear separation between healthy, overweight and obese BMI. Shading represents 95% prediction ellipse. D) Comparison of Obese to Overweight shows that the majority of detected peptide domains are significantly increased in obese donors compared to overweight. E) When comparing all measurements of collagen domain peptides, collagen α-1(I) chain shows an increase in unmodified peptides in obese compared to overweight donors (Mann-Whitney Test, p-values). F) P1343 and G) P1313 showing high sensitivity and specificity distinguishing overweight and obese. H) When comparing all collagen domain peptides from collagen α-2(I) chain, significant differences comparing either unmodified or modified between different categories, but not within BMI categories. I) Peptide P1321 alters within the overweight category by genetic ancestry. J) Peptide P1188 alters within the obese category by genetic ancestry.

### Multivariate analysis links the collagen proteome to BMI status

The molecular relationship of the normal breast collagen proteome and links to cumulative clinical characteristics were modeled as a collective signature. Using collagen peptide measurements, multivariate linear regression of the collagen proteome intensity measures extracted from stroma regions were fit to clinical characteristics, breast density, income, and fiber measurements and adjusted for covariates of ancestry. Principal components analysis was used to reduce the multiple peptide domain measurements (n=478 peptides) to 6 principal components where principal component 1 accounted for 91.2% of the variability. Collagen fiber measurements of width and length were used to validate the utility of this approach (**Figure 6 A, B**). These data demonstrate that independent of ancestry, collagen fiber width has a strong positive association with the normal breast microenvironment, corresponding to findings in fiber measurements by SHG. When clinical characteristics including income were evaluated, donors with African ancestry showed a positive association with both BMI overweight and BMI obesity (**Figure 6C**), whereas European ancestry showed a strong positive association with BMI Obese but not BMI overweight (**Figure 6D**). Altogether, cumulative data supports that BMI has a significant impact on normal breast health, with strong positive associations to changes in the collagen protein microenvironment.

**Figure 6.**
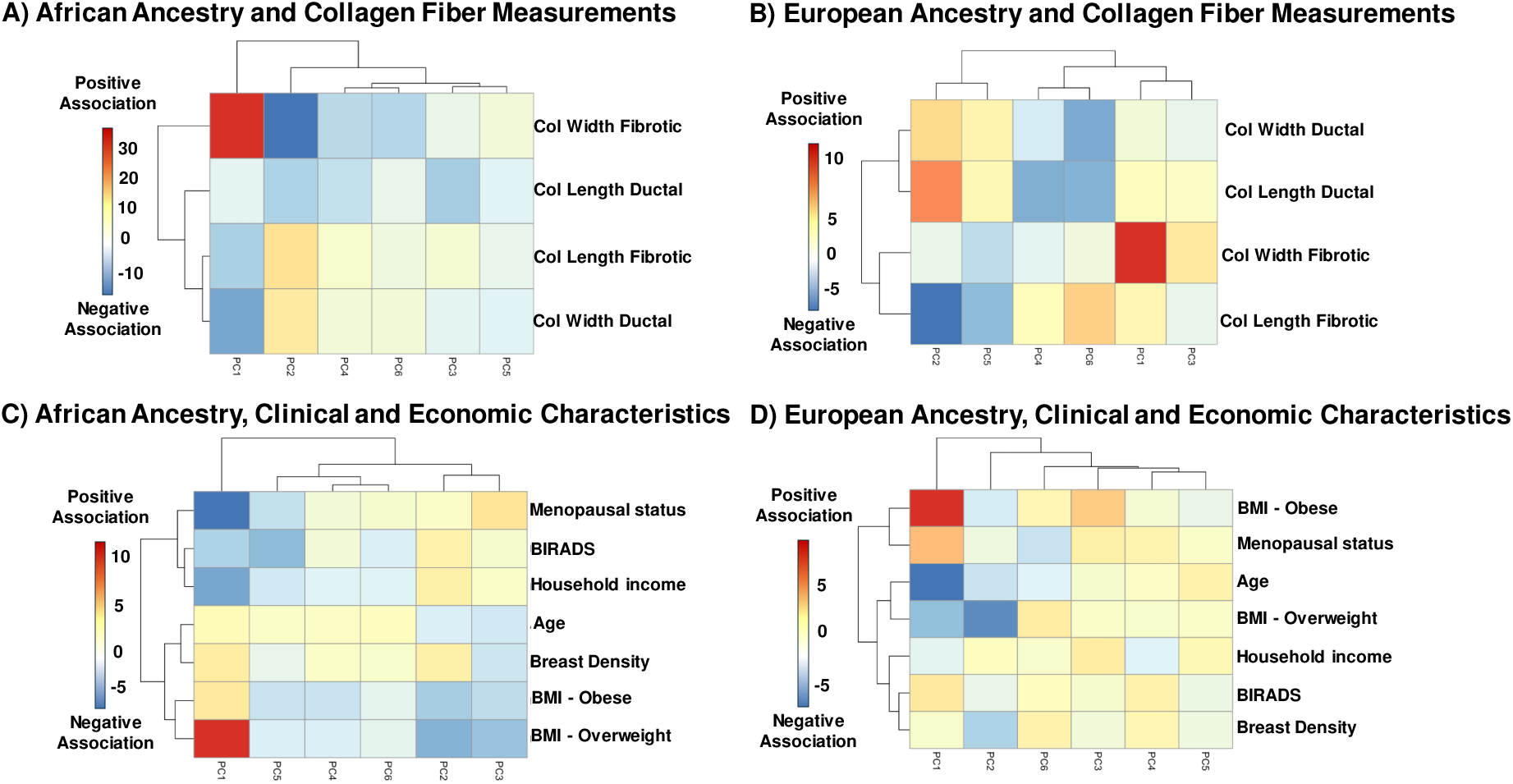
Multivariate analysis relationships between normal breast collagen proteome, economic, and clinical characteristics. Principal component analysis was used to reduce complex collagen peptide domain signatures to 6 principal components and evaluated by economic and clinical characteristics per genetic ancestry. A,B) Validation of the approach was demonstrated using collagen proteome PC compared to collagen fiber measurements. This data reported strong positive associations with collagen width, also shown in the singular second harmonic generation measurements. C) Per African ancestry (AA), normal breast collagen proteome shows a strong positive association with overweight BMI. D) Normal breast collagen proteome by European ancestry shows a lower (compared to AA) positive association obese BMI.

## Discussion

It is a dichotomy that throughout the course of clinical care, breast stroma is used to evaluate breast cancer risk, yet we know very little about the molecular composition of the breast stroma microenvironment. Previous studies have provided excellent single cell profiles from within the normal breast microenvironment(12, 79), but generally focus on transcript level; information on proteomic output that forms the working stroma microenvironment is more limited. This study makes new inroads into mechanisms of the normal breast microenvironment that include changes within the collagen stroma proteome. To this end, donor tissue from the Susan G. Komen tissue bank (n=40) classified as low risk for cancer based on Tyrer-Cuzick score and moderate risk based on Gail score were used to investigate protein cell markers, collagen fiber measurements and extracellular proteomic distribution relative to clinical characteristics. This study reports a broad association with BMI status and alterations of the normal breast proteomic microenvironment. Cell markers of breast cancer prognosis were detected, correlated with stroma markers of breast cancer risk and displayed BMI-associated alterations. Collagen fiber measurements further demonstrated a connection to BMI status with potential influences based on genetic ancestry obesity. Signatures derived from the stroma distinguished clinical measurements of breast density and further demonstrated modulation due to BMI status. Linear regression analysis including lifestyle factors and risk scoring further emphasized that BMI plays a significant role in normal breast stroma microenvironment. The normal breast stroma proteome is a complex microenvironment particularly sensitive to lifestyle factors that increase breast cancer risk.

A major finding in this study was that obesity, known to be a major clinical risk factor of breast cancer, modulates the normal breast stroma proteomic microenvironment. Collagen signatures explored by antibody staining, fiber measurements, proteomic imaging and multivariate analyses all pointed to normal breast collagen altered by BMI status. The normal breast stroma proteome showed spatial changes within stroma, demonstrating a dynamic regulation of collagen domains across breast histology. Cumulatively, overweight or obese BMI showed peptides from COL1A1 with significant decreases in modification compared to healthy BMI. Canonically, collagen α-1(I) chain forms a triple helical of 2:1 with collagen α-2(I) chain; hydroxylation of proline is a basis for chain-to-chain recognition and masks or exposes cell interactive sites within collagen domains. Therefore, changes in post-translational modification status within COL1A1 suggest differential access to domains that modulate cell-collagen interactions. Indeed, previous work has pointed to obesity-related alterations of immune evasion(39–41) (34, 42), and signaling gradients(43, 44) to increase emergent tumor, invasion and metastasis (39, 45); global proteomic studies have shown that specific sites increase in invasive breast carcinoma relative to benign breast diseases(25). It is thus likely that collagen stroma proteomic modifications within the tissue microenvironment play a fundamental role in BMI change related to immune trafficking, recruitment, and localization as well as gradient control of cytokines. This was supported by data showing that collagen stroma proteomic signatures were highly complex and that specific post-translationally modified collagen peptides could distinguish between BMI categories of overweight and obese.

Obesity status further associated with critical cellular changes and collagen fiber measurements. In this study, CD44 decreased with obese BMI status and frequently co-localized with the CD68 macrophage marker but did not show expression level correlation with CD68. CD44 is a cell surface glycoprotein binds hyaluronan, another component of the extracellular matrix with roles in regulation of inflammation and cancer(80). In the normal breast, CD44 is predominantly expressed on myoepithelial cells that surround and support luminal ductal cells. CD44 plays a critical role in normal mammary gland development, with changes in expression seen throughout reproduction and lactation, reflecting its association with mammary stem-like properties(81). Previous work has shown that CD44 can both promote and inhibit breast tumor invasion through independent structural binding sites that are dependent on localized extracellular composition(82, 83). Since the CD44+/CD24-phenotype has been identified as a marker of cancer stemness and shows increased expression in women of African genetic ancestries related to breast cancer risk(84). it is likely that other CD44 subtypes may exist that are modulated by BMI status. There are many known isoforms of CD44 due to alternative splicing. The standard form (CD44s) has been positively correlated with cancer stem cell signatures whereas variant forms (CD44v) negatively associate with these signatures(85). Additionally, obesity-related inflammation has been linked to elevated CD44 expression in white adipose tissue, particularly in the liver(86). Together with the current study, these findings suggest that obesity may modulate the expression of specific CD44 variants in normal breast tissue, with potential implications for cancer risk and progression.

Collagen fiber measurements further highlighted potential obesity changes within the stroma. Interestingly, when genetic ancestry was considered with BMI, overweight BMI showed increases in collagen fiber width for women of European genetic ancestry; in obese categories, this switched to show an increase in collagen fiber width for women of African genetic ancestry. Investigations including collagen fiber width measurements on normal breast are limited and few to none report on ancestry. Studies of ductal carcinoma in situ show that increases in collagen fiber width associate with decreased risk of recurrence(87). Breast fiber width was increased in normal or benign breast relative to breast cancer(88). Our own studies shown ancestry-dependent fiber width decreases in tumor compared to normal adjacent breast tissue(89). Contradictions could arise from the small site localization of where the SHG collagen fiber width data are collected relative to the tissue pathology. This is supported by the current study showing difference in ductal versus fibrotic regions and past reports that thicker collagen fibers localized to the invasive front of the tumor link to poor outcomes(18), pointing to the need to understand fiber regulation across the entire tissue microenvironment. In this study, when all proteomic data were assessed including ancestry, collagen width in fibrotic regions had a strong positive association with both genetic ancestries. Cumulative clinical and economic data showed a strong positive association of the collagen proteome with BMI status overweight in African genetic ancestry; in European genetic ancestry obese BMI showed the strongest positive association compared to clinical and economic measurements. The collective data imply that genetic ancestry along with BMI status has a significant role in homeostasis and control of the collagen stroma within the normal breast microenvironment.

This study suggests that clinical categories of breast density by BI-RADS may be predicted based on molecular composition of the breast stroma. Multiple collagen domains derived as peptides specific to breast stroma regions appeared altered based on density category of “scattered fibroglandular”, sparce glandular features within mostly fatty tissue versus “heterogeneously dense”, tissue with higher proportions of fibrous tissue. All collagen domain peptides derived from stroma showed an increase in intensity in samples categorized as heterogeneously dense. However, although relative expression level changes were small, collagen domain peptides showed the ability to distinguish between SF and HD with high specificity and sensitivity. Thus, it is possible that small molecular changes in breast transitioning to different breast density categories are detectable with predictive value and could be further leveraged to add to assessments of breast health. In these cases of healthy normal breast, it is still unknown whether or not the donors developed breast cancer. Larger studies are needed to assess stroma composition related to changes in breast density and early cancer emergence. Previous work has shown by collagen fiber measurements shows the systematic emergence of tissue associated collagen signatures in breast cancer progression(18). The current data supports that the molecular composition of collagen in the normal breast is also likely to be systematic and predictable.

This study had limitations. The samples were selected from the Susan G. Komen tissue bank of normal breast based on similar risk categories and presence of stroma in the biopsy, resulting in a smaller cohort. Early collection procedure did not document sampling location of the donated tissue, and it is likely that normal breast shows heterogeneity across the breast. Therefore, it is unknown how the collected sample represents the entire breast. While donors are being followed for development of breast cancer, we do not yet know the connection between breast density that develops cancer versus breast density that does not develop cancer. Finally, certain genetic ancestries have high breast density and lower incidence of breast cancer(90). Studies are needed that are cross-institutional and include information on genetic ancestries with BMI status for a complete understanding of both cellular and extracellular changes in the normal breast microenvironment.

## Conclusions

An emergent story is that the normal breast microenvironment is dynamically modulated by cellular composition, microbiome changes, ancestry, menopause, and age (12, 13, 15, 79, 91). These valuable studies are yielding unexpected insights into cell populations that are harbored within the normal breast and increase breast cancer risk. Breast stroma is monitored throughout the course of a lifetime for assessment of cancer risk, yet lacks definition of the molecular underpinnings that lead to risk. Collagen breast stroma has the capacity to control cell function through cell-collagen binding with comprehensive implications particularly to emergent tumor cells and to immune recruitment, localization, and response. One gap in our knowledge is understanding cellular origins and timelines that result in specific stroma collagen composition. Our current knowledge of breast stroma is that fibroblast cell types express the largest percent of collagen protein, yet fibroblasts are not always found in regions of high collagen from normal breast or breast cancer that show immune cell streaming or cellular deserts. Therefore, collagen histology leading to current breast health status and immune capabilities are produced at a much earlier timepoint. Implications arise that suggest that 1) collagen stroma microenvironment is the frontline of breast health outcomes, either due to genetics or socioeconomic factors, and much earlier detection of a cancer permissive environment should be possible based on stroma composition; 2) progressive changes in breast density by deposition and remodeling should lead to stroma shed into circulation for noninvasive tests reporting higher breast cancer risk. Clearly more studies are required that include comprehensive assessment of both cellular and collagen stroma microenvironments in progression up to and including breast cancer.

## Supporting information

All supplemental

## Declarations

## Ethics approval and consent to participate

Normal breast tissue sections were obtained through the Susan G. Komen tissue bank with appropriate IRB protocols.

## Consent for publication

Not applicable

## Availability of data and materials

The datasets supporting the conclusions of this article are included within the article (and its supplemental files). The full LC-MS/MS dataset analyzed in the current study is available from the corresponding author on reasonable request.

## Authors’ contributions

The work reported in this paper has been performed by the authors. JBD performed MALDI-MSI and MALDI-IHC experiments and used QuPath for data analysis. HBT and JKM performed proteomic tissue preparation and LC-MS/MS workflows, respectively. HJS performed second harmonic generation microscopy data acquisition for JBD to analyze. YP developed novel models for multivariate analysis. Analysis and intellectual contributions are attributed to JBD and PMA along with technical and intellectual support from the other coauthors.

## Acknowledgements

Tissues used in this manuscript were selected from the Susan G Komen Tissue Bank; we thank the courageous donors to this bank. We thank Jill Henry for her invaluable and ongoing assistance with the SGK samples. We appreciate the work done by Jessica Lord in organizing the tables.

## References

1. Boyd NF GH, Martin LJ, Sun L, Stone J, Fishell E, Jong RA, Hislop G, Chiarelli A, Minkin S, Yaffe MJ. Mammographic Density and the Risk and Detection of Breast Cancer. N Engl J Med. 2007;356;3.

2. Li T SL, Miller N, Nicklee T, Woo J, Hulse-Smith L, Tsao MS, Khokha R, Martin L, Boyd N. The Association of Measured Breast Tissue Characteristics with Mammographic Density and Other Risk Factors for Breast Cancer. Cancer Epidemiol Biomarkers Prev. 2005;14(2).

3. Alowami S, Troup S, Al-Haddad S, Kirkpatrick I, Watson PH. Mammographic density is related to stroma and stromal proteoglycan expression. Breast Cancer Res. 2003;5(5):R129–35.

4. Bodelon C, Mullooly M, Pfeiffer RM, Fan S, Abubakar M, Lenz P, et al. Mammary collagen architecture and its association with mammographic density and lesion severity among women undergoing image-guided breast biopsy. Breast Cancer Res. 2021;23(1):105.

5. Provenzano PP, Inman DR, Eliceiri KW, Knittel JG, Yan L, Rueden CT, et al. Collagen density promotes mammary tumor initiation and progression. BMC Med. 2008;6:11.

6. Boyd NF, Martin LJ, Yaffe MJ, Minkin S. Mammographic density and breast cancer risk: current understanding and future prospects. Breast cancer research. 2011;13(6):223. PMCID: PMC3326547.

7. Control UCfD. Adult BMI Categories 2024 [

8. Watanabe K, Wilmanski T, Diener C, Earls JC, Zimmer A, Lincoln B, et al. Multiomic signatures of body mass index identify heterogeneous health phenotypes and responses to a lifestyle intervention. Nature Medicine. 2023;29(4):996–1008. PMC10115644.

9. Martinez J, Smith PC. The dynamic interaction between extracellular matrix remodeling and breast tumor progression. Cells. 2021;10(5):1046.PMC8145942.

10. Muschler J, Streuli CH. Cell–matrix interactions in mammary gland development and breast cancer. Cold Spring Harbor perspectives in biology. 2010;2(10):a003202. PMC2944360.

11. Chen R, Zhang R, Ke F, Guo X, Zeng F, Liu Q. Mechanisms of breast cancer metastasis: the role of extracellular matrix. Molecular and Cellular Biochemistry. 2024:1–26. PMID:39652293.

12. Bhat-Nakshatri P, Gao H, Sheng L, McGuire PC, Xuei X, Wan J, et al. A single-cell atlas of the healthy breast tissues reveals clinically relevant clusters of breast epithelial cells. Cell Reports Medicine. 2021;2(3):100219.PMID: 33763657.

13. Kumar T, Nee K, Wei R, He S, Nguyen QH, Bai S, et al. A spatially resolved single-cell genomic atlas of the adult human breast. Nature. 2023;620(7972):181-91. PMID: 37380767.

14. Reed AD, Pensa S, Steif A, Stenning J, Kunz DJ, Porter LJ, et al. A single-cell atlas enables mapping of homeostatic cellular shifts in the adult human breast. Nature Genetics. 2024;56(4):652–62. PMC11018528.

15. Hieken TJ, Chen J, Chen B, Johnson S, Hoskin TL, Degnim AC, et al. The breast tissue microbiome, stroma, immune cells and breast cancer. Neoplasia. 2022;27:100786. PMC8971327.

16. Ricard-Blum S. The collagen family. Cold Spring Harbor Perspectives in Biology. 2011;3(1):a004978.PMID: 21421911.

17. Provenzano PP, Eliceiri KW, Campbell JM, Inman DR, White JG, Keely PJ. Collagen reorganization at the tumor-stromal interface facilitates local invasion. BMC Med. 2006;4(1):38.

18. Conklin MW, Eickhoff JC, Riching KM, Pehlke CA, Eliceiri KW, Provenzano PP, et al. Aligned collagen is a prognostic signature for survival in human breast carcinoma. The American Journal of Pathology. 2011;178(3):1221–32. PMCID: PMC3070581.

19. Brett EA, Sauter MA, Machens H-G, Duscher D. Tumor-associated collagen signatures: pushing tumor boundaries. Cancer & Metabolism. 2020;8:1–5. PMC7331261.

20. Guo Q, Sun D, Barrett AS, Jindal S, Pennock ND, Conklin MW, et al. Mammary collagen is under reproductive control with implications for breast cancer. Matrix Biology. 2022;105:104–26. PMID: 34839002.

21. San Antonio JD, Jacenko O, Fertala A, Orgel JPRO. Collagen structure-function mapping informs applications for regenerative medicine. Bioengineering. 2020;8(1):3.PMCID: PMC7824244.

22. Sweeney SM, Orgel JP, Fertala A, McAuliffe JD, Turner KR, Di Lullo GA, et al. Candidate cell and matrix interaction domains on the collagen fibril, the predominant protein of vertebrates. J Biol Chem. 2008;283(30):21187–97.PMCID: PMC2475701.

23. Shoulders MD, Raines RT. Collagen structure and stability. Annual Review of Biochemistry. 2009;78:929–58. PMID: 19344236.

24. Gjaltema RAF, Bank RA. Molecular insights into prolyl and lysyl hydroxylation of fibrillar collagens in health and disease. Critical reviews in biochemistry and molecular biology. 2017;52(1):74–95.

25. Montgomery H, Rustogi N, Hadjisavvas A, Tanaka K, Kyriacou K, Sutton CW. Proteomic profiling of breast tissue collagens and site-specific characterization of hydroxyproline residues of collagen alpha-1-(I). Journal of Proteome Research. 2012;11(12):5890–902.

26. Gorres KL, Raines RT. Prolyl 4-hydroxylase. Critical Reviews in Biochemistry and Molecular Biology. 2010;45(2):106–24. PMCID: PMC2841224.

27. Salo AM, Myllyharju J. Prolyl and lysyl hydroxylases in collagen synthesis. Experimental Dermatology. 2021;30(1):38–49.

28. Xiong G, Stewart RL, Chen J, Gao T, Scott TL, Samayoa LM, et al. Collagen prolyl 4-hydroxylase 1 is essential for HIF-1α stabilization and TNBC chemoresistance. Nature Communications. 2018;9(1):1–16. PMCID: PMC6203834.

29. Xiong G, Deng L, Zhu J, Rychahou PG, Xu R. Prolyl-4-hydroxylase α subunit 2 promotes breast cancer progression and metastasis by regulating collagen deposition. BMC Cancer. 2014;14(1):1.PMCID: PMC3880410.

30. Gilkes DM, Chaturvedi P, Bajpai S, Wong CC, Wei H, Pitcairn S, et al. Collagen prolyl hydroxylases are essential for breast cancer metastasis. Cancer Res. 2013;73(11):3285–96.

31. Vasta JD, Raines RT, James Vasta MD, Raines RT. Collagen Prolyl 4-Hydroxylase as a Therapeutic Target. Journal of Medicinal Chemistry. 2018.

32. Shi R, Gao S, Zhang J, Xu J, Graham LM, Yang X, Li C. Collagen prolyl 4-hydroxylases modify tumor progression. Acta Biochimica et Biophysica. 2021;53(7):805–14.PMID: 34009234.

33. Hillers-Ziemer LE, Kuziel G, Williams AE, Moore BN, Arendt LM. Breast cancer microenvironment and obesity: challenges for therapy. Cancer and Metastasis Reviews. 2022;41(3):627–47. PMC9470689.

34. Savva C, Copson E, Johnson PWM, Cutress RI, Beers SA. Obesity is associated with immunometabolic changes in adipose tissue that may drive treatment resistance in breast cancer: immune-metabolic reprogramming and novel therapeutic strategies. Cancers. 2023;15(9):2440. PMC10177091.

35. Lauby-Secretan B, Scoccianti C, Loomis D, Grosse Y, Bianchini F, Straif K. Body fatness and cancer—viewpoint of the IARC Working Group. New England Journal of Medicine. 2016;375(8):794–8. PMC6754861.

36. Recalde M, Pistillo A, Davila-Batista V, Leitzmann M, Romieu I, Viallon V, et al. Longitudinal body mass index and cancer risk: a cohort study of 2.6 million Catalan adults. Nature Communications. 2023;14(1):3816. PMC10313757.

37. Sudan SK, Sharma A, Vikramdeo KS, Davis W, Deshmukh SK, Poosarla T, et al. Obesity and early-onset breast cancer and specific molecular subtype diagnosis in black and white women: NIMHD social epigenomics program. JAMA Network Open. 2024;7(7):e2421846-e. PMC11287389.

38. Nyrop KA, Damone EM, Deal AM, Carey LA, Lorentsen M, Shachar SS, et al. Obesity, comorbidities, and treatment selection in Black and White women with early breast cancer. Cancer. 2021;127(6):922–30. PMID: 33284988.

39. Lee CM, Fang S. Fat Biology in Triple-Negative Breast Cancer: Immune Regulation, Fibrosis, and Senescence. J Obes Metab Syndr. 2023;32(4):312–21. PMC10786212.

40. Floris G, Richard F, Hamy A-S, Jongen L, Wildiers H, Ardui J, et al. Body mass index and tumor-infiltrating lymphocytes in triple-negative breast cancer. JNCI: Journal of the National Cancer Institute. 2021;113(2):146–53. PMC7850533.

41. Pingili AK, Chaib M, Sipe LM, Miller EJ, Teng B, Sharma R, et al. Immune checkpoint blockade reprograms systemic immune landscape and tumor microenvironment in obesity-associated breast cancer. Cell Reports. 2021;35(12):109285. PMC8574993.

42. Russo S, Kwiatkowski M, Govorukhina N, Bischoff R, Melgert BN. Meta-inflammation and metabolic reprogramming of macrophages in diabetes and obesity: the importance of metabolites. Frontiers in immunology. 2021;12:746151. PMC8602812.

43. Zhang K, Chen L, Zheng H, Zeng Y. Cytokines secreted from adipose tissues mediate tumor proliferation and metastasis in triple negative breast cancer. BMC cancer. 2022;22(1):886. PMC9375239.

44. Lee K, Kruper L, Dieli-Conwright CM, Mortimer J. The Impact of Obesity on Breast Cancer Diagnosis and Treatment. Current Oncology Reports. 2019;21(5):41. PMC6437123.

45. Bousquenaud M, Fico F, Solinas G, Rüegg C, Santamaria-Martínez A. Obesity promotes the expansion of metastasis-initiating cells in breast cancer. Breast Cancer Research. 2018;20:1–11. PMC6123990.

46. Bhat-Nakshatri P, Marino N, Gao H, Liu Y, Storniolo AM, Nakshatri H. Acquisition, processing, and single-cell analysis of normal human breast tissues from a biobank. STAR Protoc. 2022;3(1):101047.

47. Nievergelt CM, Maihofer, A.X., Shekhtman, T., Libiger, O., Wang, X., Kidd, Kenneth K., and Kidd, Judith R. Inference of human continental origin and admixture proportions using a highly discriminative ancestry informative 41-SNP panel. Investigative Genetics. 2013;4(13).

48. Gail MH, Costantino JP, Pee D, Bondy M, Newman L, Selvan M, et al. Projecting individualized absolute invasive breast cancer risk in African American women. J Natl Cancer Inst. 2007;99(23):1782–92.

49. Gail MH BL, Byar DP, Corle DK, Green SB, Schairer C, Mulvihill JJ. Projecting individualized probabilities of developing breast cancer for white females who are being examined annually. J Natl Cancer Inst 1989;81.

50. Tyrer J, Duffy SW, Cuzick J. A breast cancer prediction model incorporating familial and personal risk factors. Stat Med. 2004;23(7):1111–30.

51. Yagnik G, Liu Z, Rothschild KJ, Lim MJ. Highly multiplexed immunohistochemical MALDI-MS imaging of biomarkers in tissues. Journal of the American Society for Mass Spectrometry. 2021;32(4):977–88.

52. Schneider CA, Rasband, Wayne S., and Eliceiri, Kevin W. NIH Image to ImageJ: 25 years of Image Analysis. Nat Methods. 2012;9(7):671–5.

53. Bredfeldt JS, Liu Y, Conklin MW, Keely PJ, Mackie TR, Eliceiri KW. Automated quantification of aligned collagen for human breast carcinoma prognosis. J Pathol Inform. 2014;5(1):28.

54. Bredfeldt JS, Liu Y, Pehlke CA, Conklin MW, Szulczewski JM, Inman DR, et al. Computational segmentation of collagen fibers from second-harmonic generation images of breast cancer. J Biomed Opt. 2014;19(1):16007.

55. Liu Y, Keikhosravi A, Mehta GS, Drifka CR, Eliceiri KW. Methods for Quantifying Fibrillar Collagen Alignment. Methods Mol Biol. 2017;1627:429–51.

56. Angel PM, Comte-Walters S, Ball LE, Talbot K, Mehta A, Brockbank KGM, Drake RR. Mapping Extracellular Matrix Proteins in Formalin-Fixed, Paraffin-Embedded Tissues by MALDI Imaging Mass Spectrometry. J Proteome Res. 2018;17(1):635–46.

57. Powers TW, Neely BA, Shao Y, Tang H, Troyer DA, Mehta AS, et al. MALDI imaging mass spectrometry profiling of N-glycans in formalin-fixed paraffin embedded clinical tissue blocks and tissue microarrays. PLoS One. 2014;9(9):e106255, pp1-11. PMID: 25184632.

58. Clift CL, Drake RR, Mehta A, Angel PM. Multiplexed imaging mass spectrometry of the extracellular matrix using serial enzyme digests from formalin-fixed paraffin-embedded tissue sections. Anal Bioanal Chem. 2021;413(10):2709–19.

59. Bankhead P, Loughrey MB, Fernández JA, Dombrowski Y, McArt DG, Dunne PD, et al. QuPath: Open source software for digital pathology image analysis. Scientific Reports. 2017;7(1):1–7.PMCID: PMC5715110.

60. Xia J, Wishart DS. Web-based inference of biological patterns, functions and pathways from metabolomic data using MetaboAnalyst. Nature Protocols. 2011;6(6):743–60.

61. da Veiga Leprevost F, Haynes, S.E., Avtonomov, D.M. et al. Philosopher: a versatile toolkit for shotgun proteomics data analysis. Nat Methods. 2020;17.

62. Venne K BE, Eng K, Thibault P. Improvement in Peptide Detection for Proteomics Analyses Using NanoLC-MS and High-Field Asymmetry Waveform Ion Mobility Mass Spectrometry. Anal Chem. 2005;77.

63. Biniossek ML, Schilling O. Enhanced identification of peptides lacking basic residues by LC-ESI-MS/MS analysis of singly charged peptides. Proteomics. 2012;12(9):1303–9.

64. Geiszler DJ, Kong AT, Avtonomov DM, Yu F, Leprevost FDV, Nesvizhskii AI. PTM-Shepherd: Analysis and Summarization of Post-Translational and Chemical Modifications From Open Search Results. Mol Cell Proteomics. 2021;20:100018.

65. Kong AT, Leprevost FV, Avtonomov DM, Mellacheruvu D, Nesvizhskii AI. MSFragger: ultrafast and comprehensive peptide identification in mass spectrometry-based proteomics. Nat Methods. 2017;14(5):513–20.

66. Li K, Vaudel M, Zhang B, Ren Y, Wen B. PDV: an integrative proteomics data viewer. Bioinformatics. 2019;35(7):1249–51.

67. Shteynberg DD, Deutsch EW, Campbell DS, Hoopmann MR, Kusebauch U, Lee D, et al. PTMProphet: Fast and Accurate Mass Modification Localization for the Trans-Proteomic Pipeline. J Proteome Res. 2019;18(12):4262–72.

68. Teo GC, Polasky DA, Yu F, Nesvizhskii AI. Fast Deisotoping Algorithm and Its Implementation in the MSFragger Search Engine. J Proteome Res. 2021;20(1):498–505.

69. Yu F, Haynes SE, Nesvizhskii AI. IonQuant Enables Accurate and Sensitive Label-Free Quantification With FDR-Controlled Match-Between-Runs. Mol Cell Proteomics. 2021;20:100077.

70. Yu F, Haynes SE, Teo GC, Avtonomov DM, Polasky DA, Nesvizhskii AI. Fast Quantitative Analysis of timsTOF PASEF Data with MSFragger and IonQuant. Mol Cell Proteomics. 2020;19(9):1575–85.

71. Yu F, Teo GC, Kong AT, Haynes SE, Avtonomov DM, Geiszler DJ, Nesvizhskii AI. Identification of modified peptides using localization-aware open search. Nat Commun. 2020;11(1):4065.

72. Jansson M, Lindberg J, Rask G, Svensson J, Billing O, Nazemroaya A, et al. Stromal type I collagen in breast cancer: correlation to prognostic biomarkers and prediction of chemotherapy response. Clinical Breast Cancer. 2024.

73. Sweeney SM, Orgel JP, Fertala A, McAuliffe JD, Turner KR, Di Lullo GA, et al. Candidate cell and matrix interaction domains on the collagen fibril, the predominant protein of vertebrates. J Biol Chem. 2008;283(30):21187–97.

74. Pettersson A, Graff RE, Ursin G, Santos Silva ID, McCormack V, Baglietto L, et al. Mammographic density phenotypes and risk of breast cancer: a meta-analysis. J Natl Cancer Inst. 2014;106(5).

75. Nazari SS, Mukherjee P. An overview of mammographic density and its association with breast cancer. Breast Cancer. 2018;25(3):259–67.

76. Pierobon M, Frankenfeld CL. Obesity as a risk factor for triple-negative breast cancers: a systematic review and meta-analysis. Breast Cancer Res Treat. 2013;137(1):307–14.

77. Sun H, Zou J, Chen L, Zu X, Wen G, Zhong J. Triple-negative breast cancer and its association with obesity. Mol Clin Oncol. 2017;7(6):935–42.

78. Hales CM CM, Fryar CD, Ogden CL. Prevalence of obesity and severe obesity among adults: United States, 2017–2018.: Centers for Disease Control and Prevention; 2020 [Available from: www.cdc.gov/nchs/products/databriefs/db360.htm

79. Lin Y, Wang J, Wang K, Bai S, Thennavan A, Wei R, et al. Normal breast tissues harbour rare populations of aneuploid epithelial cells. Nature. 2024:1–8. PMID39567687.

80. Misra S, Hascall VC, Markwald RR, Ghatak S. Interactions between Hyaluronan and Its Receptors (CD44, RHAMM) Regulate the Activities of Inflammation and Cancer. Front Immunol. 2015;6:201.

81. Louderbough JM, Brown JA, Nagle RB, Schroeder JA. CD44 Promotes Epithelial Mammary Gland Development and Exhibits Altered Localization during Cancer Progression. Genes Cancer. 2011;2(8):771–81.

82. Louderbough JMV, Schroeder JA. Understanding the dual nature of CD44 in breast cancer progression. Molecular Cancer Research. 2011;9(12):1573–86. PMID: 21970856.

83. Cirillo N. The hyaluronan/CD44 axis: A double-edged sword in cancer. International Journal of Molecular Sciences. 2023;24(21):15812. PMC10649834.

84. Nakshatri H, Anjanappa M, Bhat-Nakshatri P. Ethnicity-dependent and-independent heterogeneity in healthy normal breast hierarchy impacts tumor characterization. Scientific Reports. 2015;5:13526. PMCID: PMC4550930.

85. Zhang H, Brown RL, Wei Y, Zhao P, Liu S, Liu X, et al. CD44 splice isoform switching determines breast cancer stem cell state. Genes Dev. 2019;33(3-4):166–79.

86. Kang HS, Liao G, DeGraff LM, Gerrish K, Bortner CD, Garantziotis S, Jetten AM. CD44 plays a critical role in regulating diet-induced adipose inflammation, hepatic steatosis, and insulin resistance. PLoS One. 2013;8(3):e58417.

87. Sprague BL, Vacek PM, Mulrow SE, Evans MF, Trentham-Dietz A, Herschorn SD, et al. Collagen Organization in Relation to Ductal Carcinoma In Situ Pathology and OutcomesCollagen Organization in Relation to DCIS Outcomes. Cancer Epidemiology, Biomarkers & Prevention. 2021;30(1):80–8. PMCID: PMC7855820.

88. Bodelon C, Mullooly M, Pfeiffer RM, Fan S, Abubakar M, Lenz P, et al. Mammary collagen architecture and its association with mammographic density and lesion severity among women undergoing image-guided breast biopsy. Breast Cancer Research. 2021;23:1–14.

89. Taylor H, Spruill L, Jensen-Smith H, Rujchanarong D, Hulahan T, Ivey A, et al. Spatial Localization of Collagen Hydroxylated Proline Site Variation as an Ancestral Trait in the Breast Cancer Microenvironment. Matrix Biology. 2025;Jan 23:S0945-053X(25)00012-5:PMID: 39863086.

90. Howlader N, Noone AM, Krapcho M, Miller D, Brest A, Yu M, et al. SEER Cancer Statistics Review, 1975–2018, National Cancer Institute. Bethesda, MD, https://seer.cancer.gov/csr/1975_2018/, based on November 2020 SEER data submission, posted to the SEER web site, April 2021. 2021.

91. Pal B, Chen Y, Vaillant F, Capaldo BD, Joyce R, Song X, et al. A single-cell RNA expression atlas of normal, preneoplastic and tumorigenic states in the human breast. The EMBO journal. 2021;40(11):e107333. PMC8167363.

