## Supplementary material for "Spatial Proteomics of the Normal Breast Collagen Stroma: Links to BI-RADS Categories and Body Mass Index": All supplemental

### Normal Breast Collagen Proteome Spatially Modulates with BMI and Breast Density Categories

**Supplemental Figure 1. Collagens detected by LC-MS/MS by six replicate runs on samples pooled by genetic ancestry.** A) Percentage of collagen types for donors of African ancestry and European ancestry. B) Total fibrillar collagen coverage on COL1A1, COL1A2, COL3A1. Green – modified hydroxyproline, yellow – unmodified region.

A) Collagen Type Proteins Detected by LC-MS/MS

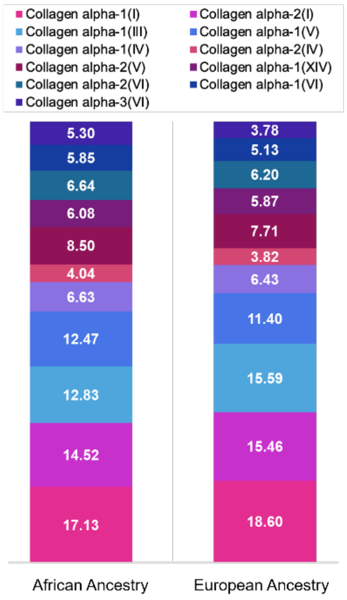

B) Sequence coverage of fibrillar collagen triple helical regions

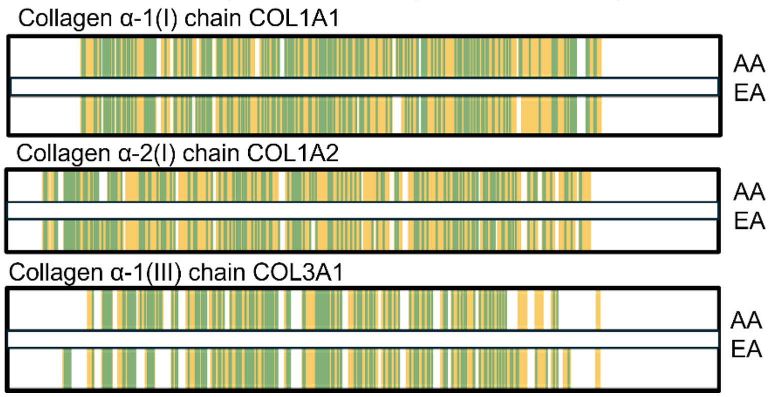

**Supplemental Figure 2. Samples and Markers Used for MALDI-IHC. A)** Donor samples used for MALDI-IHC were profiled by genetic ancestry. **B)** Markers used for MALDI-IHC and their general biological significance.

**A) Cohort used for multiplexed immunohistochemistry**

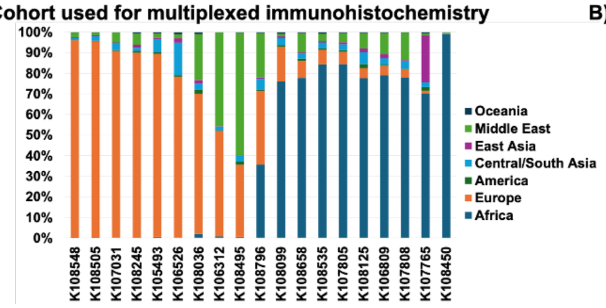

**B) Markers by multiplexed immunohistochemistry**

| Marker | Functions (Uniport.org) |
|---|---|
| CTNNB1 | Cell adhesions, Wnt signaling |
| VIM | Cell motility, orientation, mesenchymal |
| FN1 | Cell adhesion, motility |
| COL1A1 | Collagen alpha-1(I) chain |
| CD44 | Cell interactions, adhesions |
| CDH1 | Cell-cell adhesions, mobility, proliferation |
| CD68 | Macrophage marker |
| αSMA | Fibroblast marker |
| H3 | Histone |

**Supplemental Figure 3. Individual donor samples with the spatial distribution of each of the markers used in MALDI-IHC and an H&E-stained serial section.** Spatial distributions of 9 markers are shown alongside the H&E-stained tissue section.

Page 1 of 8

**K108548**

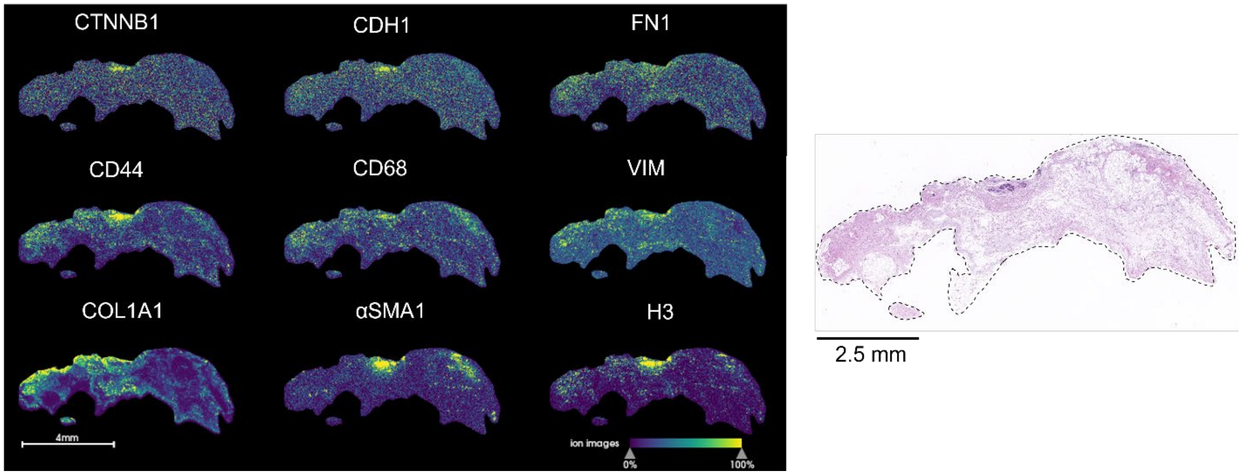

**K108505**

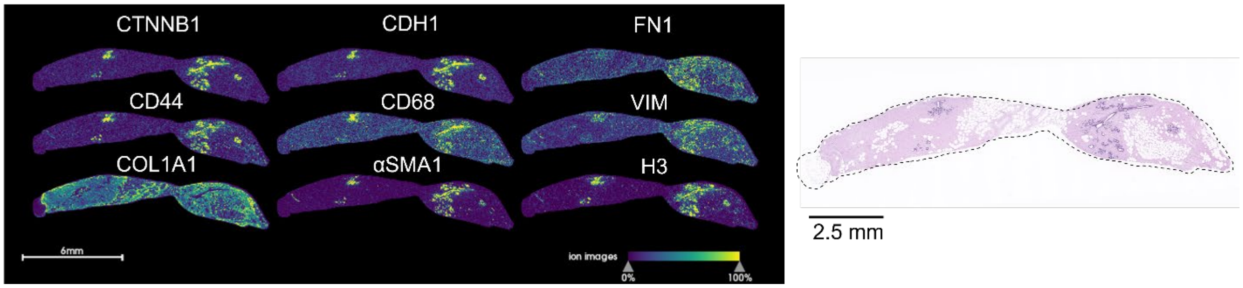

**K107031**

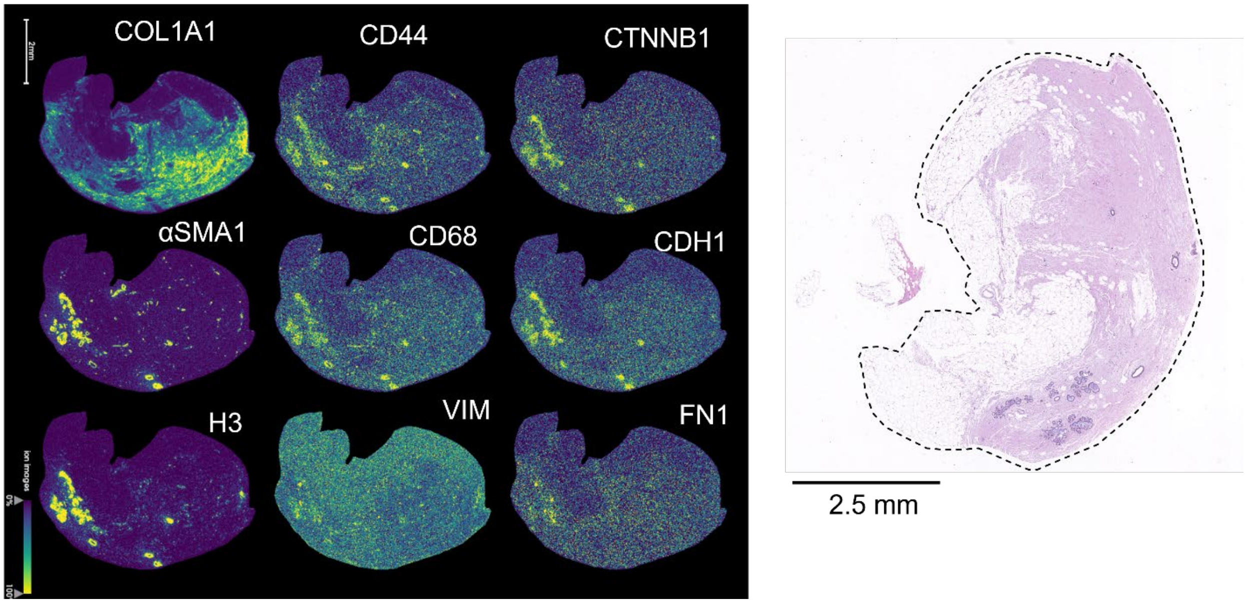

Page 2 of 8

**K107031**

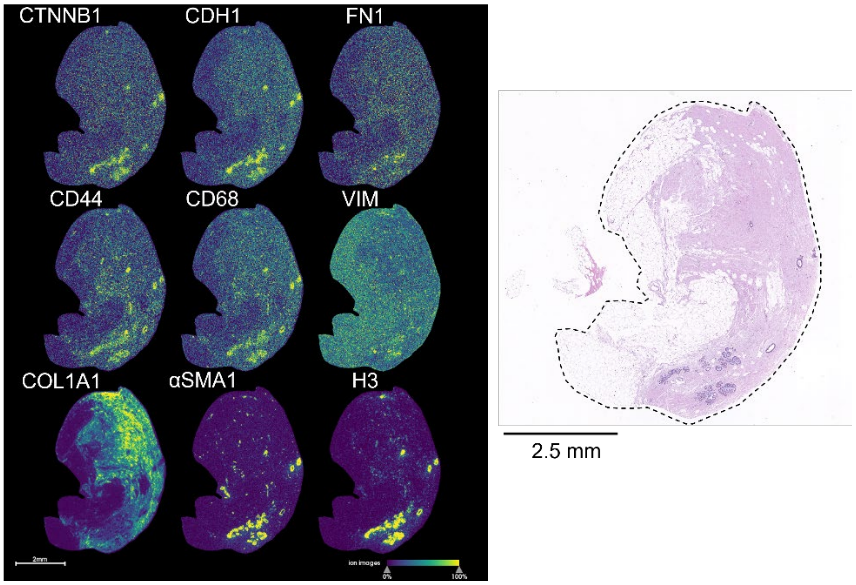

**K108245**

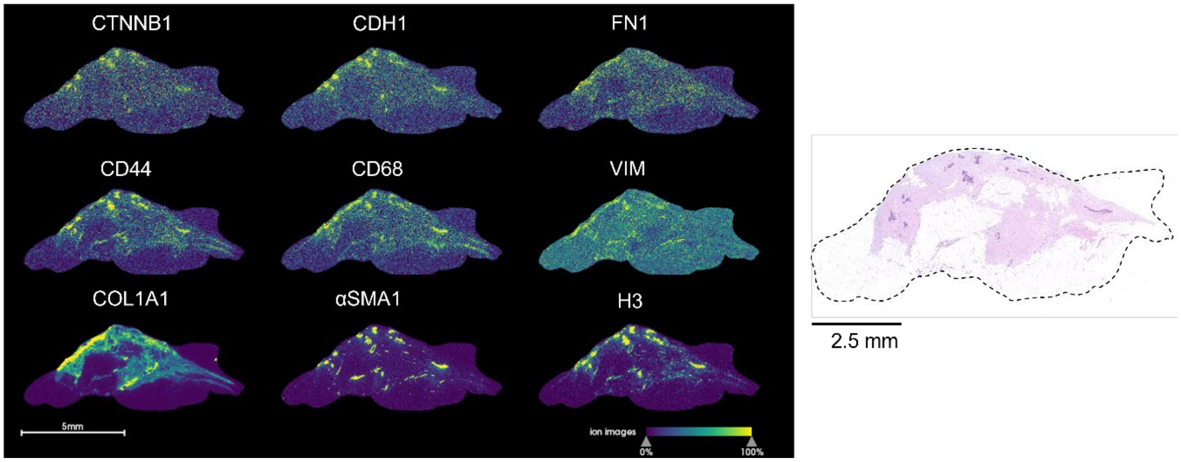

**K105493**

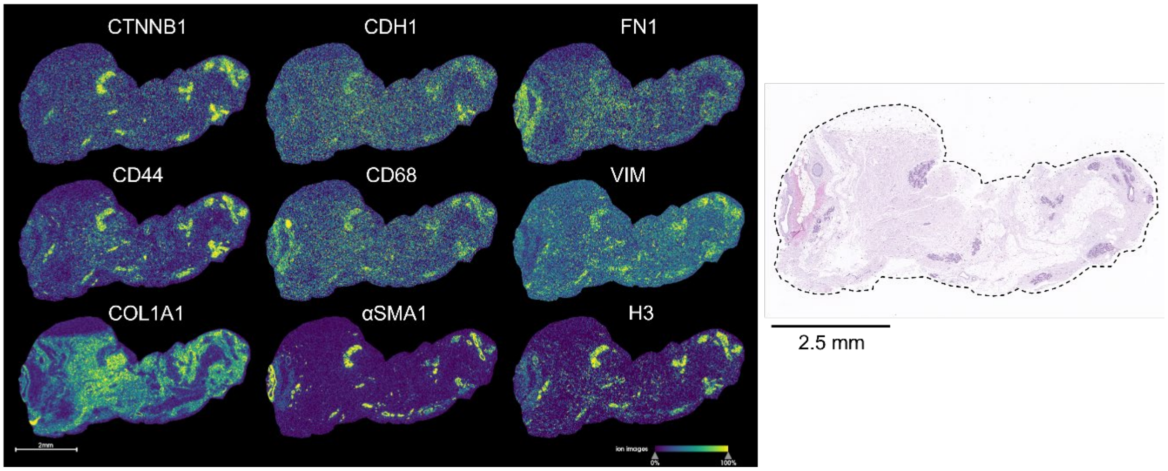

Page 3 of 8

**K106526**

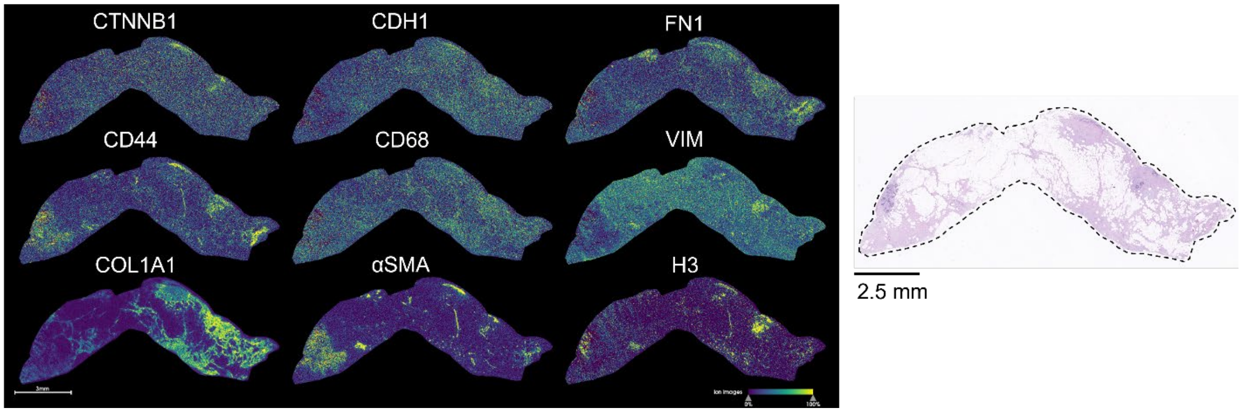

K108036

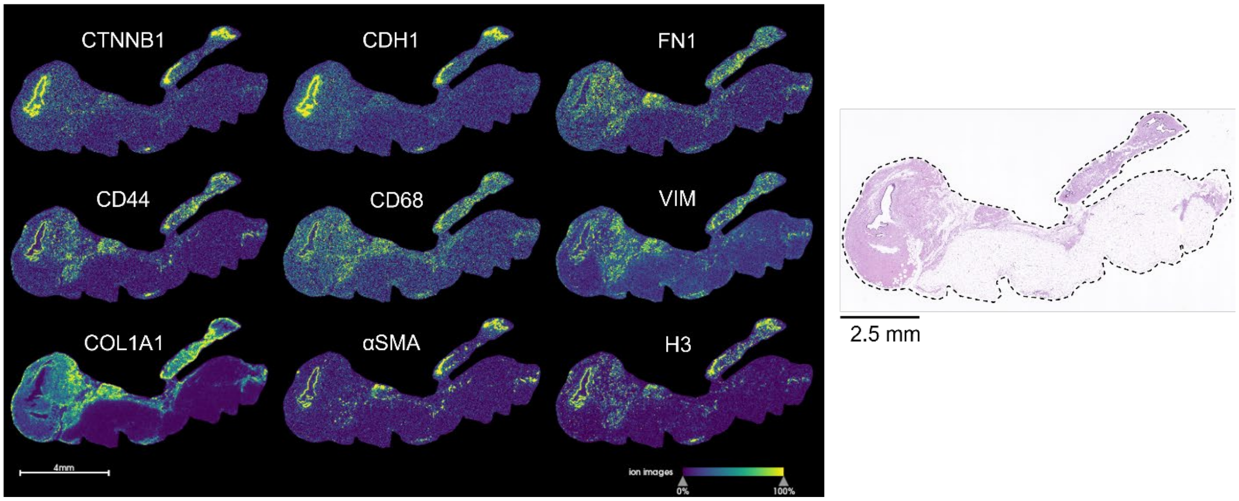

K107805

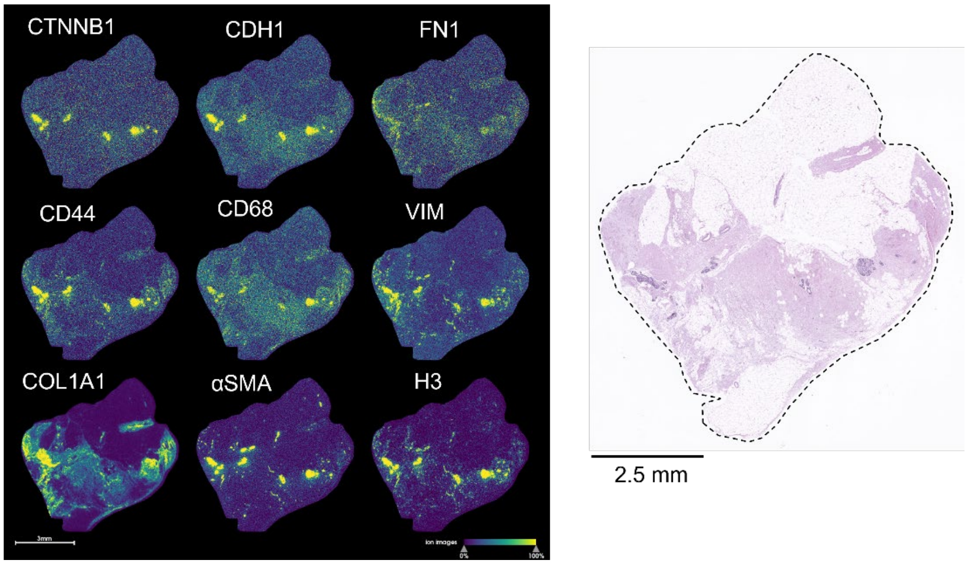

**K106312**

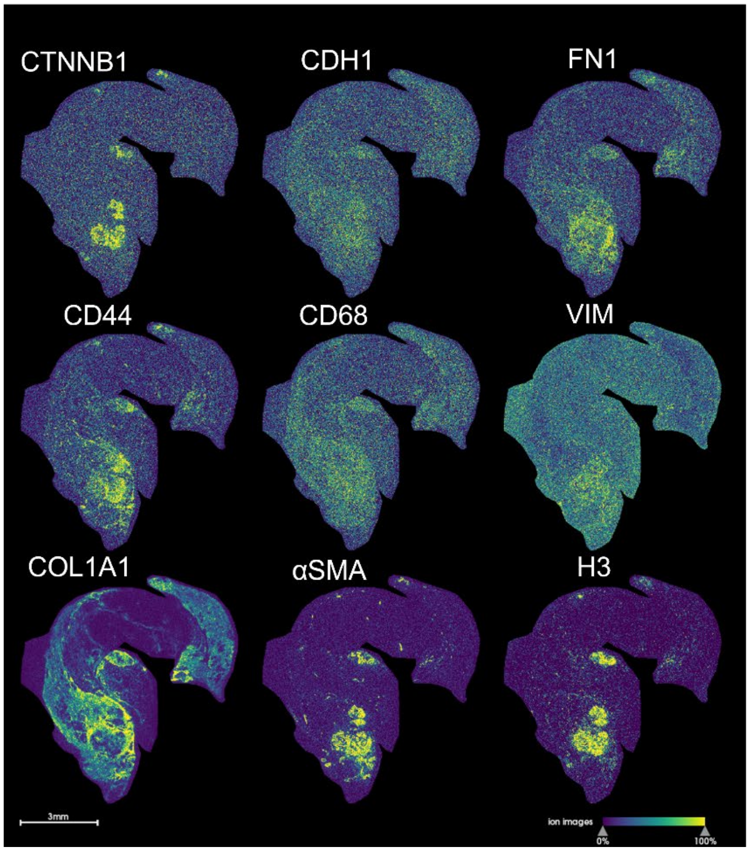

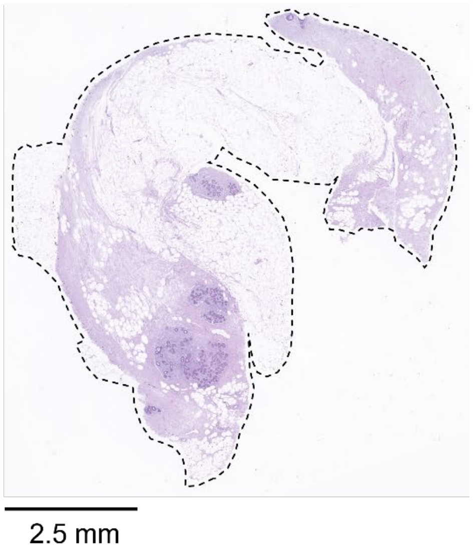

**K108495**

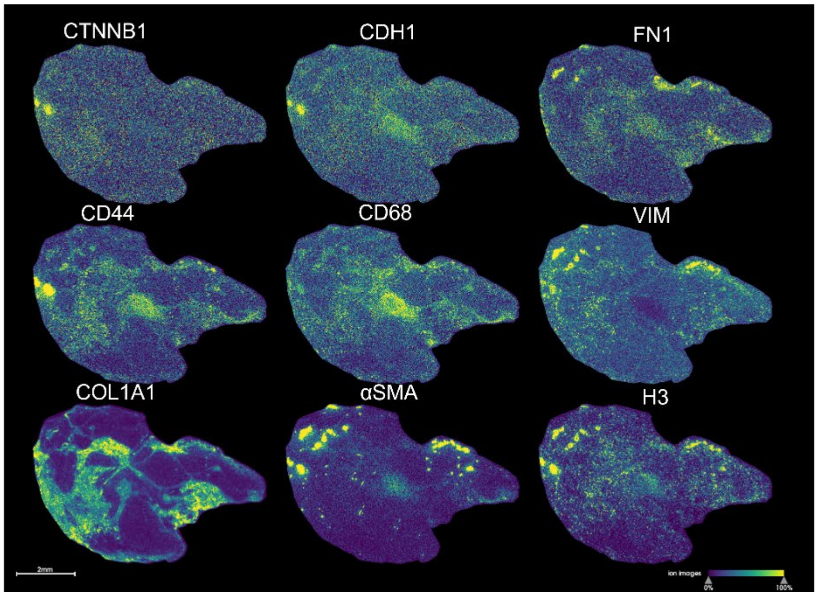

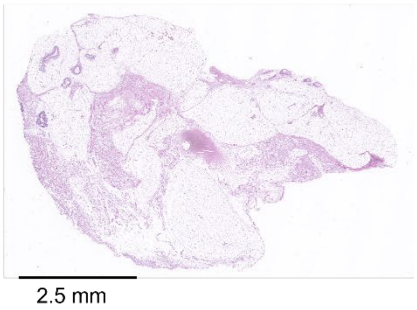

**K108796**

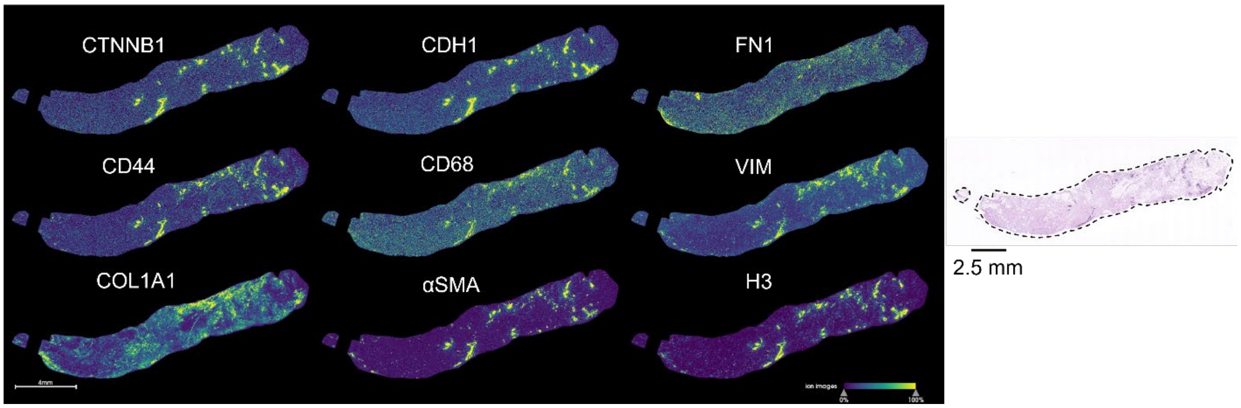

**K108099**

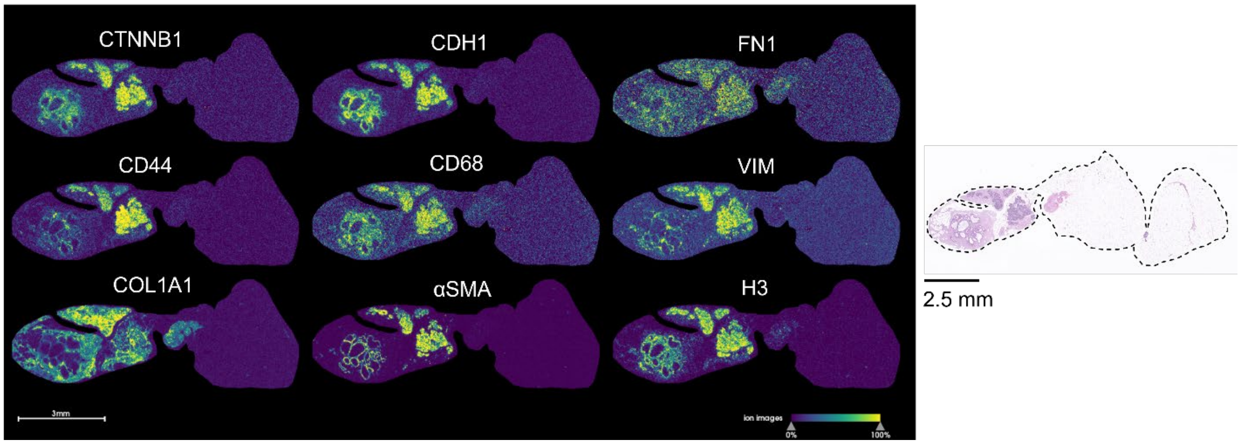

**K108658**

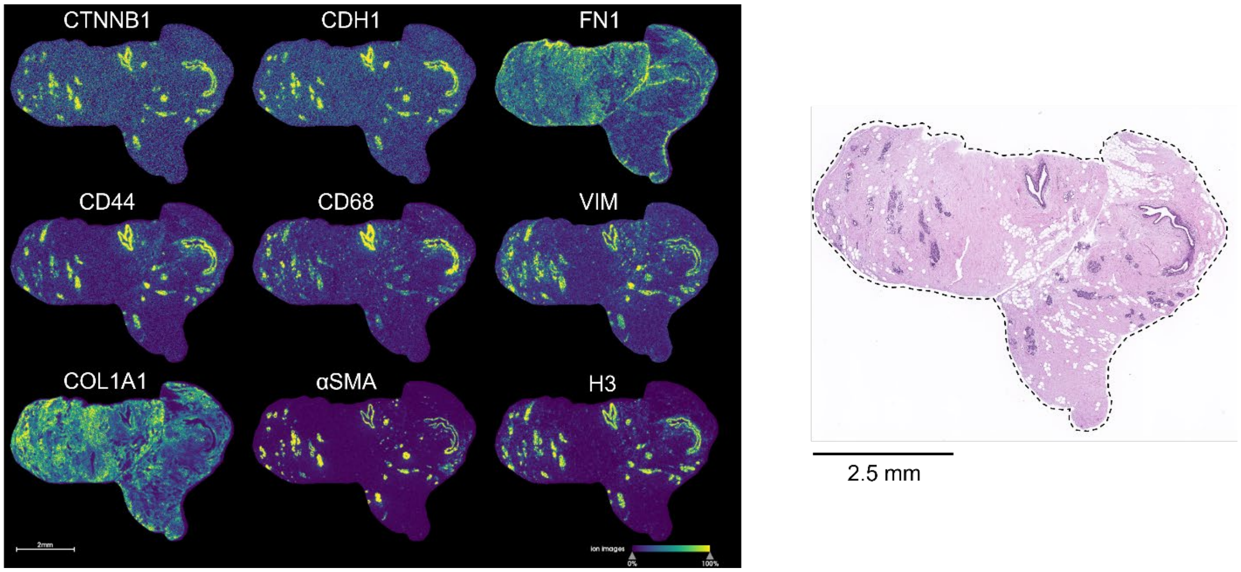

**K108535**

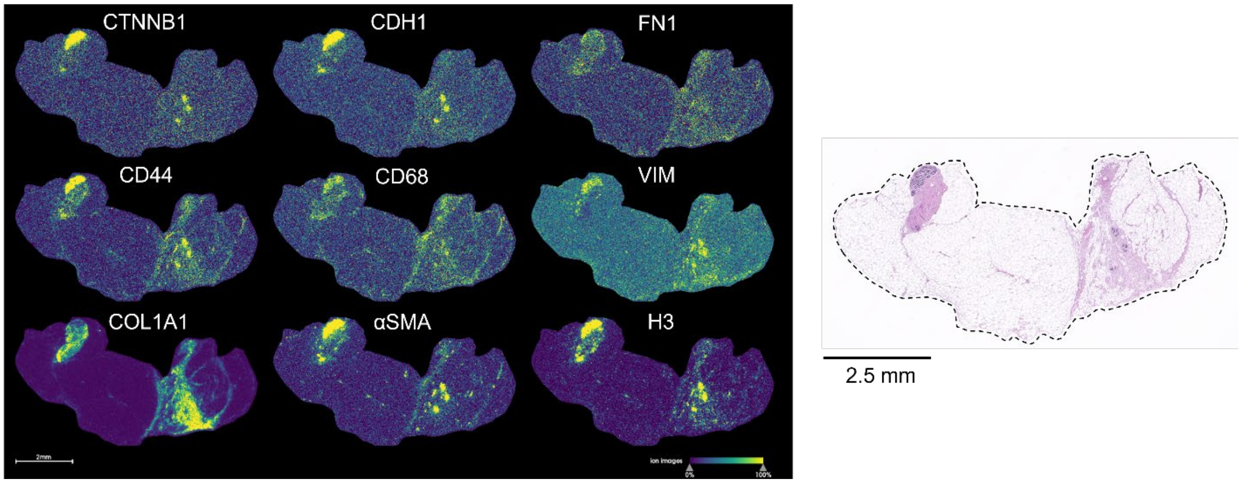

**K107765**

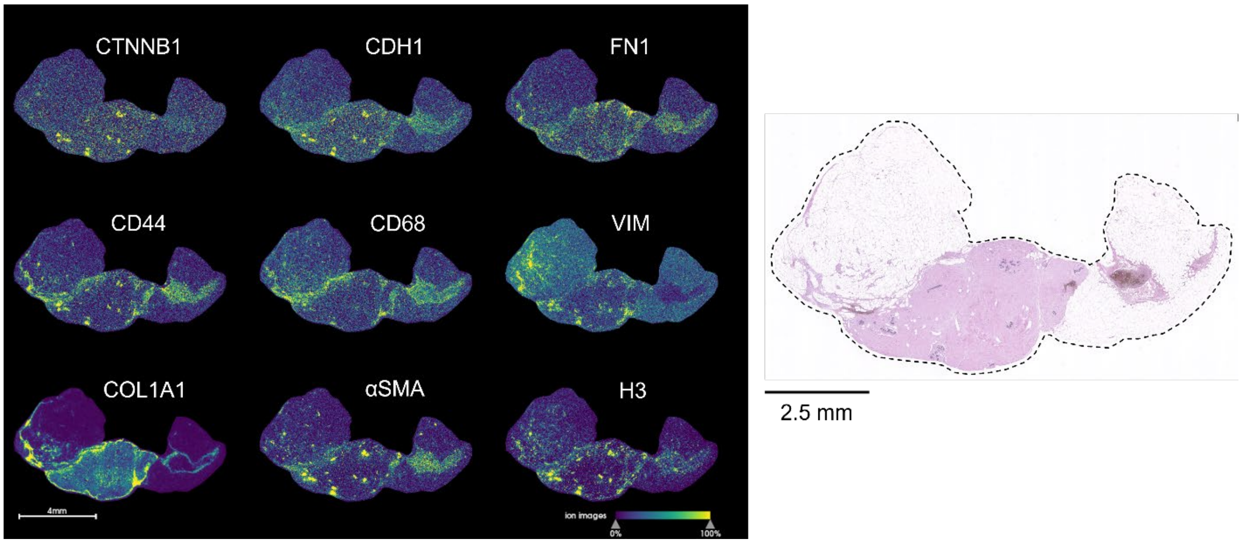

**K106809**

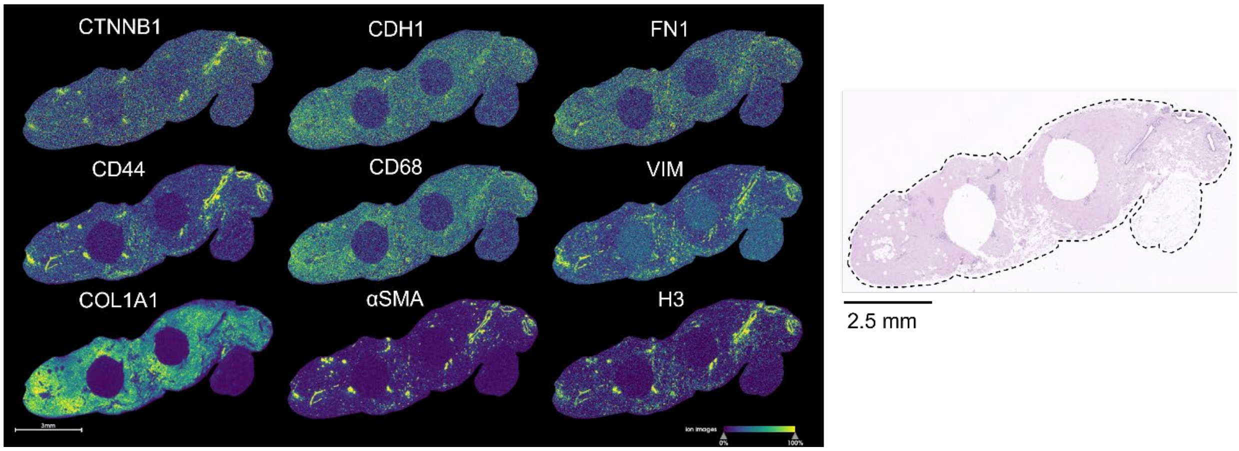

**K108125**

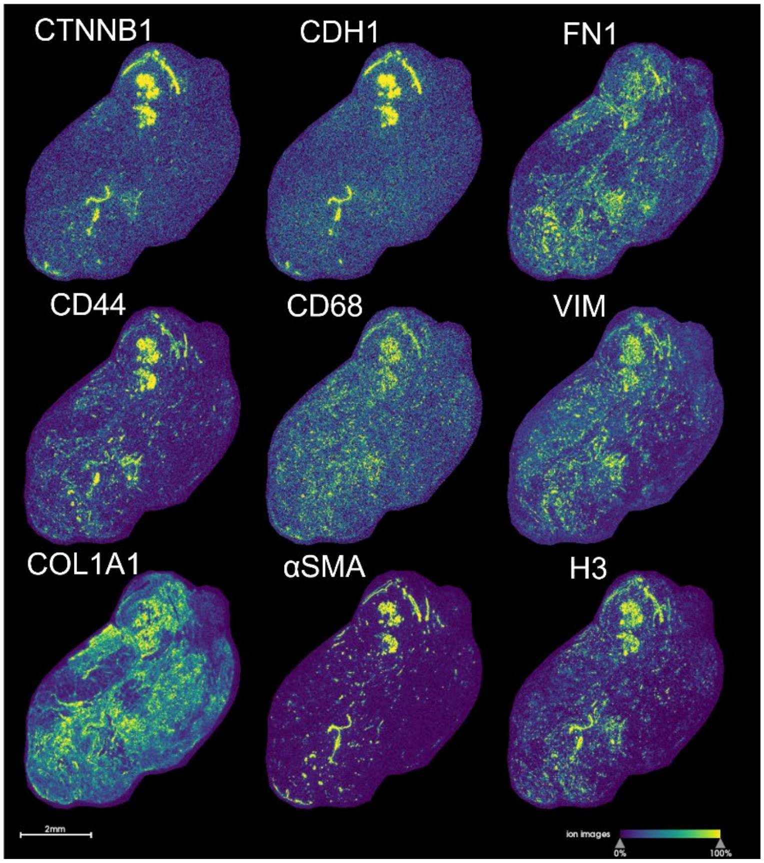

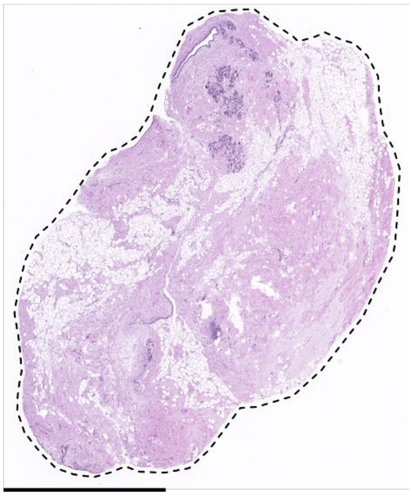

2.5 mm

**K107808**

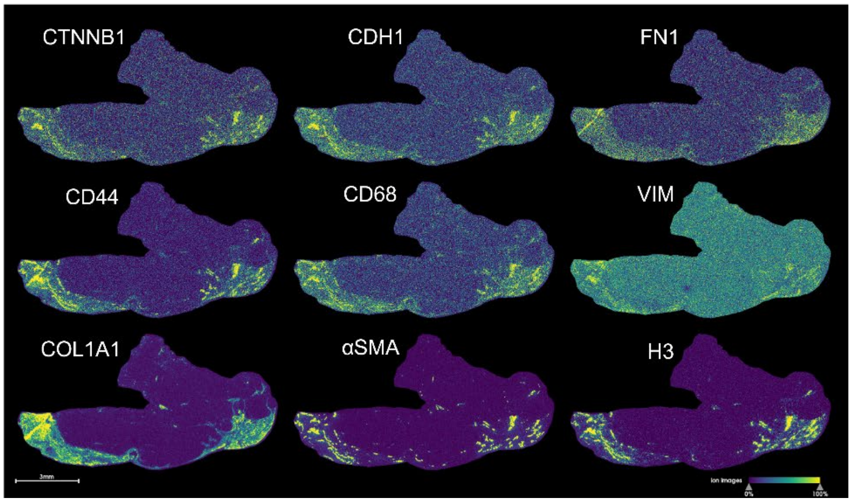

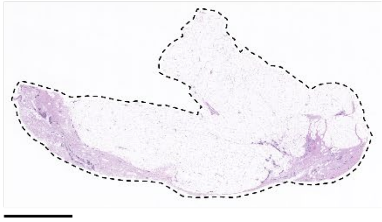

2.5 mm

**K108450**

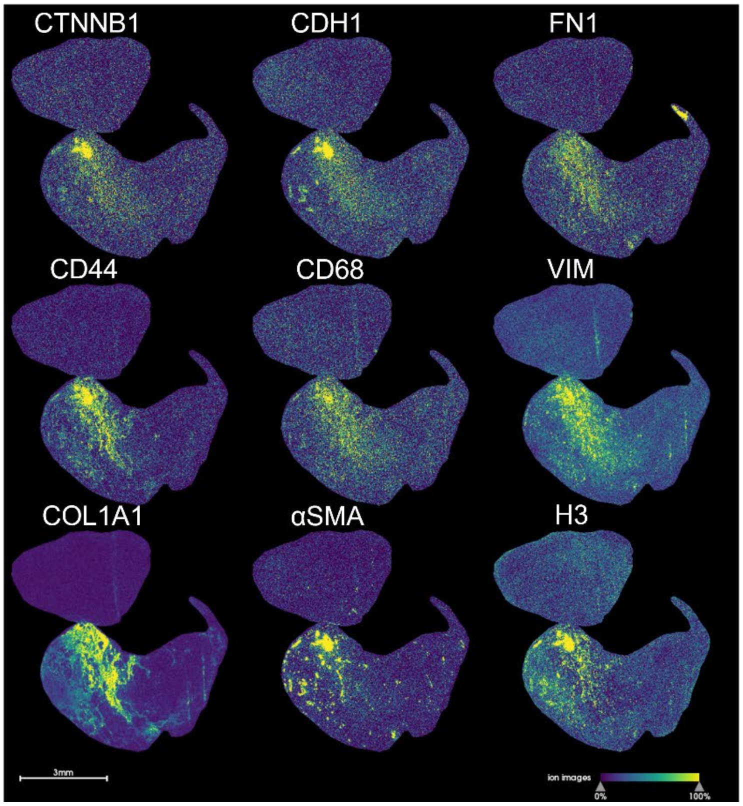
CTNNB1
CDH1
FN1
CD44
CD68
VIM
COL1A1
αSMA
H3
3mm
ion images
0%
100%

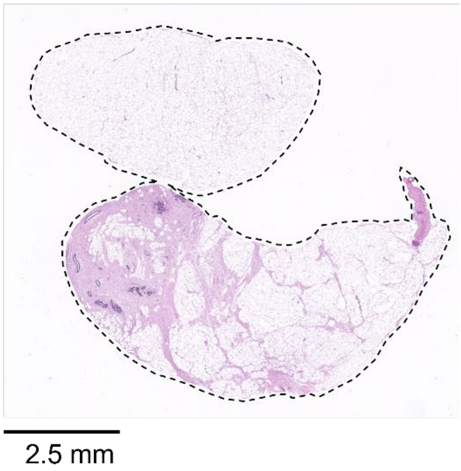
2.5 mm

**Supplemental Figure 4. Collagen fiber widths, including measurements from donors with healthy BMI.** All samples in different BMI categories show similar collagen fiber widths when measured in both A) ductal and B) fibrotic regions. Differences in collagen widths by BMI and genetic ancestry are observed only between overweight and obese samples in both C) ductal and D) fibrotic regions.

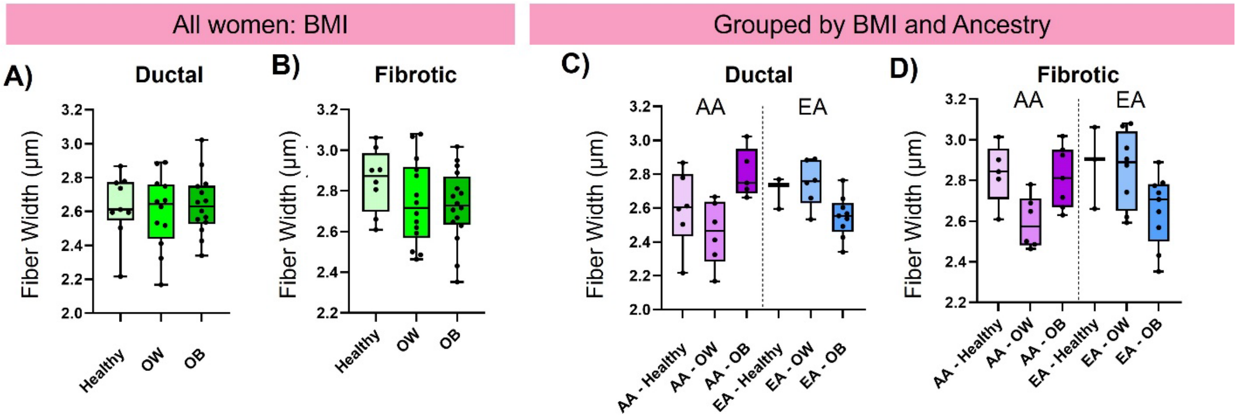

**Supplemental Figure 5. SCiLS Lab segmentation images.** Segmentation used identified COL1A1 peptides and the bisecting K-means algorithm using the Manhattan metric within SCiLS software.

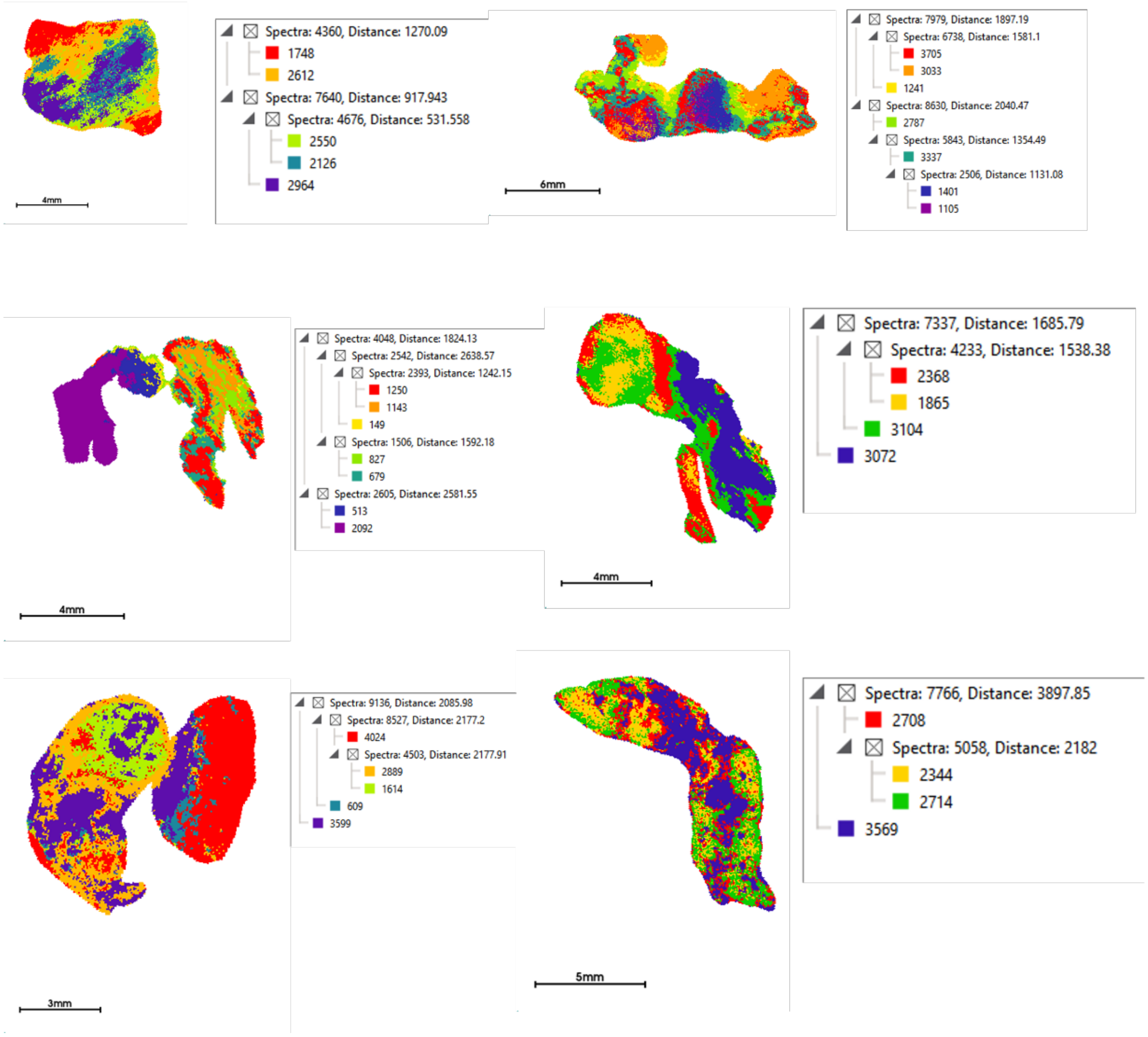

**Supplemental Figure 6. AUROC for 47 peptide domains that distinguish between density categories: scattered fibroglandular & heterogeneously dense.**

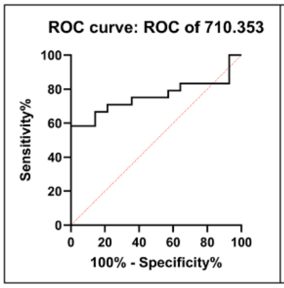

| | |
|---|---|
| AUROC | 0.7589 |
| Std. Error | 0.07849 |
| 95% CI | 0.6051 to 0.9128 |
| P value | 0.0085 |

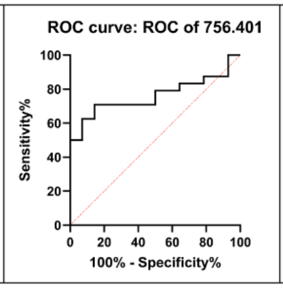

| | |
|---|---|
| AUROC | 0.7619 |
| Std. Error | 0.07767 |
| 95% CI | 0.6097 to 0.9141 |
| P value | 0.0077 |

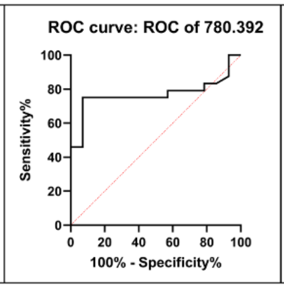

| | |
|---|---|
| AUROC | 0.7693 |
| Std. Error | 0.07946 |
| 95% CI | 0.6136 to 0.9251 |
| P value | 0.0062 |

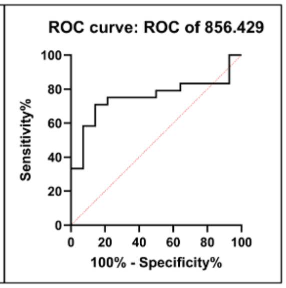

| | |
|---|---|
| AUROC | 0.7530 |
| Std. Error | 0.08059 |
| 95% CI | 0.5950 to 0.9109 |
| P value | 0.0101 |

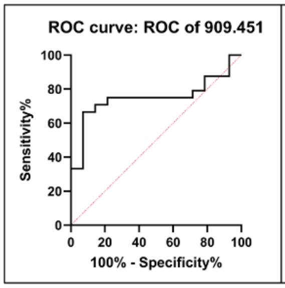

| | |
|---|---|
| AUROC | 0.7500 |
| Std. Error | 0.08157 |
| 95% CI | 0.5901 to 0.9099 |
| P value | 0.0110 |

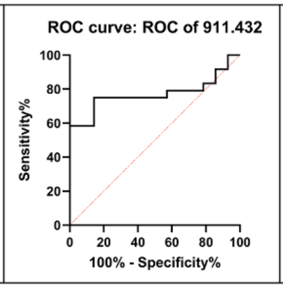

| | |
|---|---|
| AUROC | 0.7708 |
| Std. Error | 0.07771 |
| 95% CI | 0.6185 to 0.9231 |
| P value | 0.0059 |

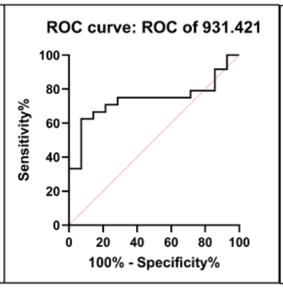

| | |
|---|---|
| AUROC | 0.7381 |
| Std. Error | 0.08240 |
| 95% CI | 0.5766 to 0.8996 |
| P value | 0.0155 |

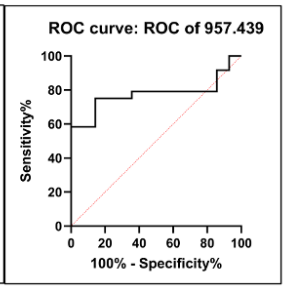

| | |
|---|---|
| AUROC | 0.7768 |
| Std. Error | 0.07734 |
| 95% CI | 0.6252 to 0.9284 |
| P value | 0.0049 |

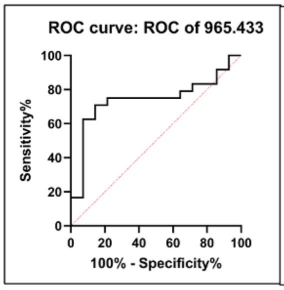

| | |
|---|---|
| AUROC | 0.7411 |
| Std. Error | 0.08430 |
| 95% CI | 0.5759 to 0.9063 |
| P value | 0.0142 |

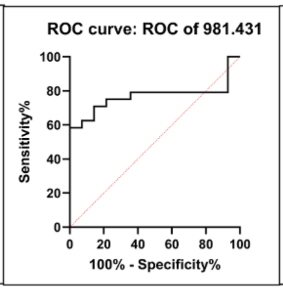

| | |
|---|---|
| AUROC | 0.7679 |
| Std. Error | 0.07951 |
| 95% CI | 0.6120 to 0.9237 |
| P value | 0.0065 |

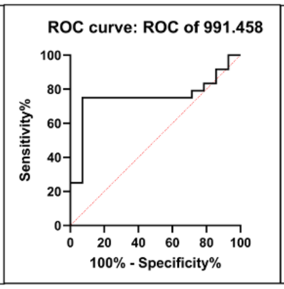

| | |
|---|---|
| AUROC | 0.7530 |
| Std. Error | 0.08345 |
| 95% CI | 0.5894 to 0.9165 |
| P value | 0.0101 |

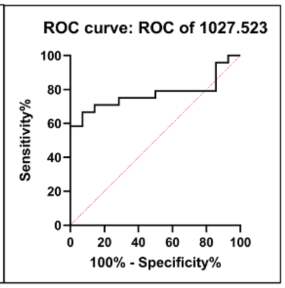

| | |
|---|---|
| AUROC | 0.7738 |
| Std. Error | 0.07721 |
| 95% CI | 0.6225 to 0.9251 |
| P value | 0.0054 |

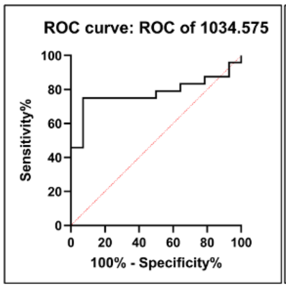

| | |
|---|---|
| AUROC | 0.7798 |
| Std. Error | 0.07716 |
| 95% CI | 0.6285 to 0.9310 |
| P value | 0.0044 |

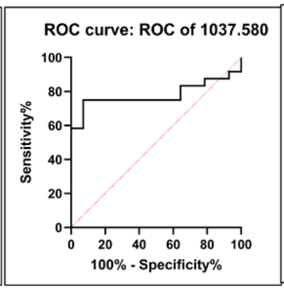

| | |
|---|---|
| AUROC | 0.7798 |
| Std. Error | 0.07772 |
| 95% CI | 0.6274 to 0.9321 |
| P value | 0.0044 |

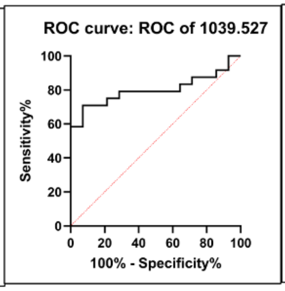

| | |
|---|---|
| AUROC | 0.8006 |
| Std. Error | 0.07258 |
| 95% CI | 0.6583 to 0.9429 |
| P value | 0.0022 |

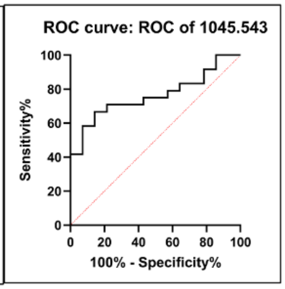

| | |
|---|---|
| AUROC | 0.7619 |
| Std. Error | 0.07694 |
| 95% CI | 0.6111 to 0.9127 |
| P value | 0.0077 |

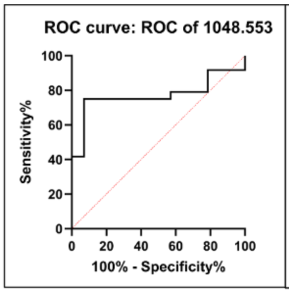

| | |
|---|---|
| AUROC | 0.7708 |
| Std. Error | 0.07901 |
| 95% CI | 0.6160 to 0.9257 |
| P value | 0.0059 |

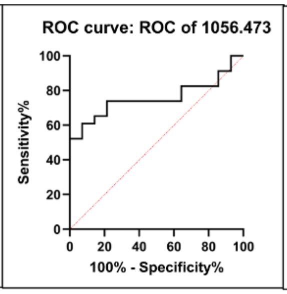

| | |
|---|---|
| AUROC | 0.7578 |
| Std. Error | 0.07994 |
| 95% CI | 0.6011 to 0.9144 |
| P value | 0.0093 |

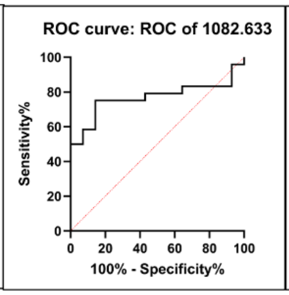

| | |
|---|---|
| AUROC | 0.7679 |
| Std. Error | 0.07830 |
| 95% CI | 0.6144 to 0.9213 |
| P value | 0.0065 |

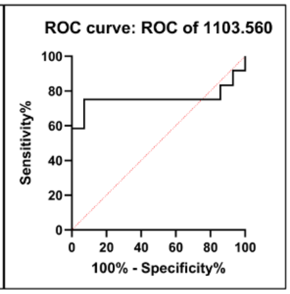

| | |
|---|---|
| AUROC | 0.7560 |
| Std. Error | 0.08339 |
| 95% CI | 0.5925 to 0.9194 |
| P value | 0.0093 |

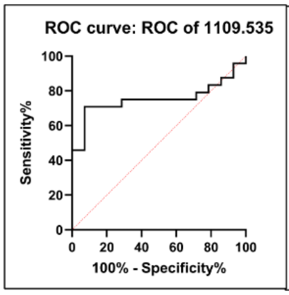

| | |
|---|---|
| AUROC | 0.7530 |
| Std. Error | 0.08087 |
| 95% CI | 0.5945 to 0.9115 |
| P value | 0.0101 |

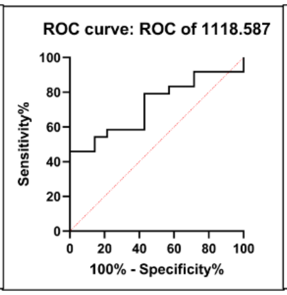

| | |
|---|---|
| AUROC | 0.7232 |
| Std. Error | 0.08172 |
| 95% CI | 0.5630 to 0.8834 |
| P value | 0.0232 |

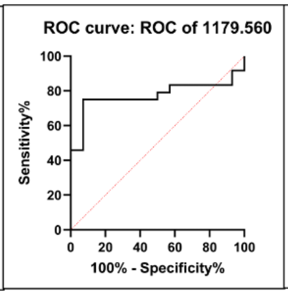

| | |
|---|---|
| AUROC | 0.7738 |
| Std. Error | 0.07889 |
| 95% CI | 0.6192 to 0.9284 |
| P value | 0.0054 |

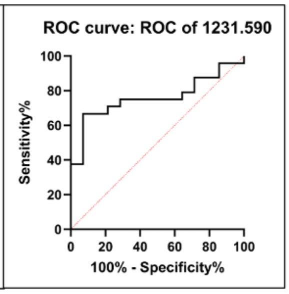

| | |
|---|---|
| AUROC | 0.7589 |
| Std. Error | 0.07871 |
| 95% CI | 0.6047 to 0.9132 |
| P value | 0.0085 |

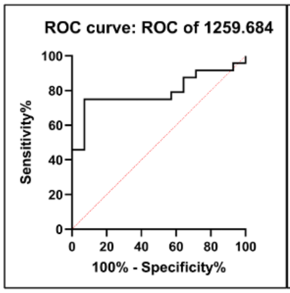

| | |
|---|---|
| AUROC | 0.7917 |
| Std. Error | 0.07453 |
| 95% CI | 0.6456 to 0.9377 |
| P value | 0.0030 |

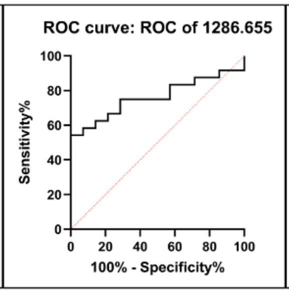

| | |
|---|---|
| AUROC | 0.7619 |
| Std. Error | 0.07714 |
| 95% CI | 0.6107 to 0.9131 |
| P value | 0.0077 |

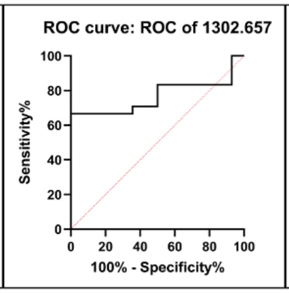

| | |
|---|---|
| AUROC | 0.7679 |
| Std. Error | 0.07807 |
| 95% CI | 0.6148 to 0.9209 |
| P value | 0.0065 |

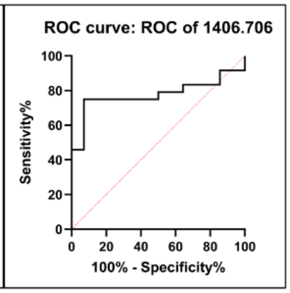

| | |
|---|---|
| AUROC | 0.7768 |
| Std. Error | 0.07788 |
| 95% CI | 0.6241 to 0.9294 |
| P value | 0.0049 |

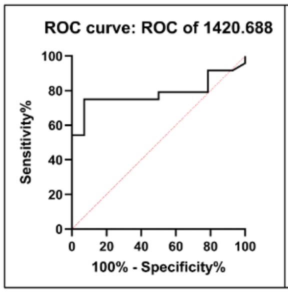

| | |
|---|---|
| AUROC | 0.7842 |
| Std. Error | 0.07651 |
| 95% CI | 0.6343 to 0.9342 |
| P value | 0.0039 |

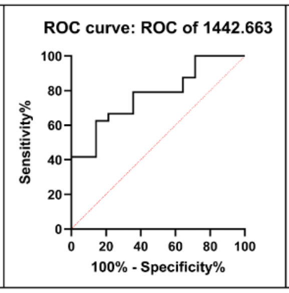

| | |
|---|---|
| AUROC | 0.7738 |
| Std. Error | 0.07564 |
| 95% CI | 0.6256 to 0.9221 |
| P value | 0.0054 |

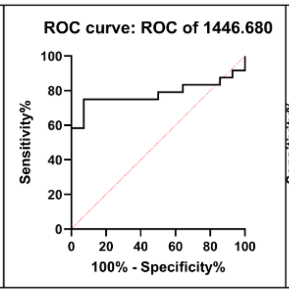

| | |
|---|---|
| AUROC | 0.7827 |
| Std. Error | 0.07729 |
| 95% CI | 0.6312 to 0.9342 |
| P value | 0.0040 |

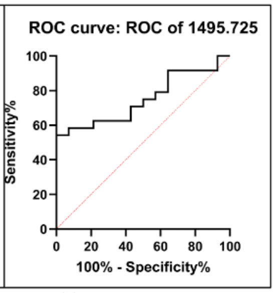

| | |
|---|---|
| AUROC | 0.7500 |
| Std. Error | 0.07811 |
| 95% CI | 0.5969 to 0.9031 |
| P value | 0.0110 |

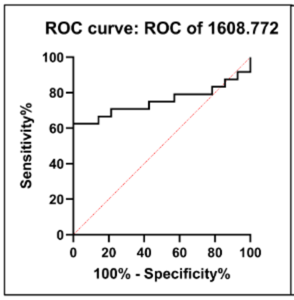

| | |
|---|---|
| AUROC | 0.7530 |
| Std. Error | 0.07999 |
| 95% CI | 0.5962 to 0.9098 |
| P value | 0.0101 |

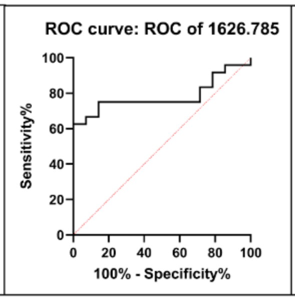

| | |
|---|---|
| AUROC | 0.7827 |
| Std. Error | 0.07627 |
| 95% CI | 0.6333 to 0.9322 |
| P value | 0.0040 |

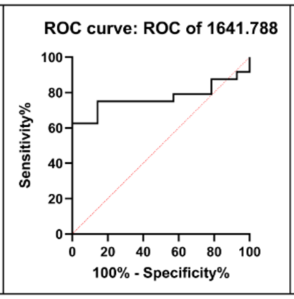

| | |
|---|---|
| AUROC | 0.7708 |
| Std. Error | 0.07849 |
| 95% CI | 0.6170 to 0.9247 |
| P value | 0.0059 |

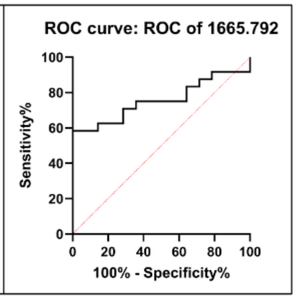

| | |
|---|---|
| AUROC | 0.7560 |
| Std. Error | 0.07790 |
| 95% CI | 0.6033 to 0.9086 |
| P value | 0.0093 |

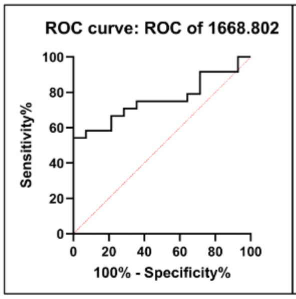

| | |
|---|---|
| AUROC | 0.7589 |
| Std. Error | 0.07711 |
| 95% CI | 0.6078 to 0.9101 |
| P value | 0.0085 |

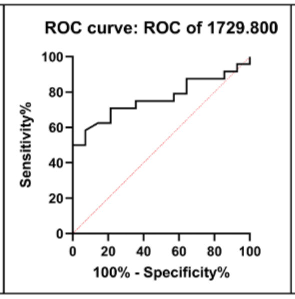

| | |
|---|---|
| AUROC | 0.7634 |
| Std. Error | 0.07670 |
| 95% CI | 0.6131 to 0.9137 |
| P value | 0.0074 |

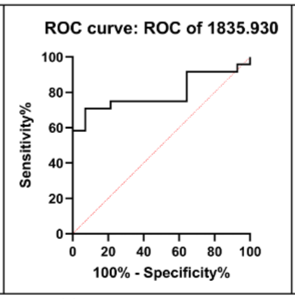

| | |
|---|---|
| AUROC | 0.7946 |
| Std. Error | 0.07343 |
| 95% CI | 0.6507 to 0.9386 |
| P value | 0.0027 |

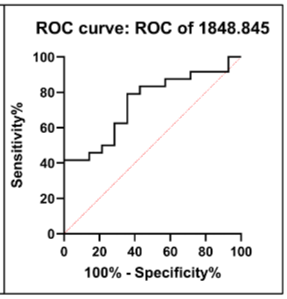

| | |
|---|---|
| AUROC | 0.7411 |
| Std. Error | 0.08092 |
| 95% CI | 0.5825 to 0.8997 |
| P value | 0.0142 |

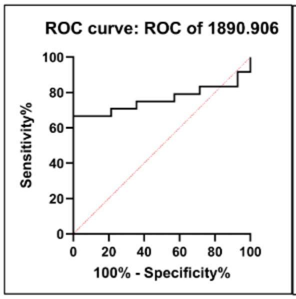

| | |
|---|---|
| AUROC | 0.7619 |
| Std. Error | 0.07964 |
| 95% CI | 0.6058 to 0.9180 |
| P value | 0.0077 |

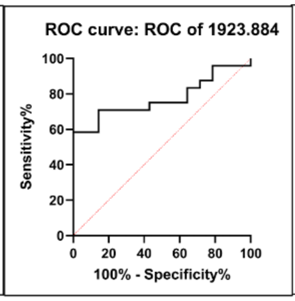

| | |
|---|---|
| AUROC | 0.7738 |
| Std. Error | 0.07555 |
| 95% CI | 0.6257 to 0.9219 |
| P value | 0.0054 |

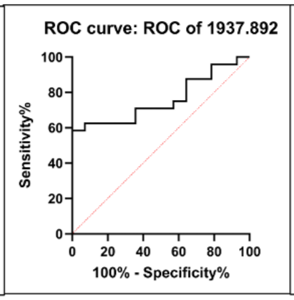

| | |
|---|---|
| AUROC | 0.7589 |
| Std. Error | 0.07724 |
| 95% CI | 0.6076 to 0.9103 |
| P value | 0.0085 |

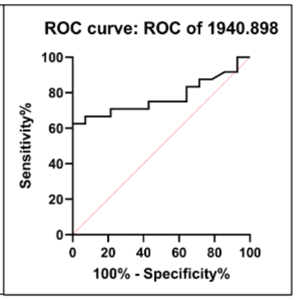

| | |
|---|---|
| AUROC | 0.7753 |
| Std. Error | 0.07582 |
| 95% CI | 0.6267 to 0.9239 |
| P value | 0.0051 |

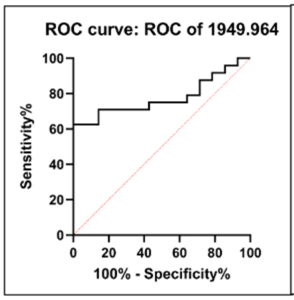

| | |
|---|---|
| AUROC | 0.7768 |
| Std. Error | 0.07555 |
| 95% CI | 0.6287 to 0.9249 |
| P value | 0.0049 |

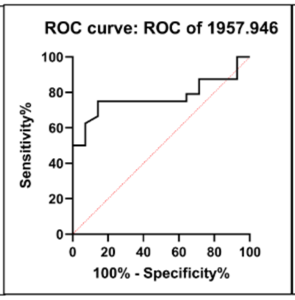

| | |
|---|---|
| AUROC | 0.7723 |
| Std. Error | 0.07731 |
| 95% CI | 0.6208 to 0.9239 |
| P value | 0.0056 |

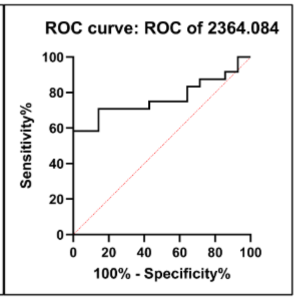

| | |
|---|---|
| AUROC | 0.7679 |
| Std. Error | 0.07663 |
| 95% CI | 0.6177 to 0.9180 |
| P value | 0.0065 |

**Supplemental Figure 7. Peptide domain P998.470 altered by ancestry within scattered fibroglandular BI-RADS category.** P-values are two-tailed t-tests after passing normality by Shapiro-Wilke (W>0.85; p-value>0.1).

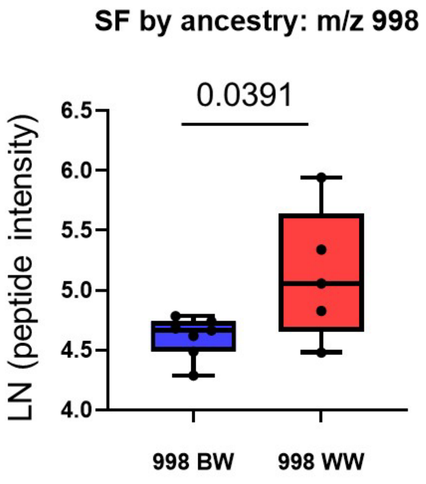

**Supplemental Figure 8. Peptide domain differences based on overweight or obese category and ancestry.** A &B) Peptides altered in obese category show higher expression in European ancestry (EA) samples compared to African ancestry (AA) samples. C) Overweight category showed increases in AA samples. p-values are two-tailed t-tests after passing normality by Shapiro-Wilke (W>0.85; p-value>0.1).

A) Obese category by ancestry

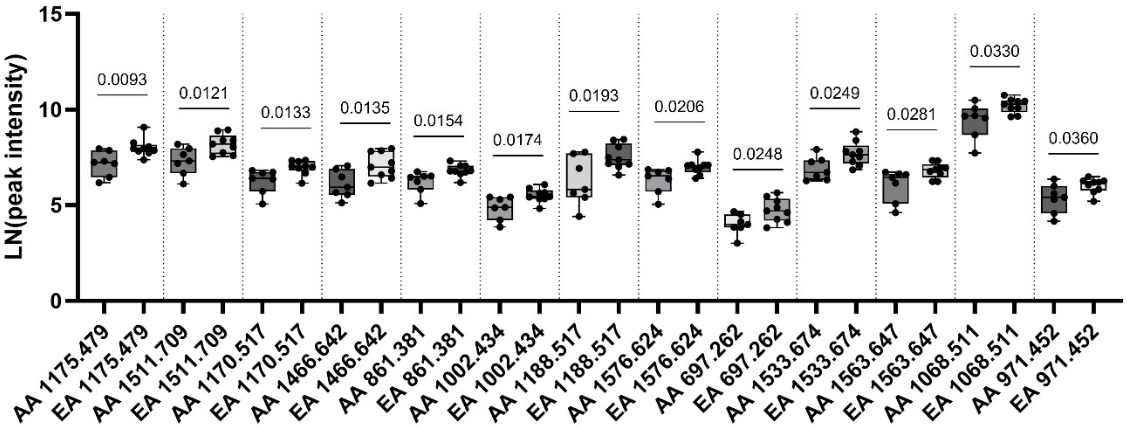

B) Obese category by ancestry

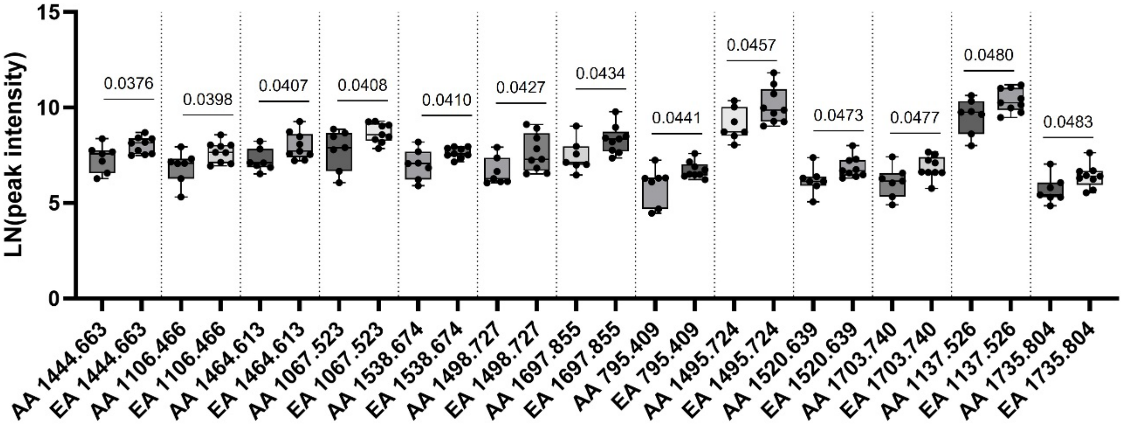

C) Overweight category by ancestry

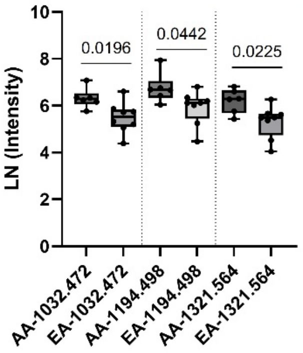

### Supplemental Table 1. Col1a1 peptide sequences from normal breast by LC-MS/MS.

| M+H | Gene | Domain Start | Domain End | Peptide | Modified Peptide [113] means HYP | Sequence with HYP probability PM:15.9949 | Hyperscore | Nextscore | Peptide Prophet Probability |
|---|---|---|---|---|---|---|---|---|---|
| 730.303 | COL1A1 | 1193 | 1199 | SAGFDFS | SAGFDFS | | 18.64 | 10.72 | 0.967 |
| 747.327 | COL1A1 | 1190 | 1197 | GPPSAGFD | GPPSAGFD | | 20.41 | 17.89 | 0.947 |
| 761.381 | COL1A1 | 405 | 412 | IAGAPGFP | IAGAP[113]GFP[113] | IAGAP(1.0000)GFP(1.0000) | 18.66 | 12.38 | 0.991 |
| 764.352 | COL1A1 | 872 | 880 | GATGFPGAA | GATGFP[113]GAA | GATGFP(1.0000)GAA | 18.25 | 14.78 | 0.994 |
| 767.428 | COL1A1 | 765 | 772 | LTGPIGPP | LTGPIGPP[113] | LTGP(0.0058)IGP(0.0064)P(0.9879) | 19.86 | 18.24 | 0.932 |
| 769.350 | COL1A1 | 887 | 895 | GPSGNAGPP | GPSGNAGPP[113] | GP(0.0053)SGNAGP(0.0062)P(0.9885) | 20.45 | 18.65 | 1.000 |
| 770.381 | COL1A1 | 809 | 817 | GPAGFAGPP | GPAGFAGPP | | 19.05 | 12.66 | 0.991 |
| 773.388 | COL1A1 | 174 | 181 | GISVPGPM | GISVPGPM[147] | GISVP(0.0057)GP(0.0100)M(0.9842) | 18.02 | 16.31 | 0.994 |
| 781.418 | COL1A1 | 851 | 859 | GPIGNVGAP | GPIGNVGAP | | 19.07 | 17.28 | 0.934 |
| 786.376 | COL1A1 | 809 | 817 | GPAGFAGPP | GPAGFAGPP[113] | GP(0.0054)AGFAGP(0.0070)P(0.9876) | 21.62 | 21.43 | 0.961 |
| 797.411 | COL1A1 | 851 | 859 | GPIGNVGAP | GPIGNVGAP[113] | GP(0.0105)IGNVGAP(0.9895) | 21.19 | 17.54 | 1.000 |
| 797.428 | COL1A1 | 878 | 886 | GAAGRVGPP | GAAGRVGPP[113] | GAAGRVGP(0.4770)P(0.5230) | 18.54 | 17.13 | 0.991 |
| 798.398 | COL1A1 | 851 | 859 | GPIGNVGAP | GPIGN[115]VGAP[113] | GP(0.0104)IGNVGAP(0.9896) | 19.43 | 14.64 | 0.958 |
| 802.410 | COL1A1 | 404 | 412 | GIAGAPGFP | GIAGAPGFP[113] | GIAGAP(0.0101)GFP(0.9899) | 18.83 | 14.08 | 0.976 |
| 806.366 | COL1A1 | 494 | 502 | GFPGADGVA | GFP[113]GADGVA | GFP(1.0000)GADGVA | 18.70 | 12.33 | 0.987 |
| 814.409 | COL1A1 | 341 | 349 | GPPGFPGAV | GPP[113]GFPGAV | GP(0.0062)P(0.9879)GFP(0.0058)GAV | 19.90 | 18.19 | 0.938 |
| 814.428 | COL1A1 | 254 | 262 | GLPGTAGLP | GLP[113]GTAGLP[113] | GLP(1.0000)GTAGLP(1.0000) | 21.38 | 13.74 | 0.975 |
| 818.396 | COL1A1 | 405 | 413 | IAGAPGFPG | IAGAP[113]GFP[113]G | IAGAP(1.0000)GFP(1.0000)G | 20.62 | 11.50 | 0.970 |
| 819.386 | COL1A1 | 1113 | 1120 | FSGLQGPP | FSGLQ[129]GPP[113] | FSGLQGP(0.0122)P(0.9878) | 18.05 | 16.20 | 0.967 |
| 823.426 | COL1A1 | 452 | 460 | GPVGVQGPP | GPVGVQGPP[113] | GP(0.0052)VGVQGP(0.0055)P(0.9893) | 21.15 | 21.15 | 0.920 |
| 827.353 | COL1A1 | 1192 | 1199 | PSAGFDFS | PSAGFDFS | | 18.55 | 11.91 | 1.000 |
| 829.369 | COL1A1 | 198 | 205 | PQGFQGPP | PQ[129]GFQ[129]GPP | | 19.44 | 12.13 | 0.994 |
| 830.407 | COL1A1 | 173 | 181 | GGISVPGPM | GGISVPGPM[147] | GGISVP(0.0050)GP(0.0103)M(0.9848) | 18.08 | 16.23 | 0.962 |
| 841.405 | COL1A1 | 548 | 556 | GPDGKTGPP | GPDGKTGPP[113] | GP(0.0050)DGKTGP(0.0082)P(0.9869) | 20.08 | 18.39 | 0.969 |
| 856.408 | COL1A1 | 473 | 481 | GEPGPTGLP | GEP[113]GPTGLP[113] | GEP(0.9872)GP(0.0256)TGLP(0.9872) | 19.04 | 14.34 | 0.946 |
| 856.448 | COL1A1 | 346 | 355 | PGAVGAKGEA | PGAVGAKGEA | | 24.85 | 13.88 | 1.000 |
| 858.394 | COL1A1 | 274 | 283 | DGAKGDAGPA | DGAKGDAGPA | | 20.74 | 18.74 | 0.952 |
| 863.399 | COL1A1 | 488 | 496 | GGPGSRGFP | GGP[113]GSRGFP[113] | GGP(1.0000)GSRGFP(1.0000) | 18.09 | 10.12 | 0.995 |
| 871.427 | COL1A1 | 341 | 350 | GPPGFPGAVG | GPP[113]GFPGAVG | GP(0.0063)P(0.9887)GFP(0.0050)GAVG | 21.46 | 19.95 | 0.988 |
| 875.390 | COL1A1 | 722 | 730 | GAPGLQGMP | GAP[113]GLQGM[147]P[113] | GAP(1.0000)GLQGM(1.0000)P(1.0000) | 19.23 | 13.53 | 0.936 |
| 875.426 | COL1A1 | 404 | 413 | GIAGAPGFPG | GIAGAP[113]GFP[113]G | GIAGAP(1.0000)GFP(1.0000)G | 19.04 | 14.11 | 0.957 |
| 876.374 | COL1A1 | 722 | 730 | GAPGLQGMP | GAP[113]GLQ[129]GM[147]P[113] | GAP(1.0000)GLQGM(1.0000)P(1.0000) | 20.05 | 10.28 | 0.999 |
| 876.405 | COL1A1 | 1112 | 1120 | GFSGLQGPP | GFSGLQ[129]GPP[113] | GFSGLQGP(0.0124)P(0.9876) | 20.79 | 18.85 | 0.964 |
| 879.397 | COL1A1 | 575 | 583 | GQAGVMGFP | GQAGVMGFP[113] | GQAGVM(0.0098)GFP(0.9902) | 19.26 | 16.21 | 0.953 |
| 883.428 | COL1A1 | 1187 | 1196 | GPPGPPSAGF | GPPGPPSAGF | | 21.58 | 14.91 | 0.994 |
| 885.408 | COL1A1 | 197 | 205 | GPQGFQGPP | GPQGFQ[129]GPP | | 22.90 | 14.76 | 0.989 |
| 886.392 | COL1A1 | 197 | 205 | GPQGFQGPP | GPQ[129]GFQ[129]GPP | | 19.65 | 13.51 | 0.961 |
| 887.423 | COL1A1 | 341 | 350 | GPPGFPGAVG | GPP[113]GFP[113]GAVG | GP(0.0611)P(0.9693)GFP(0.9696)GAVG | 22.04 | 21.73 | 0.961 |
| 891.400 | COL1A1 | 638 | 646 | GSPGFQGLP | GSP[113]GFQGLP[113] | GSP(1.0000)GFQGLP(1.0000) | 20.96 | 17.78 | 0.955 |
| 892.400 | COL1A1 | 638 | 646 | GSPGFQGLP | GSP[113]GFQ[129]GLP[113] | GSP(1.0000)GFQGLP(1.0000) | 19.33 | 15.40 | 0.974 |
| 894.392 | COL1A1 | 1190 | 1198 | GPPSAGFDF | GPPSAGFDF | | 25.66 | 11.95 | 1.000 |
| 895.381 | COL1A1 | 575 | 583 | GQAGVMGFP | GQAGVM[147]GFP[113] | GQAGVM(1.0000)GFP(1.0000) | 24.38 | 11.07 | 1.000 |
| 899.423 | COL1A1 | 1187 | 1196 | GPPGPPSAGF | GPP[113]GPPSAGF | GP(0.0036)P(0.9891)GP(0.0036)P(0.0037)SAGF | 23.26 | 21.08 | 0.961 |
| 902.383 | COL1A1 | 197 | 205 | GPQGFQGPP | GPQ[129]GFQ[129]GPP[113] | GP(0.0053)QGFQGP(0.0069)P(0.9877) | 18.36 | 18.24 | 0.949 |
| 903.425 | COL1A1 | 151 | 160 | GPPGLGGNFA | GPP[113]GLGGN[115]FA | GP(0.0115)P(0.9885)GLGGNFA | 18.38 | 16.24 | 1.000 |
| 910.389 | COL1A1 | 1190 | 1198 | GPPSAGFDF | GPP[113]SAGFDF | GP(0.0112)P(0.9888)SAGFDF | 18.43 | 16.50 | 0.985 |
| 911.445 | COL1A1 | 850 | 859 | PGPIGNVGAP | P[113]GPIGN[115]VGAP[113] | P(0.9891)GP(0.0218)IGNVGAP(0.9891) | 18.25 | 14.49 | 0.961 |
| 915.460 | COL1A1 | 172 | 181 | TGGISVPGPM | TGGISVPGPM | | 19.71 | 8.15 | 1.000 |
| 924.414 | COL1A1 | 1191 | 1199 | PPSAGFDFS | PPSAGFDFS | | 18.10 | 9.69 | 1.000 |
| 927.439 | COL1A1 | 661 | 670 | QGVPGDLGAP | Q[129]GVPGDLGAP[113] | QGVP(0.0289)GDLGAP(0.9711) | 18.08 | 12.15 | 0.975 |
| 931.450 | COL1A1 | 172 | 181 | TGGISVPGPM | TGGISVPGPM[147] | TGGISVP(0.0055)GP(0.0424)M(0.9521) | 18.83 | 17.68 | 1.000 |
| 941.476 | COL1A1 | 175 | 184 | ISVPGPMGPS | ISVPGPMGPS | | 19.10 | 10.50 | 0.999 |
| 942.448 | COL1A1 | 661 | 670 | QGVPGDLGAP | QGVP[113]GDLGAP[113] | QGVP(1.0000)GDLGAP(1.0000) | 19.79 | 13.29 | 0.977 |
| 943.410 | COL1A1 | 197 | 206 | GPQGFQGPPG | GPQ[129]GFQ[129]GPPG | | 25.60 | 18.40 | 0.955 |
| 946.460 | COL1A1 | 404 | 414 | GIAGAPGFPGA | GIAGAP[113]GFP[113]GA | GIAGAP(1.0000)GFP(1.0000)GA | 19.22 | 12.33 | 0.981 |
| 947.374 | COL1A1 | 208 | 217 | PGEPGASGPM | P[113]GEP[113]GASGPM[147] | P(0.9864)GEP(0.9864)GASGP(0.0409)M(0.9864) | 23.09 | 21.17 | 0.984 |
| 948.440 | COL1A1 | 638 | 647 | GSPGFQGLPG | GSP[113]GFQGLP[113]G | GSP(1.0000)GFQGLP(1.0000)G | 18.46 | 15.50 | 0.958 |
| 955.488 | COL1A1 | 1200 | 1207 | FLPQPPQE | FLPQPPQE | | 19.83 | 11.72 | 0.986 |
| 956.474 | COL1A1 | 1200 | 1207 | FLPQPPQE | FLPQPPQ[129]E | | 19.84 | 12.79 | 0.998 |
| 957.468 | COL1A1 | 175 | 184 | ISVPGPMGPS | ISVPGPM[147]GPS | ISVP(0.0035)GP(0.0048)M(0.9879)GP(0.0037)S | 22.24 | 18.46 | 1.000 |
| 959.411 | COL1A1 | 197 | 206 | GPQGFQGPPG | GPQ[129]GFQ[129]GPP[113]G | GP(0.0056)QGFQGP(0.0073)P(0.9871)G | 20.13 | 18.29 | 0.946 |
| 964.473 | COL1A1 | 1001 | 1011 | GPPGLAGPPGE | GPPGLAGPP[113]GE | GP(0.0033)P(0.0033)GLAGP(0.0037)P(0.9897)GE | 26.67 | 22.45 | 0.982 |
| 966.449 | COL1A1 | 270 | 279 | FSGLDGAKGD | FSGLDGAKGD | | 24.31 | 12.14 | 1.000 |
| 971.476 | COL1A1 | 273 | 283 | LDGAKGDAGPA | LDGAKGDAGPA | | 21.32 | 12.56 | 0.987 |
| 981.410 | COL1A1 | 1190 | 1199 | GPPSAGFDFS | GPPSAGFDFS | | 26.01 | 14.81 | 1.000 |
| 984.496 | COL1A1 | 1150 | 1159 | PGKDGLNGLP | PGKDGLN[115]GLP[113] | P(0.0126)GKDGLNGLP(0.9874) | 18.12 | 13.41 | 0.955 |
| 984.549 | COL1A1 | 760 | 769 | DGVRGLTGPI | DGVRGLTGPI | | 19.82 | 11.64 | 0.990 |
| 985.453 | COL1A1 | 398 | 409 | GANGAPGIAGAP | GAN[115]GAP[113]GIAGAP[113] | GANGAP(1.0000)GIAGAP(1.0000) | 21.36 | 13.30 | 0.978 |
| 997.419 | COL1A1 | 1190 | 1199 | GPPSAGFDFS | GPP[113]SAGFDFS | GP(0.0113)P(0.9887)SAGFDFS | 30.26 | 27.65 | 1.000 |
| 998.450 | COL1A1 | 1187 | 1197 | GPPGPPSAGFD | GPPGPPSAGFD | | 28.97 | 18.69 | 1.000 |
| 1000.476 | COL1A1 | 1196 | 1203 | FDFSFLPQ | FDFSFLPQ | | 18.56 | 9.96 | 0.994 |
| 1002.490 | COL1A1 | 171 | 181 | STGGISVPGPM | STGGISVPGPM | | 18.23 | 11.59 | 0.991 |
| 1014.447 | COL1A1 | 1187 | 1197 | GPPGPPSAGFD | GPP[113]GPPSAGFD | GP(0.0053)P(0.9876)GP(0.0035)P(0.0035)SAGFD | 23.82 | 23.51 | 0.996 |
| 1015.437 | COL1A1 | 196 | 205 | PGPQGFQGPP | P[113]GPQ[129]GFQ[129]GPP[113] | P(0.9879)GP(0.0110)QGFQGP(0.0132)P(0.9878) | 21.40 | 19.47 | 0.993 |
| 1017.467 | COL1A1 | 320 | 331 | GARGNDGATGAA | GARGNDGATGAA | | 18.48 | 11.02 | 0.986 |
| 1018.458 | COL1A1 | 320 | 331 | GARGNDGATGAA | GARGN[115]DGATGAA | | 18.81 | 12.53 | 0.997 |
| 1018.483 | COL1A1 | 171 | 181 | STGGISVPGPM | STGGISVPGP[113]M | STGGISVP(0.0041)GP(0.4980)M(0.4980) | 18.53 | 18.53 | 1.000 |
| 1028.493 | COL1A1 | 272 | 283 | GLDGAKGDAGPA | GLDGAKGDAGPA | | 21.06 | 17.92 | 0.977 |
| 1030.443 | COL1A1 | 1187 | 1197 | GPPGPPSAGFD | GPP[113]GPP[113]SAGFD | GP(0.0110)P(0.9886)GP(0.0118)P(0.9886)SAGFD | 21.96 | 19.95 | 0.996 |
| 1034.573 | COL1A1 | 1076 | 1087 | GPVGPVGARGPA | GPVGPVGARGPA | | 21.55 | 12.82 | 0.980 |

| | | | | | | | | | |
|---|---|---|---|---|---|---|---|---|---|
| 1037.482 | COL1A1 | 270 | 280 | FSGLDGAKGDA | FSGLDGAKGDA | | 20.13 | 12.38 | 0.983 |
| 1041.529 | COL1A1 | 347 | 358 | GAVGAKGEAGPQ | GAVGAKGEAGPQ | | 27.59 | 15.29 | 0.998 |
| 1042.475 | COL1A1 | 932 | 943 | GEKGSPGADGPA | GEKGSPGADGPA | | 21.63 | 13.53 | 0.981 |
| 1042.513 | COL1A1 | 347 | 358 | GAVGAKGEAGPQ | GAVGAKGEAGPQ[129] | | 25.64 | 14.56 | 1.000 |
| 1043.507 | COL1A1 | 1145 | 1156 | GSAGAPGKDGLN | GSAGAPGKDGLN | | 19.03 | 13.57 | 0.991 |
| 1043.509 | COL1A1 | 401 | 412 | GAPGIAGAPGFP | GAP[113]GIAGAPGFP[113] | GAP(0.9888)GIAGAP(0.0224)GFP(0.9888) | 20.97 | 14.98 | 0.988 |
| 1044.494 | COL1A1 | 1145 | 1156 | GSAGAPGKDGLN | GSAGAPGKDGLN[115] | | 19.71 | 9.94 | 1.000 |
| 1045.544 | COL1A1 | 405 | 415 | IAGAPGFPGAR | IAGAP[113]GFP[113]GAR | IAGAP(1.0000)GFP(1.0000)GAR | 19.91 | 15.87 | 0.988 |
| 1057.484 | COL1A1 | 608 | 619 | GPAGKDGEAGAQ | GPAGKDGEAGAQ | | 23.27 | 9.80 | 1.000 |
| 1058.472 | COL1A1 | 776 | 787 | GAPGDKGESGPS | GAPGDKGESGPS | | 19.11 | 12.43 | 1.000 |
| 1059.504 | COL1A1 | 1145 | 1156 | GSAGAPGKDGLN | GSAGAP[113]GKDGLN | GSAGAP(1.0000)GKDGLN | 19.15 | 13.84 | 1.000 |
| 1059.505 | COL1A1 | 401 | 412 | GAPGIAGAPGFP | GAP[113]GIAGAP[113]GFP[113] | GAP(1.0000)GIAGAP(1.0000)GFP(1.0000) | 35.88 | 17.42 | 1.000 |
| 1060.524 | COL1A1 | 536 | 547 | GAKGLTGSPGSP | GAKGLTGSP[113]GSP[113] | GAKGLTGSP(1.0000)GSP(1.0000) | 21.38 | 13.16 | 0.981 |
| 1066.637 | COL1A1 | 954 | 964 | IAGQRGVVGLP | IAGQRGVVGLP | | 18.72 | 14.08 | 0.977 |
| 1067.625 | COL1A1 | 954 | 964 | IAGQRGVVGLP | IAGQ[129]RGVVGLP | | 19.94 | 11.16 | 0.985 |
| 1071.492 | COL1A1 | 660 | 670 | EQGVPGDLGAP | EQGVP[113]GDLGAP[113] | EQGVP(1.0000)GDLGAP(1.0000) | 20.90 | 14.15 | 0.973 |
| 1072.455 | COL1A1 | 197 | 207 | GPQGFQGPPGE | GPQ[129]GFQ[129]GPPGE | | 22.25 | 15.16 | 0.987 |
| 1074.465 | COL1A1 | 776 | 787 | GAPGDKGESGPS | GAP[113]GDKGESGPS | GAP(0.9858)GDKGESGP(0.0142)S | 20.87 | 13.65 | 0.991 |
| 1076.553 | COL1A1 | 872 | 883 | GATGFPGAAGRV | GATGFP[113]GAAGRV | GATGFP(1.0000)GAAGRV | 19.91 | 15.80 | 1.000 |
| 1078.477 | COL1A1 | 1189 | 1199 | PGPPSAGFDFS | PGPPSAGFDFS | | 18.31 | 9.94 | 0.997 |
| 1082.623 | COL1A1 | 954 | 964 | IAGQRGVVGLP | IAGQRGVVGLP[113] | IAGQRGVVGLP(1.0000) | 21.83 | 16.05 | 0.977 |
| 1083.610 | COL1A1 | 954 | 964 | IAGQRGVVGLP | IAGQ[129]RGVVGLP[113] | IAGQRGVVGLP(1.0000) | 23.71 | 13.38 | 0.999 |
| 1084.500 | COL1A1 | 986 | 997 | GPSGASGERGPP | GPSGASGERGPP[113] | GP(0.0057)SGASGERGP(0.0080)P(0.9863) | 22.17 | 22.17 | 0.999 |
| 1088.447 | COL1A1 | 197 | 207 | GPQGFQGPPGE | GPQ[129]GFQ[129]GPP[113]GE | GP(0.0053)QGFQGP(0.0071)P(0.9876)GE | 21.21 | 17.39 | 0.939 |
| 1094.477 | COL1A1 | 1189 | 1199 | PGPPSAGFDFS | P[113]GPPSAGFDFS | P(0.9897)GP(0.0052)P(0.0051)SAGFDFS | 20.64 | 15.94 | 1.000 |
| 1094.512 | COL1A1 | 269 | 280 | GFSGLDGAKGDA | GFSGLDGAKGDA | | 21.65 | 7.48 | 1.000 |
| 1097.517 | COL1A1 | 977 | 988 | GPSGEPGKQGPS | GPSGEPGKQGPS | | 22.27 | 22.27 | 1.000 |
| 1098.505 | COL1A1 | 977 | 988 | GPSGEPGKQGPS | GPSGEPGKQ[129]GPS | | 19.88 | 19.88 | 0.938 |
| 1098.592 | COL1A1 | 251 | 262 | GARGLPGTAGLP | GARGLP[113]GTAGLP[113] | GARGLP(1.0000)GTAGLP(1.0000) | 20.58 | 11.40 | 0.998 |
| 1102.563 | COL1A1 | 404 | 415 | GIAGAPGFPGAR | GIAGAP[113]GFP[113]GAR | GIAGAP(1.0000)GFP(1.0000)GAR | 19.93 | 9.48 | 1.000 |
| 1106.525 | COL1A1 | 491 | 502 | GSRGFPGADGVA | GSRGFP[113]GADGVA | GSRGFP(1.0000)GADGVA | 19.79 | 10.63 | 1.000 |
| 1110.514 | COL1A1 | 1007 | 1018 | GPPGESGREGAP | GPPGESGREGAP | | 20.38 | 12.77 | 0.982 |
| 1112.513 | COL1A1 | 659 | 670 | GEQGVPGDLGAP | GEQGVPGDLGAP[113] | GEQGVP(0.0165)GDLGAP(0.9835) | 23.21 | 14.58 | 0.985 |
| 1112.547 | COL1A1 | 1148 | 1159 | GAPGKDGLNGLP | GAPGKDGLN[115]GLP[113] | GAP(0.0107)GKDGLNGLP(0.9893) | 23.61 | 13.15 | 0.999 |
| 1113.492 | COL1A1 | 659 | 670 | GEQGVPGDLGAP | GEQ[129]GVPGDLGAP[113] | GEQGVP(0.0106)GDLGAP(0.9894) | 23.71 | 14.34 | 0.998 |
| 1113.514 | COL1A1 | 977 | 988 | GPSGEPGKQGPS | GPSGEP[113]GKQGPS | GP(0.0055)SGEP(0.9841)GKQGP(0.0104)S | 24.06 | 24.06 | 0.972 |
| 1114.501 | COL1A1 | 977 | 988 | GPSGEPGKQGPS | GPSGEP[113]GKQ[129]GPS | GP(0.0051)SGEP(0.9874)GKQGP(0.0075)S | 21.20 | 21.20 | 0.981 |
| 1115.525 | COL1A1 | 271 | 283 | SGLDGAKGDAGPA | SGLDGAKGDAGPA | | 26.44 | 17.24 | 1.000 |
| 1115.540 | COL1A1 | 1031 | 1042 | GAKGDRGETGPA | GAKGDRGETGPA | | 19.88 | 9.57 | 0.990 |
| 1116.527 | COL1A1 | 635 | 646 | GPAGSPGFQGLP | GPAGSP[113]GFQGLP[113] | GP(0.0232)AGSP(0.9884)GFQGLP(0.9884) | 19.06 | 14.27 | 0.979 |
| 1117.498 | COL1A1 | 635 | 646 | GPAGSPGFQGLP | GPAGSP[113]GFQ[129]GLP[113] | GP(0.0208)AGSP(0.9896)GFQGLP(0.9896) | 21.06 | 17.67 | 0.996 |
| 1118.473 | COL1A1 | 1019 | 1030 | GAEGSPGRDGSP | GAEGSP[113]GRDGSP[113] | GAEGSP(1.0000)GRDGSP(1.0000) | 18.80 | 12.15 | 0.984 |
| 1122.557 | COL1A1 | 914 | 925 | GPAGRPGEVGPP | GPAGRP[113]GEVGPP[113] | GP(0.0116)AGRP(0.9881)GEVGP(0.0122)P(0.9881) | 21.37 | 21.37 | 0.922 |
| 1125.525 | COL1A1 | 626 | 637 | GPAGERGEQGPA | GPAGERGEQGPA | | 22.35 | 22.35 | 0.957 |
| 1126.507 | COL1A1 | 1007 | 1018 | GPPGESGREGAP | GPPGESGREGAP[113] | GP(0.0063)P(0.0076)GESGREGAP(0.9861) | 21.48 | 13.18 | 0.967 |
| 1128.496 | COL1A1 | 1190 | 1200 | GPPSAGFDFSF | GPPSAGFDFSF | | 25.93 | 11.64 | 1.000 |
| 1128.512 | COL1A1 | 659 | 670 | GEQGVPGDLGAP | GEQGVP[113]GDLGAP[113] | GEQGVP(1.0000)GDLGAP(1.0000) | 24.32 | 13.01 | 0.997 |
| 1128.524 | COL1A1 | 151 | 162 | GPPGLGGNFAPQ | GPP[113]GLGGN[115]FAPQ | GP(0.0062)P(0.9879)GLGGNFAP(0.0059)Q | 20.95 | 18.69 | 0.989 |
| 1128.552 | COL1A1 | 1148 | 1159 | GAPGKDGLNGLP | GAP[113]GKDGLN[115]GLP[113] | GAP(1.0000)GKDGLNGLP(1.0000) | 19.77 | 16.88 | 0.926 |
| 1131.536 | COL1A1 | 1061 | 1072 | GKSGDRGETGPA | GKSGDRGETGPA | | 21.20 | 7.42 | 1.000 |
| 1139.656 | COL1A1 | 954 | 965 | IAGQRGVVGLPG | IAGQRGVVGLP[113]G | IAGQRGVVGLP(1.0000)G | 21.49 | 10.78 | 0.998 |
| 1140.636 | COL1A1 | 953 | 964 | GIAGQRGVVGLP | GIAGQ[129]RGVVGLP[113] | GIAGQRGVVGLP(1.0000) | 20.38 | 15.87 | 0.966 |
| 1142.507 | COL1A1 | 1007 | 1018 | GPPGESGREGAP | GPP[113]GESGREGAP[113] | GP(0.0198)P(0.9901)GESGREGAP(0.9901) | 20.04 | 17.43 | 0.980 |
| 1142.544 | COL1A1 | 524 | 535 | GEAGRPGEAGLP | GEAGRP[113]GEAGLP[113] | GEAGRP(1.0000)GEAGLP(1.0000) | 18.32 | 12.86 | 0.945 |
| 1145.520 | COL1A1 | 1187 | 1198 | GPPGPPSAGFDF | GPPGPPSAGFDF | | 24.00 | 17.58 | 0.995 |
| 1154.507 | COL1A1 | 797 | 808 | GAPGDRGEPGPP | GAP[113]GDRGEP[113]GPP[113] | GAP(0.9880)GDRGEP(0.9880)GP(0.0359)P(0.9880) | 20.81 | 20.81 | 0.961 |
| 1154.561 | COL1A1 | 179 | 190 | GPMGPSGPRGLP | GPM[147]GPSGPRGLP[113] | GP(0.0113)M(0.9864)GP(0.0072)SGP(0.0086)RGLP(0.9866) | 20.08 | 18.22 | 0.984 |
| 1154.595 | COL1A1 | 740 | 751 | GPKGDRGDAGPK | GPKGDRGDAGPK | | 19.12 | 15.32 | 0.986 |
| 1155.526 | COL1A1 | 650 | 661 | GPPGEAGKPGEQ | GPP[113]GEAGKP[113]GEQ | GP(0.0205)P(0.9897)GEAGKP(0.9897)GEQ | 23.64 | 20.90 | 0.945 |
| 1156.549 | COL1A1 | 607 | 619 | VGPAGKDGEAGAQ | VGPAGKDGEAGAQ | | 22.49 | 17.68 | 0.967 |
| 1161.512 | COL1A1 | 1187 | 1198 | GPPGPPSAGFDF | GPP[113]GPPSAGFDF | GP(0.0058)P(0.9829)GP(0.0069)P(0.0043)SAGFDF | 32.45 | 30.28 | 1.000 |
| 1163.564 | COL1A1 | 572 | 583 | GARGQAGVMGFP | GARGQAGVMGFP[113] | GARGQAGVM(0.0107)GFP(0.9893) | 20.86 | 11.96 | 0.998 |
| 1164.550 | COL1A1 | 572 | 583 | GARGQAGVMGFP | GARGQ[129]AGVMGFP[113] | GARGQAGVM(0.0107)GFP(0.9893) | 21.07 | 12.49 | 0.998 |
| 1172.575 | COL1A1 | 752 | 764 | GADGSPGKDGVRG | GADGSPGKDGVRG | | 18.76 | 10.35 | 1.000 |
| 1175.544 | COL1A1 | 725 | 736 | GLQGMPGERGAA | GLQGM[147]P[113]GERGAA | GLQGM(1.0000)P(1.0000)GERGAA | 18.86 | 11.15 | 0.965 |
| 1177.508 | COL1A1 | 1187 | 1198 | GPPGPPSAGFDF | GPP[113]GPP[113]SAGFDF | GP(0.0127)P(0.9861)GP(0.0152)P(0.9860)SAGFDF | 24.60 | 22.48 | 0.999 |
| 1179.560 | COL1A1 | 572 | 583 | GARGQAGVMGFP | GARGQAGVM[147]GFP[113] | GARGQAGVM(1.0000)GFP(1.0000) | 22.67 | 12.36 | 0.999 |
| 1179.573 | COL1A1 | 365 | 376 | GPQGVRGEPGPP | GPQGVRGEP[113]GPP[113] | GP(0.0135)QGVRGEP(0.9876)GP(0.0113)P(0.9876) | 23.68 | 23.68 | 0.988 |
| 1180.531 | COL1A1 | 572 | 583 | GARGQAGVMGFP | GARGQ[129]AGVM[147]GFP[113] | GARGQAGVM(1.0000)GFP(1.0000) | 22.83 | 11.28 | 0.972 |
| 1186.528 | COL1A1 | 362 | 373 | GSEGPQGVRGEP | GSEGPQ[129]GVRGEP[113] | GSEGP(0.0193)QGVRGEP(0.9807) | 18.62 | 13.56 | 1.000 |
| 1188.557 | COL1A1 | 752 | 764 | GADGSPGKDGVRG | GADGSP[113]GKDGVRG | GADGSP(1.0000)GKDGVRG | 18.05 | 4.15 | 1.000 |
| 1189.596 | COL1A1 | 1198 | 1207 | FSFLPQPPQE | FSFLPQPPQE | | 20.79 | 12.22 | 0.974 |
| 1196.562 | COL1A1 | 625 | 637 | AGPAGERGEQGPA | AGPAGERGEQGPA | | 20.34 | 20.34 | 0.988 |
| 1205.559 | COL1A1 | 485 | 496 | GERGGPGSRGFP | GERGGP[113]GSRGFP[113] | GERGGP(1.0000)GSRGFP(1.0000) | 20.76 | 10.03 | 1.000 |
| 1212.548 | COL1A1 | 650 | 662 | GPPGEAGKPGEQG | GPP[113]GEAGKP[113]GEQG | GP(0.0210)P(0.9895)GEAGKP(0.9895)GEQG | 21.95 | 19.21 | 0.995 |
| 1215.614 | COL1A1 | 403 | 415 | PGIAGAPGFPGAR | P[113]GIAGAP[113]GFP[113]GAR | P(1.0000)GIAGAP(1.0000)GFP(1.0000)GAR | 28.74 | 16.62 | 1.000 |
| 1223.596 | COL1A1 | 913 | 925 | TGPAGRPGEVGPP | TGPAGRP[113]GEVGP[113]P | TGP(0.0101)AGRP(0.7601)GEVGP(0.6149)P(0.6149) | 22.03 | 22.03 | 0.983 |
| 1227.591 | COL1A1 | 606 | 619 | AVGPAGKDGEAGAQ | AVGPAGKDGEAGAQ | | 23.44 | 11.50 | 1.000 |
| 1229.584 | COL1A1 | 680 | 691 | GFPGERGVQGPP | GFP[113]GERGVQGPP[113] | GFP(0.9161)GERGVQGP(0.1697)P(0.9142) | 24.24 | 24.24 | 0.991 |
| 1230.582 | COL1A1 | 680 | 691 | GFPGERGVQGPP | GFP[113]GERGVQ[129]GP[113]P | GFP(0.7197)GERGVQGP(0.6402)P(0.6402) | 21.18 | 21.18 | 0.988 |
| 1232.547 | COL1A1 | 1187 | 1199 | GPPGPPSAGFDFS | GPPGPPSAGFDFS | | 27.11 | 11.83 | 0.968 |
| 1241.609 | COL1A1 | 395 | 409 | GAKGANGAPGIAGAP | GAKGAN[115]GAP[113]GIAGAP[113] | GAKGANGAP(1.0000)GIAGAP(1.0000) | 19.09 | 12.20 | 0.987 |
| 1242.576 | COL1A1 | 317 | 331 | GPAGARGNDGATGAA | GPAGARGNDGATGAA | | 20.61 | 12.22 | 0.997 |

| | | | | | | | | | |
|---|---|---|---|---|---|---|---|---|---|
| 1243.560 | COL1A1 | 317 | 331 | GPAGARGNDGATGAA | GPAGARGN[115]DGATGAA | | 18.47 | 11.43 | 0.982 |
| 1248.542 | COL1A1 | 1187 | 1199 | GPPGPPSAGFDFS | GPPGPP[113]SAGFDFS | GP(0.0079)P(0.0134)GP(0.3492)P(0.6294)SAGFDFS | 20.31 | 20.29 | 0.977 |
| 1251.650 | COL1A1 | 175 | 187 | ISVPGPMGPSGPR | ISVPGPMGPSGPR | | 18.83 | 9.23 | 0.995 |
| 1262.586 | COL1A1 | 270 | 283 | FSGLDGAKGDAGPA | FSGLDGAKGDAGPA | | 43.16 | 18.37 | 1.000 |
| 1264.539 | COL1A1 | 1187 | 1199 | GPPGPPSAGFDFS | GPP[113]GP[113]PSAGFDFS | GP(0.0134)P(0.9826)GP(0.9823)P(0.0217)SAGFDFS | 19.41 | 18.37 | 1.000 |
| 1267.582 | COL1A1 | 929 | 943 | GPAGEKGSPGADGPA | GPAGEKGSPGADGPA | | 29.11 | 12.24 | 1.000 |
| 1267.649 | COL1A1 | 175 | 187 | ISVPGPMGPSGPR | ISVPGPM[147]GPSGPR | ISVP(0.0028)GP(0.0049)M(0.9860)GP(0.0032)SGP(0.0031)R | 19.87 | 17.53 | 0.995 |
| 1267.716 | COL1A1 | 954 | 966 | IAGQRGVVGLPGQ | IAGQRGVVGLP[113]GQ | IAGQRGVVGLP(1.0000)GQ | 24.39 | 12.30 | 0.998 |
| 1272.533 | COL1A1 | 201 | 213 | FQGPPGEPGEPGA | FQ[129]GPPGEP[113]GEP[113]GA | FQGP(0.0120)P(0.0122)GEP(0.9879)GEP(0.9879)GA | 20.35 | 14.76 | 0.942 |
| 1278.595 | COL1A1 | 270 | 283 | FSGLDGAKGDAGPA | FSGLDGAKGDAGP[113]A | FSGLDGAKGDAGP(1.0000)A | 23.23 | 17.63 | 0.942 |
| 1280.533 | COL1A1 | 1187 | 1199 | GPPGPPSAGFDFS | GP[113]P[113]GPP[113]SAGFDFS | GP(0.9813)P(0.9813)GP(0.0561)P(0.9812)SAGFDFS | 25.32 | 21.53 | 1.000 |
| 1283.568 | COL1A1 | 932 | 946 | GEKGSPGADGPAGAP | GEKGSPGADGP[113]AGAP | GEKGSP(0.0080)GADGP(0.9864)AGAP(0.0056) | 20.05 | 15.79 | 0.991 |
| 1283.582 | COL1A1 | 773 | 787 | GPAGAPGDKGESGPS | GPAGAPGDKGESGPS | | 32.67 | 18.00 | 0.998 |
| 1284.607 | COL1A1 | 605 | 619 | GAVGPAGKDGEAGAQ | GAVGPAGKDGEAGAQ | | 32.77 | 13.40 | 1.000 |
| 1285.595 | COL1A1 | 605 | 619 | GAVGPAGKDGEAGAQ | GAVGPAGKDGEAGAQ[129] | | 23.49 | 15.19 | 0.961 |
| 1286.553 | COL1A1 | 701 | 715 | GAPGNDGAKGDAGAP | GAP[113]GNDGAKGDAGAP[113] | GAP(1.0000)GNDGAKGDAGAP(1.0000) | 18.39 | 11.69 | 0.998 |
| 1286.644 | COL1A1 | 405 | 418 | IAGAPGFPGARGPS | IAGAP[113]GFP[113]GARGPS | IAGAP(0.9826)GFP(0.9825)GARGP(0.0350)S | 18.82 | 18.82 | 0.977 |
| 1287.598 | COL1A1 | 680 | 692 | GFPGERGVQGPPG | GFP[113]GERGVQ[129]GPP[113]G | GFP(0.9682)GERGVQGP(0.0641)P(0.9678)G | 22.46 | 22.46 | 0.999 |
| 1299.575 | COL1A1 | 773 | 787 | GPAGAPGDKGESGPS | GPAGAP[113]GDKGESGPS | GP(0.0053)AGAP(0.9895)GDKGESGP(0.0052)S | 30.00 | 16.85 | 0.993 |
| 1299.575 | COL1A1 | 929 | 943 | GPAGEKGSPGADGPA | GP[113]AGEKGSP[113]GADGPA | GP(0.9852)AGEKGSP(0.9858)GADGP(0.0290)A | 21.02 | 12.88 | 0.980 |
| 1302.587 | COL1A1 | 398 | 412 | GANGAPGIAGAPGFP | GAN[115]GAP[113]GIAGAP[113]GFP[113] | GANGAP(1.0000)GIAGAP(1.0000)GFP(1.0000) | 26.39 | 14.87 | 1.000 |
| 1304.628 | COL1A1 | 677 | 688 | GERGFPGERGVQ | GERGFP[113]GERGVQ | GERGFP(1.0000)GERGVQ | 20.44 | 10.72 | 0.993 |
| 1311.652 | COL1A1 | 1145 | 1159 | GSAGAPGKDGLNGLP | GSAGAPGKDGLN[115]GLP | | 22.42 | 17.68 | 0.961 |
| 1313.551 | COL1A1 | 1121 | 1135 | GPPGSPGEQGPSGAS | GPP[113]GSP[113]GEQGPSGAS | GP(0.0322)P(0.9759)GSP(0.9764)GEQGP(0.0154)SGAS | 24.74 | 24.74 | 0.984 |
| 1313.597 | COL1A1 | 653 | 666 | GEAGKPGEQGVPGD | GEAGKP[113]GEQGVPGD | GEAGKP(0.9875)GEQGVP(0.0125)GD | 19.33 | 12.48 | 0.985 |
| 1315.572 | COL1A1 | 773 | 787 | GPAGAPGDKGESGPS | GPAGAP[113]GDKGESGP[113]S | GP(0.1205)AGAP(0.9403)GDKGESGP(0.9392)S | 18.21 | 10.59 | 0.936 |
| 1319.610 | COL1A1 | 269 | 283 | GFSGLDGAKGDAGPA | GFSGLDGAKGDAGPA | | 40.45 | 23.25 | 1.000 |
| 1322.639 | COL1A1 | 1196 | 1206 | FDFSFLPQPPQ | FDFSFLPQPPQ | | 19.53 | 12.89 | 0.958 |
| 1326.666 | COL1A1 | 1145 | 1159 | GSAGAPGKDGLNGLP | GSAGAPGKDGLNGLP[113] | GSAGAP(0.0189)GKDGLNGLP(0.9811) | 29.12 | 13.86 | 1.000 |
| 1327.640 | COL1A1 | 1145 | 1159 | GSAGAPGKDGLNGLP | GSAGAPGKDGLN[115]GLP[113] | GSAGAP(0.0103)GKDGLNGLP(0.9897) | 18.14 | 10.28 | 0.991 |
| 1333.609 | COL1A1 | 488 | 502 | GGPGSRGFPGADGVA | GGP[113]GSRGFP[113]GADGVA | GGP(1.0000)GSRGFP(1.0000)GADGVA | 19.71 | 12.18 | 0.997 |
| 1335.610 | COL1A1 | 269 | 283 | GFSGLDGAKGDAGPA | GFSGLDGAKGDAGP[113]A | GFSGLDGAKGDAGP(1.0000)A | 27.24 | 13.43 | 1.000 |
| 1342.623 | COL1A1 | 522 | 535 | SPGEAGRPGEAGLP | SP[113]GEAGRP[113]GEAGLP[113] | SP(1.0000)GEAGRP(1.0000)GEAGLP(1.0000) | 18.71 | 10.03 | 0.997 |
| 1342.651 | COL1A1 | 344 | 358 | GFPGAVGAKGEAGPQ | GFPGAVGAKGEAGPQ | | 28.20 | 14.26 | 0.998 |
| 1342.656 | COL1A1 | 1145 | 1159 | GSAGAPGKDGLNGLP | GSAGAP[113]GKDGLNGLP[113] | GSAGAP(1.0000)GKDGLNGLP(1.0000) | 18.29 | 12.75 | 0.937 |
| 1343.621 | COL1A1 | 1145 | 1159 | GSAGAPGKDGLNGLP | GSAGAP[113]GKDGLN[115]GLP[113] | GSAGAP(1.0000)GKDGLNGLP(1.0000) | 24.90 | 13.38 | 1.000 |
| 1343.656 | COL1A1 | 404 | 418 | GIAGAPGFPGARGPS | GIAGAP[113]GFP[113]GARGPS | GIAGAP(0.9632)GFP(0.9627)GARGP(0.0741)S | 28.40 | 22.55 | 0.994 |
| 1343.672 | COL1A1 | 401 | 415 | GAPGIAGAPGFPGAR | GAP[113]GIAGAP[113]GFP[113]GAR | GAP(1.0000)GIAGAP(1.0000)GFP(1.0000)GAR | 35.77 | 21.66 | 1.000 |
| 1350.633 | COL1A1 | 623 | 637 | GPAGPAGERGEQGPA | GPAGPAGERGEQGPA | | 18.09 | 18.09 | 0.959 |
| 1352.589 | COL1A1 | 203 | 217 | GPPGEPGEPGASGPM | GPPGEPGEPGASGP[113]M | GP(0.0033)P(0.0028)GEP(0.0029)GEP(0.0030)GASGP(0.4940)M(0.4940) | 30.09 | 30.09 | 1.000 |
| 1355.608 | COL1A1 | 542 | 556 | GSPGSPGPDGKTGPP | GSP[113]GSP[113]GPDGKTGPP[113] | GSP(0.9856)GSP(0.9855)GP(0.0239)DGKTGP(0.0194)P(0.9856) | 18.82 | 18.82 | 1.000 |
| 1358.577 | COL1A1 | 201 | 214 | FQGPPGEPGEPGAS | FQGPPGEP[113]GEP[113]GAS | FQGP(0.0099)P(0.0099)GEP(0.9901)GEP(0.9901)GAS | 30.04 | 20.84 | 1.000 |
| 1358.668 | COL1A1 | 344 | 358 | GFPGAVGAKGEAGPQ | GFPGAVGAKGEAGP[113]Q | GFP(0.0792)GAVGAKGEAGP(0.9208)Q | 19.96 | 11.71 | 0.998 |
| 1359.637 | COL1A1 | 404 | 418 | GIAGAPGFPGARGPS | GIAGAP[113]GFP[113]GARGP[113]S | GIAGAP(1.0000)GFP(1.0000)GARGP(1.0000)S | 28.48 | 11.76 | 1.000 |
| 1367.651 | COL1A1 | 1004 | 1018 | GLAGPPGESGREGAP | GLAGPPGESGREGAP[113] | GLAGP(0.0071)P(0.0087)GESGREGAP(0.9841) | 30.34 | 21.77 | 1.000 |
| 1368.568 | COL1A1 | 203 | 217 | GPPGEPGEPGASGPM | GPPGEP[113]GEPGASGP[113]M | GP(0.0052)P(0.0047)GEP(0.8117)GEP(0.1022)GASGP(0.5381)M(0.5381) | 29.53 | 29.53 | 0.994 |
| 1368.691 | COL1A1 | 587 | 601 | GAAGEPGKAGERGVP | GAAGEP[113]GKAGERGVP | GAAGEP(0.7636)GKAGERGVP(0.2364) | 18.51 | 12.11 | 0.997 |
| 1371.566 | COL1A1 | 197 | 210 | GPQGFQGPPGEPGE | GPQ[129]GFQ[129]GPPGEP[113]GE | GP(0.0040)QGFQGP(0.0062)P(0.0062)GEP(0.9837)GE | 18.92 | 18.27 | 0.990 |
| 1374.661 | COL1A1 | 344 | 358 | GFPGAVGAKGEAGPQ | GFP[113]GAVGAKGEAGP[113]Q | GFP(1.0000)GAVGAKGEAGP(1.0000)Q | 18.15 | 12.87 | 0.954 |
| 1381.703 | COL1A1 | 248 | 262 | GPQGARGLPGTAGLP | GPQ[129]GARGLP[113]GTAGLP[113] | GP(0.0222)QGARGLP(0.9889)GTAGLP(0.9889) | 21.23 | 13.71 | 0.982 |
| 1383.648 | COL1A1 | 1004 | 1018 | GLAGPPGESGREGAP | GLAGPP[113]GESGREGAP[113] | GLAGP(0.0314)P(0.9843)GESGREGAP(0.9843) | 23.78 | 19.42 | 0.992 |
| 1384.558 | COL1A1 | 203 | 217 | GPPGEPGEPGASGPM | GPPGEP[113]GEP[113]GASGP[113]M | GP(0.0101)P(0.0101)GEP(0.8273)GEP(0.8273)GASGP(0.6626)M(0.6626) | 27.79 | 27.79 | 0.980 |
| 1384.670 | COL1A1 | 587 | 601 | GAAGEPGKAGERGVP | GAAGEP[113]GKAGERGVP[113] | GAAGEP(1.0000)GKAGERGVP(1.0000) | 20.17 | 10.29 | 0.975 |
| 1385.675 | COL1A1 | 726 | 739 | LQGMPGERGAAGLP | LQGMP[113]GERGAAGLP[113] | LQGM(0.0235)P(0.9883)GERGAAGLP(0.9883) | 20.80 | 18.29 | 0.997 |
| 1386.691 | COL1A1 | 752 | 766 | GADGSPGKDGVRGLT | GADGSPGKDGVRGLT | | 21.06 | 8.51 | 1.000 |
| 1394.696 | COL1A1 | 656 | 670 | GKPGEQGVPGDLGAP | GKPGEQGVPGDLGAP[113] | GKP(0.0100)GEQGVP(0.0090)GDLGAP(0.9810) | 21.01 | 15.00 | 0.959 |
| 1397.706 | COL1A1 | 899 | 913 | GPAGKEGGKGPRGET | GPAGKEGGKGPRGET | | 20.06 | 11.89 | 0.989 |
| 1398.603 | COL1A1 | 284 | 298 | GPKGEPGSPGENGAP | GPKGEP[113]GSP[113]GENGAP[113] | GP(0.0321)KGEP(0.9893)GSP(0.9893)GENGAP(0.9893) | 19.69 | 13.98 | 0.985 |
| 1399.584 | COL1A1 | 284 | 298 | GPKGEPGSPGENGAP | GPKGEP[113]GSP[113]GEN[115]GAP[113] | GP(0.0322)KGEP(0.9893)GSP(0.9893)GENGAP(0.9893) | 25.40 | 19.13 | 0.981 |
| 1399.627 | COL1A1 | 521 | 535 | GSPGEAGRPGEAGLP | GSP[113]GEAGRP[113]GEAGLP[113] | GSP(1.0000)GEAGRP(1.0000)GEAGLP(1.0000) | 24.24 | 13.67 | 1.000 |
| 1400.551 | COL1A1 | 203 | 217 | GPPGEPGEPGASGPM | GP[113]P[113]GEP[113]GEP[113]GASGPM | GP(0.6402)P(0.6402)GEP(0.7197)GEP(0.7197)GASGP(0.6402)M(0.6402) | 25.98 | 25.98 | 0.934 |
| 1401.669 | COL1A1 | 726 | 739 | LQGMPGERGAAGLP | LQGM[147]P[113]GERGAAGLP[113] | LQGM(1.0000)P(1.0000)GERGAAGLP(1.0000) | 21.67 | 13.50 | 0.960 |
| 1402.684 | COL1A1 | 752 | 766 | GADGSPGKDGVRGLT | GADGSP[113]GKDGVRGLT | GADGSP(1.0000)GKDGVRGLT | 18.25 | 8.12 | 1.000 |
| 1409.655 | COL1A1 | 911 | 925 | GETGPAGRPGEVGPP | GETGPAGRP[113]GEVGPP[113] | GETGP(0.0104)AGRP(0.9872)GEVGP(0.0152)P(0.9872) | 22.43 | 22.43 | 0.938 |
| 1410.684 | COL1A1 | 656 | 670 | GKPGEQGVPGDLGAP | GKP[113]GEQGVPGDLGAP[113] | GKP(0.9879)GEQGVP(0.0242)GDLGAP(0.9879) | 19.77 | 13.86 | 0.969 |
| 1411.669 | COL1A1 | 656 | 670 | GKPGEQGVPGDLGAP | GKP[113]GEQ[129]GVPGDLGAP[113] | GKP(0.9867)GEQGVP(0.0267)GDLGAP(0.9865) | 18.82 | 12.83 | 0.977 |
| 1412.623 | COL1A1 | 224 | 238 | GPPGKNGDDGEAGKP | GPP[113]GKN[115]GDDGEAGKP | GP(0.0051)P(0.9860)GKNGDDGEAGKP(0.0089) | 29.10 | 25.84 | 1.000 |
| 1414.605 | COL1A1 | 284 | 298 | GPKGEPGSPGENGAP | GP[113]KGEP[113]GSP[113]GENGAP[113] | GP(1.0000)KGEP(1.0000)GSP(1.0000)GENGAP(1.0000) | 21.71 | 11.24 | 0.997 |
| 1415.584 | COL1A1 | 284 | 298 | GPKGEPGSPGENGAP | GP[113]KGEP[113]GSP[113]GEN[115]GAP[113] | GP(1.0000)KGEP(1.0000)GSP(1.0000)GENGAP(1.0000) | 26.64 | 10.96 | 1.000 |
| 1415.610 | COL1A1 | 200 | 214 | GFQGPPGEPGEPGAS | GFQGPPGEP[113]GEP[113]GAS | GFQGP(0.0127)P(0.0129)GEP(0.9872)GEP(0.9872)GAS | 20.23 | 13.44 | 1.000 |
| 1421.779 | COL1A1 | 950 | 964 | GPQGIAGQRGVVGLP | GPQGIAGQRGVVGLP[113] | GP(0.0103)QGIAGQRGVVGLP(0.9897) | 46.36 | 24.00 | 1.000 |
| 1422.765 | COL1A1 | 950 | 964 | GPQGIAGQRGVVGLP | GPQ[129]GIAGQRGVVGLP[113] | GP(0.0132)QGIAGQRGVVGLP(0.9868) | 19.81 | 13.63 | 0.996 |
| 1423.753 | COL1A1 | 950 | 964 | GPQGIAGQRGVVGLP | GPQ[129]GIAGQ[129]RGVVGLP[113] | GP(0.0161)QGIAGQRGVVGLP(0.9839) | 25.87 | 11.79 | 1.000 |
| 1424.656 | COL1A1 | 607 | 622 | VGPAGKDGEAGAQGPP | VGPAGKDGEAGAQ[129]GP[113]P | VGP(0.0042)AGKDGEAGAQGP(0.4979)P(0.4979) | 23.94 | 23.94 | 0.939 |
| 1424.670 | COL1A1 | 650 | 664 | GPPGEAGKPGEQGVP | GPP[113]GEAGKP[113]GEQGVP[113] | GP(0.1147)P(0.9615)GEAGKP(0.9619)GEQGVP(0.9619) | 18.47 | 16.73 | 0.986 |
| 1424.793 | COL1A1 | 954 | 967 | IAGQRGVVGLPGQR | IAGQRGVVGLP[113]GQ[129]R | IAGQRGVVGLP(1.0000)GQR | 21.52 | 9.46 | 1.000 |
| 1426.672 | COL1A1 | 656 | 670 | GKPGEQGVPGDLGAP | GKP[113]GEQGVP[113]GDLGAP[113] | GKP(1.0000)GEQGVP(1.0000)GDLGAP(1.0000) | 36.08 | 19.13 | 0.979 |
| 1427.663 | COL1A1 | 656 | 670 | GKPGEQGVPGDLGAP | GKP[113]GEQ[129]GVP[113]GDLGAP[113] | GKP(1.0000)GEQGVP(1.0000)GDLGAP(1.0000) | 19.42 | 14.46 | 0.949 |
| 1428.622 | COL1A1 | 224 | 238 | GPPGKNGDDGEAGKP | GPP[113]GKN[115]GDDGEAGKP[113] | GP(0.0222)P(0.9889)GKNGDDGEAGKP(0.9889) | 22.57 | 19.15 | 1.000 |
| 1430.685 | COL1A1 | 569 | 583 | GPPGARGQAGVMGFP | GPP[113]GARGQAGVMGFP[113] | GP(0.0325)P(0.9717)GARGQAGVM(0.0232)GFP(0.9725) | 18.62 | 16.84 | 0.994 |
| 1431.646 | COL1A1 | 632 | 646 | GEQGPAGSPGFQGLP | GEQGPAGSP[113]GFQ[129]GLP[113] | GEQGP(0.0335)AGSP(0.9832)GFQGLP(0.9833) | 22.90 | 18.67 | 1.000 |
| 1432.631 | COL1A1 | 632 | 646 | GEQGPAGSPGFQGLP | GEQ[129]GPAGSP[113]GFQ[129]GLP[113] | GEQGP(0.0408)AGSP(0.9795)GFQGLP(0.9797) | 18.31 | 14.59 | 1.000 |
| 1442.701 | COL1A1 | 725 | 739 | GLQGMPGERGAAGLP | GLQGM[147]PGERGAAGLP[113] | GLQGM(0.9850)P(0.0300)GERGAAGLP(0.9850) | 18.22 | 13.88 | 0.923 |

| | | | | | | | | | |
|---|---|---|---|---|---|---|---|---|---|
| 1443.684 | COL1A1 | 725 | 739 | GLQGMPGERGAAGLP | GLQ[129]GMP[113]GERGAAGLP[113] | GLQGM(0.0234)P(0.9883)GERGAAGLP(0.9883) | 20.64 | 15.69 | 0.986 |
| 1446.671 | COL1A1 | 569 | 583 | GPPGARGQAGVMGFP | GPP[113]GARGQAGVM[147]GFP[113] | GP(0.0662)P(0.9779)GARGQAGVM(0.9779)GFP(0.9780) | 19.88 | 17.93 | 0.981 |
| 1447.676 | COL1A1 | 569 | 583 | GPPGARGQAGVMGFP | GPP[113]GARGQ[129]AGVM[147]GFP[113] | GP(0.0727)P(0.9756)GARGQAGVM(0.9758)GFP(0.9758) | 18.63 | 16.69 | 0.993 |
| 1454.707 | COL1A1 | 749 | 764 | GPKGADGSPGKDGVRG | GPKGADGSPGKDGVRG | | 26.13 | 11.19 | 1.000 |
| 1458.685 | COL1A1 | 725 | 739 | GLQGMPGERGAAGLP | GLQGM[147]P[113]GERGAAGLP[113] | GLQGM(1.0000)P(1.0000)GERGAAGLP(1.0000) | 43.01 | 17.59 | 1.000 |
| 1459.678 | COL1A1 | 725 | 739 | GLQGMPGERGAAGLP | GLQ[129]GM[147]P[113]GERGAAGLP[113] | GLQGM(1.0000)P(1.0000)GERGAAGLP(1.0000) | 22.78 | 13.48 | 1.000 |
| 1465.688 | COL1A1 | 650 | 665 | GPPGEAGKPGEQGVPG | GPP[113]GEAGKP[113]GEQGVPG | GP(0.0143)P(0.9874)GEAGKP(0.9876)GEQGVP(0.0107)G | 19.61 | 17.77 | 0.978 |
| 1466.686 | COL1A1 | 1190 | 1203 | GPPSAGFDFSFLPQ | GPPSAGFDFSFLPQ | | 20.17 | 13.90 | 0.955 |
| 1468.692 | COL1A1 | 656 | 671 | GKPGEQGVPGDLGAPG | GKP[113]GEQ[129]GVPGDLGAP[113]G | GKP(0.9899)GEQGVP(0.0206)GDLGAP(0.9895)G | 20.93 | 15.35 | 0.960 |
| 1470.724 | COL1A1 | 749 | 764 | GPKGADGSPGKDGVRG | GPKGADGSP[113]GKDGVRG | GP(0.0114)KGADGSP(0.9886)GKDGVRG | 25.59 | 12.59 | 1.000 |
| 1475.714 | COL1A1 | 268 | 283 | RGFSGLDGAKGDAGPA | RGFSGLDGAKGDAGPA | | 26.45 | 11.49 | 1.000 |
| 1481.694 | COL1A1 | 650 | 665 | GPPGEAGKPGEQGVPG | GPP[113]GEAGKP[113]GEQGVP[113]G | GP(0.0351)P(0.9883)GEAGKP(0.9883)GEQGVP(0.9883)G | 18.68 | 16.84 | 0.968 |
| 1482.708 | COL1A1 | 655 | 670 | AGKPGEQGVPGDLGAP | AGKP[113]GEQ[129]GVPGDLGAP[113] | AGKP(0.9884)GEQGVP(0.0232)GDLGAP(0.9884) | 19.87 | 15.44 | 0.978 |
| 1498.703 | COL1A1 | 655 | 670 | AGKPGEQGVPGDLGAP | AGKP[113]GEQ[129]GVP[113]GDLGAP[113] | AGKP(1.0000)GEQGVP(1.0000)GDLGAP(1.0000) | 18.25 | 11.99 | 0.984 |
| 1529.627 | COL1A1 | 698 | 715 | GANGAPGNDGAKGDAGAP | GAN[115]GAP[113]GNDGAKGDAGAP[113] | GANGAP(1.0000)GNDGAKGDAGAP(1.0000) | 27.22 | 14.44 | 1.000 |
| 1535.731 | COL1A1 | 605 | 622 | GAVGPAGKDGEAGAQGPP | GAVGPAGKDGEAGAQGPP | | 25.58 | 14.11 | 0.971 |
| 1536.730 | COL1A1 | 605 | 622 | GAVGPAGKDGEAGAQGPP | GAVGPAGKDGEAGAQ[129]GPP | | 18.78 | 13.85 | 0.975 |
| 1538.665 | COL1A1 | 1121 | 1138 | GPPGSPGEQGPSGASGPA | GP[113]PGSP[113]GEQGPSGASGPA | GP(0.5874)P(0.5874)GSP(0.8113)GEQGP(0.0065)SGASGP(0.0074)A | 30.35 | 30.35 | 1.000 |
| 1550.799 | COL1A1 | 175 | 190 | ISVPGPMGPSGPRGLP | ISVPGPM[147]GPSGPRGLP[113] | ISVP(0.0100)GP(0.1433)M(0.8944)GP(0.0137)SGP(0.0139)RGLP(0.9246) | 21.36 | 21.33 | 1.000 |
| 1551.724 | COL1A1 | 602 | 619 | GPPGAVGPAGKDGEAGAQ | GPP[113]GAVGPAGKDGEAGAQ | GP(0.0100)P(0.9840)GAVGP(0.0060)AGKDGEAGAQ | 21.93 | 20.03 | 0.965 |
| 1552.708 | COL1A1 | 605 | 622 | GAVGPAGKDGEAGAQGPP | GAVGPAGKDGEAGAQ[129]GPP[113] | GAVGP(0.0074)AGKDGEAGAQGP(0.0084)P(0.9842) | 32.46 | 32.46 | 0.997 |
| 1552.818 | COL1A1 | 851 | 868 | GPIGNVGAPGAKGARGSA | GPIGNVGAP[113]GAKGARGSA | GP(0.0138)IGNVGAP(0.9862)GAKGARGSA | 27.01 | 11.34 | 1.000 |
| 1553.721 | COL1A1 | 167 | 181 | YDEKSTGGISVPGPM | YDEKSTGGISVPGP[113]M | YDEKSTGGISVP(0.0056)GP(0.4972)M(0.4972) | 26.59 | 26.59 | 1.000 |
| 1557.757 | COL1A1 | 395 | 412 | GAKGANGAPGIAGAPGFP | GAKGANGAP[113]GIAGAP[113]GFP[113] | GAKGANGAP(1.0000)GIAGAP(1.0000)GFP(1.0000) | 18.12 | 9.71 | 0.993 |
| 1558.719 | COL1A1 | 1091 | 1105 | GPRGDKGETGEQGDR | GPRGDKGETGEQGDR | | 19.05 | 8.71 | 0.999 |
| 1558.739 | COL1A1 | 395 | 412 | GAKGANGAPGIAGAPGFP | GAKGAN[115]GAP[113]GIAGAP[113]GFP[113] | GAKGANGAP(1.0000)GIAGAP(1.0000)GFP(1.0000) | 24.68 | 10.81 | 1.000 |
| 1562.774 | COL1A1 | 1142 | 1159 | GPPGSAGAPGKDGLNGLP | GPPGSAGAPGKDGLN[115]GLP | | 22.73 | 11.98 | 0.998 |
| 1567.724 | COL1A1 | 824 | 841 | GAKGEPGDAGAKGDAGPP | GAKGEPGDAGAKGDAGP[113]P | GAKGEP(0.0044)GDAGAKGDAGP(0.4978)P(0.4978) | 21.09 | 21.09 | 0.977 |
| 1569.725 | COL1A1 | 540 | 556 | LTGSPGSPGPDGKTGPP | LTGSP[113]GSP[113]GPDGKTGPP[113] | LTGSP(0.9620)GSP(0.9605)GP(0.0961)DGKTGP(0.0194)P(0.9620) | 30.77 | 30.77 | 0.998 |
| 1572.741 | COL1A1 | 677 | 691 | GERGFPGERGVQGPP | GERGFP[113]GERGVQ[129]GPP[113] | GERGFP(0.9864)GERGVQGP(0.0273)P(0.9863) | 20.41 | 18.45 | 0.951 |
| 1574.710 | COL1A1 | 1091 | 1105 | GPRGDKGETGEQGDR | GP[113]RGDKGETGEQGDR | GP(1.0000)RGDKGETGEQGDR | 21.04 | 12.12 | 0.998 |
| 1578.770 | COL1A1 | 1142 | 1159 | GPPGSAGAPGKDGLNGLP | GPPGSAGAPGKDGLN[115]GLP[113] | GP(0.0074)P(0.0077)GSAGAP(0.0130)GKDGLNGLP(0.9718) | 24.33 | 16.57 | 0.941 |
| 1580.708 | COL1A1 | 650 | 666 | GPPGEAGKPGEQGVPGD | GPP[113]GEAGKP[113]GEQGVPGD | GP(0.0144)P(0.9866)GEAGKP(0.9867)GEQGVP(0.0123)GD | 22.54 | 20.55 | 1.000 |
| 1581.699 | COL1A1 | 650 | 666 | GPPGEAGKPGEQGVPGD | GPP[113]GEAGKP[113]GEQ[129]GVPGD | GP(0.0221)P(0.9823)GEAGKP(0.9826)GEQGVP(0.0130)GD | 22.08 | 20.10 | 1.000 |
| 1583.719 | COL1A1 | 824 | 841 | GAKGEPGDAGAKGDAGPP | GAKGEP[113]GDAGAKGDAGPP[113] | GAKGEP(0.9556)GDAGAKGDAGP(0.0896)P(0.9548) | 21.51 | 21.51 | 0.960 |
| 1584.766 | COL1A1 | 401 | 418 | GAPGIAGAPGFPGARGPS | GAP[113]GIAGAP[113]GFP[113]GARGPS | GAP(0.9484)GIAGAP(0.9484)GFP(0.9478)GARGP(0.1553)S | 22.73 | 20.76 | 0.990 |
| 1586.744 | COL1A1 | 398 | 415 | GANGAPGIAGAPGFPGAR | GAN[115]GAP[113]GIAGAP[113]GFP[113]GAR | GANGAP(1.0000)GIAGAP(1.0000)GFP(1.0000)GAR | 30.09 | 17.40 | 1.000 |
| 1588.794 | COL1A1 | 674 | 688 | GARGERGFPGERGVQ | GARGERGFP[113]GERGVQ | GARGERGFP(1.0000)GERGVQ | 18.24 | 11.69 | 0.949 |
| 1593.776 | COL1A1 | 1142 | 1159 | GPPGSAGAPGKDGLNGLP | GPP[113]GSAGAPGKDGLNGLP[113] | GP(0.0394)P(0.9730)GSAGAP(0.0136)GKDGLNGLP(0.9740) | 35.86 | 33.69 | 1.000 |
| 1594.765 | COL1A1 | 1142 | 1159 | GPPGSAGAPGKDGLNGLP | GPP[113]GSAGAPGKDGLN[115]GLP[113] | GP(0.0138)P(0.9843)GSAGAP(0.0178)GKDGLNGLP(0.9841) | 45.88 | 43.38 | 1.000 |
| 1596.707 | COL1A1 | 650 | 666 | GPPGEAGKPGEQGVPGD | GP[113]PGEAGKP[113]GEQGVP[113]GD | GP(0.7216)P(0.7216)GEAGKP(0.7784)GEQGVP(0.7784)GD | 24.07 | 24.07 | 1.000 |
| 1600.761 | COL1A1 | 401 | 418 | GAPGIAGAPGFPGARGPS | GAP[113]GIAGAP[113]GFP[113]GARGP[113]S | GAP(1.0000)GIAGAP(1.0000)GFP(1.0000)GARGP(1.0000)S | 24.07 | 12.84 | 0.996 |
| 1609.768 | COL1A1 | 1142 | 1159 | GPPGSAGAPGKDGLNGLP | GPP[113]GSAGAP[113]GKDGLNGLP[113] | GP(0.0371)P(0.9876)GSAGAP(0.9876)GKDGLNGLP(0.9876) | 19.85 | 17.96 | 0.999 |
| 1609.781 | COL1A1 | 1055 | 1072 | GPVGPAGKSGDRGETGPA | GPVGPAGKSGDRGETGPA | | 20.09 | 12.66 | 1.000 |
| 1609.791 | COL1A1 | 341 | 358 | GPPGFPGAVGAKGEAGPQ | GPP[113]GFPGAVGAKGEAGPQ | GP(0.0069)P(0.9840)GFP(0.0039)GAVGAKGEAGP(0.0052)Q | 22.34 | 20.47 | 0.980 |
| 1610.762 | COL1A1 | 1142 | 1159 | GPPGSAGAPGKDGLNGLP | GPP[113]GSAGAP[113]GKDGLN[115]GLP[113] | GP(0.0524)P(0.9825)GSAGAP(0.9826)GKDGLNGLP(0.9826) | 20.05 | 20.05 | 0.995 |
| 1612.672 | COL1A1 | 197 | 213 | GPQGFQGPPGEPGEPGA | GPQ[129]GFQ[129]GPPGEP[113]GEP[113]GA | GP(0.0109)QGFQGP(0.0092)P(0.0097)GEP(0.9852)GEP(0.9851)GA | 32.35 | 23.63 | 0.997 |
| 1615.750 | COL1A1 | 1091 | 1106 | GPRGDKGETGEQGDRG | GPRGDKGETGEQGDRG | | 25.67 | 8.26 | 1.000 |
| 1622.820 | COL1A1 | 899 | 916 | GPAGKEGGKGPRGETGPA | GPAGKEGGKGPRGETGPA | | 19.44 | 10.68 | 1.000 |
| 1625.783 | COL1A1 | 341 | 358 | GPPGFPGAVGAKGEAGPQ | GPP[113]GFP[113]GAVGAKGEAGPQ | GP(0.0194)P(0.9786)GFP(0.9785)GAVGAKGEAGP(0.0235)Q | 36.40 | 36.40 | 1.000 |
| 1628.661 | COL1A1 | 197 | 213 | GPQGFQGPPGEPGEPGA | GPQ[129]GFQ[129]GPP[113]GEP[113]GEP[113]GA | GP(0.0163)QGFQGP(0.0214)P(0.9874)GEP(0.9875)GEP(0.9875)GA | 22.90 | 21.01 | 0.992 |
| 1631.729 | COL1A1 | 1091 | 1106 | GPRGDKGETGEQGDRG | GP[113]RGDKGETGEQGDRG | GP(1.0000)RGDKGETGEQGDRG | 29.89 | 10.39 | 1.000 |
| 1634.771 | COL1A1 | 1001 | 1018 | GPPGLAGPPGESGREGAP | GPPGLAGPP[113]GESGREGAP[113] | GP(0.0070)P(0.0070)GLAGP(0.0087)P(0.9890)GESGREGAP(0.9883) | 23.92 | 19.92 | 0.995 |
| 1641.783 | COL1A1 | 341 | 358 | GPPGFPGAVGAKGEAGPQ | GPP[113]GFP[113]GAVGAKGEAGP[113]Q | GP(0.0538)P(0.9824)GFP(0.9825)GAVGAKGEAGP(0.9814)Q | 18.60 | 17.12 | 0.998 |
| 1643.695 | COL1A1 | 201 | 217 | FQGPPGEPGEPGASGPM | FQGPPGEP[113]GEP[113]GASGPM | FQGP(0.0059)P(0.0060)GEP(0.9878)GEP(0.9877)GASGP(0.0063)M(0.0063) | 32.02 | 22.99 | 1.000 |
| 1650.845 | COL1A1 | 584 | 601 | GPKGAAGEPGKAGERGVP | GPKGAAGEP[113]GKAGERGVP | GP(0.0091)KGAAGEP(0.9825)GKAGERGVP(0.0084) | 26.56 | 14.75 | 1.000 |
| 1651.792 | COL1A1 | 653 | 670 | GEAGKPGEQGVPGDLGAP | GEAGKPGEQGVPGDLGAP[113] | GEAGKP(0.0058)GEQGVP(0.0056)GDLGAP(0.9886) | 22.92 | 15.81 | 0.954 |
| 1656.811 | COL1A1 | 440 | 457 | GSKGDTGAKGEPGPVGVQ | GSKGDTGAKGEP[113]GPVGVQ | GSKGDTGAKGEP(0.9890)GP(0.0110)VGVQ | 24.09 | 18.53 | 1.000 |
| 1659.688 | COL1A1 | 201 | 217 | FQGPPGEPGEPGASGPM | FQGPPGEP[113]GEP[113]GASGP[113]M | FQGP(0.0101)P(0.0101)GEP(0.8241)GEP(0.8226)GASGP(0.6666)M(0.6666) | 19.97 | 19.97 | 0.982 |
| 1660.675 | COL1A1 | 201 | 217 | FQGPPGEPGEPGASGPM | FQ[129]GPPGEP[113]GEP[113]GASGP[113]M | FQGP(0.0095)P(0.0095)GEP(0.8292)GEP(0.8292)GASGP(0.6613)M(0.6613) | 31.43 | 31.43 | 1.000 |
| 1666.835 | COL1A1 | 584 | 601 | GPKGAAGEPGKAGERGVP | GPKGAAGEP[113]GKAGERGVP[113] | GP(0.0359)KGAAGEP(0.9820)GKAGERGVP(0.9821) | 27.40 | 17.07 | 1.000 |
| 1667.777 | COL1A1 | 653 | 670 | GEAGKPGEQGVPGDLGAP | GEAGKPGEQGVP[113]GDLGAP[113] | GEAGKP(0.0368)GEQGVP(0.9816)GDLGAP(0.9816) | 25.02 | 14.59 | 0.986 |
| 1668.762 | COL1A1 | 653 | 670 | GEAGKPGEQGVPGDLGAP | GEAGKP[113]GEQ[129]GVPGDLGAP[113] | GEAGKP(0.9800)GEQGVP(0.0400)GDLGAP(0.9800) | 23.61 | 17.91 | 1.000 |
| 1668.848 | COL1A1 | 749 | 766 | GPKGADGSPGKDGVRGLT | GPKGADGSPGKDGVRGLT | | 18.65 | 12.16 | 1.000 |
| 1669.779 | COL1A1 | 266 | 283 | GHRGFSGLDGAKGDAGPA | GHRGFSGLDGAKGDAGPA | | 19.71 | 11.46 | 1.000 |
| 1675.671 | COL1A1 | 201 | 217 | FQGPPGEPGEPGASGPM | FQGPP[113]GEP[113]GEP[113]GASGPM[147] | FQGP(0.0882)P(0.8731)GEP(0.8738)GEP(0.8738)GASGP(0.4279)M(0.8632) | 34.16 | 34.16 | 0.997 |
| 1676.677 | COL1A1 | 201 | 217 | FQGPPGEPGEPGASGPM | FQ[129]GP[113]P[113]GEP[113]GEP[113]GASGPM | FQGP(0.6414)P(0.6414)GEP(0.7172)GEP(0.7171)GASGP(0.6414)M(0.6414) | 20.28 | 20.28 | 0.999 |
| 1676.860 | COL1A1 | 1200 | 1214 | FLPQPPQEKAHDGGR | FLPQPPQEKAHDGGR | | 19.49 | 10.34 | 1.000 |
| 1681.799 | COL1A1 | 518 | 535 | GPKGSPGEAGRPGEAGLP | GPKGSP[113]GEAGRP[113]GEAGLP[113] | GP(0.0377)KGSP(0.9874)GEAGRP(0.9874)GEAGLP(0.9874) | 21.26 | 15.67 | 0.983 |
| 1682.828 | COL1A1 | 584 | 601 | GPKGAAGEPGKAGERGVP | GP[113]KGAAGEP[113]GKAGERGVP[113] | GP(1.0000)KGAAGEP(1.0000)GKAGERGVP(1.0000) | 29.71 | 16.62 | 1.000 |
| 1683.767 | COL1A1 | 653 | 670 | GEAGKPGEQGVPGDLGAP | GEAGKP[113]GEQGVP[113]GDLGAP[113] | GEAGKP(1.0000)GEQGVP(1.0000)GDLGAP(1.0000) | 25.92 | 14.15 | 0.992 |
| 1684.759 | COL1A1 | 653 | 670 | GEAGKPGEQGVPGDLGAP | GEAGKP[113]GEQ[129]GVP[113]GDLGAP[113] | GEAGKP(1.0000)GEQGVP(1.0000)GDLGAP(1.0000) | 28.13 | 14.89 | 1.000 |
| 1684.846 | COL1A1 | 749 | 766 | GPKGADGSPGKDGVRGLT | GPKGADGSP[113]GKDGVRGLT | GP(0.0297)KGADGSP(0.9703)GKDGVRGLT | 25.71 | 12.94 | 1.000 |
| 1697.726 | COL1A1 | 197 | 214 | GPQGFQGPPGEPGEPGAS | GPQGFQGPPGEP[113]GEP[113]GAS | GP(0.0100)QGFQGP(0.0071)P(0.0071)GEP(0.9879)GEP(0.9878)GAS | 22.26 | 14.40 | 0.989 |
| 1698.708 | COL1A1 | 197 | 214 | GPQGFQGPPGEPGEPGAS | GPQGFQ[129]GPPGEP[113]GEP[113]GAS | GP(0.0111)QGFQGP(0.0078)P(0.0080)GEP(0.9865)GEP(0.9865)GAS | 31.49 | 25.30 | 0.999 |
| 1699.702 | COL1A1 | 197 | 214 | GPQGFQGPPGEPGEPGAS | GPQ[129]GFQ[129]GPPGEP[113]GEP[113]GAS | GP(0.0089)QGFQGP(0.0071)P(0.0071)GEP(0.9884)GEP(0.9885)GAS | 25.75 | 17.05 | 1.000 |
| 1699.799 | COL1A1 | 722 | 739 | GAPGLQGMPGERGAAGLP | GAP[113]GLQGM[147]P[113]GERGAAGLP[113] | GAP(1.0000)GLQGM(1.0000)P(1.0000)GERGAAGLP(1.0000) | 24.04 | 13.88 | 0.999 |
| 1700.790 | COL1A1 | 722 | 739 | GAPGLQGMPGERGAAGLP | GAP[113]GLQ[129]GM[147]P[113]GERGAAGLP[113] | GAP(1.0000)GLQGM(1.0000)P(1.0000)GERGAAGLP(1.0000) | 22.51 | 11.38 | 1.000 |
| 1700.797 | COL1A1 | 1025 | 1042 | GRDGSPGAKGDRGETGPA | GRDGSP[113]GAKGDRGETGPA | GRDGSP(0.9868)GAKGDRGETGP(0.0132)A | 19.44 | 7.74 | 1.000 |
| 1713.734 | COL1A1 | 197 | 214 | GPQGFQGPPGEPGEPGAS | GPQGFQGPP[113]GEP[113]GEP[113]GAS | GP(0.0275)QGFQGP(0.4284)P(0.8363)GEP(0.8539)GEP(0.8539)GAS | 22.11 | 20.27 | 0.997 |
| 1716.711 | COL1A1 | 200 | 217 | GFQGPPGEPGEPGASGPM | GFQGPPGEP[113]GEP[113]GASGP[113]M | GFQGP(0.0091)P(0.0091)GEP(0.8275)GEP(0.8275)GASGP(0.6634)M(0.6634) | 36.31 | 36.31 | 1.000 |
| 1719.833 | COL1A1 | 908 | 925 | GPRGETGPAGRPGEVGPP | GPRGETGPAGRP[113]GEVGPP[113] | GP(0.0142)RGETGP(0.0190)AGRP(0.9732)GEVGP(0.0204)P(0.9731) | 21.63 | 21.63 | 0.969 |
| 1725.770 | COL1A1 | 653 | 671 | GEAGKPGEQGVPGDLGAPG | GEAGKP[113]GEQ[129]GVPGDLGAP[113]G | GEAGKP(0.9801)GEQGVP(0.0400)GDLGAP(0.9799)G | 22.02 | 11.63 | 1.000 |

### Supplemental Table 1. Col1a1 peptide sequences from normal breast by LC-MS/MS.

| | | | | | | | | | |
|---|---|---|---|---|---|---|---|---|---|
| 1731.710 | COL1A1 | 284 | 301 | GPKGEPGSPGENGAPGQM | GPKGEP[113]GSP[113]GEN[115]GAP[113]GQM[147] | GP(0.0635)KGEP(0.9841)GSP(0.9841)GENGAP(0.9841)GQM(0.9841) | 19.33 | 15.49 | 0.947 |
| 1738.846 | COL1A1 | 461 | 478 | GPAGEEGKRGARGEPGPT | GPAGEEGKRGARGEP[113]GPT | GP(0.0061)AGEEGKRGARGEP(0.9885)GP(0.0054)T | 18.10 | 14.02 | 0.998 |
| 1742.834 | COL1A1 | 1202 | 1216 | PQPPQEKAHDGGRYY | PQPPQEKAHDGGRYY | | 20.23 | 11.26 | 1.000 |
| 1762.846 | COL1A1 | 359 | 376 | GPRGSEGPQGVRGEPGPP | GPRGSEGPQGVRGEP[113]GPP[113] | GP(0.0070)RGSEGP(0.0070)QGVRGEP(0.9890)GP(0.0080)P(0.9890) | 18.66 | 18.66 | 1.000 |
| 1768.931 | COL1A1 | 960 | 976 | VVGLPGQRGERGFPGLP | VVGLP[113]GQ[129]RGERGFPGLP[113] | VVGLP(0.9807)GQRGERGFP(0.0385)GLP(0.9809) | 18.91 | 15.64 | 0.987 |
| 1774.781 | COL1A1 | 629 | 646 | GERGEQGPAGSPGFQGLP | GERGEQ[129]GPAGSP[113]GFQ[129]GLP[113] | GERGEQGP(0.0295)AGSP(0.9853)GFQGLP(0.9853) | 21.35 | 16.61 | 1.000 |
| 1788.837 | COL1A1 | 1190 | 1206 | GPPSAGFDFSFLPQPPQ | GPPSAGFDFSFLPQPPQ | | 21.37 | 11.43 | 0.942 |
| 1802.857 | COL1A1 | 602 | 622 | GPPGAVGPAGKDGEAGAQGPP | GPPGAVGPAGKDGEAGAQGPP[113] | GP(0.0028)P(0.0028)GAVGP(0.0025)AGKDGEAGAQGP(0.0038)P(0.9880) | 20.22 | 20.22 | 0.990 |
| 1818.847 | COL1A1 | 602 | 622 | GPPGAVGPAGKDGEAGAQGPP | GP[113]P[113]GAVGPAGKDGEAGAQGPP | GP(0.4987)P(0.4987)GAVGP(0.0052)AGKDGEAGAQGP(0.4987)P(0.4987) | 18.11 | 18.11 | 0.979 |
| 1819.945 | COL1A1 | 851 | 871 | GPIGNVGAPGAKGARGSAGPP | GPIGNVGAP[113]GAKGARGSAGP[113]P | GP(0.0109)IGNVGAP(0.7639)GAKGARGSAGP(0.6126)P(0.6126) | 19.57 | 19.57 | 0.981 |
| 1827.859 | COL1A1 | 398 | 418 | GANGAPGIAGAPGFPGARGPS | GAN[115]GAP[113]GIAGAP[113]GFP[113]GARGPS | GANGAP(0.9088)GIAGAP(0.9087)GFP(0.9072)GARGP(0.2753)S | 24.55 | 24.55 | 0.999 |
| 1839.807 | COL1A1 | 695 | 715 | GPRGANGAPGNDGAKGDAGAP | GPRGAN[115]GAP[113]GNDGAKGDAGAP[113] | GP(0.0398)RGANGAP(0.9801)GNDGAKGDAGAP(0.9801) | 27.31 | 24.09 | 1.000 |
| 1840.783 | COL1A1 | 695 | 715 | GPRGANGAPGNDGAKGDAGAP | GPRGAN[115]GAP[113]GN[115]DGAKGDAGAP[113] | GP(0.0288)RGANGAP(0.9856)GNDGAKGDAGAP(0.9856) | 20.20 | 12.38 | 0.998 |
| 1841.893 | COL1A1 | 395 | 415 | GAKGANGAPGIAGAPGFPGAR | GAKGANGAP[113]GIAGAP[113]GFP[113]GAR | GAKGANGAP(1.0000)GIAGAP(1.0000)GFP(1.0000)GAR | 23.65 | 10.58 | 1.000 |
| 1842.889 | COL1A1 | 395 | 415 | GAKGANGAPGIAGAPGFPGAR | GAKGAN[115]GAP[113]GIAGAP[113]GFP[113]GAR | GAKGANGAP(1.0000)GIAGAP(1.0000)GFP(1.0000)GAR | 33.31 | 20.48 | 1.000 |
| 1876.930 | COL1A1 | 1145 | 1165 | GSAGAPGKDGLNGLPGPIGPP | GSAGAPGKDGLNGLP[113]GPIGP[113]P[113] | GSAGAP(0.0169)GKDGLNGLP(0.9668)GP(0.0856)IGP(0.9654)P(0.9654) | 30.16 | 24.10 | 0.982 |
| 1877.879 | COL1A1 | 1145 | 1165 | GSAGAPGKDGLNGLPGPIGPP | GSAGAPGKDGLN[115]GLP[113]GPIGP[113]P[113] | GSAGAP(0.0425)GKDGLNGLP(0.9628)GP(0.0676)IGP(0.9638)P(0.9633) | 18.97 | 17.12 | 0.998 |
| 1882.886 | COL1A1 | 536 | 556 | GAKGLTGSPGSPGPDGKTGPP | GAKGLTGSP[113]GSP[113]GPDGKTGPP[113] | GAKGLTGSP(0.9824)GSP(0.9824)GP(0.0228)DGKTGP(0.0300)P(0.9822) | 28.03 | 28.03 | 1.000 |
| 1899.995 | COL1A1 | 956 | 973 | GQRGVVGLPGQRGERGFP | GQRGVVGLP[113]GQ[129]RGERGFP[113] | GQRGVVGLP(1.0000)GQRGERGFP(1.0000) | 19.44 | 12.14 | 1.000 |
| 1917.847 | COL1A1 | 431 | 451 | GNSGEPGAPGSKGDTGAKGEP | GNSGEP[113]GAP[113]GSKGDTGAKGEP[113] | GNSGEP(1.0000)GAP(1.0000)GSKGDTGAKGEP(1.0000) | 20.62 | 11.29 | 0.998 |
| 1918.018 | COL1A1 | 944 | 964 | GAPGTPGPQGIAGQRGVVGLP | GAP[113]GTPGPQGIAGQRGVVGLP[113] | GAP(0.9888)GTP(0.0106)GP(0.0119)QGIAGQRGVVGLP(0.9888) | 22.83 | 18.25 | 1.000 |
| 1918.903 | COL1A1 | 650 | 670 | GPPGEAGKPGEQGVPGDLGAP | GP[113]PGEAGKPGEQGVPGDLGAP[113] | GP(0.5882)P(0.5882)GEAGKP(0.0055)GEQGVP(0.0084)GDLGAP(0.8097) | 24.93 | 24.93 | 0.981 |
| 1921.887 | COL1A1 | 977 | 997 | GPSGEPGKQGPSGASGERGPP | GPSGEPGKQGPSGASGERGPP[113] | GP(0.0038)SGEP(0.0065)GKQGP(0.0049)SGASGERGP(0.0068)P(0.9780) | 25.49 | 25.49 | 1.000 |
| 1923.918 | COL1A1 | 440 | 460 | GSKGDTGAKGEPGPVGVQGPP | GSKGDTGAKGEP[113]GPVGVQGPP[113] | GSKGDTGAKGEP(0.9681)GP(0.0376)VGVQGP(0.0258)P(0.9686) | 21.85 | 21.85 | 0.995 |
| 1934.881 | COL1A1 | 650 | 670 | GPPGEAGKPGEQGVPGDLGAP | GPP[113]GEAGKP[113]GEQGVPGDLGAP[113] | GP(0.0200)P(0.9874)GEAGKP(0.9874)GEQGVP(0.0177)GDLGAP(0.9875) | 50.97 | 50.97 | 1.000 |
| 1935.849 | COL1A1 | 650 | 670 | GPPGEAGKPGEQGVPGDLGAP | GPP[113]GEAGKP[113]GEQ[129]GVP[113]GDLGAP | GP(0.1042)P(0.8323)GEAGKP(0.8342)GEQGVP(0.8342)GDLGAP(0.3950) | 19.84 | 19.84 | 0.999 |
| 1937.877 | COL1A1 | 977 | 997 | GPSGEPGKQGPSGASGERGPP | GPSGEP[113]GKQGPSGASGERGP[113]P | GP(0.0063)SGEP(0.8035)GKQGP(0.0123)SGASGERGP(0.5890)P(0.5890) | 19.99 | 19.99 | 1.000 |
| 1939.916 | COL1A1 | 440 | 460 | GSKGDTGAKGEPGPVGVQGPP | GSKGDTGAKGEP[113]GP[113]VGVQGP[113]P | GSKGDTGAKGEP(0.7798)GP(0.7798)VGVQGP(0.7202)P(0.7202) | 22.44 | 22.44 | 0.977 |
| 1942.896 | COL1A1 | 482 | 502 | GPPGERGGPGSRGFPGADGVA | GPP[113]GERGGP[113]GSRGFP[113]GADGVA | GP(0.0311)P(0.9896)GERGGP(0.9896)GSRGFP(0.9896)GADGVA | 22.50 | 20.31 | 0.996 |
| 1950.856 | COL1A1 | 650 | 670 | GPPGEAGKPGEQGVPGDLGAP | GPP[113]GEAGKP[113]GEQGVP[113]GDLGAP[113] | GP(0.0510)P(0.9872)GEAGKP(0.9873)GEQGVP(0.9872)GDLGAP(0.9872) | 47.82 | 47.82 | 1.000 |
| 1951.882 | COL1A1 | 650 | 670 | GPPGEAGKPGEQGVPGDLGAP | GP[113]PGEAGKP[113]GEQ[129]GVP[113]GDLGAP[113] | GP(0.7706)P(0.7706)GEAGKP(0.8196)GEQGVP(0.8196)GDLGAP(0.8196) | 18.97 | 18.97 | 0.976 |
| 1982.844 | COL1A1 | 197 | 217 | GPQGFQGPPGEPGEPGASGPM | GPQGFQGPPGEP[113]GEP[113]GASGPM | GP(0.0045)QGFQGP(0.0044)P(0.0042)GEP(0.9872)GEP(0.9867)GASGP(0.0065)M(0.0065) | 30.34 | 21.92 | 0.998 |
| 1998.847 | COL1A1 | 197 | 217 | GPQGFQGPPGEPGEPGASGPM | GPQGFQGPPGEP[113]GEP[113]GASGPM[147] | GP(0.0080)QGFQGP(0.0078)P(0.0078)GEP(0.9876)GEP(0.9876)GASGP(0.0140)M(0.9872) | 63.88 | 63.88 | 1.000 |
| 1999.833 | COL1A1 | 197 | 217 | GPQGFQGPPGEPGEPGASGPM | GPQGFQ[129]GPPGEP[113]GEP[113]GASGP[113]M | GP(0.0122)QGFQGP(0.0075)P(0.0075)GEP(0.8470)GEP(0.8468)GASGP(0.6395)M(0.6395) | 18.10 | 18.10 | 0.965 |
| 1999.901 | COL1A1 | 626 | 646 | GPAGERGEQGPAGSPGFQGLP | GPAGERGEQ[129]GPAGSP[113]GFQ[129]GLP[113] | GP(0.0277)AGERGEQGP(0.0304)AGSP(0.9707)GFQGLP(0.9712) | 18.89 | 14.16 | 1.000 |
| 2000.802 | COL1A1 | 197 | 217 | GPQGFQGPPGEPGEPGASGPM | GPQ[129]GFQ[129]GPPGEP[113]GEP[113]GASGP[113]M | GP(0.0085)QGFQGP(0.0092)P(0.0093)GEP(0.8443)GEP(0.8444)GASGP(0.6421)M(0.6421) | 24.36 | 24.36 | 1.000 |
| 2002.966 | COL1A1 | 1200 | 1216 | FLPQPPQEKAHDGGRYY | FLPQPPQEKAHDGGRYY | | 20.01 | 11.18 | 1.000 |
| 2003.967 | COL1A1 | 1200 | 1216 | FLPQPPQEKAHDGGRYY | FLPQPPQ[129]EKAHDGGRYY | | 20.94 | 20.94 | 1.000 |
| 2014.838 | COL1A1 | 197 | 217 | GPQGFQGPPGEPGEPGASGPM | GPQGFQGPP[113]GEP[113]GEP[113]GASGP[113]M | GP(0.0121)QGFQGP(0.0198)P(0.8489)GEP(0.8491)GEP(0.8491)GASGP(0.7105)M(0.7105) | 27.04 | 27.04 | 0.952 |
| 2015.821 | COL1A1 | 197 | 217 | GPQGFQGPPGEPGEPGASGPM | GPQGFQ[129]GP[113]PGEP[113]GEP[113]GASGP[113]M | GP(0.0139)QGFQGP(0.8432)P(0.0291)GEP(0.8436)GEP(0.8435)GASGP(0.7133)M(0.7133) | 40.35 | 40.35 | 1.000 |
| 2030.839 | COL1A1 | 197 | 217 | GPQGFQGPPGEPGEPGASGPM | GPQGFQGP[113]P[113]GEP[113]GEP[113]GASGP[113]M | GP(0.0335)QGFQGP(0.8534)P(0.8534)GEP(0.8534)GEP(0.8535)GASGP(0.7763)M(0.7763) | 23.89 | 18.28 | 0.999 |
| 2041.877 | COL1A1 | 284 | 304 | GPKGEPGSPGENGAPGQMGPR | GPKGEP[113]GSP[113]GEN[115]GAP[113]GQM[147]GPR | GP(0.0302)KGEP(0.8318)GSP(0.8319)GENGAP(0.8181)GQM(0.7440)GP(0.7440)R | 21.40 | 21.40 | 0.989 |
| 2051.915 | COL1A1 | 386 | 409 | GNPGADGQPGAKGANGAPGIAGAP | GNP[113]GADGQ[129]PGAKGAN[115]GAP[113]GIAGAP[113] | GNP(0.9896)GADGQP(0.0312)GAKGANGAP(0.9895)GIAGAP(0.9896) | 19.05 | 15.14 | 0.941 |
| 2079.956 | COL1A1 | 227 | 247 | GKNGDDGEAGKPGRPGERGPP | GKNGDDGEAGKPGRP[113]GERGP[113]P | GKNGDDGEAGKP(0.0108)GRP(0.7642)GERGP(0.6125)P(0.6125) | 23.96 | 23.96 | 1.000 |
| 2080.954 | COL1A1 | 227 | 247 | GKNGDDGEAGKPGRPGERGPP | GKN[115]GDDGEAGKPGRP[113]GERGP[113]P | GKNGDDGEAGKP(0.0104)GRP(0.7649)GERGP(0.6123)P(0.6123) | 29.97 | 29.97 | 1.000 |
| 2083.012 | COL1A1 | 395 | 418 | GAKGANGAPGIAGAPGFPGARGPS | GAKGANGAP[113]GIAGAP[113]GFP[113]GARGPS | GAKGANGAP(0.9366)GIAGAP(0.9366)GFP(0.9357)GARGP(0.1911)S | 29.19 | 29.19 | 1.000 |
| 2083.966 | COL1A1 | 395 | 418 | GAKGANGAPGIAGAPGFPGARGPS | GAKGAN[115]GAP[113]GIAGAP[113]GFP[113]GARGPS | GAKGANGAP(0.9397)GIAGAP(0.9393)GFP(0.9389)GARGP(0.1821)S | 23.64 | 22.20 | 1.000 |
| 2084.098 | COL1A1 | 954 | 973 | IAGQRGVVGLPGQRGERGFP | IAGQRGVVGLP[113]GQ[129]RGERGFP[113] | IAGQRGVVGLP(1.0000)GQRGERGFP(1.0000) | 19.22 | 12.28 | 0.996 |
| 2085.087 | COL1A1 | 954 | 973 | IAGQRGVVGLPGQRGERGFP | IAGQ[129]RGVVGLP[113]GQ[129]RGERGFP[113] | IAGQRGVVGLP(1.0000)GQRGERGFP(1.0000) | 23.31 | 16.75 | 1.000 |
| 2093.917 | COL1A1 | 818 | 841 | GADGQPGAKGEPGDAGAKGDAGPP | GADGQ[129]PGAKGEPGDAGAKGDAGP[113]P | GADGQP(0.0043)GAKGEP(0.0032)GDAGAKGDAGP(0.4963)P(0.4963) | 31.30 | 31.30 | 1.000 |
| 2096.936 | COL1A1 | 227 | 247 | GKNGDDGEAGKPGRPGERGPP | GKN[115]GDDGEAGKP[113]GRP[113]GERGP[113]P | GKNGDDGEAGKP(0.7784)GRP(0.7783)GERGP(0.7216)P(0.7216) | 27.24 | 27.24 | 1.000 |
| 2099.972 | COL1A1 | 395 | 418 | GAKGANGAPGIAGAPGFPGARGPS | GAKGAN[115]GAP[113]GIAGAP[113]GFP[113]GARGP[113]S | GAKGANGAP(1.0000)GIAGAP(1.0000)GFP(1.0000)GARGP(1.0000)S | 22.51 | 12.50 | 1.000 |
| 2101.964 | COL1A1 | 162 | 181 | QLSYGYDEKSTGGISVPGPM | QLSYGYDEKSTGGISVPGP[113]M | QLSYGYDEKSTGGISVP(0.0048)GP(0.4976)M(0.4976) | 19.94 | 19.94 | 1.000 |
| 2108.927 | COL1A1 | 818 | 841 | GADGQPGAKGEPGDAGAKGDAGPP | GADGQPGAKGEP[113]GDAGAKGDAGPP[113] | GADGQP(0.0169)GAKGEP(0.9851)GDAGAKGDAGP(0.0127)P(0.9853) | 19.76 | 19.76 | 1.000 |
| 2109.905 | COL1A1 | 818 | 841 | GADGQPGAKGEPGDAGAKGDAGPP | GADGQ[129]PGAKGEP[113]GDAGAKGDAGP[113]P | GADGQP(0.0102)GAKGEP(0.7643)GDAGAKGDAGP(0.6128)P(0.6128) | 23.21 | 23.21 | 1.000 |
| 2124.934 | COL1A1 | 818 | 841 | GADGQPGAKGEPGDAGAKGDAGPP | GADGQP[113]GAKGEP[113]GDAGAKGDAGP[113]P | GADGQP(0.7798)GAKGEP(0.7798)GDAGAKGDAGP(0.7202)P(0.7202) | 18.75 | 18.75 | 0.991 |
| 2125.892 | COL1A1 | 818 | 841 | GADGQPGAKGEPGDAGAKGDAGPP | GADGQ[129]P[113]GAKGEP[113]GDAGAKGDAGPP[113] | GADGQP(0.9869)GAKGEP(0.9869)GDAGAKGDAGP(0.0391)P(0.9871) | 33.28 | 33.28 | 1.000 |
| 2200.002 | COL1A1 | 428 | 451 | GPKGNSGEPGAPGSKGDTGAKGEP | GPKGNSGEP[113]GAP[113]GSKGDTGAKGEP[113] | GP(0.0338)KGNSGEP(0.9887)GAP(0.9887)GSKGDTGAKGEP(0.9887) | 20.20 | 14.65 | 0.987 |
| 2215.000 | COL1A1 | 1019 | 1042 | GAEGSPGRDGSPGAKGDRGETGPA | GAEGSP[113]GRDGSP[113]GAKGDRGETGPA | GAEGSP(0.9216)GRDGSP(0.9211)GAKGDRGETGP(0.1573)A | 19.42 | 11.41 | 0.985 |
| 2225.057 | COL1A1 | 482 | 505 | GPPGERGGPGSRGFPGADGVAGPK | GPP[113]GERGGP[113]GSRGFP[113]GADGVAGPK | GP(0.0284)P(0.9829)GERGGP(0.9829)GSRGFP(0.9829)GADGVAGP(0.0229)K | 21.39 | 17.66 | 0.988 |
| 2256.067 | COL1A1 | 743 | 766 | GDRGDAGPKGADGSPGKDGVRGLT | GDRGDAGPKGADGSP[113]GKDGVRGLT | GDRGDAGP(0.0124)KGADGSP(0.9876)GKDGVRGLT | 31.84 | 16.40 | 1.000 |
| 2264.126 | COL1A1 | 1091 | 1112 | GPRGDKGETGEQGDRGIKGHRG | GPRGDKGETGEQGDRGIKGHRG | | 24.44 | 12.66 | 1.000 |
| 2308.124 | COL1A1 | 740 | 764 | GPKGDRGDAGPKGADGSPGKDGVRG | GPKGDRGDAGPKGADGSPGKDGVRG | | 23.43 | 11.06 | 0.987 |
| 2324.113 | COL1A1 | 740 | 764 | GPKGDRGDAGPKGADGSPGKDGVRG | GPKGDRGDAGPKGADGSP[113]GKDGVRG | GP(0.0179)KGDRGDAGP(0.0137)KGADGSP(0.9684)GKDGVRG | 24.09 | 14.80 | 1.000 |
| 2333.112 | COL1A1 | 1049 | 1075 | GAPGAPGPVGPAGKSGDRGETGPAGPA | GAP[113]GAP[113]GPVGPAGKSGDRGETGP[113]AGPA | GAP(0.9585)GAP(0.9581)GP(0.0227)VGP(0.0306)AGKSGDRGETGP(0.9558)AGP(0.0743)A | 24.37 | 24.37 | 0.994 |
| 2347.086 | COL1A1 | 224 | 247 | GPPGKNGDDGEAGKPGRPGERGPP | GPP[113]GKNGDDGEAGKPGRP[113]GERGP[113]P | GP(0.0455)P(0.8114)GKNGDDGEAGKP(0.0158)GRP(0.8125)GERGP(0.6574)P(0.6574) | 27.59 | 27.59 | 1.000 |
| 2348.067 | COL1A1 | 224 | 247 | GPPGKNGDDGEAGKPGRPGERGPP | GPP[113]GKN[115]GDDGEAGKP[113]GRP[113]GERGPP | GP(0.0077)P(0.7118)GKNGDDGEAGKP(0.5701)GRP(0.5701)GERGP(0.5701)P(0.5701) | 20.61 | 20.61 | 1.000 |
| 2363.090 | COL1A1 | 224 | 247 | GPPGKNGDDGEAGKPGRPGERGPP | GPP[113]GKNGDDGEAGKP[113]GRPGERGP[113]P[113] | GP(0.0877)P(0.8190)GKNGDDGEAGKP(0.7274)GRP(0.7274)GERGP(0.8192)P(0.8192) | 21.07 | 21.07 | 1.000 |
| 2364.065 | COL1A1 | 224 | 247 | GPPGKNGDDGEAGKPGRPGERGPP | GPP[113]GKN[115]GDDGEAGKP[113]GRP[113]GERGPP[113] | GP(0.1320)P(0.8760)GKNGDDGEAGKP(0.8691)GRP(0.8691)GERGP(0.3767)P(0.8771) | 20.28 | 20.28 | 0.967 |
| 2412.158 | COL1A1 | 359 | 385 | GPRGSEGPQGVRGEPGPPGPAGAAGPA | GPRGSEGPQGVRGEP[113]GPP[113]GPAGAAGPA | GP(0.0071)RGSEGP(0.0055)QGVRGEP(0.9605)GP(0.0400)P(0.9744)GP(0.0079)AGAAGP(0.0046)A | 28.24 | 26.27 | 0.973 |
| 2522.254 | COL1A1 | 740 | 766 | GPKGDRGDAGPKGADGSPGKDGVRGLT | GPKGDRGDAGPKGADGSPGKDGVRGLT | | 27.39 | 15.23 | 1.000 |
| 2538.247 | COL1A1 | 740 | 766 | GPKGDRGDAGPKGADGSPGKDGVRGLT | GPKGDRGDAGPKGADGSP[113]GKDGVRGLT | GP(0.0078)KGDRGDAGP(0.0068)KGADGSP(0.9854)GKDGVRGLT | 22.99 | 12.18 | 1.000 |
| 2554.253 | COL1A1 | 740 | 766 | GPKGDRGDAGPKGADGSPGKDGVRGLT | GP[113]KGDRGDAGPKGADGSP[113]GKDGVRGLT | GP(0.9743)KGDRGDAGP(0.0512)KGADGSP(0.9745)GKDGVRGLT | 31.99 | 23.37 | 1.000 |
| 2653.235 | COL1A1 | 386 | 415 | GNPGADGQPGAKGANGAPGIAGAPGFP | GNP[113]GADGQ[129]PGAKGANGAP[113]GIAGAP[113]GFP[113]G | GNP(0.9650)GADGQP(0.1425)GAKGANGAP(0.9628)GIAGAP(0.9648)GFP(0.9648)GAR | 18.76 | 18.71 | 0.993 |
| 2805.419 | COL1A1 | 740 | 769 | GPKGDRGDAGPKGADGSPGKDGVRGLT | GPKGDRGDAGPKGADGSP[113]GKDGVRGLTGPI | GP(0.0083)KGDRGDAGP(0.0035)KGADGSP(0.9824)GKDGVRGLTGP(0.0058)I | 33.64 | 24.58 | 1.000 |

### Supplemental Table 1. Col1a2 peptide sequences from normal breast by LC-MS/MS.

| M+H | Gene | Domain Start | Domain End | Peptide | Modified Peptide [113] means HYP | Sequence with HYP probability PM:15.9949 | Hyperscore | Nextscore | Peptide Prophet Probability |
|---|---|---|---|---|---|---|---|---|---|
| 659.334 | COL1A2 | 629 | 636 | AVGTAGPS | AVGTAGPS | | 18.94 | 15.02 | 0.943 |
| 711.293 | COL1A2 | 1102 | 1109 | GVSGGGYD | GVSGGGYD | | 19.38 | 9.84 | 1.000 |
| 730.405 | COL1A2 | 625 | 633 | GVVGAVGTA | GVVGAVGTA | | 19.95 | 11.34 | 0.984 |
| 737.417 | COL1A2 | 896 | 903 | PLGIAGPP | PLGIAGPP[113] | P(0.0062)LGIAGP(0.0061)P(0.9877) | 19.79 | 17.86 | 0.969 |
| 741.375 | COL1A2 | 833 | 840 | EVGAVGPP | EVGAVGP[113]P | EVGAVGP(0.5000)P(0.5000) | 18.52 | 18.52 | 0.938 |
| 755.392 | COL1A2 | 344 | 351 | LVGEPGPA | LVGEP[113]GPA | LVGEP(0.9888)GP(0.0112)A | 19.30 | 15.45 | 0.945 |
| 755.402 | COL1A2 | 952 | 960 | GNIGPVGAA | GNIGPVGAA | | 19.71 | 12.92 | 0.975 |
| 756.387 | COL1A2 | 952 | 960 | GNIGPVGAA | GN[115]IGPVGAA | | 20.49 | 14.97 | 0.945 |
| 772.383 | COL1A2 | 886 | 894 | GVAGAVGEP | GVAGAVGEP[113] | GVAGAVGEP(1.0000) | 20.89 | 16.59 | 0.934 |
| 774.357 | COL1A2 | 910 | 918 | GAVGSPGVN | GAVGSP[113]GVN[115] | GAVGSP(1.0000)GVN | 21.65 | 12.55 | 0.968 |
| 784.382 | COL1A2 | 799 | 807 | GPSGISGPP | GPSGISGPP[113] | GP(0.0052)SGISGP(0.0056)P(0.9892) | 21.72 | 19.90 | 0.956 |
| 784.396 | COL1A2 | 86 | 94 | VGLGPGPMG | VGLGPGPMG | | 22.69 | 12.05 | 1.000 |
| 785.374 | COL1A2 | 454 | 462 | GSPGNIGPA | GSP[113]GNIGPA | GSP(0.9891)GNIGP(0.0109)A | 18.68 | 11.15 | 0.977 |
| 785.440 | COL1A2 | 1028 | 1035 | LQGLPGIA | LQ[129]GLP[113]GIA | LQGLP(1.0000)GIA | 18.68 | 14.79 | 0.954 |
| 787.369 | COL1A2 | 721 | 729 | GPNGFAGPA | GPNGFAGPA | | 19.58 | 10.35 | 0.996 |
| 788.354 | COL1A2 | 721 | 729 | GPNGFAGPA | GPN[115]GFAGPA | | 21.30 | 11.87 | 0.989 |
| 794.439 | COL1A2 | 895 | 903 | GPLGIAGPP | GPLGIAGPP[113] | GP(0.0053)LGIAGP(0.0059)P(0.9889) | 19.64 | 17.71 | 0.963 |
| 800.394 | COL1A2 | 86 | 94 | VGLGPGPMG | VGLGPGPM[147]G | VGLGP(0.0057)GP(0.0157)M(0.9786)G | 19.00 | 17.04 | 1.000 |
| 810.471 | COL1A2 | 868 | 876 | GAPGILGLP | GAPGILGLP[113] | GAP(0.0104)GILGLP(0.9896) | 21.77 | 16.98 | 0.938 |
| 812.412 | COL1A2 | 343 | 351 | GLVGEPGPA | GLVGEP[113]GPA | GLVGEP(0.9881)GP(0.0119)A | 21.16 | 15.86 | 0.944 |
| 815.419 | COL1A2 | 627 | 636 | VGAVGTAGPS | VGAVGTAGPS | | 20.53 | 12.02 | 0.988 |
| 824.439 | COL1A2 | 427 | 435 | GPAGVRGPN | GPAGVRGPN | | 19.00 | 9.83 | 0.997 |
| 825.437 | COL1A2 | 954 | 963 | IGPVGAAGAP | IGPVGAAGAP[113] | IGP(0.0104)VGAAGAP(0.9896) | 18.62 | 11.14 | 0.946 |
| 826.434 | COL1A2 | 267 | 276 | IGAVGNAGPA | IGAVGNAGPA | | 19.90 | 12.20 | 0.974 |
| 826.469 | COL1A2 | 868 | 876 | GAPGILGLP | GAP[113]GILGLP[113] | GAP(1.0000)GILGLP(1.0000) | 19.36 | 16.56 | 0.950 |
| 827.420 | COL1A2 | 267 | 276 | IGAVGNAGPA | IGAVGN[115]AGPA | | 21.65 | 12.50 | 0.997 |
| 827.426 | COL1A2 | 562 | 570 | GPAGEVGKP | GPAGEVGKP[113] | GP(0.0120)AGEVGKP(0.9880) | 19.23 | 12.26 | 0.941 |
| 830.379 | COL1A2 | 748 | 756 | GENGVVGPT | GEN[115]GVVGPT | | 19.66 | 12.97 | 0.958 |
| 840.355 | COL1A2 | 784 | 792 | GMTGFPGAA | GM[147]TGFP[113]GAA | GM(1.0000)TGFP(1.0000)GAA | 19.40 | 11.14 | 1.000 |
| 840.445 | COL1A2 | 466 | 474 | GPVGLPGID | GPVGLP[113]GID | GP(0.0104)VGLP(0.9896)GID | 18.43 | 16.78 | 0.946 |
| 841.420 | COL1A2 | 85 | 94 | GVGLGPGPMG | GVGLGPGPMG | | 20.66 | 9.57 | 1.000 |
| 843.453 | COL1A2 | 624 | 633 | PGVVGAVGTA | P[113]GVVGAVGTA | P(1.0000)GVVGAVGTA | 18.02 | 10.26 | 0.999 |
| 844.439 | COL1A2 | 283 | 291 | GEVGLPGLS | GEVGLP[113]GLS | GEVGLP(1.0000)GLS | 20.43 | 15.46 | 0.978 |
| 848.401 | COL1A2 | 1081 | 1089 | GPQGHQGPA | GPQGHQGPA | | 19.44 | 12.47 | 0.999 |
| 849.395 | COL1A2 | 409 | 417 | GADGRAGVM | GADGRAGVM[147] | GADGRAGVM(1.0000) | 19.21 | 10.56 | 0.991 |
| 857.415 | COL1A2 | 85 | 94 | GVGLGPGPMG | GVGLGPGPM[147]G | GVGLGP(0.0050)GP(0.0116)M(0.9834)G | 22.92 | 20.83 | 0.998 |
| 858.355 | COL1A2 | 1102 | 1110 | GVSGGGYDF | GVSGGGYDF | | 19.66 | 13.21 | 0.960 |
| 858.406 | COL1A2 | 109 | 117 | GPQGFQGPA | GPQGFQGPA | | 21.82 | 13.88 | 0.978 |
| 863.385 | COL1A2 | 1099 | 1108 | GPPGVSGGGY | GPP[113]GVSGGGY | GP(0.0120)P(0.9880)GVSGGGY | 21.88 | 19.46 | 0.977 |
| 868.446 | COL1A2 | 951 | 960 | PGNIGPVGAA | P[113]GNIGPVGAA | P(0.9832)GNIGP(0.0168)VGAA | 23.64 | 13.32 | 0.967 |
| 869.471 | COL1A2 | 463 | 471 | GKEGPVGLP | GKEGPVGLP[113] | GKEGP(0.0113)VGLP(0.9887) | 18.13 | 14.57 | 0.925 |
| 874.429 | COL1A2 | 166 | 174 | GFPGTPGLP | GFP[113]GTPGLP[113] | GFP(0.9901)GTP(0.0197)GLP(0.9901) | 23.10 | 13.81 | 0.998 |
| 883.453 | COL1A2 | 862 | 871 | GPQGLLGAPG | GPQ[129]GLLGAP[113]G | GP(0.0193)QGLLGAP(0.9807)G | 18.13 | 16.12 | 0.958 |
| 884.446 | COL1A2 | 562 | 571 | GPAGEVGKPG | GPAGEVGKP[113]G | GP(0.0104)AGEVGKP(0.9896)G | 19.07 | 12.03 | 0.987 |
| 884.494 | COL1A2 | 811 | 819 | GPAGKEGLR | GPAGKEGLR | | 18.16 | 10.57 | 0.997 |
| 885.445 | COL1A2 | 575 | 582 | LHGEFGLP | LHGEFGLP[113] | LHGEFGLP(1.0000) | 19.72 | 11.56 | 0.991 |
| 888.430 | COL1A2 | 88 | 96 | LGPGPMGLM | LGPGPMGLM[147] | LGP(0.0039)GP(0.0039)M(0.0040)GLM(0.9882) | 18.92 | 10.79 | 1.000 |
| 890.419 | COL1A2 | 949 | 957 | GYPGNIGPV | GYP[113]GN[115]IGPV | GYP(0.9817)GNIGP(0.0183)V | 18.53 | 14.98 | 0.999 |
| 890.427 | COL1A2 | 166 | 174 | GFPGTPGLP | GFP[113]GTP[113]GLP[113] | GFP(1.0000)GTP(1.0000)GLP(1.0000) | 18.28 | 14.60 | 0.954 |
| 897.466 | COL1A2 | 466 | 475 | GPVGLPGIDG | GPVGLP[113]GIDG | GP(0.0120)VGLP(0.9880)GIDG | 18.53 | 16.49 | 0.968 |
| 899.441 | COL1A2 | 831 | 840 | TGEVGAVGPP | TGEVGAVGPP[113] | TGEVGAVGP(0.0132)P(0.9868) | 22.07 | 20.05 | 0.973 |
| 901.442 | COL1A2 | 550 | 558 | GPPGFQGLP | GPP[113]GFQGLP[113] | GP(0.0213)P(0.9894)GFQGLP(0.9894) | 20.40 | 18.37 | 0.959 |
| 902.423 | COL1A2 | 550 | 558 | GPPGFQGLP | GPP[113]GFQ[129]GLP[113] | GP(0.0266)P(0.9867)GFQGLP(0.9867) | 19.87 | 17.87 | 0.965 |
| 904.426 | COL1A2 | 88 | 96 | LGPGPMGLM | LGPGPM[147]GLM[147] | LGP(0.0104)GP(0.0202)M(0.9845)GLM(0.9849) | 20.27 | 14.82 | 0.994 |
| 909.442 | COL1A2 | 1068 | 1077 | TGHPGTVGPA | TGHP[113]GTVGPA | TGHP(0.9891)GTVGP(0.0109)A | 18.12 | 12.12 | 0.973 |
| 911.515 | COL1A2 | 285 | 294 | VGLPGLSGPV | VGLP[113]GLSGPV | VGLP(0.9889)GLSGP(0.0111)V | 24.28 | 14.06 | 1.000 |
| 912.472 | COL1A2 | 311 | 321 | AAGLPGVAGAP | AAGLP[113]GVAGAP[113] | AAGLP(1.0000)GVAGAP(1.0000) | 19.41 | 12.60 | 0.975 |
| 914.488 | COL1A2 | 626 | 636 | VVGAVGTAGPS | VVGAVGTAGPS | | 35.01 | 15.94 | 1.000 |
| 915.376 | COL1A2 | 1102 | 1111 | GVSGGGYDFG | GVSGGGYDFG | | 20.56 | 12.74 | 0.997 |
| 924.511 | COL1A2 | 459 | 468 | IGPAGKEGPV | IGPAGKEGPV | | 19.69 | 12.35 | 0.974 |
| 925.500 | COL1A2 | 891 | 900 | VGEPGPLGIA | VGEP[113]GPLGIA | VGEP(0.9891)GP(0.0109)LGIA | 18.81 | 14.58 | 0.951 |
| 939.546 | COL1A2 | 867 | 876 | LGAPGILGLP | LGAP[113]GILGLP[113] | LGAP(1.0000)GILGLP(1.0000) | 23.06 | 15.97 | 0.964 |
| 942.468 | COL1A2 | 574 | 582 | GLHGEFGLP | GLHGEFGLP[113] | GLHGEFGLP(1.0000) | 18.07 | 9.46 | 0.998 |
| 947.447 | COL1A2 | 949 | 958 | GYPGNIGPVG | GYP[113]GN[115]IGPVG | GYP(0.9793)GNIGP(0.0207)VG | 20.72 | 14.54 | 0.999 |
| 956.468 | COL1A2 | 632 | 642 | TAGPSGPSGLP | TAGPSGPSGLP[113] | TAGP(0.0054)SGP(0.0050)SGLP(0.9896) | 20.72 | 15.09 | 0.951 |
| 959.447 | COL1A2 | 550 | 559 | GPPGFQGLPG | GPP[113]GFQ[129]GLP[113]G | GP(0.0282)P(0.9859)GFQGLP(0.9859)G | 18.14 | 16.24 | 0.965 |
| 969.497 | COL1A2 | 310 | 321 | GAAGLPGVAGAP | GAAGLP[113]GVAGAP[113] | GAAGLP(1.0000)GVAGAP(1.0000) | 20.47 | 17.36 | 0.971 |
| 971.511 | COL1A2 | 625 | 636 | GVVGAVGTAGPS | GVVGAVGTAGPS | | 32.50 | 19.07 | 1.000 |
| 972.502 | COL1A2 | 623 | 633 | EPGVVGAVGTA | EP[113]GVVGAVGTA | EP(1.0000)GVVGAVGTA | 23.12 | 10.23 | 1.000 |

| | | | | | | | | | |
|---|---|---|---|---|---|---|---|---|---|
| 974.457 | COL1A2 | 69 | 79 | GPPGLGGNFAA | GPP[113]GLGGN[115]FAA | GP(0.0119)P(0.9881)GLGGNFAA | 18.03 | 16.11 | 0.990 |
| 978.408 | COL1A2 | 1099 | 1109 | GPPGVSGGGYD | GPP[113]GVSGGGYD | GP(0.0105)P(0.9895)GVSGGGYD | 26.45 | 24.17 | 1.000 |
| 980.512 | COL1A2 | 952 | 963 | GNIGPVGAAGAP | GNIGPVGAAGAP | | 19.98 | 11.55 | 0.994 |
| 983.374 | COL1A2 | 1110 | 1117 | FGYDGDFY | FGYDGDFY | | 20.52 | 10.18 | 0.999 |
| 986.469 | COL1A2 | 721 | 732 | GPNGFAGPAGAA | GPNGFAGPAGAA | | 18.02 | 8.08 | 0.998 |
| 987.443 | COL1A2 | 721 | 732 | GPNGFAGPAGAA | GPN[115]GFAGPAGAA | | 30.56 | 18.79 | 1.000 |
| 987.504 | COL1A2 | 625 | 636 | GVVGAVGTAGPS | GVVGAVGTAGP[113]S | GVVGAVGTAGP(1.0000)S | 19.05 | 13.47 | 0.992 |
| 996.504 | COL1A2 | 952 | 963 | GNIGPVGAAGAP | GNIGPVGAAGAP[113] | GNIGP(0.0100)VGAAGAP(0.9900) | 24.21 | 12.04 | 1.000 |
| 996.535 | COL1A2 | 890 | 900 | AVGEPGPLGIA | AVGEP[113]GPLGIA | AVGEP(0.9775)GP(0.0225)LGIA | 18.68 | 17.38 | 0.990 |
| 996.571 | COL1A2 | 867 | 877 | LGAPGILGLPG | LGAP[113]GILGLP[113]G | LGAP(1.0000)GILGLP(1.0000)G | 22.68 | 14.42 | 0.988 |
| 997.476 | COL1A2 | 427 | 437 | GPAGVRGPNGD | GPAGVRGPN[115]GD | | 18.59 | 10.25 | 0.985 |
| 997.492 | COL1A2 | 952 | 963 | GNIGPVGAAGAP | GN[115]IGPVGAAGAP[113] | GNIGP(0.0117)VGAAGAP(0.9883) | 21.14 | 14.48 | 0.974 |
| 999.468 | COL1A2 | 235 | 246 | GSDGSVGPVGPA | GSDGSVGPVGPA | | 21.49 | 13.10 | 0.977 |
| 999.517 | COL1A2 | 826 | 836 | GPVGRTGEVGA | GPVGRTGEVGA | | 20.42 | 12.44 | 0.945 |
| 1012.501 | COL1A2 | 265 | 276 | GEIGAVGNAGPA | GEIGAVGNAGPA | | 24.14 | 15.49 | 0.997 |
| 1013.484 | COL1A2 | 265 | 276 | GEIGAVGNAGPA | GEIGAVGN[115]AGPA | | 27.72 | 21.09 | 1.000 |
| 1015.464 | COL1A2 | 910 | 921 | GAVGSPGVNGAP | GAVGSP[113]GVN[115]GAP[113] | GAVGSP(1.0000)GVNGAP(1.0000) | 22.19 | 15.69 | 0.988 |
| 1021.423 | COL1A2 | 1103 | 1112 | VSGGGYDFGY | VSGGGYDFGY | | 18.55 | 11.34 | 0.996 |
| 1029.515 | COL1A2 | 622 | 633 | GEPGVVGAVGTA | GEP[113]GVVGAVGTA | GEP(1.0000)GVVGAVGTA | 24.38 | 10.63 | 1.000 |
| 1032.467 | COL1A2 | 688 | 699 | GATGDRGEAGAA | GATGDRGEAGAA | | 21.69 | 14.66 | 0.966 |
| 1032.472 | COL1A2 | 438 | 447 | AGRPGEPGLM | AGRP[113]GEP[113]GLM[147] | AGRP(1.0000)GEP(1.0000)GLM(1.0000) | 22.90 | 11.14 | 0.998 |
| 1036.635 | COL1A2 | 866 | 876 | LLGAPGILGLP | LLGAPGILGLP[113] | LLGAP(0.0105)GILGLP(0.9895) | 22.75 | 10.19 | 0.983 |
| 1039.539 | COL1A2 | 886 | 897 | GVAGAVGEPGPL | GVAGAVGEP[113]GPL | GVAGAVGEP(0.9885)GP(0.0115)L | 21.50 | 18.34 | 0.976 |
| 1040.474 | COL1A2 | 772 | 783 | GPAGSRGDGGPP | GPAGSRGDGGPP[113] | GP(0.0054)AGSRGDGGP(0.0060)P(0.9886) | 18.86 | 17.22 | 0.986 |
| 1041.571 | COL1A2 | 827 | 837 | PVGRTGEVGAV | PVGRTGEVGAV | | 18.94 | 11.65 | 0.974 |
| 1052.625 | COL1A2 | 866 | 876 | LLGAPGILGLP | LLGAP[113]GILGLP[113] | LLGAP(1.0000)GILGLP(1.0000) | 27.58 | 14.21 | 0.977 |
| 1053.551 | COL1A2 | 889 | 900 | GAVGEPGPLGIA | GAVGEP[113]GPLGIA | GAVGEP(0.9888)GP(0.0112)LGIA | 25.79 | 18.11 | 1.000 |
| 1055.531 | COL1A2 | 883 | 894 | GLPGVAGAVGEP | GLP[113]GVAGAVGEP[113] | GLP(1.0000)GVAGAVGEP(1.0000) | 27.54 | 17.77 | 1.000 |
| 1058.520 | COL1A2 | 232 | 243 | GARGSDGSVGPV | GARGSDGSVGPV | | 25.69 | 12.01 | 0.998 |
| 1060.520 | COL1A2 | 86 | 96 | VGLGPGPMGLM | VGLGPGPM[147]GLM[147] | VGLGP(0.0133)GP(0.0356)M(0.9752)GLM(0.9759) | 25.92 | 19.89 | 1.000 |
| 1068.500 | COL1A2 | 427 | 438 | GPAGVRGPNGDA | GPAGVRGPN[115]GDA | | 19.24 | 12.60 | 0.966 |
| 1068.528 | COL1A2 | 451 | 462 | GLPGSPGNIGPA | GLP[113]GSP[113]GNIGPA | GLP(0.9876)GSP(0.9876)GNIGP(0.0248)A | 18.37 | 12.63 | 0.963 |
| 1069.565 | COL1A2 | 262 | 273 | GPKGEIGAVGNA | GPKGEIGAVGNA | | 19.95 | 12.32 | 0.983 |
| 1072.547 | COL1A2 | 949 | 960 | GYPGNIGPVGAA | GYPGNIGPVGAA | | 20.75 | 13.83 | 0.992 |
| 1075.492 | COL1A2 | 1034 | 1044 | IAGHHGDQGAP | IAGHHGDQGAP[113] | IAGHHGDQGAP(1.0000) | 18.13 | 11.23 | 0.994 |
| 1078.434 | COL1A2 | 1102 | 1112 | GVSGGGYDFGY | GVSGGGYDFGY | | 21.81 | 10.66 | 1.000 |
| 1080.571 | COL1A2 | 886 | 898 | GVAGAVGEPGPLG | GVAGAVGEPGPLG | | 23.99 | 20.06 | 0.980 |
| 1085.524 | COL1A2 | 247 | 258 | GPIGSAGPPGFP | GPIGSAGPP[113]GFP[113] | GP(0.0118)IGSAGP(0.0127)P(0.9877)GFP(0.9878) | 21.09 | 17.39 | 1.000 |
| 1088.533 | COL1A2 | 949 | 960 | GYPGNIGPVGAA | GYP[113]GNIGPVGAA | GYP(0.9896)GNIGP(0.0104)VGAA | 28.67 | 14.31 | 1.000 |
| 1089.513 | COL1A2 | 949 | 960 | GYPGNIGPVGAA | GYP[113]GN[115]IGPVGAA | GYP(0.9886)GNIGP(0.0114)VGAA | 22.81 | 14.46 | 0.964 |
| 1096.558 | COL1A2 | 886 | 898 | GVAGAVGEPGPLG | GVAGAVGEP[113]GPLG | GVAGAVGEP(0.9830)GP(0.0170)LG | 28.23 | 20.29 | 0.999 |
| 1096.577 | COL1A2 | 340 | 351 | GARGLVGEPGPA | GARGLVGEP[113]GPA | GARGLVGEP(0.9896)GP(0.0104)A | 22.22 | 16.52 | 0.994 |
| 1097.577 | COL1A2 | 283 | 294 | GEVGLPGLSGPV | GEVGLP[113]GLSGPV | GEVGLP(0.9863)GLSGP(0.0137)V | 20.21 | 11.39 | 0.966 |
| 1097.599 | COL1A2 | 281 | 291 | PRGEVGLPGLS | PRGEVGLP[113]GLS | P(0.0112)RGEVGLP(0.9888)GLS | 19.07 | 10.69 | 0.958 |
| 1098.571 | COL1A2 | 826 | 837 | GPVGRTGEVGAV | GPVGRTGEVGAV | | 18.01 | 10.28 | 0.967 |
| 1099.536 | COL1A2 | 454 | 465 | GSPGNIGPAGKE | GSP[113]GNIGPAGKE | GSP(0.9886)GNIGP(0.0114)AGKE | 18.94 | 10.14 | 1.000 |
| 1100.520 | COL1A2 | 454 | 465 | GSPGNIGPAGKE | GSP[113]GN[115]IGPAGKE | GSP(0.9891)GNIGP(0.0109)AGKE | 18.31 | 11.04 | 0.997 |
| 1101.519 | COL1A2 | 247 | 258 | GPIGSAGPPGFP | GPIGSAGP[113]P[113]GFP[113] | GP(0.0344)IGSAGP(0.9885)P(0.9886)GFP(0.9885) | 20.75 | 12.12 | 1.000 |
| 1101.569 | COL1A2 | 158 | 168 | VVGPQGARGFP | VVGPQ[129]GARGFP[113] | VVGP(0.0119)QGARGFP(0.9881) | 18.00 | 10.19 | 0.999 |
| 1109.606 | COL1A2 | 1072 | 1083 | GTVGPAGIRGPQ | GTVGPAGIRGPQ | | 22.06 | 10.09 | 1.000 |
| 1109.652 | COL1A2 | 865 | 876 | GLLGAPGILGLP | GLLGAP[113]GILGLP[113] | GLLGAP(1.0000)GILGLP(1.0000) | 27.51 | 13.70 | 1.000 |
| 1110.561 | COL1A2 | 575 | 585 | LHGEFGLPGPA | LHGEFGLP[113]GPA | LHGEFGLP(0.9891)GP(0.0109)A | 20.75 | 15.87 | 0.961 |
| 1112.532 | COL1A2 | 709 | 720 | GSPGERGEVGPA | GSPGERGEVGPA | | 20.41 | 13.18 | 0.990 |
| 1112.555 | COL1A2 | 745 | 756 | GPKGENGVVGPT | GPKGEN[115]GVVGPT | | 19.56 | 15.14 | 0.920 |
| 1112.570 | COL1A2 | 829 | 840 | GRTGEVGAVGPP | GRTGEVGAVGPP[113] | GRTGEVGAVGP(0.0131)P(0.9869) | 20.27 | 20.27 | 0.952 |
| 1116.505 | COL1A2 | 409 | 420 | GADGRAGVMGPP | GADGRAGVM[147]GPP[113] | GADGRAGVM(0.9857)GP(0.0286)P(0.9856) | 22.89 | 22.89 | 0.959 |
| 1117.475 | COL1A2 | 113 | 123 | FQGPAGEPGEP | FQGPAGEP[113]GEP[113] | FQGP(0.0195)AGEP(0.9902)GEP(0.9902) | 19.17 | 11.85 | 0.981 |
| 1117.532 | COL1A2 | 85 | 96 | GVGLGPGPMGLM | GVGLGPGP[113]MGLM[147] | GVGLGP(0.0144)GP(0.6113)M(0.6113)GLM(0.7631) | 18.10 | 14.55 | 0.975 |
| 1123.558 | COL1A2 | 856 | 867 | GPPGTPGPQGLL | GPP[113]GTP[113]GPQ[129]GLL | GP(0.0117)P(0.9887)GTP(0.9887)GP(0.0109)QGLL | 21.58 | 19.55 | 0.995 |
| 1125.474 | COL1A2 | 1099 | 1110 | GPPGVSGGGYDF | GPP[113]GVSGGGYDF | GP(0.0106)P(0.9894)GVSGGGYDF | 21.64 | 19.61 | 1.000 |
| 1127.572 | COL1A2 | 745 | 756 | GPKGENGVVGPT | GP[113]KGENGVVGPT | GP(0.9843)KGENGVVGP(0.0157)T | 21.57 | 15.72 | 0.976 |
| 1128.524 | COL1A2 | 709 | 720 | GSPGERGEVGPA | GSP[113]GERGEVGPA | GSP(0.9888)GERGEVGP(0.0112)A | 19.21 | 11.86 | 0.978 |
| 1128.544 | COL1A2 | 745 | 756 | GPKGENGVVGPT | GP[113]KGEN[115]GVVGPT | GP(0.9864)KGENGVVGP(0.0136)T | 20.40 | 11.51 | 0.990 |
| 1132.507 | COL1A2 | 1033 | 1044 | GIAGHHGDQGAP | GIAGHHGDQGAP[113] | GIAGHHGDQGAP(1.0000) | 22.05 | 11.66 | 0.996 |
| 1136.445 | COL1A2 | 1103 | 1113 | VSGGGYDFGYD | VSGGGYDFGYD | | 22.40 | 10.45 | 1.000 |
| 1139.483 | COL1A2 | 1110 | 1118 | FGYDGDFYR | FGYDGDFYR | | 18.29 | 10.03 | 0.974 |
| 1142.549 | COL1A2 | 550 | 561 | GPPGFQGLPGPS | GPP[113]GFQGLP[113]GPS | GP(0.1566)P(0.8685)GFQGLP(0.8721)GP(0.1028)S | 22.04 | 21.92 | 0.999 |
| 1150.617 | COL1A2 | 466 | 477 | GPVGLPGIDGRP | GPVGLP[113]GIDGRP | GP(0.0052)VGLP(0.9779)GIDGRP(0.0169) | 29.36 | 14.42 | 0.998 |
| 1154.522 | COL1A2 | 784 | 795 | GMTGFPGAAGRT | GM[147]TGFP[113]GAAGRT | GM(1.0000)TGFP(1.0000)GAAGRT | 19.70 | 9.35 | 1.000 |
| 1154.590 | COL1A2 | 280 | 291 | GPRGEVGLPGLS | GPRGEVGLP[113]GLS | GP(0.0350)RGEVGLP(0.9650)GLS | 21.03 | 11.52 | 1.000 |
| 1154.603 | COL1A2 | 463 | 474 | GKEGPVGLPGID | GKEGPVGLP[113]GID | GKEGP(0.0110)VGLP(0.9890)GID | 27.21 | 16.04 | 0.989 |
| 1166.607 | COL1A2 | 466 | 477 | GPVGLPGIDGRP | GPVGLP[113]GIDGRP[113] | GP(0.0212)VGLP(0.9894)GIDGRP(0.9894) | 29.59 | 13.39 | 0.988 |

| | | | | | | | | | |
|---|---|---|---|---|---|---|---|---|---|
| 1169.584 | COL1A2 | 562 | 573 | GPAGEVGKPGER | GPAGEVGKP[113]GER | GP(0.0109)AGEVGKP(0.9891)GER | 18.81 | 10.02 | 0.946 |
| 1174.590 | COL1A2 | 163 | 174 | GARGFPGTPGLP | GARGFP[113]GTP[113]GLP[113] | GARGFP(1.0000)GTP(1.0000)GLP(1.0000) | 19.56 | 13.90 | 0.973 |
| 1174.596 | COL1A2 | 1078 | 1089 | GIRGPQGHQGPA | GIRGPQGHQGPA | | 26.76 | 10.68 | 1.000 |
| 1182.478 | COL1A2 | 1099 | 1111 | GPPGVSGGGYDFG | GPP[113]GVSGGGYDFG | GP(0.0105)P(0.9895)GVSGGGYDFG | 39.16 | 36.56 | 1.000 |
| 1193.464 | COL1A2 | 1102 | 1113 | GVSGGGYDFGYD | GVSGGGYDFGYD | | 32.81 | 15.51 | 1.000 |
| 1198.506 | COL1A2 | 1099 | 1111 | GPPGVSGGGYDFG | GP[113]P[113]GVSGGGYDFG | GP(1.0000)P(1.0000)GVSGGGYDFG | 18.70 | 8.04 | 1.000 |
| 1203.568 | COL1A2 | 484 | 495 | GARGEPGNIGFP | GARGEP[113]GNIGFP[113] | GARGEP(1.0000)GNIGFP(1.0000) | 19.76 | 11.75 | 0.946 |
| 1204.520 | COL1A2 | 436 | 447 | GDAGRPGEPGLM | GDAGRP[113]GEP[113]GLM[147] | GDAGRP(1.0000)GEP(1.0000)GLM(1.0000) | 21.23 | 11.31 | 0.988 |
| 1204.558 | COL1A2 | 484 | 495 | GARGEPGNIGFP | GARGEP[113]GN[115]IGFP[113] | GARGEP(1.0000)GNIGFP(1.0000) | 18.40 | 12.72 | 0.973 |
| 1207.670 | COL1A2 | 459 | 471 | IGPAGKEGPVGLP | IGPAGKEGPVGLP[113] | IGP(0.0097)AGKEGP(0.0081)VGLP(0.9822) | 27.41 | 21.97 | 0.996 |
| 1210.516 | COL1A2 | 1110 | 1119 | FGYDGDFYRA | FGYDGDFYRA | | 19.12 | 9.81 | 0.999 |
| 1211.623 | COL1A2 | 463 | 475 | GKEGPVGLPGIDG | GKEGPVGLP[113]GIDG | GKEGP(0.0106)VGLP(0.9894)GIDG | 25.56 | 13.77 | 0.999 |
| 1220.649 | COL1A2 | 219 | 231 | PGERGRVGAPGPA | PGERGRVGAPGPA | | 19.61 | 14.94 | 0.926 |
| 1222.657 | COL1A2 | 1071 | 1083 | PGTVGPAGIRGPQ | P[113]GTVGPAGIRGPQ | P(0.9892)GTVGP(0.0055)AGIRGP(0.0054)Q | 21.35 | 13.44 | 0.994 |
| 1223.654 | COL1A2 | 887 | 900 | VAGAVGEPGPLGIA | VAGAVGEP[113]GPLGIA | VAGAVGEP(0.9528)GP(0.0472)LGIA | 26.13 | 22.26 | 0.956 |
| 1226.606 | COL1A2 | 562 | 574 | GPAGEVGKPGERG | GPAGEVGKP[113]GERG | GP(0.0103)AGEVGKP(0.9897)GERG | 20.70 | 12.84 | 0.991 |
| 1235.655 | COL1A2 | 1068 | 1080 | TGHPGTVGPAGIR | TGHP[113]GTVGPAGIR | TGHP(0.9886)GTVGP(0.0114)AGIR | 19.95 | 9.80 | 1.000 |
| 1237.667 | COL1A2 | 178 | 189 | GIRGHNGLDGLK | GIRGHN[115]GLDGLK | | 22.72 | 14.50 | 1.000 |
| 1250.483 | COL1A2 | 1102 | 1114 | GVSGGGYDFGYDG | GVSGGGYDFGYDG | | 30.21 | 10.73 | 1.000 |
| 1257.569 | COL1A2 | 685 | 699 | GPAGATGDRGEAGAA | GPAGATGDRGEAGAA | | 36.58 | 13.08 | 1.000 |
| 1263.655 | COL1A2 | 890 | 903 | AVGEPGPLGIAGPP | AVGEP[113]GPLGIAGPP[113] | AVGEP(0.6118)GP(0.6118)LGIAGP(0.0093)P(0.7671) | 19.14 | 19.14 | 0.986 |
| 1270.621 | COL1A2 | 622 | 636 | GEPGVVGAVGTAGPS | GEP[113]GVVGAVGTAGPS | GEP(0.9896)GVVGAVGTAGP(0.0104)S | 27.92 | 15.75 | 1.000 |
| 1272.609 | COL1A2 | 298 | 312 | GNPGANGLTGAKGAA | GNP[113]GAN[115]GLTGAKGAA | GNP(1.0000)GANGLTGAKGAA | 19.90 | 11.82 | 1.000 |
| 1278.590 | COL1A2 | 842 | 855 | FAGEKGPSGEAGTA | FAGEKGPSGEAGTA | | 35.71 | 11.33 | 1.000 |
| 1280.679 | COL1A2 | 886 | 900 | GVAGAVGEPGPLGIA | GVAGAVGEP[113]GPLGIA | GVAGAVGEP(0.9870)GP(0.0130)LGIA | 29.73 | 23.51 | 1.000 |
| 1283.620 | COL1A2 | 232 | 246 | GARGSDGSVGPVGPA | GARGSDGSVGPVGPA | | 26.19 | 13.93 | 0.993 |
| 1284.630 | COL1A2 | 571 | 582 | GERGLHGEFGLP | GERGLHGEFGLP[113] | GERGLHGEFGLP(1.0000) | 18.27 | 10.82 | 0.976 |
| 1294.587 | COL1A2 | 842 | 855 | FAGEKGPSGEAGTA | FAGEKGP[113]SGEAGTA | FAGEKGP(1.0000)SGEAGTA | 24.87 | 12.51 | 1.000 |
| 1294.667 | COL1A2 | 262 | 276 | GPKGEIGAVGNAGPA | GPKGEIGAVGNAGPA | | 20.02 | 13.53 | 0.987 |
| 1295.647 | COL1A2 | 262 | 276 | GPKGEIGAVGNAGPA | GPKGEIGAVGN[115]AGPA | | 34.89 | 21.74 | 1.000 |
| 1296.566 | COL1A2 | 116 | 129 | PAGEPGEPGQTGPA | PAGEP[113]GEP[113]GQTGPA | P(0.0281)AGEP(0.9802)GEP(0.9809)GQTGP(0.0109)A | 22.72 | 20.36 | 0.991 |
| 1310.622 | COL1A2 | 247 | 261 | GPIGSAGPPGFPGAP | GPIGSAGP[113]PGFPGAP[113] | GP(0.0162)IGSAGP(0.5833)P(0.5833)GFP(0.0062)GAP(0.8109) | 18.87 | 18.87 | 0.992 |
| 1310.669 | COL1A2 | 262 | 276 | GPKGEIGAVGNAGPA | GP[113]KGEIGAVGNAGPA | GP(0.9897)KGEIGAVGNAGP(0.0103)A | 35.47 | 24.11 | 1.000 |
| 1311.648 | COL1A2 | 262 | 276 | GPKGEIGAVGNAGPA | GP[113]KGEIGAVGN[115]AGPA | GP(0.9886)KGEIGAVGNAGP(0.0114)A | 34.60 | 23.60 | 0.999 |
| 1320.679 | COL1A2 | 889 | 903 | GAVGEPGPLGIAGPP | GAVGEP[113]GPLGIAGPP[113] | GAVGEP(0.9631)GP(0.0150)LGIAGP(0.0609)P(0.9610) | 28.11 | 25.89 | 0.999 |
| 1325.678 | COL1A2 | 337 | 351 | GATGARGLVGEPGPA | GATGARGLVGEP[113]GPA | GATGARGLVGEP(0.9885)GP(0.0115)A | 24.47 | 18.78 | 0.997 |
| 1326.628 | COL1A2 | 247 | 261 | GPIGSAGPPGFPGAP | GPIGSAGPP[113]GFP[113]GAP[113] | GP(0.0175)IGSAGP(0.0332)P(0.9829)GFP(0.9832)GAP(0.9832) | 19.96 | 19.96 | 0.990 |
| 1328.687 | COL1A2 | 619 | 633 | GNKGEPGVVGAVGTA | GNKGEP[113]GVVGAVGTA | GNKGEP(1.0000)GVVGAVGTA | 18.38 | 9.16 | 0.998 |
| 1329.636 | COL1A2 | 949 | 963 | GYPGNIGPVGAAGAP | GYP[113]GNIGPVGAAGAP[113] | GYP(0.9866)GNIGP(0.0268)VGAAGAP(0.9866) | 27.85 | 18.91 | 1.000 |
| 1330.620 | COL1A2 | 949 | 963 | GYPGNIGPVGAAGAP | GYP[113]GN[115]IGPVGAAGAP[113] | GYP(0.9891)GNIGP(0.0219)VGAAGAP(0.9890) | 23.36 | 14.70 | 0.992 |
| 1335.604 | COL1A2 | 841 | 855 | GFAGEKGPSGEAGTA | GFAGEKGPSGEAGTA | | 45.78 | 22.55 | 1.000 |
| 1339.692 | COL1A2 | 562 | 575 | GPAGEVGKPGERGL | GPAGEVGKP[113]GERGL | GP(0.0103)AGEVGKP(0.9897)GERGL | 18.23 | 11.70 | 1.000 |
| 1345.564 | COL1A2 | 1099 | 1112 | GPPGVSGGGYDFGY | GP[113]PGVSGGGYDFGY | GP(0.5000)P(0.5000)GVSGGGYDFGY | 26.21 | 26.21 | 1.000 |
| 1351.610 | COL1A2 | 841 | 855 | GFAGEKGPSGEAGTA | GFAGEKGP[113]SGEAGTA | GFAGEKGP(1.0000)SGEAGTA | 25.16 | 15.31 | 0.999 |
| 1351.694 | COL1A2 | 637 | 651 | GPSGLPGERGAAGIP | GPSGLPGERGAAGIP[113] | GP(0.0063)SGLP(0.0073)GERGAAGIP(0.9864) | 29.31 | 19.86 | 0.994 |
| 1352.679 | COL1A2 | 454 | 468 | GSPGNIGPAGKEGPV | GSP[113]GNIGPAGKEGPV | GSP(0.9887)GNIGP(0.0056)AGKEGP(0.0057)V | 28.86 | 16.09 | 1.000 |
| 1353.580 | COL1A2 | 115 | 129 | GPAGEPGEPGQTGPA | GPAGEP[113]GEP[113]GQTGPA | GP(0.0669)AGEP(0.9601)GEP(0.9625)GQTGP(0.0105)A | 30.06 | 29.77 | 0.999 |
| 1353.641 | COL1A2 | 454 | 468 | GSPGNIGPAGKEGPV | GSP[113]GN[115]IGPAGKEGPV | GSP(0.9868)GNIGP(0.0063)AGKEGP(0.0069)V | 23.46 | 17.91 | 1.000 |
| 1354.563 | COL1A2 | 115 | 129 | GPAGEPGEPGQTGPA | GPAGEP[113]GEP[113]GQ[129]TGPA | GP(0.0107)AGEP(0.9880)GEP(0.9879)GQTGP(0.0133)A | 25.91 | 21.86 | 0.969 |
| 1365.693 | COL1A2 | 826 | 840 | GPVGRTGEVGAVGPP | GPVGRTGEVGAVGPP[113] | GP(0.0053)VGRTGEVGAVGP(0.0057)P(0.9890) | 45.20 | 45.20 | 1.000 |
| 1367.694 | COL1A2 | 637 | 651 | GPSGLPGERGAAGIP | GPSGLP[113]GERGAAGIP[113] | GP(0.0268)SGLP(0.9866)GERGAAGIP(0.9865) | 22.81 | 17.72 | 0.980 |
| 1370.618 | COL1A2 | 115 | 129 | GPAGEPGEPGQTGPA | GP[113]AGEP[113]GEPGQTGP[113]A | GP(0.7641)AGEP(0.7621)GEP(0.6819)GQTGP(0.7919)A | 19.36 | 19.18 | 0.983 |
| 1372.683 | COL1A2 | 158 | 171 | VVGPQGARGFPGTP | VVGPQ[129]GARGFP[113]GTP[113] | VVGP(0.0406)QGARGFP(0.9796)GTP(0.9797) | 18.47 | 10.75 | 0.997 |
| 1378.731 | COL1A2 | 457 | 471 | GNIGPAGKEGPVGLP | GNIGPAGKEGPVGLP[113] | GNIGP(0.0117)AGKEGP(0.0055)VGLP(0.9828) | 30.46 | 25.53 | 1.000 |
| 1379.711 | COL1A2 | 883 | 898 | GLPGVAGAVGEPGPLG | GLP[113]GVAGAVGEPGP[113]LG | GLP(0.7895)GVAGAVGEP(0.4335)GP(0.7770)LG | 31.81 | 29.46 | 0.993 |
| 1379.711 | COL1A2 | 460 | 474 | GPAGKEGPVGLPGID | GPAGKEGPVGLP[113]GID | GP(0.0057)AGKEGP(0.0055)VGLP(0.9888)GID | 19.60 | 16.33 | 0.952 |
| 1381.707 | COL1A2 | 826 | 840 | GPVGRTGEVGAVGPP | GPVGRTGEVGAVGP[113]P[113] | GP(0.0331)VGRTGEVGAVGP(0.9835)P(0.9835) | 27.29 | 17.93 | 0.992 |
| 1384.693 | COL1A2 | 661 | 674 | GLRGEIGNPGRDGA | GLRGEIGNP[113]GRDGA | GLRGEIGNP(1.0000)GRDGA | 19.86 | 11.87 | 1.000 |
| 1384.708 | COL1A2 | 300 | 315 | PGANGLTGAKGAAGLP | P[113]GAN[115]GLTGAKGAAGLP[113] | P(1.0000)GANGLTGAKGAAGLP(1.0000) | 27.72 | 11.49 | 1.000 |
| 1391.782 | COL1A2 | 862 | 876 | GPQGLLGAPGILGLP | GPQGLLGAP[113]GILGLP[113] | GP(0.0316)QGLLGAP(0.9842)GILGLP(0.9842) | 23.27 | 12.26 | 0.987 |
| 1397.699 | COL1A2 | 880 | 894 | GERGLPGVAGAVGEP | GERGLP[113]GVAGAVGEP[113] | GERGLP(1.0000)GVAGAVGEP(1.0000) | 21.60 | 17.98 | 0.933 |
| 1398.685 | COL1A2 | 823 | 837 | GDQGPVGRTGEVGAV | GDQGPVGRTGEVGAV | | 34.61 | 13.37 | 1.000 |
| 1398.703 | COL1A2 | 621 | 636 | KGEPGVVGAVGTAGPS | KGEP[113]GVVGAVGTAGPS | KGEP(0.9894)GVVGAVGTAGP(0.0106)S | 31.22 | 11.90 | 1.000 |
| 1399.623 | COL1A2 | 196 | 210 | GVKGEPGAPGENGTP | GVKGEP[113]GAP[113]GEN[115]GTP | GVKGEP(0.9835)GAP(0.9834)GENGTP(0.0330) | 23.89 | 14.66 | 0.982 |
| 1403.599 | COL1A2 | 113 | 126 | FQGPAGEPGEPGQT | FQGPAGEP[113]GEP[113]GQT | FQGP(0.0215)AGEP(0.9893)GEP(0.9893)GQT | 26.33 | 18.37 | 1.000 |
| 1407.737 | COL1A2 | 280 | 294 | GPRGEVGLPGLSGPV | GPRGEVGLP[113]GLSGPV | GP(0.0059)RGEVGLP(0.9885)GLSGP(0.0056)V | 27.46 | 14.34 | 1.000 |
| 1408.672 | COL1A2 | 535 | 549 | GPQGVQGGKGEQGPP | GPQGVQGGKGEQGPP[113] | GP(0.0060)QGVQGGKGEQGP(0.0063)P(0.9876) | 22.51 | 22.51 | 1.000 |
| 1409.655 | COL1A2 | 535 | 549 | GPQGVQGGKGEQGPP | GPQGVQ[129]GGKGEQGPP[113] | GP(0.0142)QGVQGGKGEQGP(0.0064)P(0.9794) | 24.96 | 24.96 | 0.957 |
| 1415.613 | COL1A2 | 196 | 210 | GVKGEPGAPGENGTP | GVKGEP[113]GAP[113]GEN[115]GTP[113] | GVKGEP(1.0000)GAP(1.0000)GENGTP(1.0000) | 23.45 | 9.97 | 0.986 |
| 1416.723 | COL1A2 | 1069 | 1083 | GHPGTVGPAGIRGPQ | GHP[113]GTVGPAGIRGPQ | GHP(0.9871)GTVGP(0.0068)AGIRGP(0.0061)Q | 27.36 | 18.13 | 1.000 |
| 1417.716 | COL1A2 | 1069 | 1083 | GHPGTVGPAGIRGPQ | GHP[113]GTVGPAGIRGPQ[129] | GHP(0.9893)GTVGP(0.0053)AGIRGP(0.0054)Q | 19.28 | 11.83 | 0.999 |
| 1421.685 | COL1A2 | 970 | 984 | GPAGKHGNRGETGPS | GPAGKHGNRGETGPS | | 23.46 | 12.61 | 1.000 |

| | | | | | | | | | |
|---|---|---|---|---|---|---|---|---|---|
| 1422.700 | COL1A2 | 1060 | 1074 | GPAGKDGRTGHPGTV | GPAGKDGRTGHP[113]GTV | GP(0.0157)AGKDGRTGHP(0.9843)GTV | 22.71 | 11.47 | 1.000 |
| 1422.734 | COL1A2 | 826 | 841 | GPVGRTGEVGAVGPPG | GPVGRTGEVGAVGPP[113]G | GP(0.0057)VGRTGEVGAVGP(0.0115)P(0.9828)G | 27.49 | 27.49 | 0.992 |
| 1425.681 | COL1A2 | 616 | 631 | GPDGNKGEPGVVGAVG | GPDGNKGEP[113]GVVGAVG | GP(0.0388)DGNKGEP(0.9612)GVVGAVG | 19.71 | 15.81 | 0.975 |
| 1428.685 | COL1A2 | 481 | 495 | GPAGARGEPGNIGFP | GPAGARGEP[113]GNIGFP[113] | GP(0.0277)AGARGEP(0.9861)GNIGFP(0.9861) | 18.32 | 12.41 | 0.971 |
| 1431.678 | COL1A2 | 946 | 960 | GERGYPGNIGPVGAA | GERGYP[113]GN[115]IGPVGAA | GERGYP(0.9703)GNIGP(0.0297)VGAA | 27.83 | 15.74 | 0.996 |
| 1432.689 | COL1A2 | 403 | 417 | GSRGLPGADGRAGVM | GSRGLP[113]GADGRAGVM[147] | GSRGLP(1.0000)GADGRAGVM(1.0000) | 24.05 | 11.75 | 1.000 |
| 1435.748 | COL1A2 | 457 | 472 | GNIGPAGKEGPVGLPG | GNIGPAGKEGPVGLP[113]G | GNIGP(0.0053)AGKEGP(0.0052)VGLP(0.9895)G | 25.01 | 16.14 | 0.996 |
| 1437.630 | COL1A2 | 784 | 798 | GMTGFPGAAGRTGPP | GM[147]TGFP[113]GAAGRTGP[113]P[113] | GM(1.0000)TGFP(1.0000)GAAGRTGP(1.0000)P(1.0000) | 28.03 | 10.42 | 1.000 |
| 1441.593 | COL1A2 | 922 | 936 | GEAGRDGNPGNDGPP | GEAGRDGNP[113]GNDGPP[113] | GEAGRDGNP(0.9883)GNDGP(0.0234)P(0.9883) | 18.40 | 18.40 | 0.976 |
| 1443.694 | COL1A2 | 618 | 633 | DGNKGEPGVVGAVGTA | DGNKGEP[113]GVVGAVGTA | DGNKGEP(1.0000)GVVGAVGTA | 22.19 | 11.86 | 1.000 |
| 1457.630 | COL1A2 | 433 | 447 | GPNGDAGRPGEPGLM | GPN[115]GDAGRP[113]GEP[113]GLM | GP(0.0115)NGDAGRP(0.9808)GEP(0.9802)GLM(0.0275) | 30.18 | 26.05 | 1.000 |
| 1460.589 | COL1A2 | 1099 | 1113 | GPPGVSGGGYDFGYD | GPP[113]GVSGGGYDFGYD | GP(0.0202)P(0.9798)GVSGGGYDFGYD | 18.18 | 16.30 | 0.996 |
| 1464.747 | COL1A2 | 463 | 477 | GKEGPVGLPGIDGRP | GKEGPVGLP[113]GIDGRP | GKEGP(0.0052)VGLP(0.9895)GIDGRP(0.0052) | 23.94 | 14.20 | 1.000 |
| 1465.660 | COL1A2 | 499 | 514 | GPTGDPGKNGDKGHAG | GPTGDPGKN[115]GDKGHAG | | 19.35 | 12.59 | 0.960 |
| 1472.637 | COL1A2 | 433 | 447 | GPNGDAGRPGEPGLM | GPNGDAGRP[113]GEP[113]GLM[147] | GP(0.0432)NGDAGRP(0.9856)GEP(0.9856)GLM(0.9856) | 20.20 | 7.08 | 1.000 |
| 1473.618 | COL1A2 | 433 | 447 | GPNGDAGRPGEPGLM | GPN[115]GDAGRP[113]GEP[113]GLM[147] | GP(0.0327)NGDAGRP(0.9891)GEP(0.9891)GLM(0.9891) | 28.92 | 11.74 | 1.000 |
| 1476.747 | COL1A2 | 562 | 576 | GPAGEVGKPGERGLH | GPAGEVGKP[113]GERGLH | GP(0.0123)AGEVGKP(0.9877)GERGLH | 18.34 | 10.54 | 0.996 |
| 1480.757 | COL1A2 | 463 | 477 | GKEGPVGLPGIDGRP | GKEGPVGLP[113]GIDGRP[113] | GKEGP(0.0224)VGLP(0.9888)GIDGRP(0.9888) | 25.12 | 13.83 | 1.000 |
| 1481.669 | COL1A2 | 499 | 514 | GPTGDPGKNGDKGHAG | GPTGDP[113]GKN[115]GDKGHAG | GP(0.0100)TGDP(0.9900)GKNGDKGHAG | 19.40 | 10.51 | 0.999 |
| 1482.686 | COL1A2 | 685 | 702 | GPAGATGDRGEAGAAGPA | GPAGATGDRGEAGAAGPA | | 32.71 | 12.15 | 1.000 |
| 1490.786 | COL1A2 | 887 | 903 | VAGAVGEPGPLGIAGPP | VAGAVGEP[113]GPLGIAGPP[113] | VAGAVGEP(0.9692)GP(0.0219)LGIAGP(0.0404)P(0.9685) | 28.50 | 28.50 | 0.997 |
| 1492.787 | COL1A2 | 459 | 474 | IGPAGKEGPVGLPGID | IGPAGKEGPVGLP[113]GID | IGP(0.0057)AGKEGP(0.0054)VGLP(0.9889)GID | 20.85 | 12.02 | 0.987 |
| 1498.677 | COL1A2 | 685 | 702 | GPAGATGDRGEAGAAGPA | GPAGATGDRGEAGAAGP[113]A | GP(0.0105)AGATGDRGEAGAAGP(0.9895)A | 31.29 | 13.24 | 1.000 |
| 1501.777 | COL1A2 | 1068 | 1083 | TGHPGTVGPAGIRGPQ | TGHPGTVGPAGIRGPQ | | 27.39 | 9.51 | 1.000 |
| 1508.736 | COL1A2 | 229 | 246 | GPAGARGSDGSVGPVGPA | GPAGARGSDGSVGPVGPA | | 28.30 | 12.96 | 1.000 |
| 1511.803 | COL1A2 | 1004 | 1017 | IRGDKGEPGEKGPR | IRGDKGEP[113]GEKGPR | IRGDKGEP(0.9829)GEKGP(0.0171)R | 20.76 | 14.84 | 1.000 |
| 1517.615 | COL1A2 | 1099 | 1114 | GPPGVSGGGYDFGYDG | GP[113]PGVSGGGYDFGYDG | GP(0.5000)P(0.5000)GVSGGGYDFGYDG | 35.81 | 35.81 | 1.000 |
| 1517.757 | COL1A2 | 1068 | 1083 | TGHPGTVGPAGIRGPQ | TGHP[113]GTVGPAGIRGPQ | TGHP(0.9861)GTVGP(0.0075)AGIRGP(0.0064)Q | 22.07 | 15.38 | 1.000 |
| 1518.747 | COL1A2 | 1068 | 1083 | TGHPGTVGPAGIRGPQ | TGHP[113]GTVGPAGIRGPQ[129] | TGHP(0.9891)GTVGP(0.0052)AGIRGP(0.0057)Q | 21.86 | 13.14 | 1.000 |
| 1524.805 | COL1A2 | 661 | 675 | GLRGEIGNPGRDGAR | GLRGEIGNPGRDGAR | | 21.13 | 12.29 | 0.994 |
| 1527.724 | COL1A2 | 1034 | 1050 | IAGHHGDQGAPGSVGPA | IAGHHGDQGAPGSVGPA | | 19.03 | 12.33 | 0.955 |
| 1533.775 | COL1A2 | 562 | 577 | GPAGEVGKPGERGLHG | GPAGEVGKP[113]GERGLHG | GP(0.0103)AGEVGKP(0.9897)GERGLHG | 19.99 | 12.32 | 0.997 |
| 1538.760 | COL1A2 | 820 | 835 | GPRGDQGPVGRTGEVG | GPRGDQGPVGRTGEVG | | 21.58 | 11.99 | 1.000 |
| 1539.750 | COL1A2 | 298 | 315 | GNPGANGLTGAKGAAGLP | GNPGAN[115]GLTGAKGAAGLP[113] | GNP(0.0189)GANGLTGAKGAAGLP(0.9811) | 24.50 | 17.97 | 1.000 |
| 1540.756 | COL1A2 | 617 | 633 | PDGNKGEPGVVGAVGTA | PDGNKGEP[113]GVVGAVGTA | P(0.0142)DGNKGEP(0.9858)GVVGAVGTA | 19.39 | 15.25 | 0.993 |
| 1540.794 | COL1A2 | 661 | 675 | GLRGEIGNPGRDGAR | GLRGEIGNP[113]GRDGAR | GLRGEIGNP(1.0000)GRDGAR | 18.28 | 8.44 | 1.000 |
| 1541.776 | COL1A2 | 661 | 675 | GLRGEIGNPGRDGAR | GLRGEIGN[115]P[113]GRDGAR | GLRGEIGNP(1.0000)GRDGAR | 19.33 | 14.46 | 1.000 |
| 1543.718 | COL1A2 | 1034 | 1050 | IAGHHGDQGAPGSVGPA | IAGHHGDQGAP[113]GSVGPA | IAGHHGDQGAP(0.9892)GSVGP(0.0108)A | 21.90 | 13.17 | 1.000 |
| 1545.710 | COL1A2 | 842 | 858 | FAGEKGPSGEAGTAGPP | FAGEKGPSGEAGTAGPP[113] | FAGEKGP(0.0058)SGEAGTAGP(0.0062)P(0.9880) | 45.77 | 45.77 | 1.000 |
| 1547.786 | COL1A2 | 886 | 903 | GVAGAVGEPGPLGIAGPP | GVAGAVGEP[113]GPLGIAGP[113]P | GVAGAVGEP(0.7372)GP(0.0663)LGIAGP(0.5982)P(0.5982) | 41.35 | 41.35 | 1.000 |
| 1552.807 | COL1A2 | 331 | 348 | GPVGAAGATGARGLVGEP | GPVGAAGATGARGLVGEP[113] | GP(0.0124)VGAAGATGARGLVGEP(0.9876) | 43.71 | 13.85 | 1.000 |
| 1555.766 | COL1A2 | 298 | 315 | GNPGANGLTGAKGAAGLP | GNP[113]GAN[115]GLTGAKGAAGLP[113] | GNP(1.0000)GANGLTGAKGAAGLP(1.0000) | 26.86 | 18.05 | 1.000 |
| 1556.724 | COL1A2 | 298 | 315 | GNPGANGLTGAKGAAGLP | GN[115]P[113]GAN[115]GLTGAKGAAGLP[113] | GNP(1.0000)GANGLTGAKGAAGLP(1.0000) | 22.99 | 13.20 | 1.000 |
| 1563.816 | COL1A2 | 883 | 900 | GLPGVAGAVGEPGPLGIA | GLP[113]GVAGAVGEP[113]GPLGIA | GLP(0.9714)GVAGAVGEP(0.9711)GP(0.0575)LGIA | 32.07 | 27.90 | 1.000 |
| 1569.777 | COL1A2 | 619 | 636 | GNKGEPGVVGAVGTAGPS | GNKGEP[113]GVVGAVGTAGPS | GNKGEP(0.9892)GVVGAVGTAGP(0.0108)S | 35.31 | 13.40 | 1.000 |
| 1581.779 | COL1A2 | 616 | 633 | GPDGNKGEPGVVGAVGTA | GPDGNKGEPGVVGAVGTA | | 30.21 | 11.53 | 1.000 |
| 1582.766 | COL1A2 | 616 | 633 | GPDGNKGEPGVVGAVGTA | GPDGN[115]KGEPGVVGAVGTA | | 19.44 | 10.48 | 0.993 |
| 1584.745 | COL1A2 | 1033 | 1050 | GIAGHHGDQGAPGSVGPA | GIAGHHGDQGAPGSVGPA | | 18.30 | 12.64 | 0.954 |
| 1597.752 | COL1A2 | 616 | 633 | GPDGNKGEPGVVGAVGTA | GPDGNKGEP[113]GVVGAVGTA | GP(0.0106)DGNKGEP(0.9894)GVVGAVGTA | 49.74 | 28.14 | 1.000 |
| 1598.759 | COL1A2 | 616 | 633 | GPDGNKGEPGVVGAVGTA | GPDGN[115]KGEP[113]GVVGAVGTA | GP(0.0103)DGNKGEP(0.9897)GVVGAVGTA | 47.84 | 31.64 | 1.000 |
| 1599.730 | COL1A2 | 910 | 927 | GAVGSPGVNGAPGEAGRD | GAVGSP[113]GVNGAP[113]GEAGRD | GAVGSP(1.0000)GVNGAP(1.0000)GEAGRD | 25.58 | 9.66 | 1.000 |
| 1600.699 | COL1A2 | 910 | 927 | GAVGSPGVNGAPGEAGRD | GAVGSP[113]GVN[115]GAP[113]GEAGRD | GAVGSP(1.0000)GVNGAP(1.0000)GEAGRD | 23.72 | 10.04 | 1.000 |
| 1600.733 | COL1A2 | 1033 | 1050 | GIAGHHGDQGAPGSVGPA | GIAGHHGDQGAP[113]GSVGPA | GIAGHHGDQGAP(0.9883)GSVGP(0.0117)A | 21.78 | 13.37 | 0.997 |
| 1602.723 | COL1A2 | 841 | 858 | GFAGEKGPSGEAGTAGPP | GFAGEKGPSGEAGTAGPP[113] | GFAGEKGP(0.0057)SGEAGTAGP(0.0062)P(0.9881) | 66.98 | 66.98 | 1.000 |
| 1602.735 | COL1A2 | 349 | 366 | GPAGSKGESGNKGEPGSA | GPAGSKGESGNKGEP[113]GSA | GP(0.0133)AGSKGESGNKGEP(0.9867)GSA | 21.61 | 12.75 | 1.000 |
| 1609.794 | COL1A2 | 820 | 836 | GPRGDQGPVGRTGEVGA | GPRGDQGPVGRTGEVGA | | 27.27 | 10.75 | 1.000 |
| 1610.782 | COL1A2 | 820 | 836 | GPRGDQGPVGRTGEVGA | GPRGDQ[129]GPVGRTGEVGA | | 19.27 | 11.23 | 0.993 |
| 1613.767 | COL1A2 | 616 | 633 | GPDGNKGEPGVVGAVGTA | GP[113]DGNKGEP[113]GVVGAVGTA | GP(1.0000)DGNKGEP(1.0000)GVVGAVGTA | 33.34 | 12.18 | 1.000 |
| 1614.757 | COL1A2 | 616 | 633 | GPDGNKGEPGVVGAVGTA | GP[113]DGN[115]KGEP[113]GVVGAVGTA | GP(1.0000)DGNKGEP(1.0000)GVVGAVGTA | 21.51 | 12.08 | 0.997 |
| 1618.635 | COL1A2 | 1103 | 1117 | VSGGGYDFGYDGDFY | VSGGGYDFGYDGDFY | | 33.04 | 12.49 | 1.000 |
| 1618.713 | COL1A2 | 841 | 858 | GFAGEKGPSGEAGTAGPP | GFAGEKGPSGEAGTAGP[113]P[113] | GFAGEKGP(0.0255)SGEAGTAGP(0.9872)P(0.9872) | 37.54 | 15.30 | 1.000 |
| 1628.698 | COL1A2 | 113 | 129 | FQGPAGEPGEPGQTGPA | FQGPAGEP[113]GEP[113]GQTGPA | FQGP(0.0118)AGEP(0.9836)GEP(0.9833)GQTGP(0.0213)A | 31.81 | 20.90 | 1.000 |
| 1629.693 | COL1A2 | 113 | 129 | FQGPAGEPGEPGQTGPA | FQGPAGEP[113]GEP[113]GQ[129]TGPA | FQGP(0.0110)AGEP(0.9863)GEP(0.9861)GQTGP(0.0167)A | 23.07 | 19.50 | 1.000 |
| 1631.814 | COL1A2 | 1060 | 1077 | GPAGKDGRTGHPGTVGPA | GPAGKDGRTGHPGTVGPA | | 20.22 | 9.84 | 1.000 |
| 1635.827 | COL1A2 | 454 | 471 | GSPGNIGPAGKEGPVGLP | GSP[113]GNIGPAGKEGPVGLP[113] | GSP(0.9865)GNIGP(0.0126)AGKEGP(0.0146)VGLP(0.9863) | 30.47 | 28.25 | 1.000 |
| 1636.815 | COL1A2 | 454 | 471 | GSPGNIGPAGKEGPVGLP | GSP[113]GN[115]IGPAGKEGPVGLP[113] | GSP(0.9818)GNIGP(0.0185)AGKEGP(0.0182)VGLP(0.9815) | 18.18 | 18.18 | 0.972 |
| 1640.724 | COL1A2 | 193 | 210 | GAPGVKGEPGAPGENGTP | GAP[113]GVKGEP[113]GAP[113]GEN[115]GTP | GAP(0.9885)GVKGEP(0.9885)GAP(0.9885)GENGTP(0.0344) | 22.38 | 15.31 | 1.000 |
| 1640.733 | COL1A2 | 712 | 729 | GERGEVGPAGPNGFAGPA | GERGEVGPAGPN[115]GFAGPA | | 24.73 | 11.97 | 1.000 |
| 1645.728 | COL1A2 | 113 | 129 | FQGPAGEPGEPGQTGPA | FQGP[113]AGEP[113]GEPGQTGP[113]A | FQGP(0.7368)AGEP(0.7673)GEP(0.7140)GQTGP(0.7819)A | 25.22 | 21.73 | 0.982 |
| 1646.728 | COL1A2 | 113 | 129 | FQGPAGEPGEPGQTGPA | FQGP[113]AGEP[113]GEPGQ[129]TGP[113]A | FQGP(0.7597)AGEP(0.7702)GEP(0.6721)GQTGP(0.7979)A | 21.10 | 20.66 | 0.955 |
| 1647.810 | COL1A2 | 1060 | 1077 | GPAGKDGRTGHPGTVGPA | GPAGKDGRTGHP[113]GTVGPA | GP(0.0078)AGKDGRTGHP(0.9861)GTVGP(0.0060)A | 21.77 | 14.53 | 0.977 |
| 1649.778 | COL1A2 | 499 | 516 | GPTGDPGKNGDKGHAGLA | GPTGDPGKN[115]GDKGHAGLA | | 20.50 | 9.60 | 1.000 |

Supplemental Table 1. Col1a1 peptide sequences from normal breast by LC-MS/MS.

| | | | | | | | | | |
|---|---|---|---|---|---|---|---|---|---|
| 1656.728 | COL1A2 | 193 | 210 | GAPGVKGEPGAPGENGTP | GAP[113]GVKGEP[113]GAP[113]GEN[115]GTP[113] | GAP(1.0000)GVKGEP(1.0000)GAP(1.0000)GENGTP(1.0000) | 24.81 | 16.27 | 1.000 |
| 1656.753 | COL1A2 | 373 | 390 | GPSGEEGKRGPNGEAGSA | GPSGEEGKRGPNGEAGSA | | 23.55 | 12.30 | 1.000 |
| 1657.734 | COL1A2 | 373 | 390 | GPSGEEGKRGPNGEAGSA | GPSGEEGKRGPN[115]GEAGSA | | 20.22 | 11.68 | 0.994 |
| 1659.746 | COL1A2 | 352 | 369 | GSKGESGNKGEPGSAGPQ | GSKGESGNKGEP[113]GSAGPQ | GSKGESGNKGEP(0.9895)GSAGP(0.0105)Q | 24.33 | 15.53 | 0.991 |
| 1660.733 | COL1A2 | 352 | 369 | GSKGESGNKGEPGSAGPQ | GSKGESGN[115]KGEP[113]GSAGPQ | GSKGESGNKGEP(0.9898)GSAGP(0.0102)Q | 23.49 | 14.97 | 0.986 |
| 1662.815 | COL1A2 | 562 | 578 | GPAGEVGKPGERGLHGE | GPAGEVGKP[113]GERGLHGE | GP(0.0136)AGEVGKP(0.9864)GERGLHGE | 23.70 | 11.81 | 1.000 |
| 1664.810 | COL1A2 | 499 | 516 | GPTGDPGKNGDKGHAGLA | GPTGDP[113]GKNGDKGHAGLA | GP(0.0342)TGDP(0.9658)GKNGDKGHAGLA | 19.08 | 11.22 | 0.999 |
| 1665.774 | COL1A2 | 499 | 516 | GPTGDPGKNGDKGHAGLA | GPTGDP[113]GKN[115]GDKGHAGLA | GP(0.0142)TGDP(0.9858)GKNGDKGHAGLA | 19.31 | 12.87 | 0.989 |
| 1665.802 | COL1A2 | 586 | 603 | GPRGERGPPGESGAAGPT | GPRGERGP[113]PGESGAAGPT | GP(0.0042)RGERGP(0.9690)P(0.0234)GESGAAGP(0.0035)T | 19.82 | 19.82 | 0.966 |
| 1665.812 | COL1A2 | 823 | 840 | GDQGPVGRTGEVGAVGPP | GDQGPVGRTGEVGAVGPP[113] | GDQGP(0.0061)VGRTGEVGAVGP(0.0193)P(0.9746) | 41.94 | 41.94 | 1.000 |
| 1672.778 | COL1A2 | 946 | 963 | GERGYPGNIGPVGAAGAP | GERGYP[113]GN[115]IGPVGAAGAP[113] | GERGYP(0.9782)GNIGP(0.0436)VGAAGAP(0.9782) | 25.25 | 15.40 | 0.983 |
| 1674.819 | COL1A2 | 967 | 984 | GPVGPAGKHGNRGETGPS | GPVGPAGKHGNRGETGPS | | 20.04 | 12.26 | 0.974 |
| 1674.877 | COL1A2 | 280 | 297 | GPRGEVGLPGLSGPVGPP | GPRGEVGLP[113]GLSGPVGPP[113] | GP(0.0077)RGEVGLP(0.9877)GLSGP(0.0070)VGP(0.0101)P(0.9876) | 31.51 | 31.51 | 1.000 |
| 1675.644 | COL1A2 | 1102 | 1117 | GVSGGGYDFGYDGDFY | GVSGGGYDFGYDGDFY | | 22.92 | 10.70 | 1.000 |
| 1675.810 | COL1A2 | 967 | 984 | GPVGPAGKHGNRGETGPS | GPVGPAGKHGN[115]RGETGPS | | 20.70 | 10.77 | 1.000 |
| 1682.697 | COL1A2 | 919 | 936 | GAPGEAGRDGNPGNDGPP | GAP[113]GEAGRDGNP[113]GNDGPP[113] | GAP(0.9779)GEAGRDGNP(0.9779)GNDGP(0.0666)P(0.9777) | 18.71 | 18.71 | 1.000 |
| 1684.788 | COL1A2 | 618 | 636 | DGNKGEPGVVGAVGTAGPS | DGNKGEP[113]GVVGAVGTAGPS | DGNKGEP(0.9886)GVVGAVGTAGP(0.0114)S | 31.12 | 14.46 | 1.000 |
| 1685.732 | COL1A2 | 113 | 130 | FQGPAGEPGEPGQTGPAG | FQGPAGEP[113]GEP[113]GQTGPAG | FQGP(0.0133)AGEP(0.9749)GEP(0.9739)GQTGP(0.0378)AG | 33.99 | 24.85 | 1.000 |
| 1686.706 | COL1A2 | 113 | 130 | FQGPAGEPGEPGQTGPAG | FQGPAGEP[113]GEP[113]GQ[129]TGPAG | FQGP(0.0114)AGEP(0.9850)GEP(0.9847)GQTGP(0.0189)AG | 28.57 | 19.98 | 0.980 |
| 1692.836 | COL1A2 | 454 | 472 | GSPGNIGPAGKEGPVGLPG | GSP[113]GNIGPAGKEGPVGLP[113]G | GSP(0.9869)GNIGP(0.0146)AGKEGP(0.0115)VGLP(0.9869)G | 21.06 | 17.46 | 1.000 |
| 1693.832 | COL1A2 | 454 | 472 | GSPGNIGPAGKEGPVGLPG | GSP[113]GN[115]IGPAGKEGPVGLP[113]G | GSP(0.9826)GNIGP(0.0227)AGKEGP(0.0117)VGLP(0.9830)G | 19.73 | 15.63 | 0.943 |
| 1698.827 | COL1A2 | 643 | 660 | GERGAAGIPGGKGEKGEP | GERGAAGIP[113]GGKGEKGEP[113] | GERGAAGIP(1.0000)GGKGEKGEP(1.0000) | 19.40 | 11.49 | 0.993 |
| 1704.832 | COL1A2 | 1060 | 1078 | GPAGKDGRTGHPGTVGPAG | GPAGKDGRTGHP[113]GTVGPAG | GP(0.0122)AGKDGRTGHP(0.9814)GTVGP(0.0064)AG | 22.97 | 17.31 | 0.981 |
| 1707.803 | COL1A2 | 685 | 705 | GPAGATGDRGEAGAAGPAGPA | GPAGATGDRGEAGAAGPAGPA | | 20.88 | 14.05 | 0.983 |
| 1708.853 | COL1A2 | 820 | 837 | GPRGDQGPVGRTGEVGAV | GPRGDQGPVGRTGEVGAV | | 23.91 | 12.08 | 1.000 |
| 1709.848 | COL1A2 | 820 | 837 | GPRGDQGPVGRTGEVGAV | GPRGDQ[129]GPVGRTGEVGAV | | 24.30 | 12.54 | 1.000 |
| 1712.798 | COL1A2 | 431 | 447 | VRGPNGDAGRPGEPGLM | VRGPN[115]GDAGRP[113]GEPGLM[147] | VRGP(0.0130)NGDAGRP(0.9883)GEP(0.0102)GLM(0.9884) | 19.88 | 15.41 | 0.982 |
| 1724.882 | COL1A2 | 662 | 678 | LRGEIGNPGRDGARGAP | LRGEIGNP[113]GRDGARGAP[113] | LRGEIGNP(1.0000)GRDGARGAP(1.0000) | 20.33 | 11.34 | 0.994 |
| 1728.782 | COL1A2 | 431 | 447 | VRGPNGDAGRPGEPGLM | VRGPN[115]GDAGRP[113]GEP[113]GLM[147] | VRGP(0.0316)NGDAGRP(0.9895)GEP(0.9895)GLM(0.9895) | 20.41 | 11.69 | 0.970 |
| 1742.756 | COL1A2 | 109 | 126 | GPQGFQGPAGEPGEPGQT | GPQGFQGPAGEP[113]GEP[113]GQT | GP(0.0247)QGFQGP(0.0130)AGEP(0.9812)GEP(0.9811)GQT | 24.89 | 16.44 | 1.000 |
| 1743.737 | COL1A2 | 109 | 126 | GPQGFQGPAGEPGEPGQT | GPQGFQ[129]GPAGEP[113]GEP[113]GQT | GP(0.0283)QGFQGP(0.0140)AGEP(0.9808)GEP(0.9768)GQT | 28.16 | 24.28 | 1.000 |
| 1750.804 | COL1A2 | 427 | 445 | GPAGVRGPNGDAGRPGEPG | GPAGVRGPN[115]GDAGRP[113]GEP[113]G | GP(0.0190)AGVRGP(0.0440)NGDAGRP(0.9682)GEP(0.9688)G | 19.63 | 10.08 | 0.987 |
| 1753.847 | COL1A2 | 943 | 960 | GHKGERGYPGNIGPVGAA | GHKGERGYP[113]GN[115]IGPVGAA | GHKGERGYP(0.9894)GNIGP(0.0106)VGAA | 21.69 | 15.58 | 0.966 |
| 1769.811 | COL1A2 | 430 | 447 | GVRGPNGDAGRPGEPGLM | GVRGPN[115]GDAGRP[113]GEP[113]GLM | GVRGP(0.0106)NGDAGRP(0.9895)GEP(0.9896)GLM(0.0103) | 22.59 | 15.83 | 0.997 |
| 1771.769 | COL1A2 | 910 | 929 | GAVGSPGVNGAPGEAGRDGN | GAVGSP[113]GVN[115]GAP[113]GEAGRDGN | GAVGSP(1.0000)GVNGAP(1.0000)GEAGRDGN | 21.13 | 11.10 | 1.000 |
| 1772.740 | COL1A2 | 910 | 929 | GAVGSPGVNGAPGEAGRDGN | GAVGSP[113]GVN[115]GAP[113]GEAGRDGN[115] | GAVGSP(1.0000)GVNGAP(1.0000)GEAGRDGN | 19.88 | 12.75 | 0.989 |
| 1774.735 | COL1A2 | 1103 | 1118 | VSGGGYDFGYDGDFYR | VSGGGYDFGYDGDFYR | | 23.96 | 9.49 | 1.000 |
| 1777.910 | COL1A2 | 331 | 351 | GPVGAAGATGARGLVGEPGPA | GPVGAAGATGARGLVGEP[113]GPA | GP(0.0071)VGAAGATGARGLVGEP(0.9512)GP(0.0417)A | 35.50 | 35.50 | 1.000 |
| 1781.881 | COL1A2 | 661 | 678 | GLRGEIGNPGRDGARGAP | GLRGEIGNP[113]GRDGARGAP[113] | GLRGEIGNP(1.0000)GRDGARGAP(1.0000) | 23.03 | 18.77 | 1.000 |
| 1784.830 | COL1A2 | 430 | 447 | GVRGPNGDAGRPGEPGLM | GVRGPNGDAGRP[113]GEP[113]GLM[147] | GVRGP(0.0298)NGDAGRP(0.9901)GEP(0.9901)GLM(0.9900) | 19.66 | 12.14 | 0.996 |
| 1784.872 | COL1A2 | 661 | 678 | GLRGEIGNPGRDGARGAP | GLRGEIGN[115]P[113]GRDGARGAP[113] | GLRGEIGNP(1.0000)GRDGARGAP(1.0000) | 21.47 | 15.72 | 1.000 |
| 1785.808 | COL1A2 | 430 | 447 | GVRGPNGDAGRPGEPGLM | GVRGPN[115]GDAGRP[113]GEP[113]GLM[147] | GVRGP(0.0399)NGDAGRP(0.9867)GEP(0.9867)GLM(0.9867) | 20.54 | 12.81 | 0.999 |
| 1809.883 | COL1A2 | 562 | 579 | GPAGEVGKPGERGLHGEF | GPAGEVGKP[113]GERGLHGEF | GP(0.0132)AGEVGKP(0.9868)GERGLHGEF | 37.82 | 18.46 | 1.000 |
| 1816.821 | COL1A2 | 842 | 861 | FAGEKGPSGEAGTAGPPGTP | FAGEKGPSGEAGTAGPP[113]GTP[113] | FAGEKGP(0.0188)SGEAGTAGP(0.0181)P(0.9814)GTP(0.9817) | 18.15 | 18.15 | 0.942 |
| 1818.940 | COL1A2 | 459 | 477 | IGPAGKEGPVGLPGIDGRP | IGPAGKEGPVGLP[113]GIDGRP[113] | IGP(0.0247)AGKEGP(0.0118)VGLP(0.9818)GIDGRP(0.9818) | 23.48 | 13.83 | 0.966 |
| 1822.892 | COL1A2 | 616 | 636 | GPDGNKGEPGVVGAVGTAGPS | GPDGNKGEPGVVGAVGTAGPS | | 33.39 | 14.10 | 1.000 |
| 1830.953 | COL1A2 | 883 | 903 | GLPGVAGAVGEPGPLGIAGPP | GLP[113]GVAGAVGEP[113]GPLGIAGP[113]P | GLP(0.8001)GVAGAVGEP(0.8006)GP(0.0159)LGIAGP(0.6917)P(0.6917) | 48.41 | 48.41 | 1.000 |
| 1831.744 | COL1A2 | 1102 | 1118 | GVSGGGYDFGYDGDFYR | GVSGGGYDFGYDGDFYR | | 24.58 | 11.90 | 1.000 |
| 1838.852 | COL1A2 | 616 | 636 | GPDGNKGEPGVVGAVGTAGPS | GPDGNKGEP[113]GVVGAVGTAGPS | GP(0.0057)DGNKGEP(0.9882)GVVGAVGTAGP(0.0061)S | 54.93 | 34.46 | 1.000 |
| 1839.870 | COL1A2 | 616 | 636 | GPDGNKGEPGVVGAVGTAGPS | GPDGN[115]KGEP[113]GVVGAVGTAGPS | GP(0.0050)DGNKGEP(0.9901)GVVGAVGTAGP(0.0049)S | 42.74 | 32.46 | 1.000 |
| 1845.769 | COL1A2 | 1103 | 1119 | VSGGGYDFGYDGDFYRA | VSGGGYDFGYDGDFYRA | | 31.74 | 8.49 | 1.000 |
| 1850.946 | COL1A2 | 1000 | 1017 | GPQGIRGDKGEPGEKGPR | GPQGIRGDKGEP[113]GEKGPR | GP(0.0087)QGIRGDKGEP(0.9746)GEKGP(0.0167)R | 22.11 | 15.50 | 1.000 |
| 1851.921 | COL1A2 | 565 | 582 | GEVGKPGERGLHGEFGLP | GEVGKPGERGLHGEFGLP[113] | GEVGKP(0.0109)GERGLHGEFGLP(0.9891) | 19.76 | 13.31 | 0.938 |
| 1854.872 | COL1A2 | 616 | 636 | GPDGNKGEPGVVGAVGTAGPS | GP[113]DGNKGEP[113]GVVGAVGTAGPS | GP(0.9901)DGNKGEP(0.9901)GVVGAVGTAGP(0.0198)S | 24.19 | 10.80 | 1.000 |
| 1855.859 | COL1A2 | 616 | 636 | GPDGNKGEPGVVGAVGTAGPS | GPDGN[115]KGEP[113]GVVGAVGTAGP[113]S | GP(0.0470)DGNKGEP(0.9764)GVVGAVGTAGP(0.9766)S | 24.16 | 17.31 | 1.000 |
| 1856.839 | COL1A2 | 429 | 447 | AGVRGPNGDAGRPGEPGLM | AGVRGPN[115]GDAGRP[113]GEP[113]GLM[147] | AGVRGP(0.0355)NGDAGRP(0.9882)GEP(0.9882)GLM(0.9882) | 19.67 | 11.19 | 1.000 |
| 1858.898 | COL1A2 | 950 | 970 | YPGNIGPVGAAGAPGPHGPVG | YPGN[115]IGPVGAAGAP[113]GPHGPVG | YP(0.0025)GNIGP(0.0025)VGAAGAP(0.9892)GP(0.0029)HGP(0.0029)VG | 18.52 | 16.87 | 0.939 |
| 1866.941 | COL1A2 | 1000 | 1017 | GPQGIRGDKGEPGEKGPR | GP[113]QGIRGDKGEP[113]GEKGPR | GP(0.9888)QGIRGDKGEP(0.9905)GEKGP(0.0206)R | 21.63 | 13.15 | 1.000 |
| 1867.929 | COL1A2 | 565 | 582 | GEVGKPGERGLHGEFGLP | GEVGKP[113]GERGLHGEFGLP[113] | GEVGKP(1.0000)GERGLHGEFGLP(1.0000) | 30.27 | 16.62 | 1.000 |
| 1867.934 | COL1A2 | 1000 | 1017 | GPQGIRGDKGEPGEKGPR | GP[113]Q[129]GIRGDKGEP[113]GEKGPR | GP(0.9883)QGIRGDKGEP(0.9907)GEKGP(0.0210)R | 20.80 | 19.28 | 0.977 |
| 1868.825 | COL1A2 | 910 | 930 | GAVGSPGVNGAPGEAGRDGNP | GAVGSP[113]GVN[115]GAP[113]GEAGRDGNP | GAVGSP(0.9306)GVNGAP(0.9293)GEAGRDGNP(0.1401) | 22.23 | 19.98 | 0.999 |
| 1869.854 | COL1A2 | 838 | 858 | GPPGFAGEKGPSGEAGTAGPP | GPP[113]GFAGEKGPSGEAGTAGP[113]P | GP(0.0108)P(0.8094)GFAGEKGP(0.0066)SGEAGTAGP(0.5866)P(0.5866) | 26.77 | 26.77 | 1.000 |
| 1873.847 | COL1A2 | 841 | 861 | GFAGEKGPSGEAGTAGPPGTP | GFAGEKGPSGEAGTAGPP[113]GTP[113] | GFAGEKGP(0.0113)SGEAGTAGP(0.0120)P(0.9883)GTP(0.9884) | 35.55 | 33.28 | 1.000 |
| 1883.851 | COL1A2 | 910 | 930 | GAVGSPGVNGAPGEAGRDGNP | GAVGSP[113]GVNGAP[113]GEAGRDGNP[113] | GAVGSP(1.0000)GVNGAP(1.0000)GEAGRDGNP(1.0000) | 21.03 | 8.55 | 1.000 |
| 1884.809 | COL1A2 | 910 | 930 | GAVGSPGVNGAPGEAGRDGNP | GAVGSP[113]GVN[115]GAP[113]GEAGRDGNP[113] | GAVGSP(1.0000)GVNGAP(1.0000)GEAGRDGNP(1.0000) | 30.28 | 15.40 | 1.000 |
| 1884.853 | COL1A2 | 349 | 369 | GPAGSKGESGNKGEPGSAGPQ | GPAGSKGESGNKGEP[113]GSAGPQ | GP(0.0058)AGSKGESGNKGEP(0.9870)GSAGP(0.0072)Q | 27.91 | 21.19 | 1.000 |
| 1889.839 | COL1A2 | 841 | 861 | GFAGEKGPSGEAGTAGPPGTP | GFAGEKGP[113]SGEAGTAGPP[113]GTP[113] | GFAGEKGP(0.9854)SGEAGTAGP(0.0440)P(0.9854)GTP(0.9852) | 22.10 | 19.84 | 0.987 |
| 1902.791 | COL1A2 | 1102 | 1119 | GVSGGGYDFGYDGDFYRA | GVSGGGYDFGYDGDFYRA | | 22.85 | 9.59 | 1.000 |
| 1912.871 | COL1A2 | 113 | 132 | FQGPAGEPGEPGQTGPAGAR | FQGPAGEP[113]GEP[113]GQTGPAGAR | FQGP(0.0099)AGEP(0.9894)GEP(0.9894)GQTGP(0.0113)AGAR | 22.49 | 13.47 | 0.997 |
| 1913.848 | COL1A2 | 113 | 132 | FQGPAGEPGEPGQTGPAGAR | FQGPAGEP[113]GEP[113]GQ[129]TGPAGAR | FQGP(0.0100)AGEP(0.9885)GEP(0.9884)GQTGP(0.0131)AGAR | 26.45 | 19.43 | 0.976 |
| 1924.857 | COL1A2 | 373 | 393 | GPSGEEGKRGPNGEAGSAGPP | GPSGEEGKRGPN[115]GEAGSAGPP[113] | GP(0.0034)SGEEGKRGP(0.0034)NGEAGSAGP(0.0044)P(0.9888) | 19.37 | 19.37 | 0.940 |
| 1933.937 | COL1A2 | 499 | 519 | GPTGDPGKNGDKGHAGLAGAR | GPTGDPGKN[115]GDKGHAGLAGAR | | 28.35 | 20.74 | 1.000 |

### Supplemental Table 1. Col1a1 peptide sequences from normal breast by LC-MS/MS.

| | | | | | | | | | |
|---|---|---|---|---|---|---|---|---|---|
| 1937.822 | COL1A2 | 916 | 936 | GVNGAPGEAGRDGNPGNDGPP | GVN[115]GAP[113]GEAGRDGNPGNDGPP[113] | GVNGAP(0.9836)GEAGRDGNP(0.0135)GNDGP(0.0195)P(0.9834) | 19.37 | 19.37 | 1.000 |
| 1938.963 | COL1A2 | 564 | 582 | AGEVGKPGERGLHGEFGLP | AGEVGKP[113]GERGLHGEFGLP[113] | AGEVGKP(1.0000)GERGLHGEFGLP(1.0000) | 22.68 | 15.04 | 0.975 |
| 1942.768 | COL1A2 | 1099 | 1117 | GPPGVSGGGYDFGYDGDFY | GP[113]PGVSGGGYDFGYDGDFY | GP(0.5000)P(0.5000)GVSGGGYDFGYDGDFY | 39.51 | 39.51 | 1.000 |
| 1949.936 | COL1A2 | 499 | 519 | GPTGDPGKNGDKGHAGLAGAR | GPTGDP[113]GKN[115]GDKGHAGLAGAR | GP(0.0102)TGDP(0.9898)GKNGDKGHAGLAGAR | 28.19 | 22.84 | 1.000 |
| 1952.826 | COL1A2 | 916 | 936 | GVNGAPGEAGRDGNPGNDGPP | GVNGAP[113]GEAGRDGNP[113]GNDGPP[113] | GVNGAP(0.9840)GEAGRDGNP(0.9840)GNDGP(0.0480)P(0.9840) | 18.59 | 18.59 | 1.000 |
| 1953.801 | COL1A2 | 916 | 936 | GVNGAPGEAGRDGNPGNDGPP | GVN[115]GAP[113]GEAGRDGNP[113]GNDGP[113]P | GVNGAP(0.7802)GEAGRDGNP(0.7801)GNDGP(0.7198)P(0.7198) | 18.62 | 18.62 | 0.979 |
| 1959.807 | COL1A2 | 1099 | 1117 | GPPGVSGGGYDFGYDGDFY | GP[113]P[113]GVSGGGYDFGYDGDFY | GP(1.0000)P(1.0000)GVSGGGYDFGYDGDFY | 21.47 | 10.82 | 1.000 |
| 1967.856 | COL1A2 | 109 | 129 | GPQGFQGPAGEPGEPGQTGPA | GPQGFQGPAGEP[113]GEP[113]GQTGPA | GP(0.0131)QGFQGP(0.0071)AGEP(0.9612)GEP(0.9557)GQTGP(0.0628)A | 22.68 | 22.65 | 0.963 |
| 1968.837 | COL1A2 | 109 | 129 | GPQGFQGPAGEPGEPGQTGPA | GPQGFQGPAGEP[113]GEP[113]GQ[129]TGPA | GP(0.0068)QGFQGP(0.0068)AGEP(0.9291)GEP(0.9128)GQTGP(0.1445)A | 31.11 | 26.71 | 1.000 |
| 1969.835 | COL1A2 | 109 | 129 | GPQGFQGPAGEPGEPGQTGPA | GPQGFQ[129]GPAGEP[113]GEP[113]GQ[129]TGPA | GP(0.0068)QGFQGP(0.0068)AGEP(0.9864)GEP(0.9858)GQTGP(0.0141)A | 29.07 | 29.07 | 1.000 |
| 1974.005 | COL1A2 | 457 | 477 | GNIGPAGKEGPVGLPGIDGRP | GNIGPAGKEGPVGLP[113]GIDGRP | GNIGP(0.0042)AGKEGP(0.0040)VGLP(0.9825)GIDGRP(0.0093) | 35.33 | 30.31 | 1.000 |
| 1974.031 | COL1A2 | 1060 | 1080 | GPAGKDGRTGHPGTVGPAGIR | GPAGKDGRTGHP[113]GTVGPAGIR | GP(0.0054)AGKDGRTGHP(0.9895)GTVGP(0.0051)AGIR | 27.46 | 16.72 | 1.000 |
| 1975.989 | COL1A2 | 820 | 840 | GPRGDQGPVGRTGEVGAVGPP | GPRGDQGPVGRTGEVGAVGPP[113] | GP(0.0036)RGDQGP(0.0034)VGRTGEVGAVGP(0.1073)P(0.8858) | 31.21 | 31.21 | 1.000 |
| 1976.977 | COL1A2 | 820 | 840 | GPRGDQGPVGRTGEVGAVGPP | GPRGDQ[129]GPVGRTGEVGAVGP[113]P | GP(0.0100)RGDQGP(0.0033)VGRTGEVGAVGP(0.9783)P(0.0084) | 24.17 | 24.17 | 0.989 |
| 1983.850 | COL1A2 | 109 | 129 | GPQGFQGPAGEPGEPGQTGPA | GPQGFQGP[113]AGEP[113]GEP[113]GQTGPA | GP(0.0645)QGFQGP(0.9721)AGEP(0.9729)GEP(0.9729)GQTGP(0.0176)A | 33.01 | 26.35 | 1.000 |
| 1990.001 | COL1A2 | 457 | 477 | GNIGPAGKEGPVGLPGIDGRP | GNIGPAGKEGPVGLP[113]GIDGRP[113] | GNIGP(0.0116)AGKEGP(0.0116)VGLP(0.9884)GIDGRP(0.9884) | 22.53 | 13.57 | 1.000 |
| 1993.942 | COL1A2 | 427 | 447 | GPAGVRGPNGDAGRPGEPGLM | GPAGVRGPNGDAGRP[113]GEP[113]GLM | GP(0.0086)AGVRGP(0.0100)NGDAGRP(0.9869)GEP(0.9871)GLM(0.0074) | 27.68 | 22.46 | 1.000 |
| 1993.972 | COL1A2 | 943 | 963 | GHKGERGYPGNIGPVGAAGAP | GHKGERGYP[113]GNIGPVGAAGAP[113] | GHKGERGYP(0.9894)GNIGP(0.0211)VGAAGAP(0.9895) | 22.01 | 14.65 | 1.000 |
| 1994.913 | COL1A2 | 427 | 447 | GPAGVRGPNGDAGRPGEPGLM | GPAGVRGPN[115]GDAGRP[113]GEP[113]GLM | GP(0.0072)AGVRGP(0.0073)NGDAGRP(0.9892)GEP(0.9892)GLM(0.0072) | 44.41 | 36.74 | 1.000 |
| 1994.966 | COL1A2 | 943 | 963 | GHKGERGYPGNIGPVGAAGAP | GHKGERGYP[113]GN[115]IGPVGAAGAP[113] | GHKGERGYP(0.9746)GNIGP(0.0505)VGAAGAP(0.9749) | 19.64 | 14.60 | 0.998 |
| 2009.931 | COL1A2 | 427 | 447 | GPAGVRGPNGDAGRPGEPGLM | GPAGVRGPNGDAGRP[113]GEP[113]GLM[147] | GP(0.0325)AGVRGP(0.0164)NGDAGRP(0.9837)GEP(0.9837)GLM(0.9837) | 18.61 | 10.85 | 0.976 |
| 2010.914 | COL1A2 | 427 | 447 | GPAGVRGPNGDAGRPGEPGLM | GPAGVRGPN[115]GDAGRP[113]GEP[113]GLM[147] | GP(0.0181)AGVRGP(0.0191)NGDAGRP(0.9876)GEP(0.9876)GLM(0.9876) | 23.48 | 15.64 | 0.965 |
| 2023.990 | COL1A2 | 298 | 321 | GNPGANGLTGAKGAAGLPGVAGAP | GNP[113]GAN[115]GLTGAKGAAGLP[113]GVAGAP[113] | GNP(1.0000)GANGLTGAKGAAGLP(1.0000)GVAGAP(1.0000) | 18.62 | 18.62 | 0.942 |
| 2026.912 | COL1A2 | 427 | 447 | GPAGVRGPNGDAGRPGEPGLM | GP[113]AGVRGPN[115]GDAGRP[113]GEP[113]GLM[147] | GP(0.8410)AGVRGP(0.6147)NGDAGRP(0.8481)GEP(0.8481)GLM(0.8481) | 18.42 | 16.69 | 0.990 |
| 2031.057 | COL1A2 | 1063 | 1083 | GKDGRTGHPGTVGPAGIRGPQ | GKDGRTGHP[113]GTVGPAGIRGPQ | GKDGRTGHP(0.9872)GTVGP(0.0069)AGIRGP(0.0058)Q | 27.83 | 17.60 | 1.000 |
| 2033.012 | COL1A2 | 820 | 841 | GPRGDQGPVGRTGEVGAVGPPG | GPRGDQGPVGRTGEVGAVGPP[113]G | GP(0.0036)RGDQGP(0.0036)VGRTGEVGAVGP(0.0765)P(0.9163)G | 23.27 | 23.27 | 0.958 |
| 2034.009 | COL1A2 | 820 | 841 | GPRGDQGPVGRTGEVGAVGPPG | GPRGDQ[129]GPVGRTGEVGAVGP[113]PG | GP(0.0034)RGDQGP(0.0033)VGRTGEVGAVGP(0.4967)P(0.4967)G | 27.16 | 27.16 | 1.000 |
| 2067.052 | COL1A2 | 649 | 669 | GIPGGKGEKGEPGLRGEIGNP | GIP[113]GGKGEKGEP[113]GLRGEIGNP[113] | GIP(1.0000)GGKGEKGEP(1.0000)GLRGEIGNP(1.0000) | 24.05 | 12.11 | 1.000 |
| 2077.038 | COL1A2 | 562 | 582 | GPAGEVGKPGERGLHGEFGLP | GPAGEVGKPGERGLHGEFGLP[113] | GP(0.0060)AGEVGKP(0.0059)GERGLHGEFGLP(0.9881) | 20.48 | 14.92 | 0.945 |
| 2093.011 | COL1A2 | 562 | 582 | GPAGEVGKPGERGLHGEFGLP | GPAGEVGKP[113]GERGLHGEFGLP[113] | GP(0.0199)AGEVGKP(0.9900)GERGLHGEFGLP(0.9901) | 31.25 | 15.91 | 1.000 |
| 2098.877 | COL1A2 | 1099 | 1118 | GPPGVSGGGYDFGYDGDFYR | GP[113]PGVSGGGYDFGYDGDFYR | GP(0.5000)P(0.5000)GVSGGGYDFGYDGDFYR | 18.36 | 18.36 | 0.998 |
| 2100.006 | COL1A2 | 950 | 973 | YPGNIGPVGAAGAPGPHGPVGPAG | YPGN[115]IGPVGAAGAP[113]GPHGPVGP[113]AG | YP(0.0046)GNIGP(0.0047)VGAAGAP(0.8140)GP(0.5192)HGP(0.0886)VGP(0.5689)AG | 27.07 | 27.07 | 1.000 |
| 2109.030 | COL1A2 | 562 | 582 | GPAGEVGKPGERGLHGEFGLP | GP[113]AGEVGKP[113]GERGLHGEFGLP[113] | GP(1.0000)AGEVGKP(1.0000)GERGLHGEFGLP(1.0000) | 20.58 | 14.24 | 0.978 |
| 2154.959 | COL1A2 | 113 | 135 | FQGPAGEPGEPGQTGPAGARGPA | FQGPAGEP[113]GEP[113]GQ[129]TGPAGARGP[113]A | FQGP(0.0220)AGEP(0.9860)GEP(0.9859)GQTGP(0.0202)AGARGP(0.9859)A | 18.69 | 14.52 | 0.979 |
| 2169.916 | COL1A2 | 1099 | 1119 | GPPGVSGGGYDFGYDGDFYRA | GPP[113]GVSGGGYDFGYDGDFYRA | GP(0.0135)P(0.9865)GVSGGGYDFGYDGDFYRA | 26.90 | 26.90 | 1.000 |
| 2170.914 | COL1A2 | 910 | 933 | GAVGSPGVNGAPGEAGRDGNPGND | GAVGSP[113]GVN[115]GAP[113]GEAGRDGNP[113]GND | GAVGSP(1.0000)GVNGAP(1.0000)GEAGRDGNP(1.0000)GND | 24.18 | 10.63 | 1.000 |
| 2175.057 | COL1A2 | 499 | 522 | GPTGDPGKNGDKGHAGLAGARGAP | GPTGDPGKN[115]GDKGHAGLAGARGAP[113] | GP(0.0064)TGDP(0.0069)GKNGDKGHAGLAGARGAP(0.9867) | 21.05 | 12.50 | 1.000 |
| 2180.087 | COL1A2 | 561 | 582 | SGPAGEVGKPGERGLHGEFGLP | SGPAGEVGKP[113]GERGLHGEFGLP[113] | SGP(0.0474)AGEVGKP(0.9762)GERGLHGEFGLP(0.9763) | 18.85 | 13.92 | 0.926 |
| 2191.035 | COL1A2 | 499 | 522 | GPTGDPGKNGDKGHAGLAGARGAP | GPTGDP[113]GKN[115]GDKGHAGLAGARGAP[113] | GP(0.0350)TGDP(0.9826)GKNGDKGHAGLAGARGAP(0.9825) | 19.19 | 14.55 | 0.995 |
| 2207.065 | COL1A2 | 706 | 729 | GPRGSPGERGEVGPAGPNGFAGPA | GPRGSPGERGEVGP[113]AGPNGFAGPA | GP(0.2615)RGSP(0.2489)GERGEVGP(0.4729)AGP(0.0123)NGFAGP(0.0045)A | 33.56 | 31.44 | 1.000 |
| 2207.113 | COL1A2 | 637 | 660 | GPSGLPGERGAAGIPGGKGEKGEP | GPSGLPGERGAAGIP[113]GGKGEKGEP[113] | GP(0.0159)SGLP(0.0148)GERGAAGIP(0.9848)GGKGEKGEP(0.9845) | 26.19 | 17.77 | 0.998 |
| 2223.104 | COL1A2 | 637 | 660 | GPSGLPGERGAAGIPGGKGEKGEP | GPSGLP[113]GERGAAGIP[113]GGKGEKGEP[113] | GP(0.1832)SGLP(0.9385)GERGAAGIP(0.9392)GGKGEKGEP(0.9391) | 22.66 | 18.12 | 0.962 |
| 2225.996 | COL1A2 | 424 | 447 | GASGPAGVRGPNGDAGRPGEPGLM | GASGPAGVRGPN[115]GDAGRP[113]GEP[113]GLM[147] | GASGP(0.0197)AGVRGP(0.0238)NGDAGRP(0.9854)GEP(0.9855)GLM(0.9855) | 27.22 | 19.04 | 1.000 |
| 2231.131 | COL1A2 | 454 | 477 | GSPGNIGPAGKEGPVGLPGIDGRP | GSP[113]GNIGPAGKEGPVGLP[113]GIDGRP | GSP(0.9834)GNIGP(0.0136)AGKEGP(0.0088)VGLP(0.9831)GIDGRP(0.0111) | 25.89 | 21.62 | 0.989 |
| 2232.110 | COL1A2 | 454 | 477 | GSPGNIGPAGKEGPVGLPGIDGRP | GSP[113]GN[115]IGPAGKEGPVGLP[113]GIDGRP | GSP(0.9798)GNIGP(0.0080)AGKEGP(0.0083)VGLP(0.9782)GIDGRP(0.0258) | 29.99 | 23.74 | 1.000 |
| 2247.112 | COL1A2 | 454 | 477 | GSPGNIGPAGKEGPVGLPGIDGRP | GSP[113]GNIGPAGKEGPVGLP[113]GIDGRP[113] | GSP(0.9881)GNIGP(0.0190)AGKEGP(0.0168)VGLP(0.9881)GIDGRP(0.9881) | 28.87 | 19.52 | 1.000 |
| 2248.082 | COL1A2 | 454 | 477 | GSPGNIGPAGKEGPVGLPGIDGRP | GSP[113]GN[115]IGPAGKEGPVGLP[113]GIDGRP[113] | GSP(0.9886)GNIGP(0.0170)AGKEGP(0.0172)VGLP(0.9886)GIDGRP(0.9886) | 24.69 | 17.92 | 1.000 |
| 2256.104 | COL1A2 | 1060 | 1083 | GPAGKDGRTGHPGTVGPAGIRGPQ | GPAGKDGRTGHP[113]GTVGPAGIRGPQ | GP(0.0050)AGKDGRTGHP(0.9829)GTVGP(0.0065)AGIRGP(0.0057)Q | 26.21 | 20.30 | 1.000 |
| 2257.130 | COL1A2 | 1060 | 1083 | GPAGKDGRTGHPGTVGPAGIRGPQ | GPAGKDGRTGHP[113]GTVGPAGIRGPQ[129] | GP(0.0127)AGKDGRTGHP(0.9799)GTVGP(0.0038)AGIRGP(0.0036)Q | 20.08 | 15.15 | 1.000 |
| 2264.130 | COL1A2 | 637 | 661 | GPSGLPGERGAAGIPGGKGEKGEPG | GPSGLPGERGAAGIP[113]GGKGEKGEP[113]G | GP(0.0742)SGLP(0.0196)GERGAAGIP(0.9559)GGKGEKGEP(0.9503)G | 23.82 | 18.86 | 1.000 |
| 2266.133 | COL1A2 | 646 | 669 | GAAGIPGGKGEKGEPGLRGEIGNP | GAAGIP[113]GGKGEKGEP[113]GLRGEIGNP[113] | GAAGIP(1.0000)GGKGEKGEP(1.0000)GLRGEIGNP(1.0000) | 23.07 | 11.32 | 1.000 |
| 2266.154 | COL1A2 | 651 | 673 | PGGKGEKGEPGLRGEIGNPGRDG | P[113]GGKGEKGEP[113]GLRGEIGNPGRDG | P(0.9887)GGKGEKGEP(0.9887)GLRGEIGNP(0.0227)GRDG | 27.60 | 15.64 | 1.000 |
| 2280.125 | COL1A2 | 637 | 661 | GPSGLPGERGAAGIPGGKGEKGEPG | GPSGLP[113]GERGAAGIP[113]GGKGEKGEP[113]G | GP(0.0371)SGLP(0.9877)GERGAAGIP(0.9876)GGKGEKGEP(0.9876)G | 23.30 | 16.31 | 0.994 |
| 2318.160 | COL1A2 | 562 | 585 | GPAGEVGKPGERGLHGEFGLPGPA | GPAGEVGKP[113]GERGLHGEFGLP[113]GPA | GP(0.0281)AGEVGKP(0.9778)GERGLHGEFGLP(0.9783)GP(0.0159)A | 28.19 | 25.63 | 1.000 |
| 2355.173 | COL1A2 | 139 | 162 | GKAGEDGHPGKPGRPGERGVVGPQ | GKAGEDGHPGKPGRP[113]GERGVVGPQ | GKAGEDGHP(0.0038)GKP(0.0050)GRP(0.9840)GERGVVGP(0.0072)Q | 24.18 | 24.18 | 1.000 |
| 2371.172 | COL1A2 | 139 | 162 | GKAGEDGHPGKPGRPGERGVVGPQ | GKAGEDGHP[113]GKPGRP[113]GERGVVGPQ | GKAGEDGHP(0.7631)GKP(0.5979)GRP(0.6255)GERGVVGP(0.0135)Q | 30.10 | 28.14 | 1.000 |
| 2372.159 | COL1A2 | 139 | 162 | GKAGEDGHPGKPGRPGERGVVGPQ | GKAGEDGHP[113]GKPGRP[113]GERGVVGPQ[129] | GKAGEDGHP(0.9753)GKP(0.0370)GRP(0.9744)GERGVVGP(0.0132)Q | 23.68 | 23.68 | 1.000 |
| 2381.098 | COL1A2 | 937 | 960 | GRDGQPGHKGERGYPGNIGPVGAA | GRDGQ[129]P[113]GHKGERGYP[113]GN[115]IGPVGAA | GRDGQP(0.9876)GHKGERGYP(0.9875)GNIGP(0.0249)VGAA | 19.50 | 12.65 | 0.998 |
| 2387.168 | COL1A2 | 139 | 162 | GKAGEDGHPGKPGRPGERGVVGPQ | GKAGEDGHP[113]GKP[113]GRP[113]GERGVVGPQ | GKAGEDGHP(0.9339)GKP(0.9333)GRP(0.9331)GERGVVGP(0.1997)Q | 30.23 | 22.99 | 1.000 |
| 2388.169 | COL1A2 | 139 | 162 | GKAGEDGHPGKPGRPGERGVVGPQ | GKAGEDGHP[113]GKP[113]GRP[113]GERGVVGPQ[129] | GKAGEDGHP(0.8960)GKP(0.8944)GRP(0.8944)GERGVVGP(0.3153)Q | 18.80 | 17.00 | 0.983 |
| 2396.107 | COL1A2 | 344 | 369 | LVGEPGPAGSKGESGNKGEPGSAGPQ | LVGEP[113]GPAGSKGESGNKGEP[113]GSAGPQ | LVGEP(0.7826)GP(0.3862)AGSKGESGNKGEP(0.8220)GSAGP(0.0092)Q | 31.76 | 27.44 | 1.000 |
| 2396.198 | COL1A2 | 652 | 675 | GGKGEKGEPGLRGEIGNPGRDGAR | GGKGEKGEP[113]GLRGEIGNP[113]GRDGAR | GGKGEKGEP(1.0000)GLRGEIGNP(1.0000)GRDGAR | 29.04 | 15.23 | 1.000 |
| 2397.088 | COL1A2 | 344 | 369 | LVGEPGPAGSKGESGNKGEPGSAGPQ | LVGEP[113]GP[113]AGSKGESGN[115]KGEPGSAGPQ | LVGEP(0.7667)GP(0.7275)AGSKGESGNKGEP(0.4838)GSAGP(0.0220)Q | 20.00 | 20.00 | 0.983 |
| 2412.111 | COL1A2 | 344 | 369 | LVGEPGPAGSKGESGNKGEPGSAGPQ | LVGEP[113]GP[113]AGSKGESGNKGEP[113]GSAGPQ | LVGEP(0.9737)GP(0.9737)AGSKGESGNKGEP(0.9742)GSAGP(0.0784)Q | 18.62 | 12.52 | 0.947 |
| 2421.025 | COL1A2 | 910 | 936 | GAVGSPGVNGAPGEAGRDGNPGNDGPP | GAVGSP[113]GVNGAP[113]GEAGRDGNPGNDGP[113]P | GAVGSP(0.8016)GVNGAP(0.8017)GEAGRDGNP(0.0243)GNDGP(0.6861)P(0.6861) | 26.78 | 26.78 | 1.000 |
| 2422.033 | COL1A2 | 910 | 936 | GAVGSPGVNGAPGEAGRDGNPGNDGPP | GAVGSP[113]GVN[115]GAP[113]GEAGRDGNPGNDGPP[113] | GAVGSP(0.9874)GVNGAP(0.9874)GEAGRDGNP(0.0197)GNDGP(0.0182)P(0.9873) | 53.62 | 53.62 | 1.000 |
| 2423.033 | COL1A2 | 910 | 936 | GAVGSPGVNGAPGEAGRDGNPGNDGPP | GAVGSP[113]GVN[115]GAP[113]GEAGRDGNPGN[115]DGP[113]P | GAVGSP(0.8049)GVNGAP(0.8049)GEAGRDGNP(0.0139)GNDGP(0.6881)P(0.6881) | 38.64 | 38.64 | 1.000 |
| 2437.041 | COL1A2 | 910 | 936 | GAVGSPGVNGAPGEAGRDGNPGNDGPP | GAVGSP[113]GVNGAP[113]GEAGRDGNP[113]GNDGPP[113] | GAVGSP(0.9827)GVNGAP(0.9827)GEAGRDGNP(0.9827)GNDGP(0.0692)P(0.9827) | 25.93 | 25.93 | 1.000 |
| 2438.015 | COL1A2 | 910 | 936 | GAVGSPGVNGAPGEAGRDGNPGNDGPP | GAVGSP[113]GVN[115]GAPGEAGRDGNP[113]GNDGP[113]P[113] | GAVGSP(0.8385)GVNGAP(0.6731)GEAGRDGNP(0.8351)GNDGP(0.8267)P(0.8267) | 24.14 | 24.14 | 1.000 |
| 2453.148 | COL1A2 | 343 | 369 | GLVGEPGPAGSKGESGNKGEPGSAGPQ | GLVGEP[113]GPAGSKGESGNKGEP[113]GSAGPQ | GLVGEP(0.9109)GP(0.1587)AGSKGESGNKGEP(0.9187)GSAGP(0.0118)Q | 30.26 | 27.93 | 1.000 |
| 2454.119 | COL1A2 | 343 | 369 | GLVGEPGPAGSKGESGNKGEPGSAGPQ | GLVGEP[113]GPAGSKGESGN[115]KGEP[113]GSAGPQ | GLVGEP(0.9770)GP(0.0338)AGSKGESGNKGEP(0.9772)GSAGP(0.0120)Q | 29.34 | 27.21 | 1.000 |
| 2459.093 | COL1A2 | 772 | 798 | GPAGSRGDGGPPGMTGFPGAAGRTGPP | GPAGSRGDGGPP[113]GM[147]TGFP[113]GAAGRTGP[113]P[113] | GP(0.1304)AGSRGDGGP(0.3347)P(0.9049)GM(0.9049)TGFP(0.9085)GAAGRTGP(0.9084)P(0.908 | 18.69 | 18.69 | 0.994 |

| | | | | | | | | | |
|---|---|---|---|---|---|---|---|---|---|
| 2469.140 | COL1A2 | 343 | 369 | GLVGEPGPAGSKGESGNKGEPGSAGPQ | GLVGEP[113]GP[113]AGSKGESGNKGEP[113]GSAGPQ | GLVGEP(0.9878)GP(0.9878)AGSKGESGNKGEP(0.9881)GSAGP(0.0362)Q | 19.30 | 12.20 | 0.976 |
| 2473.265 | COL1A2 | 877 | 903 | GSRGERGLPGVAGAVGEPGPLGIAGPP | GSRGERGLP[113]GVAGAVGEP[113]GPLGIAGP[113]P | GSRGERGLP(0.7900)GVAGAVGEP(0.7889)GP(0.0697)LGIAGP(0.6757)P(0.6757) | 28.22 | 28.22 | 1.000 |
| 2477.105 | COL1A2 | 109 | 135 | GPQGFQGPAGEPGEPGQTGPAGARGPA | GPQGFQGPAGEP[113]GEP[113]GQTGPAGARGPA | GP(0.0187)QGFQGP(0.0113)AGEP(0.9769)GEP(0.9776)GQTGP(0.0066)AGARGP(0.0088)A | 31.37 | 29.22 | 1.000 |
| 2493.105 | COL1A2 | 109 | 135 | GPQGFQGPAGEPGEPGQTGPAGARGPA | GPQGFQGPAGEP[113]GEP[113]GQTGPAGARGP[113]A | GP(0.0209)QGFQGP(0.0142)AGEP(0.9808)GEP(0.9802)GQTGP(0.0235)AGARGP(0.9804)A | 30.82 | 26.48 | 1.000 |
| 2494.112 | COL1A2 | 109 | 135 | GPQGFQGPAGEPGEPGQTGPAGARGPA | GPQGFQGPAGEP[113]GEP[113]GQ[129]TGPAGARGP[113]A | GP(0.0151)QGFQGP(0.0127)AGEP(0.9810)GEP(0.9808)GQTGP(0.0301)AGARGP(0.9803)A | 22.40 | 16.95 | 0.971 |
| 2509.194 | COL1A2 | 421 | 447 | GSRGASGPAGVRGPNGDAGRPGEPGLM | GSRGASGPAGVRGPNGDAGRP[113]GEP[113]GLM | GSRGASGP(0.3010)AGVRGP(0.1787)NGDAGRP(0.7447)GEP(0.7664)GLM(0.0092) | 26.63 | 22.19 | 1.000 |
| 2525.181 | COL1A2 | 421 | 447 | GSRGASGPAGVRGPNGDAGRPGEPGLM | GSRGASGPAGVRGPNGDAGRP[113]GEP[113]GLM[147] | GSRGASGP(0.6596)AGVRGP(0.1260)NGDAGRP(0.6659)GEP(0.7735)GLM(0.7750) | 19.86 | 19.07 | 0.995 |
| 2592.305 | COL1A2 | 643 | 669 | GERGAAGIPGGKGEKGEPGLRGEIGNP | GERGAAGIP[113]GGKGEKGEPGLRGEIGNP[113] | GERGAAGIP(0.9865)GGKGEKGEP(0.0271)GLRGEIGNP(0.9865) | 26.93 | 21.17 | 1.000 |
| 2608.278 | COL1A2 | 643 | 669 | GERGAAGIPGGKGEKGEPGLRGEIGNP | GERGAAGIP[113]GGKGEKGEP[113]GLRGEIGNP[113] | GERGAAGIP(1.0000)GGKGEKGEP(1.0000)GLRGEIGNP(1.0000) | 26.86 | 14.79 | 1.000 |
| 2622.320 | COL1A2 | 136 | 162 | GPPGKAGEDGHPGKPGRPGERGVVGPQ | GP[113]PGKAGEDGHPGKPGRP[113]GERGVVGPQ | GP(0.8867)P(0.0093)GKAGEDGHP(0.0050)GKP(0.0067)GRP(0.8006)GERGVVGP(0.2917)Q | 20.92 | 19.40 | 1.000 |
| 2638.296 | COL1A2 | 136 | 162 | GPPGKAGEDGHPGKPGRPGERGVVGPQ | GPP[113]GKAGEDGHP[113]GKPGRP[113]GERGVVGPQ | GP(0.0513)P(0.8400)GKAGEDGHP(0.8412)GKP(0.0131)GRP(0.7842)GERGVVGP(0.4702)Q | 24.43 | 24.43 | 1.000 |
| 2654.298 | COL1A2 | 136 | 162 | GPPGKAGEDGHPGKPGRPGERGVVGPQ | GPP[113]GKAGEDGHP[113]GKP[113]GRP[113]GERGVVGPQ | GP(0.0225)P(0.9843)GKAGEDGHP(0.9843)GKP(0.9842)GRP(0.9841)GERGVVGP(0.0406)Q | 18.06 | 18.06 | 1.000 |
| 2728.440 | COL1A2 | 168 | 195 | PGTPGLPGFKGIRGHNGLDGLKGQPGAP | P[113]GTPGLPGFKGIRGHNGLDGLKGQ[129]P[113]GAP | P(0.8045)GTP(0.0146)GLP(0.0146)GFKGIRGHNGLDGLKGQP(0.5832)GAP(0.5832) | 20.17 | 20.17 | 0.983 |
| 2803.401 | COL1A2 | 1062 | 1091 | AGKDGRTGHPGTVGPAGIRGPQGHQGPAGP | AGKDGRTGHPGTVGP[113]AGIRGPQGHQGPAGP | AGKDGRTGHP(0.0122)GTVGP(0.9301)AGIRGP(0.0192)QGHQGP(0.0214)AGP(0.0171) | 18.28 | 17.81 | 0.972 |
| 2878.476 | COL1A2 | 646 | 675 | GAAGIPGGKGEKGEPGLRGEIGNPGRDGAR | GAAGIP[113]GGKGEKGEP[113]GLRGEIGNP[113]GRDGAR | GAAGIP(1.0000)GGKGEKGEP(1.0000)GLRGEIGNP(1.0000)GRDGAR | 24.17 | 11.15 | 1.000 |

| M+H | Gene | Domain Start | Domain End | Peptide | Modified Peptide [113] means HYP | Sequence with HYP probability PM:15.9949 | Hyperscore | Nextscore | Peptide Prophet Probability |
|---|---|---|---|---|---|---|---|---|---|
| 741.452 | COL3A1 | 946 | 953 | PLGIAGIT | PLGIAGIT | | 18.07 | 13.03 | 0.984 |
| 743.370 | COL3A1 | 702 | 710 | GPEGGKGAA | GPEGGKGAA | | 19.36 | 12.34 | 0.992 |
| 762.340 | COL3A1 | 867 | 875 | GGPGAAGFP | GGP[113]GAAGFP[113] | GGP(1.0000)GAAGFP(1.0000) | 19.14 | 13.16 | 0.966 |
| 774.344 | COL3A1 | 1215 | 1221 | GFAPYYG | GFAPYYG | | 18.41 | 11.17 | 0.999 |
| 788.377 | COL3A1 | 1002 | 1010 | GLAGTAGEP | GLAGTAGEP[113] | GLAGTAGEP(1.0000) | 19.37 | 12.39 | 0.987 |
| 798.467 | COL3A1 | 945 | 953 | GPLGIAGIT | GPLGIAGIT | | 21.90 | 16.36 | 0.960 |
| 806.438 | COL3A1 | 164 | 172 | VAVGGLAGY | VAVGGLAGY | | 19.21 | 11.33 | 1.000 |
| 815.428 | COL3A1 | 639 | 647 | GLQGLPGTG | GLQGLP[113]GTG | GLQGLP(1.0000)GTG | 18.05 | 14.96 | 0.964 |
| 830.396 | COL3A1 | 1125 | 1133 | GQQGAIGSP | GQQGAIGSP[113] | GQQGAIGSP(1.0000) | 21.06 | 14.63 | 1.000 |
| 832.381 | COL3A1 | 336 | 344 | GPPGTAGFP | GPP[113]GTAGFP[113] | GP(0.0309)P(0.9845)GTAGFP(0.9846) | 22.18 | 21.72 | 0.993 |
| 844.421 | COL3A1 | 252 | 260 | GPAGIPGFP | GPAGIP[113]GFP[113] | GP(0.0208)AGIP(0.9896)GFP(0.9896) | 20.63 | 20.63 | 0.922 |
| 845.400 | COL3A1 | 1002 | 1011 | GLAGTAGEPG | GLAGTAGEP[113]G | GLAGTAGEP(1.0000)G | 19.48 | 10.80 | 0.993 |
| 855.489 | COL3A1 | 945 | 954 | GPLGIAGITG | GPLGIAGITG | | 22.78 | 15.51 | 0.956 |
| 858.436 | COL3A1 | 972 | 980 | GPQGVKGES | GPQGVKGES | | 18.74 | 10.03 | 0.997 |
| 891.384 | COL3A1 | 195 | 203 | GSPGYQGPP | GSP[113]GYQGPP[113] | GSP(0.9879)GYQGP(0.0223)P(0.9898) | 19.25 | 17.43 | 0.994 |
| 914.384 | COL3A1 | 644 | 653 | PGTGGPPGEN | P[113]GTGGPP[113]GEN | P(0.9886)GTGGP(0.0228)P(0.9886)GEN | 21.08 | 17.29 | 0.996 |
| 919.467 | COL3A1 | 164 | 173 | VAVGGLAGYP | VAVGGLAGYP[113] | VAVGGLAGYP(1.0000) | 24.34 | 11.63 | 1.000 |
| 921.412 | COL3A1 | 573 | 581 | GQPGVMGFP | GQPGVM[147]GFP[113] | GQP(0.0493)GVM(0.9752)GFP(0.9755) | 18.50 | 13.65 | 1.000 |
| 923.446 | COL3A1 | 840 | 851 | GVAGPPGGSGPA | GVAGPPGGSGPA | | 19.78 | 10.04 | 0.983 |
| 926.524 | COL3A1 | 945 | 955 | GPLGIAGITGA | GPLGIAGITGA | | 25.32 | 19.37 | 1.000 |
| 937.401 | COL3A1 | 573 | 581 | GQPGVMGFP | GQP[113]GVM[147]GFP[113] | GQP(1.0000)GVM(1.0000)GFP(1.0000) | 22.22 | 11.69 | 1.000 |
| 976.506 | COL3A1 | 163 | 173 | GVAVGGLAGYP | GVAVGGLAGYP[113] | GVAVGGLAGYP(1.0000) | 23.02 | 17.83 | 0.983 |
| 985.579 | COL3A1 | 947 | 957 | LGIAGITGARG | LGIAGITGARG | | 21.82 | 14.55 | 0.998 |
| 999.591 | COL3A1 | 949 | 959 | IAGITGARGLA | IAGITGARGLA | | 19.73 | 10.11 | 0.981 |
| 1019.437 | COL3A1 | 864 | 875 | GSPGGPGAAGFP | GSP[113]GGP[113]GAAGFP[113] | GSP(1.0000)GGP(1.0000)GAAGFP(1.0000) | 22.02 | 14.34 | 0.987 |
| 1025.518 | COL3A1 | 640 | 650 | LQGLPGTGGPP | LQGLP[113]GTGGPP[113] | LQGLP(0.9884)GTGGP(0.0232)P(0.9884) | 19.31 | 17.23 | 0.995 |
| 1025.610 | COL3A1 | 946 | 956 | PLGIAGITGAR | PLGIAGITGAR | | 25.85 | 11.85 | 0.998 |
| 1027.499 | COL3A1 | 780 | 791 | GEGGAPGLPGIA | GEGGAP[113]GLP[113]GIA | GEGGAP(1.0000)GLP(1.0000)GIA | 19.12 | 11.64 | 0.971 |
| 1042.502 | COL3A1 | 636 | 646 | GPQGLQGLPGT | GPQ[129]GLQ[129]GLP[113]GT | GP(0.0115)QGLQGLP(0.9885)GT | 20.38 | 11.81 | 1.000 |
| 1055.516 | COL3A1 | 1122 | 1133 | GPAGQQGAIGSP | GPAGQQGAIGSP[113] | GP(0.0113)AGQQGAIGSP(0.9887) | 18.12 | 11.84 | 0.991 |
| 1056.491 | COL3A1 | 1122 | 1133 | GPAGQQGAIGSP | GPAGQ[129]QGAIGSP[113] | GP(0.0111)AGQQGAIGSP(0.9889) | 18.04 | 16.02 | 0.994 |
| 1056.617 | COL3A1 | 948 | 959 | GIAGITGARGLA | GIAGITGARGLA | | 18.56 | 9.59 | 0.996 |
| 1059.505 | COL3A1 | 1003 | 1013 | LAGTAGEPGRD | LAGTAGEP[113]GRD | LAGTAGEP(1.0000)GRD | 20.11 | 9.86 | 1.000 |
| 1063.537 | COL3A1 | 162 | 173 | SGVAVGGLAGYP | SGVAVGGLAGYP[113] | SGVAVGGLAGYP(1.0000) | 29.31 | 13.55 | 1.000 |
| 1070.471 | COL3A1 | 624 | 635 | GPGGDKGDTGPP | GPGGDKGDTGPP[113] | GP(0.0056)GGDKGDTGP(0.0064)P(0.9880) | 18.66 | 18.66 | 0.979 |
| 1073.486 | COL3A1 | 336 | 347 | GPPGTAGFPGSP | GP[113]PGTAGFPGSP[113] | GP(0.6149)P(0.6149)GTAGFP(0.0099)GSP(0.7602) | 21.88 | 21.88 | 1.000 |
| 1082.531 | COL3A1 | 639 | 650 | GLQGLPGTGGPP | GLQGLP[113]GTGGPP[113] | GLQGLP(0.9891)GTGGP(0.0218)P(0.9891) | 30.12 | 27.94 | 1.000 |
| 1082.617 | COL3A1 | 945 | 956 | GPLGIAGITGAR | GPLGIAGITGAR | | 32.53 | 17.65 | 1.000 |
| 1083.522 | COL3A1 | 639 | 650 | GLQGLPGTGGPP | GLQ[129]GLP[113]GTGGP[113]P | GLQGLP(0.7197)GTGGP(0.6401)P(0.6401) | 18.33 | 18.33 | 0.963 |
| 1084.486 | COL3A1 | 894 | 905 | GPSGSPGKDGPP | GPSGSP[113]GKDGP[113]P | GP(0.0087)SGSP(0.7649)GKDGP(0.6132)P(0.6132) | 24.71 | 24.71 | 0.961 |
| 1085.600 | COL3A1 | 250 | 260 | IKGPAGIPGFP | IKGPAGIP[113]GFP[113] | IKGP(0.0198)AGIP(0.9901)GFP(0.9901) | 18.57 | 12.65 | 0.988 |
| 1088.502 | COL3A1 | 717 | 728 | GAAGTPGLQGMP | GAAGTP[113]GLQGMP[113] | GAAGTP(0.9885)GLQGM(0.0228)P(0.9887) | 18.81 | 16.58 | 0.993 |
| 1089.471 | COL3A1 | 336 | 347 | GPPGTAGFPGSP | GPP[113]GTAGFP[113]GSP[113] | GP(0.0397)P(0.9868)GTAGFP(0.9868)GSP(0.9868) | 22.10 | 20.09 | 0.976 |
| 1097.555 | COL3A1 | 636 | 647 | GPQGLQGLPGTG | GPQGLQGLP[113]GTG | GP(0.0103)QGLQGLP(0.9897)GTG | 21.69 | 9.13 | 1.000 |
| 1098.535 | COL3A1 | 636 | 647 | GPQGLQGLPGTG | GPQGLQ[129]GLP[113]GTG | GP(0.0112)QGLQGLP(0.9888)GTG | 20.50 | 13.14 | 1.000 |
| 1101.468 | COL3A1 | 468 | 479 | GSPGEPGANGLP | GSP[113]GEP[113]GAN[115]GLP[113] | GSP(1.0000)GEP(1.0000)GANGLP(1.0000) | 19.37 | 12.41 | 0.996 |
| 1101.587 | COL3A1 | 250 | 260 | IKGPAGIPGFP | IKGP[113]AGIP[113]GFP[113] | IKGP(1.0000)AGIP(1.0000)GFP(1.0000) | 18.11 | 11.82 | 0.974 |
| 1106.521 | COL3A1 | 1146 | 1157 | GPPGKDGTSGHP | GPPGKDGTSGHP | | 18.57 | 12.13 | 0.965 |
| 1109.563 | COL3A1 | 495 | 506 | GPNGIPGEKGPA | GPNGIP[113]GEKGPA | GP(0.0050)NGIP(0.9865)GEKGP(0.0085)A | 19.22 | 12.62 | 0.985 |
| 1110.540 | COL3A1 | 495 | 506 | GPNGIPGEKGPA | GPN[115]GIP[113]GEKGPA | GP(0.0056)NGIP(0.9888)GEKGP(0.0056)A | 20.14 | 13.38 | 0.965 |
| 1116.525 | COL3A1 | 1002 | 1013 | GLAGTAGEPGRD | GLAGTAGEP[113]GRD | GLAGTAGEP(1.0000)GRD | 24.38 | 12.73 | 0.995 |
| 1117.520 | COL3A1 | 1059 | 1070 | GKSGDRGESGPA | GKSGDRGESGPA | | 18.63 | 11.75 | 0.959 |
| 1125.566 | COL3A1 | 750 | 761 | GADGVPGKDGPR | GADGVPGKDGPR | | 21.60 | 12.59 | 0.999 |
| 1127.541 | COL3A1 | 225 | 236 | GPAGKDGESGRP | GPAGKDGESGRP | | 19.80 | 11.22 | 1.000 |
| 1128.520 | COL3A1 | 984 | 995 | GANGLSGERGPP | GAN[115]GLSGERGPP[113] | GANGLSGERGP(0.0489)P(0.9511) | 25.69 | 23.84 | 0.997 |
| 1138.556 | COL3A1 | 1149 | 1160 | GKDGTSGHPGPI | GKDGTSGHP[113]GPI | GKDGTSGHP(0.9841)GP(0.0159)I | 18.74 | 15.59 | 0.990 |
| 1139.636 | COL3A1 | 945 | 957 | GPLGIAGITGARG | GPLGIAGITGARG | | 32.60 | 8.57 | 1.000 |
| 1142.618 | COL3A1 | 249 | 260 | GIKGPAGIPGFP | GIKGPAGIP[113]GFP[113] | GIKGP(0.0204)AGIP(0.9898)GFP(0.9898) | 20.63 | 9.50 | 1.000 |
| 1143.545 | COL3A1 | 225 | 236 | GPAGKDGESGRP | GPAGKDGESGRP[113] | GP(0.0100)AGKDGESGRP(0.9900) | 18.38 | 9.57 | 0.998 |
| 1154.537 | COL3A1 | 606 | 617 | GPPGKNGETGPQ | GPP[113]GKNGETGPQ | GP(0.0051)P(0.9899)GKNGETGP(0.0051)Q | 20.48 | 17.29 | 0.968 |
| 1155.524 | COL3A1 | 606 | 617 | GPPGKNGETGPQ | GPP[113]GKN[115]GETGPQ | GP(0.0058)P(0.9887)GKNGETGP(0.0055)Q | 24.20 | 23.00 | 0.993 |
| 1156.588 | COL3A1 | 972 | 983 | GPQGVKGESGKP | GPQGVKGESGKP[113] | GP(0.0103)QGVKGESGKP(0.9897) | 24.61 | 9.97 | 1.000 |
| 1158.614 | COL3A1 | 249 | 260 | GIKGPAGIPGFP | GIKGP[113]AGIP[113]GFP[113] | GIKGP(1.0000)AGIP(1.0000)GFP(1.0000) | 18.07 | 12.86 | 0.980 |
| 1161.571 | COL3A1 | 480 | 491 | GAAGERGAPGFR | GAAGERGAP[113]GFR | GAAGERGAP(1.0000)GFR | 25.22 | 8.34 | 1.000 |
| 1162.512 | COL3A1 | 726 | 737 | GMPGERGGLGSP | GM[147]P[113]GERGGLGSP[113] | GM(1.0000)P(1.0000)GERGGLGSP(1.0000) | 18.22 | 9.17 | 0.998 |
| 1169.703 | COL3A1 | 947 | 959 | LGIAGITGARGLA | LGIAGITGARGLA | | 20.84 | 9.72 | 0.999 |
| 1172.510 | COL3A1 | 795 | 806 | GSPGERGETGPP | GSP[113]GERGETGPP[113] | GSP(0.9874)GERGETGP(0.0253)P(0.9874) | 18.76 | 18.76 | 0.947 |
| 1173.544 | COL3A1 | 1002 | 1014 | GLAGTAGEPGRDG | GLAGTAGEP[113]GRDG | GLAGTAGEP(1.0000)GRDG | 21.27 | 10.74 | 0.992 |
| 1184.602 | COL3A1 | 855 | 866 | GPQGVKGERGSP | GPQGVKGERGSP[113] | GP(0.0100)QGVKGERGSP(0.9900) | 18.45 | 6.76 | 1.000 |
| 1231.548 | COL3A1 | 1003 | 1015 | LAGTAGEPGRDGN | LAGTAGEP[113]GRDGN[115] | LAGTAGEP(1.0000)GRDGN | 22.99 | 10.61 | 1.000 |
| 1232.573 | COL3A1 | 570 | 581 | GPRGQPGVMGFP | GPRGQ[129]PGVM[147]GFP[113] | GP(0.0119)RGQP(0.0119)GVM(0.9881)GFP(0.9881) | 18.19 | 13.97 | 0.983 |
| 1248.567 | COL3A1 | 570 | 581 | GPRGQPGVMGFP | GPRGQ[129]P[113]GVM[147]GFP[113] | GP(0.0367)RGQP(0.9877)GVM(0.9878)GFP(0.9878) | 21.43 | 14.99 | 0.998 |
| 1252.732 | COL3A1 | 945 | 958 | GPLGIAGITGARGL | GPLGIAGITGARGL | | 21.27 | 9.83 | 1.000 |
| 1266.752 | COL3A1 | 946 | 959 | PLGIAGITGARGLA | PLGIAGITGARGLA | | 28.85 | 13.45 | 1.000 |
| 1287.586 | COL3A1 | 1002 | 1015 | GLAGTAGEPGRDGN | GLAGTAGEP[113]GRDGN | GLAGTAGEP(1.0000)GRDGN | 19.35 | 10.02 | 0.993 |
| 1288.573 | COL3A1 | 1002 | 1015 | GLAGTAGEPGRDGN | GLAGTAGEP[113]GRDGN[115] | GLAGTAGEP(1.0000)GRDGN | 25.28 | 11.94 | 1.000 |
| 1295.582 | COL3A1 | 771 | 785 | GPAGQPGDKGEGGAP | GPAGQ[129]PGDKGEGGAP | | 22.17 | 15.21 | 0.937 |
| 1311.658 | COL3A1 | 777 | 791 | GDKGEGGAPGLPGIA | GDKGEGGAPGLP[113]GIA | GDKGEGGAP(0.0143)GLP(0.9857)GIA | 22.95 | 18.09 | 0.996 |
| 1323.756 | COL3A1 | 945 | 959 | GPLGIAGITGARGLA | GPLGIAGITGARGLA | | 42.31 | 20.20 | 1.000 |
| 1325.588 | COL3A1 | 621 | 635 | GPTGPGGDKGDTGPP | GPTGPGGDKGDTGPP[113] | GP(0.0052)TGP(0.0053)GGDKGDTGP(0.0036)P(0.9859) | 22.42 | 22.42 | 0.996 |

| | | | | | | | | | |
|---|---|---|---|---|---|---|---|---|---|
| 1325.618 | COL3A1 | 640 | 653 | LQGLPGTGGPPGEN | LQGLP[113]GTGGPP[113]GEN | LQGLP(0.9127)GTGGP(0.1765)P(0.9107)GEN | 18.59 | 18.50 | 0.953 |
| 1327.568 | COL3A1 | 540 | 554 | GSPGGPGSDGKPGPP | GSP[113]GGP[113]GSDGKP[113]GP[113]P | GSP(0.8205)GGP(0.8204)GSDGKP(0.8161)GP(0.7715)P(0.7715) | 21.32 | 21.32 | 0.987 |
| 1327.636 | COL3A1 | 393 | 407 | GGKGEMGPAGIPGAP | GGKGEMGPAGIP[113]GAP[113] | GGKGEM(0.0132)GP(0.0137)AGIP(0.9865)GAP(0.9865) | 24.68 | 17.34 | 0.979 |
| 1327.652 | COL3A1 | 777 | 791 | GDKGEGGAPGLPGIA | GDKGEGGAP[113]GLP[113]GIA | GDKGEGGAP(1.0000)GLP(1.0000)GIA | 21.62 | 14.63 | 0.989 |
| 1329.663 | COL3A1 | 481 | 494 | AAGERGAPGFRGPA | AAGERGAP[113]GFRGPA | AAGERGAP(0.9761)GFRGP(0.0239)A | 18.20 | 13.87 | 1.000 |
| 1336.646 | COL3A1 | 747 | 761 | GGPGADGVPGKDGPR | GGPGADGVPGKDGPR | | 24.92 | 8.06 | 1.000 |
| 1341.599 | COL3A1 | 519 | 533 | GAAGEPGRDGVPGGP | GAAGEP[113]GRDGVP[113]GGP[113] | GAAGEP(1.0000)GRDGVP(1.0000)GGP(1.0000) | 19.94 | 13.16 | 0.973 |
| 1342.618 | COL3A1 | 915 | 929 | GSPGVSGPKGDAGQP | GSP[113]GVSGPKGDAGQP[113] | GSP(0.9895)GVSGP(0.0211)KGDAGQP(0.9895) | 23.28 | 13.72 | 1.000 |
| 1343.605 | COL3A1 | 1003 | 1016 | LAGTAGEPGRDGNP | LAGTAGEP[113]GRDGNP[113] | LAGTAGEP(1.0000)GRDGNP(1.0000) | 28.66 | 13.17 | 1.000 |
| 1343.628 | COL3A1 | 393 | 407 | GGKGEMGPAGIPGAP | GGKGEM[147]GPAGIP[113]GAP[113] | GGKGEM(0.9789)GP(0.0630)AGIP(0.9790)GAP(0.9790) | 22.52 | 17.08 | 0.993 |
| 1352.640 | COL3A1 | 747 | 761 | GGPGADGVPGKDGPR | GGP[113]GADGVPGKDGPR | GGP(0.9884)GADGVP(0.0059)GKDGP(0.0057)R | 20.60 | 11.10 | 1.000 |
| 1354.630 | COL3A1 | 672 | 686 | GGKGDAGAPGERGPP | GGKGDAGAP[113]GERGP[113]P | GGKGDAGAP(0.7195)GERGP(0.6403)P(0.6403) | 19.44 | 19.44 | 0.950 |
| 1358.622 | COL3A1 | 429 | 443 | GGAGEPGKNGAKGEP | GGAGEP[113]GKN[115]GAKGEP[113] | GGAGEP(1.0000)GKNGAKGEP(1.0000) | 18.20 | 13.66 | 0.971 |
| 1361.590 | COL3A1 | 387 | 401 | GINGSPGGKGEMGPA | GIN[115]GSP[113]GGKGEM[147]GPA | GINGSP(0.9890)GGKGEM(0.9890)GP(0.0220)A | 24.40 | 18.79 | 0.985 |
| 1361.601 | COL3A1 | 861 | 875 | GERGSPGGPGAAGFP | GERGSP[113]GGP[113]GAAGFP[113] | GERGSP(1.0000)GGP(1.0000)GAAGFP(1.0000) | 23.42 | 15.01 | 0.995 |
| 1364.674 | COL3A1 | 636 | 650 | GPQGLQGLPGTGGPP | GPQGLQGLP[113]GTGGP[113]P | GP(0.0099)QGLQGLP(0.7648)GTGGP(0.6126)P(0.6126) | 23.96 | 21.80 | 1.000 |
| 1365.664 | COL3A1 | 636 | 650 | GPQGLQGLPGTGGPP | GPQGLQ[129]GLP[113]GTGGPP[113] | GP(0.0129)QGLQGLP(0.9720)GTGGP(0.0444)P(0.9706) | 27.16 | 25.05 | 0.996 |
| 1371.613 | COL3A1 | 714 | 728 | GPPGAAGTPGLQGMP | GP[113]PGAAGTP[113]GLQGM[147]P[113] | GP(0.8600)P(0.5465)GAAGTP(0.8646)GLQGM(0.8645)P(0.8645) | 19.83 | 19.83 | 0.994 |
| 1382.643 | COL3A1 | 639 | 653 | GLQGLPGTGGPPGEN | GLQGLP[113]GTGGPP[113]GEN | GLQGLP(0.9843)GTGGP(0.0315)P(0.9842)GEN | 24.02 | 20.15 | 1.000 |
| 1384.635 | COL3A1 | 1002 | 1016 | GLAGTAGEPGRDGNP | GLAGTAGEP[113]GRDGNP | GLAGTAGEP(0.9887)GRDGNP(0.0113) | 24.31 | 15.97 | 0.999 |
| 1385.645 | COL3A1 | 283 | 296 | LKGENGLPGENGAP | LKGEN[115]GLP[113]GENGAP[113] | LKGENGLP(1.0000)GENGAP(1.0000) | 19.07 | 12.00 | 0.980 |
| 1386.627 | COL3A1 | 283 | 296 | LKGENGLPGENGAP | LKGEN[115]GLP[113]GEN[115]GAP[113] | LKGENGLP(1.0000)GENGAP(1.0000) | 20.42 | 12.64 | 0.991 |
| 1386.681 | COL3A1 | 480 | 494 | GAAGERGAPGFRGPA | GAAGERGAP[113]GFRGPA | GAAGERGAP(0.9853)GFRGP(0.0147)A | 18.22 | 15.72 | 0.990 |
| 1400.630 | COL3A1 | 1002 | 1016 | GLAGTAGEPGRDGNP | GLAGTAGEP[113]GRDGNP[113] | GLAGTAGEP(1.0000)GRDGNP(1.0000) | 37.77 | 13.29 | 1.000 |
| 1401.611 | COL3A1 | 465 | 479 | GKDGSPGEPGANGLP | GKDGSP[113]GEP[113]GAN[115]GLP[113] | GKDGSP(1.0000)GEP(1.0000)GANGLP(1.0000) | 19.19 | 12.06 | 0.991 |
| 1403.601 | COL3A1 | 384 | 398 | GPPGINGSPGGKGEM | GPP[113]GIN[115]GSP[113]GGKGEM[147] | GP(0.0325)P(0.9892)GINGSP(0.9892)GGKGEM(0.9892) | 20.98 | 18.70 | 0.964 |
| 1403.658 | COL3A1 | 724 | 737 | LQGMPGERGGLGSP | LQGM[147]P[113]GERGGLGSP[113] | LQGM(1.0000)P(1.0000)GERGGLGSP(1.0000) | 19.86 | 13.51 | 0.977 |
| 1409.740 | COL3A1 | 246 | 260 | GPPGIKGPAGIPGFP | GPP[113]GIKGPAGIP[113]GFP[113] | GP(0.3884)P(0.8539)GIKGP(0.0193)AGIP(0.8692)GFP(0.8692) | 21.82 | 20.65 | 0.983 |
| 1410.689 | COL3A1 | 981 | 995 | GKPGANGLSGERGPP | GKPGAN[115]GLSGERGP[113]P | GKP(0.0041)GANGLSGERGP(0.4979)P(0.4979) | 19.93 | 19.93 | 0.969 |
| 1413.663 | COL3A1 | 921 | 935 | GPKGDAGQPGEKGSP | GPKGDAGQP[113]GEKGSP[113] | GP(0.0194)KGDAGQP(0.9903)GEKGSP(0.9903) | 19.80 | 11.68 | 0.994 |
| 1425.735 | COL3A1 | 246 | 260 | GPPGIKGPAGIPGFP | GP[113]P[113]GIKGPAGIP[113]GFP[113] | GP(0.8174)P(0.8244)GIKGP(0.6961)AGIP(0.8310)GFP(0.8310) | 24.61 | 22.91 | 1.000 |
| 1426.679 | COL3A1 | 282 | 296 | GLKGENGLPGENGAP | GLKGENGLPGEN[115]GAP[113] | GLKGENGLP(0.0197)GENGAP(0.9803) | 19.81 | 13.08 | 0.989 |
| 1426.680 | COL3A1 | 981 | 995 | GKPGANGLSGERGPP | GKP[113]GAN[115]GLSGERGP[113]P | GKP(0.7186)GANGLSGERGP(0.6407)P(0.6407) | 21.45 | 21.45 | 1.000 |
| 1427.664 | COL3A1 | 282 | 296 | GLKGENGLPGENGAP | GLKGEN[115]GLPGEN[115]GAP[113] | GLKGENGLP(0.0195)GENGAP(0.9805) | 27.86 | 15.15 | 1.000 |
| 1441.660 | COL3A1 | 1002 | 1017 | GLAGTAGEPGRDGNPG | GLAGTAGEP[113]GRDGNPG | GLAGTAGEP(0.9890)GRDGNP(0.0110)G | 24.53 | 20.40 | 1.000 |
| 1441.690 | COL3A1 | 282 | 296 | GLKGENGLPGENGAP | GLKGENGLP[113]GENGAP[113] | GLKGENGLP(1.0000)GENGAP(1.0000) | 21.29 | 13.36 | 0.999 |
| 1442.670 | COL3A1 | 282 | 296 | GLKGENGLPGENGAP | GLKGENGLP[113]GEN[115]GAP[113] | GLKGENGLP(1.0000)GENGAP(1.0000) | 18.61 | 10.63 | 0.993 |
| 1443.654 | COL3A1 | 282 | 296 | GLKGENGLPGENGAP | GLKGEN[115]GLP[113]GEN[115]GAP[113] | GLKGENGLP(1.0000)GENGAP(1.0000) | 18.07 | 10.73 | 0.999 |
| 1444.678 | COL3A1 | 723 | 737 | GLQGMPGERGGLGSP | GLQGM[147]PGERGGLGSP[113] | GLQGM(0.9790)P(0.0422)GERGGLGSP(0.9788) | 26.72 | 21.31 | 0.966 |
| 1446.601 | COL3A1 | 195 | 209 | GSPGYQGPPGEPGQA | GSP[113]GYQGPP[113]GEP[113]GQA | GSP(0.9461)GYQGP(0.1617)P(0.9458)GEP(0.9465)GQA | 26.46 | 24.77 | 0.987 |
| 1455.727 | COL3A1 | 972 | 987 | GPQGVKGESGKPGANG | GPQGVKGESGKP[113]GANG | GP(0.0468)QGVKGESGKP(0.9532)GANG | 21.46 | 12.40 | 1.000 |
| 1457.650 | COL3A1 | 1002 | 1017 | GLAGTAGEPGRDGNPG | GLAGTAGEP[113]GRDGNP[113]G | GLAGTAGEP(1.0000)GRDGNP(1.0000)G | 26.62 | 10.73 | 1.000 |
| 1457.696 | COL3A1 | 1089 | 1103 | GPRGDKGETGERGAA | GPRGDKGETGERGAA | | 18.79 | 10.64 | 0.997 |
| 1460.666 | COL3A1 | 723 | 737 | GLQGMPGERGGLGSP | GLQGM[147]P[113]GERGGLGSP[113] | GLQGM(1.0000)P(1.0000)GERGGLGSP(1.0000) | 30.99 | 14.09 | 1.000 |
| 1461.659 | COL3A1 | 723 | 737 | GLQGMPGERGGLGSP | GLQ[129]GM[147]P[113]GERGGLGSP[113] | GLQGM(1.0000)P(1.0000)GERGGLGSP(1.0000) | 32.37 | 15.38 | 1.000 |
| 1462.671 | COL3A1 | 366 | 380 | GQRGEPGPQGHAGAQ | GQRGEP[113]GPQGHAGAQ | GQRGEP(0.9883)GP(0.0117)QGHAGAQ | 19.79 | 15.99 | 0.977 |
| 1473.706 | COL3A1 | 1089 | 1103 | GPRGDKGETGERGAA | GP[113]RGDKGETGERGAA | GP(1.0000)RGDKGETGERGAA | 18.89 | 12.56 | 1.000 |
| 1488.656 | COL3A1 | 525 | 539 | GRDGVPGGPGMRGMP | GRDGVP[113]GGP[113]GMRGMP[113] | GRDGVP(0.9867)GGP(0.9866)GM(0.0204)RGM(0.0197)P(0.9867) | 22.49 | 20.45 | 1.000 |
| 1504.651 | COL3A1 | 525 | 539 | GRDGVPGGPGMRGMP | GRDGVP[113]GGP[113]GM[147]RGMP[113] | GRDGVP(0.9708)GGP(0.9707)GM(0.9706)RGM(0.1172)P(0.9708) | 21.72 | 19.88 | 0.995 |
| 1508.660 | COL3A1 | 267 | 281 | GFDGRNGEKGETGAP | GFDGRN[115]GEKGETGAP[113] | GFDGRNGEKGETGAP(1.0000) | 21.56 | 12.95 | 1.000 |
| 1548.877 | COL3A1 | 942 | 959 | GAPGPLGIAGITGARGLA | GAPGPLGIAGITGARGLA | | 29.46 | 11.44 | 1.000 |
| 1564.869 | COL3A1 | 942 | 959 | GAPGPLGIAGITGARGLA | GAP[113]GPLGIAGITGARGLA | GAP(0.9886)GP(0.0114)LGIAGITGARGLA | 30.86 | 25.91 | 1.000 |
| 1578.724 | COL3A1 | 828 | 845 | GAPGEKGEGGPPGVAGPP | GAP[113]GEKGEGGPP[113]GVAGP[113]P | GAP(0.7869)GEKGEGGP(0.0637)P(0.7913)GVAGP(0.6791)P(0.6791) | 25.13 | 25.13 | 0.969 |
| 1588.770 | COL3A1 | 859 | 875 | VKGERGSPGGPGAAGFP | VKGERGSP[113]GGP[113]GAAGFP[113] | VKGERGSP(1.0000)GGP(1.0000)GAAGFP(1.0000) | 18.36 | 13.32 | 0.972 |
| 1590.894 | COL3A1 | 945 | 962 | GPLGIAGITGARGLAGPP | GPLGIAGITGARGLAGPP[113] | GP(0.0057)LGIAGITGARGLAGP(0.0183)P(0.9760) | 44.01 | 44.01 | 1.000 |
| 1593.782 | COL3A1 | 774 | 791 | GQPGDKGEGGAPGLPGIA | GQPGDKGEGGAPGLP[113]GIA | GQP(0.0078)GDKGEGGAP(0.0066)GLP(0.9856)GIA | 19.11 | 12.87 | 0.984 |
| 1595.729 | COL3A1 | 669 | 686 | GAPGGKGDAGAPGERGPP | GAP[113]GGKGDAGAP[113]GERGP[113]P | GAP(0.7800)GGKGDAGAP(0.7800)GERGP(0.7200)P(0.7200) | 25.40 | 25.40 | 1.000 |
| 1595.764 | COL3A1 | 1053 | 1070 | GPVGPAGKSGDRGESGPA | GPVGPAGKSGDRGESGPA | | 23.94 | 14.97 | 0.999 |
| 1602.687 | COL3A1 | 1003 | 1019 | LAGTAGEPGRDGNPGSD | LAGTAGEP[113]GRDGNP[113]GSD | LAGTAGEP(1.0000)GRDGNP(1.0000)GSD | 29.46 | 8.34 | 1.000 |
| 1609.774 | COL3A1 | 774 | 791 | GQPGDKGEGGAPGLPGIA | GQP[113]GDKGEGGAPGLP[113]GIA | GQP(0.9888)GDKGEGGAP(0.0222)GLP(0.9889)GIA | 21.48 | 15.01 | 0.992 |
| 1612.718 | COL3A1 | 384 | 401 | GPPGINGSPGGKGEMGPA | GPP[113]GIN[115]GSP[113]GGKGEMGPA | GP(0.0142)P(0.9754)GINGSP(0.9757)GGKGEM(0.0106)GP(0.0242)A | 18.98 | 18.98 | 0.941 |
| 1625.775 | COL3A1 | 774 | 791 | GQPGDKGEGGAPGLPGIA | GQP[113]GDKGEGGAP[113]GLP[113]GIA | GQP(1.0000)GDKGEGGAP(1.0000)GLP(1.0000)GIA | 33.58 | 14.72 | 1.000 |
| 1628.713 | COL3A1 | 384 | 401 | GPPGINGSPGGKGEMGPA | GPP[113]GIN[115]GSP[113]GGKGEM[147]GPA | GP(0.0173)P(0.9884)GINGSP(0.9879)GGKGEM(0.9884)GP(0.0180)A | 19.42 | 17.73 | 0.982 |
| 1637.822 | COL3A1 | 777 | 794 | GDKGEGGAPGLPGIAGPR | GDKGEGGAP[113]GLP[113]GIAGPR | GDKGEGGAP(0.9892)GLP(0.9892)GIAGP(0.0216)R | 30.84 | 20.90 | 1.000 |
| 1643.719 | COL3A1 | 1002 | 1019 | GLAGTAGEPGRDGNPGSD | GLAGTAGEP[113]GRDGNPGSD | GLAGTAGEP(0.9894)GRDGNP(0.0106)GSD | 30.07 | 15.95 | 1.000 |
| 1645.787 | COL3A1 | 858 | 875 | GVKGERGSPGGPGAAGFP | GVKGERGSP[113]GGP[113]GAAGFP[113] | GVKGERGSP(1.0000)GGP(1.0000)GAAGFP(1.0000) | 18.79 | 11.86 | 0.976 |
| 1651.779 | COL3A1 | 516 | 533 | GPRGAAGEPGRDGVPGGP | GPRGAAGEP[113]GRDGVP[113]GGP[113] | GP(0.0334)RGAAGEP(0.9889)GRDGVP(0.9889)GGP(0.9889) | 20.01 | 13.05 | 0.995 |
| 1659.705 | COL3A1 | 1002 | 1019 | GLAGTAGEPGRDGNPGSD | GLAGTAGEP[113]GRDGNP[113]GSD | GLAGTAGEP(1.0000)GRDGNP(1.0000)GSD | 34.45 | 8.58 | 1.000 |
| 1660.691 | COL3A1 | 1002 | 1019 | GLAGTAGEPGRDGNPGSD | GLAGTAGEP[113]GRDGN[115]P[113]GSD | GLAGTAGEP(1.0000)GRDGNP(1.0000)GSD | 21.32 | 7.89 | 1.000 |
| 1664.779 | COL3A1 | 636 | 653 | GPQGLQGLPGTGGPPGEN | GPQGLQGLP[113]GTGGPP[113]GEN | GP(0.0130)QGLQGLP(0.9828)GTGGP(0.0219)P(0.9824)GEN | 38.10 | 33.49 | 1.000 |
| 1665.774 | COL3A1 | 636 | 653 | GPQGLQGLPGTGGPPGEN | GPQ[129]GLQGLP[113]GTGGPP[113]GEN | GP(0.0178)QGLQGLP(0.9738)GTGGP(0.0356)P(0.9729)GEN | 24.24 | 24.24 | 1.000 |
| 1666.730 | COL3A1 | 642 | 659 | GLPGTGGPPGENGKPGEP | GLP[113]GTGGPP[113]GEN[115]GKPGEP[113] | GLP(0.9300)GTGGP(0.1621)P(0.9265)GENGKP(0.0519)GEP(0.9293) | 21.60 | 21.60 | 1.000 |
| 1667.750 | COL3A1 | 771 | 789 | GPAGQPGDKGEGGAPGLPG | GPAGQ[129]P[113]GDKGEGGAP[113]GLP[113]G | GP(0.0483)AGQP(0.9839)GDKGEGGAP(0.9839)GLP(0.9839)G | 18.09 | 12.89 | 0.992 |
| 1668.791 | COL3A1 | 822 | 839 | GGKGERGAPGEKGEGGPP | GGKGERGAP[113]GEKGEGGPP[113] | GGKGERGAP(0.9888)GEKGEGGP(0.0224)P(0.9888) | 18.90 | 18.90 | 0.990 |
| 1669.769 | COL3A1 | 921 | 938 | GPKGDAGQPGEKGSPGAQ | GPKGDAGQP[113]GEKGSP[113]GAQ | GP(0.0209)KGDAGQP(0.9895)GEKGSP(0.9895)GAQ | 27.77 | 14.37 | 1.000 |
| 1681.747 | COL3A1 | 642 | 659 | GLPGTGGPPGENGKPGEP | GLP[113]GTGGPP[113]GENGKP[113]GEP[113] | GLP(0.9883)GTGGP(0.0470)P(0.9883)GENGKP(0.9882)GEP(0.9882) | 23.42 | 19.27 | 0.959 |
| 1682.740 | COL3A1 | 642 | 659 | GLPGTGGPPGENGKPGEP | GLP[113]GTGGPP[113]GEN[115]GKP[113]GEP[113] | GLP(0.9890)GTGGP(0.0439)P(0.9890)GENGKP(0.9890)GEP(0.9890) | 25.18 | 20.88 | 1.000 |
| 1683.771 | COL3A1 | 999 | 1016 | GLPGLAGTAGEPGRDGNP | GLP[113]GLAGTAGEP[113]GRDGNP[113] | GLP(1.0000)GLAGTAGEP(1.0000)GRDGNP(1.0000) | 22.10 | 12.84 | 0.993 |
| 1684.814 | COL3A1 | 426 | 443 | GLRGGAGEPGKNGAKGEP | GLRGGAGEP[113]GKN[115]GAKGEP[113] | GLRGGAGEP(1.0000)GKNGAKGEP(1.0000) | 30.77 | 16.57 | 1.000 |
| 1685.753 | COL3A1 | 519 | 536 | GAAGEPGRDGVPGGPGMR | GAAGEP[113]GRDGVP[113]GGP[113]GMR | GAAGEP(0.7802)GRDGVP(0.7801)GGP(0.7199)GM(0.7199)R | 21.86 | 21.86 | 0.964 |
| 1687.707 | COL3A1 | 195 | 212 | GSPGYQGPPGEPGQAGPS | GSP[113]GYQGPP[113]GEP[113]GQAGPS | GSP(0.9891)GYQGP(0.0163)P(0.9893)GEP(0.9894)GQAGP(0.0159)S | 41.28 | 36.54 | 1.000 |
| 1688.686 | COL3A1 | 195 | 212 | GSPGYQGPPGEPGQAGPS | GSP[113]GYQGPP[113]GEP[113]GQ[129]AGPS | GSP(0.9875)GYQGP(0.0195)P(0.9877)GEP(0.9878)GQAGP(0.0174)S | 21.15 | 19.45 | 0.996 |
| 1692.790 | COL3A1 | 495 | 512 | GPNGIPGEKGPAGERGAP | GPNGIP[113]GEKGPAGERGAP[113] | GP(0.0081)NGIP(0.7895)GEKGP(0.4610)AGERGAP(0.7414) | 19.07 | 18.65 | 1.000 |

| | | | | | | | | | |
|---|---|---|---|---|---|---|---|---|---|
| 1693.800 | COL3A1 | 495 | 512 | GPNGIPGEKGPAGERGAP | GPN[115]GIP[113]GEKGPAGERGAP[113] | GP(0.0105)NGIP(0.9890)GEKGP(0.0116)AGERGAP(0.9889) | 30.02 | 23.65 | 1.000 |
| 1699.780 | COL3A1 | 978 | 995 | GESGKPGANGLSGERGPP | GESGKP[113]GAN[115]GLSGERGP[113]P | GESGKP(0.7196)GANGLSGERGP(0.6402)P(0.6402) | 18.12 | 18.12 | 0.996 |
| 1701.752 | COL3A1 | 519 | 536 | GAAGEPGRDGVPGGPGMR | GAAGEP[113]GRDGVP[113]GGP[113]GM[147]R | GAAGEP(1.0000)GRDGVP(1.0000)GGP(1.0000)GM(1.0000)R | 20.21 | 11.79 | 0.998 |
| 1703.703 | COL3A1 | 192 | 209 | GSPGSPGYQGPPGEPGQA | GSP[113]GSP[113]GYQGPP[113]GEP[113]GQA | GSP(0.9870)GSP(0.9871)GYQGP(0.0518)P(0.9871)GEP(0.9871)GQA | 28.61 | 26.81 | 1.000 |
| 1709.809 | COL3A1 | 495 | 512 | GPNGIPGEKGPAGERGAP | GPN[115]GIP[113]GEKGP[113]AGERGAP[113] | GP(0.4293)NGIP(0.8593)GEKGP(0.8554)AGERGAP(0.8560) | 22.08 | 13.44 | 0.996 |
| 1716.732 | COL3A1 | 1002 | 1020 | GLAGTAGEPGRDGNPGSDG | GLAGTAGEP[113]GRDGNP[113]GSDG | GLAGTAGEP(1.0000)GRDGNP(1.0000)GSDG | 18.53 | 9.53 | 1.000 |
| 1731.799 | COL3A1 | 720 | 737 | GTPGLQGMPGERGGLGSP | GTP[113]GLQGM[147]P[113]GERGGLGSP[113] | GTP(1.0000)GLQGM(1.0000)P(1.0000)GERGGLGSP(1.0000) | 19.54 | 12.47 | 0.993 |
| 1744.762 | COL3A1 | 282 | 299 | GLKGENGLPGENGAPGPM | GLKGEN[115]GLP[113]GEN[115]GAP[113]GP[113]M | GLKGENGLP(0.9627)GENGAP(0.9626)GP(0.9623)M(0.1124) | 23.52 | 23.52 | 0.998 |
| 1750.865 | COL3A1 | 776 | 794 | PGDKGEGGAPGLPGIAGPR | P[113]GDKGEGGAP[113]GLP[113]GIAGPR | P(0.9715)GDKGEGGAP(0.9716)GLP(0.9713)GIAGP(0.0855)R | 18.97 | 13.50 | 1.000 |
| 1834.899 | COL3A1 | 771 | 791 | GPAGQPGDKGEGGAPGLPGIA | GPAGQP[113]GDKGEGGAPGLP[113]GIA | GP(0.0166)AGQP(0.9814)GDKGEGGAP(0.0209)GLP(0.9812)GIA | 18.15 | 14.49 | 0.998 |
| 1835.885 | COL3A1 | 771 | 791 | GPAGQPGDKGEGGAPGLPGIA | GPAGQ[129]P[113]GDKGEGGAPGLP[113]GIA | GP(0.0115)AGQP(0.9878)GDKGEGGAP(0.0129)GLP(0.9878)GIA | 20.60 | 16.77 | 0.983 |
| 1850.881 | COL3A1 | 771 | 791 | GPAGQPGDKGEGGAPGLPGIA | GPAGQP[113]GDKGEGGAP[113]GLP[113]GIA | GP(0.1961)AGQP(0.9340)GDKGEGGAP(0.9349)GLP(0.9349)GIA | 19.79 | 18.63 | 0.986 |
| 1851.862 | COL3A1 | 771 | 791 | GPAGQPGDKGEGGAPGLPGIA | GPAGQ[129]P[113]GDKGEGGAP[113]GLP[113]GIA | GP(0.0453)AGQP(0.9849)GDKGEGGAP(0.9849)GLP(0.9849)GIA | 20.13 | 16.31 | 0.963 |
| 1853.864 | COL3A1 | 1003 | 1022 | LAGTAGEPGRDGNPGSDGLP | LAGTAGEP[113]GRDGNPGSDGLP | LAGTAGEP(0.9854)GRDGNP(0.0081)GSDGLP(0.0065) | 19.21 | 12.76 | 0.974 |
| 1869.854 | COL3A1 | 1003 | 1022 | LAGTAGEPGRDGNPGSDGLP | LAGTAGEP[113]GRDGNPGSDGLP[113] | LAGTAGEP(0.9879)GRDGNP(0.0242)GSDGLP(0.9879) | 20.23 | 16.16 | 0.995 |
| 1870.844 | COL3A1 | 1003 | 1022 | LAGTAGEPGRDGNPGSDGLP | LAGTAGEP[113]GRDGN[115]PGSDGLP[113] | LAGTAGEP(0.9723)GRDGNP(0.0556)GSDGLP(0.9720) | 20.33 | 16.23 | 1.000 |
| 1885.833 | COL3A1 | 1003 | 1022 | LAGTAGEPGRDGNPGSDGLP | LAGTAGEP[113]GRDGNP[113]GSDGLP[113] | LAGTAGEP(1.0000)GRDGNP(1.0000)GSDGLP(1.0000) | 20.55 | 11.52 | 0.960 |
| 1885.860 | COL3A1 | 387 | 407 | GINGSPGGKGEMGPAGIPGAP | GIN[115]GSP[113]GGKGEM[147]GPAGIP[113]GAP[113] | GINGSP(0.8811)GGKGEM(0.8783)GP(0.4779)AGIP(0.8814)GAP(0.8814) | 22.62 | 19.16 | 0.997 |
| 1886.848 | COL3A1 | 1003 | 1022 | LAGTAGEPGRDGNPGSDGLP | LAGTAGEP[113]GRDGN[115]P[113]GSDGLP[113] | LAGTAGEP(1.0000)GRDGNP(1.0000)GSDGLP(1.0000) | 19.12 | 12.89 | 0.986 |
| 1907.890 | COL3A1 | 640 | 659 | LQGLPGTGGPPGENGKPGEP | LQGLP[113]GTGGPP[113]GEN[115]GKPGEP[113] | LQGLP(0.9773)GTGGP(0.0344)P(0.9771)GENGKP(0.0341)GEP(0.9771) | 20.81 | 20.81 | 0.976 |
| 1907.906 | COL3A1 | 738 | 758 | GPKGDKGEPGGPGADGVPGKD | GPKGDKGEP[113]GGPGADGVPGKD | GP(0.0034)KGDKGEP(0.9887)GGP(0.0040)GADGVP(0.0038)GKD | 21.55 | 17.45 | 1.000 |
| 1910.882 | COL3A1 | 1002 | 1022 | GLAGTAGEPGRDGNPGSDGLP | GLAGTAGEP[113]GRDGNPGSDGLP | GLAGTAGEP(0.9859)GRDGNP(0.0082)GSDGLP(0.0059) | 20.92 | 12.82 | 0.976 |
| 1911.873 | COL3A1 | 384 | 404 | GPPGINGSPGGKGEMGPAGIP | GPP[113]GIN[115]GSP[113]GGKGEM[147]GPAGIP[113] | GP(0.0265)P(0.9876)GINGSP(0.9876)GGKGEM(0.9876)GP(0.0231)AGIP(0.9876) | 26.24 | 26.24 | 1.000 |
| 1913.873 | COL3A1 | 915 | 935 | GSPGVSGPKGDAGQPGEKGSP | GSP[113]GVSGPKGDAGQP[113]GEKGSP[113] | GSP(0.9666)GVSGP(0.0996)KGDAGQP(0.9669)GEKGSP(0.9669) | 24.35 | 18.59 | 1.000 |
| 1914.864 | COL3A1 | 915 | 935 | GSPGVSGPKGDAGQPGEKGSP | GSP[113]GVSGPKGDAGQ[129]P[113]GEKGSP[113] | GSP(0.9898)GVSGP(0.0306)KGDAGQP(0.9898)GEKGSP(0.9898) | 20.59 | 16.45 | 1.000 |
| 1914.892 | COL3A1 | 717 | 737 | GAAGTPGLQGMPGERGGLGSP | GAAGTPGLQGM[147]P[113]GERGGLGSP[113] | GAAGTP(0.0561)GLQGM(0.9812)P(0.9813)GERGGLGSP(0.9813) | 20.69 | 12.69 | 0.992 |
| 1919.918 | COL3A1 | 741 | 761 | GDKGEPGGPGADGVPGKDGPR | GDKGEPGGPGADGVPGKDGPR | | 18.76 | 10.64 | 1.000 |
| 1919.932 | COL3A1 | 774 | 794 | GQPGDKGEGGAPGLPGIAGPR | GQPGDKGEGGAP[113]GLP[113]GIAGPR | GQP(0.0115)GDKGEGGAP(0.9881)GLP(0.9880)GIAGP(0.0124)R | 21.83 | 14.80 | 1.000 |
| 1922.901 | COL3A1 | 640 | 659 | LQGLPGTGGPPGENGKPGEP | LQGLP[113]GTGGP[113]PGENGKP[113]GEP[113] | LQGLP(0.8194)GTGGP(0.7710)P(0.7710)GENGKP(0.8193)GEP(0.8194) | 22.83 | 22.83 | 1.000 |
| 1923.885 | COL3A1 | 640 | 659 | LQGLPGTGGPPGENGKPGEP | LQGLP[113]GTGGP[113]PGEN[115]GKP[113]GEP[113] | LQGLP(0.8192)GTGGP(0.7715)P(0.7715)GENGKP(0.8190)GEP(0.8188) | 21.68 | 21.68 | 1.000 |
| 1926.874 | COL3A1 | 1002 | 1022 | GLAGTAGEPGRDGNPGSDGLP | GLAGTAGEP[113]GRDGNPGSDGLP[113] | GLAGTAGEP(0.9857)GRDGNP(0.0286)GSDGLP(0.9858) | 18.03 | 14.48 | 0.985 |
| 1926.974 | COL3A1 | 976 | 995 | VKGESGKPGANGLSGERGPP | VKGESGKP[113]GAN[115]GLSGERGP[113]P | VKGESGKP(0.7208)GANGLSGERGP(0.6396)P(0.6396) | 20.90 | 20.90 | 0.999 |
| 1927.858 | COL3A1 | 1002 | 1022 | GLAGTAGEPGRDGNPGSDGLP | GLAGTAGEP[113]GRDGN[115]PGSDGLP[113] | GLAGTAGEP(0.9902)GRDGNP(0.0197)GSDGLP(0.9901) | 34.04 | 23.33 | 1.000 |
| 1927.917 | COL3A1 | 855 | 875 | GPQGVKGERGSPGGPGAAGFP | GPQGVKGERGSP[113]GGP[113]GAAGFP[113] | GP(0.5330)QGVKGERGSP(0.8174)GGP(0.8240)GAAGFP(0.8256) | 21.03 | 13.74 | 0.996 |
| 1928.908 | COL3A1 | 855 | 875 | GPQGVKGERGSPGGPGAAGFP | GPQ[129]GVKGERGSP[113]GGP[113]GAAGFP[113] | GP(0.2181)QGVKGERGSP(0.9269)GGP(0.9270)GAAGFP(0.9280) | 18.11 | 12.89 | 0.976 |
| 1930.885 | COL3A1 | 717 | 737 | GAAGTPGLQGMPGERGGLGSP | GAAGTP[113]GLQGM[147]P[113]GERGGLGSP[113] | GAAGTP(1.0000)GLQGM(1.0000)P(1.0000)GERGGLGSP(1.0000) | 18.95 | 10.63 | 0.999 |
| 1935.905 | COL3A1 | 741 | 761 | GDKGEPGGPGADGVPGKDGPR | GDKGEP[113]GGPGADGVPGKDGPR | GDKGEP(0.9868)GGP(0.0042)GADGVP(0.0046)GKDGP(0.0045)R | 20.19 | 15.93 | 1.000 |
| 1935.928 | COL3A1 | 774 | 794 | GQPGDKGEGGAPGLPGIAGPR | GQP[113]GDKGEGGAP[113]GLP[113]GIAGPR | GQP(0.9711)GDKGEGGAP(0.9711)GLP(0.9709)GIAGP(0.0869)R | 22.95 | 20.98 | 1.000 |
| 1942.832 | COL3A1 | 1002 | 1022 | GLAGTAGEPGRDGNPGSDGLP | GLAGTAGEP[113]GRDGNP[113]GSDGLP[113] | GLAGTAGEP(1.0000)GRDGNP(1.0000)GSDGLP(1.0000) | 62.38 | 12.85 | 1.000 |
| 1942.859 | COL3A1 | 459 | 479 | GAKGEDGKDGSPGEPGANGLP | GAKGEDGKDGSPGEP[113]GAN[115]GLP[113] | GAKGEDGKDGSP(0.1961)GEP(0.9007)GANGLP(0.9032) | 19.09 | 15.40 | 0.997 |
| 1943.851 | COL3A1 | 1002 | 1022 | GLAGTAGEPGRDGNPGSDGLP | GLAGTAGEP[113]GRDGN[115]P[113]GSDGLP[113] | GLAGTAGEP(1.0000)GRDGNP(1.0000)GSDGLP(1.0000) | 21.92 | 14.38 | 0.989 |
| 1943.919 | COL3A1 | 855 | 875 | GPQGVKGERGSPGGPGAAGFP | GP[113]GQVKGERGSP[113]GGP[113]GAAGFP[113] | GP(1.0000)QGVKGERGSP(1.0000)GGP(1.0000)GAAGFP(1.0000) | 21.46 | 12.57 | 0.996 |
| 1944.797 | COL3A1 | 192 | 212 | GSPGSPGYQGPPGEPGQAGPS | GSP[113]GSP[113]GYQGPP[113]GEP[113]GQAGPS | GSP(0.9831)GSP(0.9831)GYQGP(0.0444)P(0.9829)GEP(0.9831)GQAGP(0.0233)S | 31.94 | 29.84 | 1.000 |
| 1945.791 | COL3A1 | 192 | 212 | GSPGSPGYQGPPGEPGQAGPS | GSP[113]GSP[113]GYQGPP[113]GEP[113]GQ[129]AGPS | GSP(0.9502)GSP(0.9501)GYQGP(0.1784)P(0.9483)GEP(0.9504)GQAGP(0.0226)S | 35.40 | 33.03 | 0.990 |
| 1950.877 | COL3A1 | 648 | 668 | GPPGENGKPGEPGPKGDAGAP | GPP[113]GEN[115]GKP[113]GEPGP[113]KGDAGAP[113] | GP(0.0201)P(0.9798)GENGKP(0.9768)GEP(0.0633)GP(0.9797)KGDAGAP(0.9802) | 19.52 | 17.88 | 0.988 |
| 1951.912 | COL3A1 | 741 | 761 | GDKGEPGGPGADGVPGKDGPR | GDKGEP[113]GGP[113]GADGVPGKDGPR | GDKGEP(0.9883)GGP(0.9883)GADGVP(0.0108)GKDGP(0.0126)R | 25.34 | 15.14 | 1.000 |
| 1958.849 | COL3A1 | 459 | 479 | GAKGEDGKDGSPGEPGANGLP | GAKGEDGKDGSP[113]GEP[113]GAN[115]GLP[113] | GAKGEDGKDGSP(1.0000)GEP(1.0000)GANGLP(1.0000) | 19.38 | 10.27 | 0.995 |
| 1964.914 | COL3A1 | 639 | 659 | GLQGLPGTGGPPGENGKPGEP | GLQGLP[113]GTGGPP[113]GEN[115]GKP[113]GEP | GLQGLP(0.9528)GTGGP(0.0439)P(0.9525)GENGKP(0.9513)GEP(0.0995) | 23.87 | 23.87 | 0.998 |
| 1970.869 | COL3A1 | 1002 | 1023 | GLAGTAGEPGRDGNPGSDGLPG | GLAGTAGEP[113]GRDGNPGSDGLPG | GLAGTAGEP(0.9884)GRDGNP(0.0058)GSDGLP(0.0058)G | 20.05 | 19.87 | 1.000 |
| 1979.920 | COL3A1 | 639 | 659 | GLQGLPGTGGPPGENGKPGEP | GLQGLP[113]GTGGPP[113]GENGKP[113]GEP[113] | GLQGLP(0.9814)GTGGP(0.0744)P(0.9813)GENGKP(0.9814)GEP(0.9814) | 23.28 | 21.18 | 0.992 |
| 1980.890 | COL3A1 | 639 | 659 | GLQGLPGTGGPPGENGKPGEP | GLQGLP[113]GTGGPP[113]GEN[115]GKP[113]GEP[113] | GLQGLP(0.9856)GTGGP(0.0567)P(0.9859)GENGKP(0.9859)GEP(0.9859) | 29.98 | 25.99 | 1.000 |
| 1983.959 | COL3A1 | 975 | 995 | GVKGESGKPGANGLSGERGPP | GVKGESGKP[113]GAN[115]GLSGERGPP[113] | GVKGESGKP(0.9756)GANGLSGERGP(0.0490)P(0.9754) | 22.40 | 22.40 | 1.000 |
| 1986.861 | COL3A1 | 519 | 539 | GAAGEPGRDGVPGGPGMRGMP | GAAGEP[113]GRDGVP[113]GGP[113]GMRGMP[113] | GAAGEP(0.8197)GRDGVP(0.8182)GGP(0.8052)GM(0.4861)RGM(0.2497)P(0.8211) | 21.31 | 21.31 | 0.999 |
| 1998.975 | COL3A1 | 972 | 992 | GPQGVKGESGKPGANGLSGER | GPQGVKGESGKP[113]GAN[115]GLSGER | GP(0.0122)QGVKGESGKP(0.9878)GANGLSGER | 18.99 | 9.12 | 1.000 |
| 2002.844 | COL3A1 | 519 | 539 | GAAGEPGRDGVPGGPGMRGMP | GAAGEP[113]GRDGVP[113]GGP[113]GMRGM[147]P[113] | GAAGEP(0.8468)GRDGVP(0.8468)GGP(0.8065)GM(0.8065)RGM(0.8468)P(0.8466) | 21.73 | 20.08 | 1.000 |
| 2079.034 | COL3A1 | 225 | 245 | GPAGKDGESGRPGRPGERGLP | GPAGKDGESGRPGRP[113]GERGLP[113] | GP(0.0229)AGKDGESGRP(0.0127)GRP(0.9822)GERGLP(0.9822) | 18.92 | 16.83 | 1.000 |
| 2095.030 | COL3A1 | 225 | 245 | GPAGKDGESGRPGRPGERGLP | GPAGKDGESGRP[113]GRP[113]GERGLP[113] | GP(0.0516)AGKDGESGRP(0.9828)GRP(0.9828)GERGLP(0.9828) | 19.00 | 9.78 | 1.000 |
| 2113.056 | COL3A1 | 1092 | 1112 | GDKGETGERGAAGIKGHRGFP | GDKGETGERGAAGIKGHRGFP[113] | GDKGETGERGAAGIKGHRGFP(1.0000) | 18.50 | 11.25 | 1.000 |
| 2129.052 | COL3A1 | 1048 | 1070 | HPGPPGPVGPAGKSGDRGESGPA | HP[113]GP[113]P[113]GPVGPAGKSGDRGESGPA | HP(0.9840)GP(0.9840)P(0.9840)GP(0.0119)VGP(0.0159)AGKSGDRGESGP(0.0202)A | 23.00 | 18.25 | 0.981 |
| 2155.999 | COL3A1 | 1002 | 1024 | GLAGTAGEPGRDGNPGSDGLPGR | GLAGTAGEP[113]GRDGNP[113]GSDGLP[113]GR | GLAGTAGEP(1.0000)GRDGNP(1.0000)GSDGLP(1.0000)GR | 19.50 | 15.07 | 1.000 |
| 2161.041 | COL3A1 | 771 | 794 | GPAGQPGDKGEGGAPGLPGIAGPR | GPAGQP[113]GDKGEGGAP[113]GLP[113]GIAGPR | GP(0.0187)AGQP(0.9594)GDKGEGGAP(0.9594)GLP(0.9577)GIAGP(0.1048)R | 28.60 | 26.38 | 1.000 |
| 2165.949 | COL3A1 | 645 | 668 | GTGGPPGENGKPGEPGPKGDAGAP | GTGGPP[113]GEN[115]GKP[113]GEP[113]GPKGDAGAP[113] | GTGGP(0.0558)P(0.9656)GENGKP(0.9635)GEP(0.9653)GP(0.0839)KGDAGAP(0.9659) | 21.57 | 19.73 | 1.000 |
| 2169.975 | COL3A1 | 915 | 938 | GSPGVSGPKGDAGQPGEKGSPGAQ | GSP[113]GVSGPKGDAGQP[113]GEKGSP[113]GAQ | GSP(0.9894)GVSGP(0.0318)KGDAGQP(0.9894)GEKGSP(0.9894)GAQ | 24.81 | 20.43 | 1.000 |
| 2202.077 | COL3A1 | 738 | 761 | GPKGDKGEPGGPGADGVPGKDGPR | GPKGDKGEPGGPGADGVPGKDGPR | | 27.03 | 13.92 | 1.000 |
| 2210.028 | COL3A1 | 999 | 1022 | GLPGLAGTAGEPGRDGNPGSDGLP | GLP[113]GLAGTAGEP[113]GRDGNPGSDGLP[113] | GLP(0.9893)GLAGTAGEP(0.9893)GRDGNP(0.0322)GSDGLP(0.9893) | 21.32 | 21.32 | 1.000 |
| 2218.058 | COL3A1 | 738 | 761 | GPKGDKGEPGGPGADGVPGKDGPR | GPKGDKGEP[113]GGPGADGVPGKDGPR | GP(0.0031)KGDKGEP(0.9871)GGP(0.0039)GADGVP(0.0030)GKDGP(0.0030)R | 25.52 | 21.05 | 1.000 |
| 2226.024 | COL3A1 | 999 | 1022 | GLPGLAGTAGEPGRDGNPGSDGLP | GLP[113]GLAGTAGEP[113]GRDGNP[113]GSDGLP[113] | GLP(1.0000)GLAGTAGEP(1.0000)GRDGNP(1.0000)GSDGLP(1.0000) | 21.19 | 8.83 | 0.998 |
| 2228.001 | COL3A1 | 456 | 479 | GVPGAKGEDGKDGSPGEPGANGLP | GVP[113]GAKGEDGKDGSP[113]GEP[113]GAN[115]GLP[113] | GVP(1.0000)GAKGEDGKDGSP(1.0000)GEP(1.0000)GANGLP(1.0000) | 20.66 | 12.24 | 0.995 |
| 2234.062 | COL3A1 | 738 | 761 | GPKGDKGEPGGPGADGVPGKDGPR | GP[113]KGDKGEP[113]GGPGADGVPGKDGPR | GP(0.9863)KGDKGEP(0.9890)GGP(0.0080)GADGVP(0.0085)GKDGP(0.0082)R | 29.67 | 27.94 | 1.000 |
| 2250.032 | COL3A1 | 738 | 761 | GPKGDKGEPGGPGADGVPGKDGPR | GP[113]KGDKGEP[113]GGP[113]GADGVPGKDGPR | GP(0.9875)KGDKGEP(0.9878)GGP(0.9879)GADGVP(0.0187)GKDGP(0.0181)R | 24.62 | 16.27 | 0.985 |
| 2250.094 | COL3A1 | 972 | 995 | GPQGVKGESGKPGANGLSGERGPP | GPQGVKGESGKPGAN[115]GLSGERGP[113]P | GP(0.0036)QGVKGESGKP(0.0038)GANGLSGERGP(0.4963)P(0.4963) | 19.78 | 19.78 | 1.000 |
| 2266.091 | COL3A1 | 972 | 995 | GPQGVKGESGKPGANGLSGERGPP | GPQGVKGESGKP[113]GAN[115]GLSGERGP[113]P | GP(0.0406)QGVKGESGKP(0.7518)GANGLSGERGP(0.6038)P(0.6038) | 40.20 | 40.20 | 1.000 |
| 2267.087 | COL3A1 | 972 | 995 | GPQGVKGESGKPGANGLSGERGPP | GPQ[129]GVKGESGKP[113]GAN[115]GLSGERGP[113]P | GP(0.0639)QGVKGESGKP(0.7431)GANGLSGERGP(0.5965)P(0.5965) | 22.29 | 22.29 | 1.000 |
| 2271.031 | COL3A1 | 1002 | 1025 | GLAGTAGEPGRDGNPGSDGLPGRD | GLAGTAGEP[113]GRDGNP[113]GSDGLP[113]GRD | GLAGTAGEP(1.0000)GRDGNP(1.0000)GSDGLP(1.0000)GRD | 27.32 | 13.72 | 1.000 |
| 2281.101 | COL3A1 | 972 | 995 | GPQGVKGESGKPGANGLSGERGPP | GP[113]QGVKGESGKP[113]GANGLSGERGP[113]P | GP(0.7795)QGVKGESGKP(0.7797)GANGLSGERGP(0.7204)P(0.7204) | 20.09 | 20.09 | 1.000 |
| 2282.086 | COL3A1 | 972 | 995 | GPQGVKGESGKPGANGLSGERGPP | GP[113]QGVKGESGKP[113]GAN[115]GLSGERGP[113]P | GP(0.7796)QGVKGESGKP(0.7801)GANGLSGERGP(0.7201)P(0.7201) | 25.25 | 25.25 | 1.000 |
| 2297.044 | COL3A1 | 516 | 539 | GPRGAAGEPGRDGVPGGPGMRGMP | GPRGAAGEP[113]GRDGVP[113]GGP[113]GMRGMP[113] | GP(0.0161)RGAAGEP(0.9764)GRDGVP(0.9765)GGP(0.9763)GM(0.0579)RGM(0.0213)P(0.9754) | 23.45 | 21.31 | 0.981 |
| 2313.032 | COL3A1 | 516 | 539 | GPRGAAGEPGRDGVPGGPGMRGMP | GPRGAAGEP[113]GRDGVP[113]GGP[113]GMRGM[147]P[113] | GP(0.0886)RGAAGEP(0.9041)GRDGVP(0.9041)GGP(0.9045)GM(0.3952)RGM(0.8994)P(0.9042) | 27.82 | 25.81 | 1.000 |
| 2394.111 | COL3A1 | 417 | 443 | GPAGANGAPGLRGGAGEPGKNGAKGE | GPAGAN[115]GAP[113]GLRGGAGEP[113]GKN[115]GAKGEP[113] | GP(0.0506)AGANGAP(0.9832)GLRGGAGEP(0.9831)GKNGAKGEP(0.9831) | 24.06 | 18.39 | 1.000 |
| 2394.175 | COL3A1 | 417 | 443 | GPAGANGAPGLRGGAGEPGKNGAKGE | GPAGANGAP[113]GLRGGAGEP[113]GKN[115]GAKGEP[113] | GP(0.0915)AGANGAP(0.9696)GLRGGAGEP(0.9695)GKNGAKGEP(0.9694) | 19.67 | 19.06 | 1.000 |
| 2433.117 | COL3A1 | 642 | 668 | GLPGTGGPPGENGKPGEPGPKGDAGA | GLP[113]GTGGPP[113]GEN[115]GKPGEP[113]GPKGDAGAP[113] | GLP(0.8798)GTGGP(0.0206)P(0.8797)GENGKP(0.4654)GEP(0.8534)GP(0.0231)KGDAGAP(0.8780) | 27.11 | 27.11 | 1.000 |
| 2437.101 | COL3A1 | 915 | 941 | GSPGVSGPKGDAGQPGEKGSPGAQGP | GSP[113]GVSGPKGDAGQP[113]GEKGSP[113]GAQGPP[113] | GSP(0.7390)GVSGP(0.7390)KGDAGQP(0.8329)GEKGSP(0.8330)GAQGP(0.0233)P(0.8329) | 29.37 | 29.37 | 1.000 |
| 2438.090 | COL3A1 | 915 | 941 | GSPGVSGPKGDAGQPGEKGSPGAQGP | GSP[113]GVSGPKGDAGQP[113]GEKGSP[113]GAQ[129]GPP[113] | GSP(0.7364)GVSGP(0.7364)KGDAGQP(0.8279)GEKGSP(0.8279)GAQGP(0.0439)P(0.8276) | 20.74 | 20.74 | 0.961 |

Supplemental Table 1. Col3a1 peptide sequences from normal breast by LC-MS/MS.

| | | | | | | | | | |
|---|---|---|---|---|---|---|---|---|---|
| 2439.111 | COL3A1 | 1003 | 1028 | LAGTAGEPGRDGNPGSDGLPGRDGSP | LAGTAGEP[113]GRDGNPGSDGLPGRDGSP[113] | LAGTAGEP(0.7463)GRDGNP(0.2564)GSDGLP(0.2503)GRDGSP(0.7470) | 19.15 | 18.70 | 0.969 |
| 2455.091 | COL3A1 | 1003 | 1028 | LAGTAGEPGRDGNPGSDGLPGRDGSP | LAGTAGEP[113]GRDGNPGSDGLP[113]GRDGSP[113] | LAGTAGEP(0.9964)GRDGNP(0.0528)GSDGLP(0.9746)GRDGSP(0.9762) | 24.79 | 20.63 | 1.000 |
| 2471.091 | COL3A1 | 1003 | 1028 | LAGTAGEPGRDGNPGSDGLPGRDGSP | LAGTAGEP[113]GRDGNP[113]GSDGLP[113]GRDGSP[113] | LAGTAGEP(1.0000)GRDGNP(1.0000)GSDGLP(1.0000)GRDGSP(1.0000) | 25.91 | 8.00 | 1.000 |
| 2472.085 | COL3A1 | 1003 | 1028 | LAGTAGEPGRDGNPGSDGLPGRDGSP | LAGTAGEP[113]GRDGN[115]P[113]GSDGLP[113]GRDGSP[113] | LAGTAGEP(1.0000)GRDGNP(1.0000)GSDGLP(1.0000)GRDGSP(1.0000) | 23.49 | 12.04 | 1.000 |
| 2496.122 | COL3A1 | 1002 | 1028 | GLAGTAGEPGRDGNPGSDGLPGRDGSF | GLAGTAGEP[113]GRDGNPGSDGLP[113]GRDGSP | GLAGTAGEP(0.8450)GRDGNP(0.0243)GSDGLP(0.8202)GRDGSP(0.3105) | 26.97 | 24.76 | 1.000 |
| 2499.142 | COL3A1 | 459 | 485 | GAKGEDGKDGSPGEPGANGLPGAAGEF | GAKGEDGKDGSP[113]GEP[113]GANGLP[113]GAAGER | GAKGEDGKDGSP(1.0000)GEP(1.0000)GANGLP(1.0000)GAAGER | 20.25 | 10.76 | 1.000 |
| 2500.089 | COL3A1 | 459 | 485 | GAKGEDGKDGSPGEPGANGLPGAAGEF | GAKGEDGKDGSP[113]GEP[113]GAN[115]GLP[113]GAAGER | GAKGEDGKDGSP(1.0000)GEP(1.0000)GANGLP(1.0000)GAAGER | 24.43 | 17.86 | 1.000 |
| 2512.080 | COL3A1 | 1002 | 1028 | GLAGTAGEPGRDGNPGSDGLPGRDGSF | GLAGTAGEP[113]GRDGNP[113]GSDGLPGRDGSP[113] | GLAGTAGEP(0.8089)GRDGNP(0.8087)GSDGLP(0.5852)GRDGSP(0.7972) | 31.48 | 31.07 | 1.000 |
| 2513.116 | COL3A1 | 1002 | 1028 | GLAGTAGEPGRDGNPGSDGLPGRDGSF | GLAGTAGEP[113]GRDGN[115]PGSDGLP[113]GRDGSP[113] | GLAGTAGEP(0.9865)GRDGNP(0.0406)GSDGLP(0.9865)GRDGSP(0.9865) | 20.24 | 16.35 | 1.000 |
| 2528.062 | COL3A1 | 1002 | 1028 | GLAGTAGEPGRDGNPGSDGLPGRDGSF | GLAGTAGEP[113]GRDGNP[113]GSDGLP[113]GRDGSP[113] | GLAGTAGEP(1.0000)GRDGNP(1.0000)GSDGLP(1.0000)GRDGSP(1.0000) | 84.76 | 23.53 | 1.000 |
| 2529.109 | COL3A1 | 1002 | 1028 | GLAGTAGEPGRDGNPGSDGLPGRDGSF | GLAGTAGEP[113]GRDGN[115]P[113]GSDGLP[113]GRDGSP[113] | GLAGTAGEP(1.0000)GRDGNP(1.0000)GSDGLP(1.0000)GRDGSP(1.0000) | 26.73 | 21.93 | 1.000 |
| 2674.254 | COL3A1 | 640 | 668 | LQGLPGTGGPPGENGKPGEPGPKGDAC | LQGLP[113]GTGGP[113]P[113]GEN[115]GKPGEPGPKGDAGAP[1 | LQGLP(0.6943)GTGGP(0.5852)P(0.5852)GENGKP(0.3058)GEP(0.5852)GP(0.5852)KGDAGAP(0.6593) | 21.23 | 21.23 | 1.000 |
| 2725.222 | COL3A1 | 459 | 488 | GAKGEDGKDGSPGEPGANGLPGAAGEF | GAKGEDGKDGSPGEP[113]GAN[115]GLP[113]GAAGERGAP[113] | GAKGEDGKDGSP(0.0425)GEP(0.9858)GANGLP(0.9859)GAAGERGAP(0.9858) | 26.19 | 20.88 | 1.000 |
| 2730.294 | COL3A1 | 639 | 668 | GLQGLPGTGGPPGENGKPGEPGPKGDA | GLQGLP[113]GTGGPPGENGKP[113]GEP[113]GP[113]KGDAGAP | GLQGLP(0.8166)GTGGP(0.1275)P(0.1555)GENGKP(0.8129)GEP(0.8129)GP(0.7797)KGDAGAP(0.4948) | 18.14 | 18.14 | 0.997 |
| 2731.269 | COL3A1 | 639 | 668 | GLQGLPGTGGPPGENGKPGEPGPKGDA | GLQGLP[113]GTGGP[113]PGEN[115]GKPGEP[113]GPKGDAGAP[ | GLQGLP(0.7511)GTGGP(0.6307)P(0.6307)GENGKP(0.5249)GEP(0.6946)GP(0.0182)KGDAGAP(0.7497) | 25.90 | 25.90 | 1.000 |
| 2740.201 | COL3A1 | 459 | 488 | GAKGEDGKDGSPGEPGANGLPGAAGEF | GAKGEDGKDGSP[113]GEP[113]GANGLP[113]GAAGERGAP[113] | GAKGEDGKDGSP(1.0000)GEP(1.0000)GANGLP(1.0000)GAAGERGAP(1.0000) | 30.69 | 11.71 | 1.000 |
| 2741.202 | COL3A1 | 459 | 488 | GAKGEDGKDGSPGEPGANGLPGAAGEF | GAKGEDGKDGSP[113]GEP[113]GAN[115]GLP[113]GAAGERGAP[ | GAKGEDGKDGSP(1.0000)GEP(1.0000)GANGLP(1.0000)GAAGERGAP(1.0000) | 22.84 | 20.14 | 1.000 |

**Supplemental Table 2. MALDI-IHC Antibody Probes.** All antibodies used in this workflow are monoclonal antibodies. The table below includes the AmberGen catalog number, the target of the antibody, the host of the antibody, the m/z of the mass tag, the and the concentration used in the experiment.

| Catalog # | Target | Host | (M+H) | Concentration (µg/mL) |
|---|---|---|---|---|
| AP1001172 | Actin-aSM | Rabbit | 1251.68 | 2.5 |
| AP1001454 | Beta-catenin | Rabbit | 863.52 | 2.5 |
| AP1001171 | CD44 | Rabbit | 1102.59 | 2.5 |
| AP1001124 | CD68 | Rabbit | 1216.75 | 2.5 |
| AP100153 | COL1A1 | Rabbit | 1234.87 | 2.5 |
| AP1001183 | ECAD (CDH1) | Mouse | 930.56 | 2.5 |
| AP1001171 | Fibronectin (FN1) | Rabbit | 1068.60 | 3.125 |
| AP1001182 | Histone H3 | Rabbit | 1782.94 | 1.25 |
| AP1001184 | Ki67 | Mouse | 1320.76 | 1.75 |
| AP1001122 | Vimentin | Rabbit | 1230.84 | 1.25 |

**Supplemental Table 3. Summary Table of Calculated p-values for comparisons of SHG measurements by clinical characteristics.** Measurements were taken from either ductal regions (abbreviated as duct) or fibrotic regions (collagenous without duct present, abbreviated as Fibro). Alignment and orientation measures are taken with respect to the duct and are therefore only recorded for ductal regions. Significant p-values are displayed in bold.

| All Women | | | | | | | | | | | | | | |
|---|---|---|---|---|---|---|---|---|---|---|---|---|---|---|
| **Comparison** | **n1** | | **n2** | | **Width** | | **Length** | | **Curvature** | | **Density** | | **Alignment** | **Orientation** |
| Category 1 v. 2 | Duct. | Fibro. | Duct. | Fibro. | Duct. | Fibro. | Duct. | Fibro. | Duct. | Fibro. | Duct. | Fibro. | Duct. | Duct. |
| BIRADS 1 v. BIRADS 2 | 17 | 19 | 18 | 19 | 0.888 | 0.320 | 0.739 | 0.518 | 0.690 | 0.362 | 0.990 | 0.242 | 0.476 | ***0.025*** |
| Pre- v. post-menopausal | 10 | 11 | 25 | 27 | 0.718 | 0.766 | 0.386 | 0.876 | 0.897 | 0.178 | 0.512 | 0.480 | 0.121 | 0.793 |
| Age: <55 v. >55 years old | 17 | 17 | 19 | 22 | ***0.037*** | 0.334 | 0.241 | 0.774 | 0.803 | 0.800 | ***0.045*** | 0.118 | 0.444 | 0.956 |
| BMI: Average v. Overweight | 9 | 8 | 12 | 14 | 0.743 | 0.224 | 0.277 | 0.301 | 0.169 | 0.671 | 0.097 | 0.314 | 0.133 | 0.200 |
| BMI: Average v. Obese | 9 | 14 | 12 | 16 | 0.902 | 0.112 | 0.619 | 0.341 | 0.310 | 0.931 | ***0.041*** | 0.380 | 0.551 | 0.951 |
| BMI: Overweight v. Obese | 12 | 14 | 12 | 16 | 0.610 | 0.807 | 0.503 | 0.929 | 0.814 | 0.687 | 0.821 | 0.998 | 0.450 | 0.187 |
| Black Women: Overweight v. Obese | 6 | 6 | 5 | 7 | ***0.008*** | ***0.017*** | 0.131 | 0.472 | 0.370 | ***0.031*** | 0.297 | 0.841 | 0.950 | 0.521 |
| White Women: Overweight v. Obese | 6 | 8 | 9 | 9 | ***0.012*** | ***0.038*** | 0.423 | 0.531 | 0.626 | 0.412 | 0.446 | 0.805 | 0.415 | 0.355 |
| Density: Heterogeneous v. Scattered Fibro | 20 | 22 | 10 | 11 | ***0.006*** | ***0.001*** | 0.302 | 0.713 | 0.833 | 0.979 | 0.470 | 0.199 | 0.655 | 0.281 |
| Genetic Ancestry: African v. European | 18 | 19 | 18 | 20 | 0.543 | 0.754 | 0.359 | 0.810 | 0.538 | 0.763 | 0.882 | 543 | 0.800 | 0.239 |

**Supplemental Table 4. Identified peptide domains by LC-MS/MS that match MALDI-MSI within 10 ppm that are differentially expressed across BMI categories.**

| m/z | Gene | Sequence with modification |
|---|---|---|
| 1041.541 | COL1A1 | GAVGAKGEAGPQ |
| 1267.595 | COL1A1 | GPAGEKGSPGADGPA |
| 1034.575 | COL1A1 | GPVGPVGARGPA |
| 981.431 | COL1A1 | GPPSAGFDFS |
| 1125.527 | COL1A1 | GPAGERGEQGPA |
| 1115.543 | COL1A1 | SGLDGAKGDAGPA |
| 1283.635 | COL1A2 | GPAGARGSDGSVGPV |
| 1257.582 | COL1A2 | GPAGATGDRGEAGAA |
| 1098.588 | COL1A2 | GPVGRTGEVGAV |
| 1323.775 | COL3A1 | GPLGIAGITGARGLA |
| 1139.654 | COL3A1 | GPLGIAGITGARG |
| 1082.632 | COL3A1 | PLGIAGITGARG |
| 1082.632 | COL3A1 | GPLGIAGITGAR |
| 855.494 | COL3A1 | GPLGIAGITG |
| 1313.567 | COL1A1 | GP(0.0224)P(0.9828)GSP(0.9826)GEQGP(0.0121)SGAS **OR** GP(0.9697)P(0.0436)GSP(0.9708)GEQGP(0.0159)SGAS |
| 1415.618 | COL1A1 | GFQGP(0.0108)P(0.0109)GEP(0.9891)GEP(0.9892)GAS |
| 1113.520 | COL1A1 | GP(0.0054)SGEP(0.9892)GKQGP(0.0055)S |
| 1383.652 | COL1A1 | GSP(0.0229)GEAGRP(0.9886)GEAGLP(0.9885) |
| 1552.810 | COL1A1 | GP(0.0138)IGNVGAP(0.9862)GAKGARGSA |
| 1139.656 | COL1A1 | GIAGQRGVVGLP(1.0000) |
| 1343.672 | COL1A1 | GAP(1.0000)GIAGAP(1.0000)GFP(1.0000)GAR |
| 1421.789 | COL1A1 | GP(0.0108)QGIAGQRGVVGLP(0.9892) |
| 964.482 | COL1A1 | GP(0.0033)P(0.0033)GLAGP(0.0037)P(0.9897)GE |
| 1082.633 | COL1A1 | IAGQRGVVGLP(1.0000) |
| 1286.655 | COL1A1 | IAGAP(0.9852)GFP(0.9851)GARGP(0.0297)S |
| 1415.618 | COL1A2 | GVKGEP(1.0000)GAP(1.0000)GEN*GTP(1.0000) |
| 1552.81 | COL1A2 | GP(0.0159)VGAAGATGARGLVGEP(0.9841) |
| 1884.864 | COL1A2 | GP(0.0064)AGSKGESGNKGEP(0.9883)GSAGP(0.0053)Q |
| 899.444 | COL1A2 | TGEVGAVGP(0.0150)P(0.9850) |
| 1154.614 | COL1A2 | GP(0.0108)RGEVGLP(0.9892)GLS |
| 794.443 | COL1A2 | GP(0.0052)LGIAGP(0.0064)P(0.9884) |
| 1123.559 | COL1A2 | GP(0.0212)P(0.9837)GTP(0.9841)GP(0.0110)Q*GLL |
| 1310.641 | COL1A2 | GP(0.0133)IGSAGP(0.0456)P(0.9597)GFP(0.0194)GAP(0.9620) |
| 1437.645 | COL1A2 | GM(1.0000)TGFP(1.0000)GAAGRTGP(1.0000)P(1.0000) |
| 1128.53 | COL1A2 | GSP(0.9876)GERGEVGP(0.0124)A |
| 1128.528 | COL3A1 | GAN*GLSGERGP(0.0287)P(0.9713) **OR** GAN*GLSGERGP(0.8547)P(0.1453) |

### Supplemental Table 3. Col1a1 peptide sequences from normal breast by LC-MS/MS.

| M+H | Gene | Domain Start | Domain End | Peptide | Modified Peptide [113] means HYP | Sequence with HYP probability PM:15.9949 | Hyperscore | Nextscore | Peptide Prophet Probability |
|---|---|---|---|---|---|---|---|---|---|
| 730.303 | COL1A1 | 1193 | 1199 | SAGFDFS | SAGFDFS | | 18.64 | 10.72 | 0.967 |
| 747.327 | COL1A1 | 1190 | 1197 | GPPSAGFD | GPPSAGFD | | 20.41 | 17.89 | 0.947 |
| 761.381 | COL1A1 | 405 | 412 | IAGAPGFP | IAGAP[113]GFP[113] | IAGAP(1.0000)GFP(1.0000) | 18.66 | 12.38 | 0.991 |
| 764.352 | COL1A1 | 872 | 880 | GATGFPGAA | GATGFP[113]GAA | GATGFP(1.0000)GAA | 18.25 | 14.78 | 0.994 |
| 767.428 | COL1A1 | 765 | 772 | LTGPIGPP | LTGPIGPP[113] | LTGP(0.0058)IGP(0.0064)P(0.9879) | 19.86 | 18.24 | 0.932 |
| 769.350 | COL1A1 | 887 | 895 | GPSGNAGPP | GPSGNAGPP[113] | GP(0.0053)SGNAGP(0.0062)P(0.9885) | 20.45 | 18.65 | 1.000 |
| 770.381 | COL1A1 | 809 | 817 | GPAGFAGPP | GPAGFAGPP | | 19.05 | 12.66 | 0.991 |
| 773.388 | COL1A1 | 174 | 181 | GISVPGPM | GISVPGPM[147] | GISVP(0.0057)GP(0.0100)M(0.9842) | 18.02 | 16.31 | 0.994 |
| 781.418 | COL1A1 | 851 | 859 | GPIGNVGAP | GPIGNVGAP | | 19.07 | 17.28 | 0.934 |
| 786.376 | COL1A1 | 809 | 817 | GPAGFAGPP | GPAGFAGPP[113] | GP(0.0054)AGFAGP(0.0070)P(0.9876) | 21.62 | 21.43 | 0.961 |
| 797.411 | COL1A1 | 851 | 859 | GPIGNVGAP | GPIGNVGAP[113] | GP(0.0105)IGNVGAP(0.9895) | 21.19 | 17.54 | 1.000 |
| 797.428 | COL1A1 | 878 | 886 | GAAGRVGPP | GAAGRVGPP[113] | GAAGRVGP(0.4770)P(0.5230) | 18.54 | 17.13 | 0.991 |
| 798.398 | COL1A1 | 851 | 859 | GPIGNVGAP | GPIGN[115]VGAP[113] | GP(0.0104)IGNVGAP(0.9896) | 19.43 | 14.64 | 0.958 |
| 802.410 | COL1A1 | 404 | 412 | GIAGAPGFP | GIAGAPGFP[113] | GIAGAP(0.0101)GFP(0.9899) | 18.83 | 14.08 | 0.976 |
| 806.366 | COL1A1 | 494 | 502 | GFPGADGVA | GFP[113]GADGVA | GFP(1.0000)GADGVA | 18.70 | 12.33 | 0.987 |
| 814.409 | COL1A1 | 341 | 349 | GPPGFPGAV | GPP[113]GFPGAV | GP(0.0062)P(0.9879)GFP(0.0058)GAV | 19.90 | 18.19 | 0.938 |
| 814.428 | COL1A1 | 254 | 262 | GLPGTAGLP | GLP[113]GTAGLP[113] | GLP(1.0000)GTAGLP(1.0000) | 21.38 | 13.74 | 0.975 |
| 818.396 | COL1A1 | 405 | 413 | IAGAPGFPG | IAGAP[113]GFP[113]G | IAGAP(1.0000)GFP(1.0000)G | 20.62 | 11.50 | 0.970 |
| 819.386 | COL1A1 | 1113 | 1120 | FSGLQGPP | FSGLQ[129]GPP[113] | FSGLQGP(0.0122)P(0.9878) | 18.05 | 16.20 | 0.967 |
| 823.426 | COL1A1 | 452 | 460 | GPVGVQGPP | GPVGVQGPP[113] | GP(0.0052)VGVQGP(0.0055)P(0.9893) | 21.15 | 21.15 | 0.920 |
| 827.353 | COL1A1 | 1192 | 1199 | PSAGFDFS | PSAGFDFS | | 18.55 | 11.91 | 1.000 |
| 829.369 | COL1A1 | 198 | 205 | PQGFQGPP | PQ[129]GFQ[129]GPP | | 19.44 | 12.13 | 0.994 |
| 830.407 | COL1A1 | 173 | 181 | GGISVPGPM | GGISVPGPM[147] | GGISVP(0.0050)GP(0.0103)M(0.9848) | 18.08 | 16.23 | 0.962 |
| 841.405 | COL1A1 | 548 | 556 | GPDGKTGPP | GPDGKTGPP[113] | GP(0.0050)DGKTGP(0.0082)P(0.9869) | 20.08 | 18.39 | 0.969 |
| 856.408 | COL1A1 | 473 | 481 | GEPGPTGLP | GEP[113]GPTGLP[113] | GEP(0.9872)GP(0.0256)TGLP(0.9872) | 19.04 | 14.34 | 0.946 |
| 856.448 | COL1A1 | 346 | 355 | PGAVGAKGEA | PGAVGAKGEA | | 24.85 | 13.88 | 1.000 |
| 858.394 | COL1A1 | 274 | 283 | DGAKGDAGPA | DGAKGDAGPA | | 20.74 | 18.74 | 0.952 |
| 863.399 | COL1A1 | 488 | 496 | GGPGSRGFP | GGP[113]GSRGFP[113] | GGP(1.0000)GSRGFP(1.0000) | 18.09 | 10.12 | 0.995 |
| 871.427 | COL1A1 | 341 | 350 | GPPGFPGAVG | GPP[113]GFPGAVG | GP(0.0063)P(0.9887)GFP(0.0050)GAVG | 21.46 | 19.95 | 0.988 |
| 875.390 | COL1A1 | 722 | 730 | GAPGLQGMP | GAP[113]GLQGM[147]P[113] | GAP(1.0000)GLQGM(1.0000)P(1.0000) | 19.23 | 13.53 | 0.936 |
| 875.426 | COL1A1 | 404 | 413 | GIAGAPGFPG | GIAGAP[113]GFP[113]G | GIAGAP(1.0000)GFP(1.0000)G | 19.04 | 14.11 | 0.957 |
| 876.374 | COL1A1 | 722 | 730 | GAPGLQGMP | GAP[113]GLQ[129]GM[147]P[113] | GAP(1.0000)GLQGM(1.0000)P(1.0000) | 20.05 | 10.28 | 0.999 |
| 876.405 | COL1A1 | 1112 | 1120 | GFSGLQGPP | GFSGLQ[129]GPP[113] | GFSGLQGP(0.0124)P(0.9876) | 20.79 | 18.85 | 0.964 |
| 879.397 | COL1A1 | 575 | 583 | GQAGVMGFP | GQAGVMGFP[113] | GQAGVM(0.0098)GFP(0.9902) | 19.26 | 16.21 | 0.953 |
| 883.428 | COL1A1 | 1187 | 1196 | GPPGPPSAGF | GPPGPPSAGF | | 21.58 | 14.91 | 0.994 |
| 885.408 | COL1A1 | 197 | 205 | GPQGFQGPP | GPQGFQ[129]GPP | | 22.90 | 14.76 | 0.989 |
| 886.392 | COL1A1 | 197 | 205 | GPQGFQGPP | GPQ[129]GFQ[129]GPP | | 19.65 | 13.51 | 0.961 |
| 887.423 | COL1A1 | 341 | 350 | GPPGFPGAVG | GPP[113]GFP[113]GAVG | GP(0.0611)P(0.9693)GFP(0.9696)GAVG | 22.04 | 21.73 | 0.961 |
| 891.400 | COL1A1 | 638 | 646 | GSPGFQGLP | GSP[113]GFQGLP[113] | GSP(1.0000)GFQGLP(1.0000) | 20.96 | 17.78 | 0.955 |
| 892.400 | COL1A1 | 638 | 646 | GSPGFQGLP | GSP[113]GFQ[129]GLP[113] | GSP(1.0000)GFQGLP(1.0000) | 19.33 | 15.40 | 0.974 |
| 894.392 | COL1A1 | 1190 | 1198 | GPPSAGFDF | GPPSAGFDF | | 25.66 | 11.95 | 1.000 |
| 895.381 | COL1A1 | 575 | 583 | GQAGVMGFP | GQAGVM[147]GFP[113] | GQAGVM(1.0000)GFP(1.0000) | 24.38 | 11.07 | 1.000 |
| 899.423 | COL1A1 | 1187 | 1196 | GPPGPPSAGF | GPP[113]GPPSAGF | GP(0.0036)P(0.9891)GP(0.0036)P(0.0037)SAGF | 23.26 | 21.08 | 0.961 |
| 902.383 | COL1A1 | 197 | 205 | GPQGFQGPP | GPQ[129]GFQ[129]GPP[113] | GP(0.0053)QGFQGP(0.0069)P(0.9877) | 18.36 | 18.24 | 0.949 |
| 903.425 | COL1A1 | 151 | 160 | GPPGLGGNFA | GPP[113]GLGGN[115]FA | GP(0.0115)P(0.9885)GLGGNFA | 18.38 | 16.24 | 1.000 |
| 910.389 | COL1A1 | 1190 | 1198 | GPPSAGFDF | GPP[113]SAGFDF | GP(0.0112)P(0.9888)SAGFDF | 18.43 | 16.50 | 0.985 |
| 911.445 | COL1A1 | 850 | 859 | PGPIGNVGAP | P[113]GPIGN[115]VGAP[113] | P(0.9891)GP(0.0218)IGNVGAP(0.9891) | 18.25 | 14.49 | 0.961 |
| 915.460 | COL1A1 | 172 | 181 | TGGISVPGPM | TGGISVPGPM | | 19.71 | 8.15 | 1.000 |
| 924.414 | COL1A1 | 1191 | 1199 | PPSAGFDFS | PPSAGFDFS | | 18.10 | 9.69 | 1.000 |
| 927.439 | COL1A1 | 661 | 670 | QGVPGDLGAP | Q[129]GVPGDLGAP[113] | QGVP(0.0289)GDLGAP(0.9711) | 18.08 | 12.15 | 0.975 |
| 931.450 | COL1A1 | 172 | 181 | TGGISVPGPM | TGGISVPGPM[147] | TGGISVP(0.0055)GP(0.0424)M(0.9521) | 18.83 | 17.68 | 1.000 |
| 941.476 | COL1A1 | 175 | 184 | ISVPGPMGPS | ISVPGPMGPS | | 19.10 | 10.50 | 0.999 |
| 942.448 | COL1A1 | 661 | 670 | QGVPGDLGAP | QGVP[113]GDLGAP[113] | QGVP(1.0000)GDLGAP(1.0000) | 19.79 | 13.29 | 0.977 |
| 943.410 | COL1A1 | 197 | 206 | GPQGFQGPPG | GPQ[129]GFQ[129]GPPG | | 25.60 | 18.40 | 0.955 |
| 946.460 | COL1A1 | 404 | 414 | GIAGAPGFPGA | GIAGAP[113]GFP[113]GA | GIAGAP(1.0000)GFP(1.0000)GA | 19.22 | 12.33 | 0.981 |
| 947.374 | COL1A1 | 208 | 217 | PGEPGASGPM | P[113]GEP[113]GASGPM[147] | P(0.9864)GEP(0.9864)GASGP(0.0409)M(0.9864) | 23.09 | 21.17 | 0.984 |
| 948.440 | COL1A1 | 638 | 647 | GSPGFQGLPG | GSP[113]GFQGLP[113]G | GSP(1.0000)GFQGLP(1.0000)G | 18.46 | 15.50 | 0.958 |
| 955.488 | COL1A1 | 1200 | 1207 | FLPQPPQE | FLPQPPQE | | 19.83 | 11.72 | 0.986 |
| 956.474 | COL1A1 | 1200 | 1207 | FLPQPPQE | FLPQPPQ[129]E | | 19.84 | 12.79 | 0.998 |
| 957.468 | COL1A1 | 175 | 184 | ISVPGPMGPS | ISVPGPM[147]GPS | ISVP(0.0035)GP(0.0048)M(0.9879)GP(0.0037)S | 22.24 | 18.46 | 1.000 |
| 959.411 | COL1A1 | 197 | 206 | GPQGFQGPPG | GPQ[129]GFQ[129]GPP[113]G | GP(0.0056)QGFQGP(0.0073)P(0.9871)G | 20.13 | 18.29 | 0.946 |
| 964.473 | COL1A1 | 1001 | 1011 | GPPGLAGPPGE | GPPGLAGPP[113]GE | GP(0.0033)P(0.0033)GLAGP(0.0037)P(0.9897)GE | 26.67 | 22.45 | 0.982 |
| 966.449 | COL1A1 | 270 | 279 | FSGLDGAKGD | FSGLDGAKGD | | 24.31 | 12.14 | 1.000 |
| 971.476 | COL1A1 | 273 | 283 | LDGAKGDAGPA | LDGAKGDAGPA | | 21.32 | 12.56 | 0.987 |
| 981.410 | COL1A1 | 1190 | 1199 | GPPSAGFDFS | GPPSAGFDFS | | 26.01 | 14.81 | 1.000 |
| 984.496 | COL1A1 | 1150 | 1159 | PGKDGLNGLP | PGKDGLN[115]GLP[113] | P(0.0126)GKDGLNGLP(0.9874) | 18.12 | 13.41 | 0.955 |
| 984.549 | COL1A1 | 760 | 769 | DGVRGLTGPI | DGVRGLTGPI | | 19.82 | 11.64 | 0.990 |
| 985.453 | COL1A1 | 398 | 409 | GANGAPGIAGAP | GAN[115]GAP[113]GIAGAP[113] | GANGAP(1.0000)GIAGAP(1.0000) | 21.36 | 13.30 | 0.978 |
| 997.419 | COL1A1 | 1190 | 1199 | GPPSAGFDFS | GPP[113]SAGFDFS | GP(0.0113)P(0.9887)SAGFDFS | 30.26 | 27.65 | 1.000 |
| 998.450 | COL1A1 | 1187 | 1197 | GPPGPPSAGFD | GPPGPPSAGFD | | 28.97 | 18.69 | 1.000 |
| 1000.476 | COL1A1 | 1196 | 1203 | FDFSFLPQ | FDFSFLPQ | | 18.56 | 9.96 | 0.994 |
| 1002.490 | COL1A1 | 171 | 181 | STGGISVPGPM | STGGISVPGPM | | 18.23 | 11.59 | 0.991 |
| 1014.447 | COL1A1 | 1187 | 1197 | GPPGPPSAGFD | GPP[113]GPPSAGFD | GP(0.0053)P(0.9876)GP(0.0035)P(0.0035)SAGFD | 23.82 | 23.51 | 0.996 |
| 1015.437 | COL1A1 | 196 | 205 | PGPQGFQGPP | P[113]GPQ[129]GFQ[129]GPP[113] | P(0.9879)GP(0.0110)QGFQGP(0.0132)P(0.9878) | 21.40 | 19.47 | 0.993 |
| 1017.467 | COL1A1 | 320 | 331 | GARGNDGATGAA | GARGNDGATGAA | | 18.48 | 11.02 | 0.986 |
| 1018.458 | COL1A1 | 320 | 331 | GARGNDGATGAA | GARGN[115]DGATGAA | | 18.81 | 12.53 | 0.997 |
| 1018.483 | COL1A1 | 171 | 181 | STGGISVPGPM | STGGISVPGP[113]M | STGGISVP(0.0041)GP(0.4980)M(0.4980) | 18.53 | 18.53 | 1.000 |
| 1028.493 | COL1A1 | 272 | 283 | GLDGAKGDAGPA | GLDGAKGDAGPA | | 21.06 | 17.92 | 0.977 |
| 1030.443 | COL1A1 | 1187 | 1197 | GPPGPPSAGFD | GPP[113]GPP[113]SAGFD | GP(0.0110)P(0.9886)GP(0.0118)P(0.9886)SAGFD | 21.96 | 19.95 | 0.996 |
| 1034.573 | COL1A1 | 1076 | 1087 | GPVGPVGARGPA | GPVGPVGARGPA | | 21.55 | 12.82 | 0.980 |

### Supplemental Table 3. Col1a1 peptide sequences from normal breast by LC-MS/MS.

| | | | | | | | | | |
|---|---|---|---|---|---|---|---|---|---|
| 1037.482 | COL1A1 | 270 | 280 | FSGLDGAKGDA | FSGLDGAKGDA | | 20.13 | 12.38 | 0.983 |
| 1041.529 | COL1A1 | 347 | 358 | GAVGAKGEAGPQ | GAVGAKGEAGPQ | | 27.59 | 15.29 | 0.998 |
| 1042.475 | COL1A1 | 932 | 943 | GEKGSPGADGPA | GEKGSPGADGPA | | 21.63 | 13.53 | 0.981 |
| 1042.513 | COL1A1 | 347 | 358 | GAVGAKGEAGPQ | GAVGAKGEAGPQ[129] | | 25.64 | 14.56 | 1.000 |
| 1043.507 | COL1A1 | 1145 | 1156 | GSAGAPGKDGLN | GSAGAPGKDGLN | | 19.03 | 13.57 | 0.991 |
| 1043.509 | COL1A1 | 401 | 412 | GAPGIAGAPGFP | GAP[113]GIAGAPGFP[113] | GAP(0.9888)GIAGAP(0.0224)GFP(0.9888) | 20.97 | 14.98 | 0.988 |
| 1044.494 | COL1A1 | 1145 | 1156 | GSAGAPGKDGLN | GSAGAPGKDGLN[115] | | 19.71 | 9.94 | 1.000 |
| 1045.544 | COL1A1 | 405 | 415 | IAGAPGFPGAR | IAGAP[113]GFP[113]GAR | IAGAP(1.0000)GFP(1.0000)GAR | 19.91 | 15.87 | 0.988 |
| 1057.484 | COL1A1 | 608 | 619 | GPAGKDGEAGAQ | GPAGKDGEAGAQ | | 23.27 | 9.80 | 1.000 |
| 1058.472 | COL1A1 | 776 | 787 | GAPGDKGESGPS | GAPGDKGESGPS | | 19.11 | 12.43 | 1.000 |
| 1059.504 | COL1A1 | 1145 | 1156 | GSAGAPGKDGLN | GSAGAP[113]GKDGLN | GSAGAP(1.0000)GKDGLN | 19.15 | 13.84 | 1.000 |
| 1059.505 | COL1A1 | 401 | 412 | GAPGIAGAPGFP | GAP[113]GIAGAP[113]GFP[113] | GAP(1.0000)GIAGAP(1.0000)GFP(1.0000) | 35.88 | 17.42 | 1.000 |
| 1060.524 | COL1A1 | 536 | 547 | GAKGLTGSPGSP | GAKGLTGSP[113]GSP[113] | GAKGLTGSP(1.0000)GSP(1.0000) | 21.38 | 13.16 | 0.981 |
| 1066.637 | COL1A1 | 954 | 964 | IAGQRGVVGLP | IAGQRGVVGLP | | 18.72 | 14.08 | 0.977 |
| 1067.625 | COL1A1 | 954 | 964 | IAGQRGVVGLP | IAGQ[129]RGVVGLP | | 19.94 | 11.16 | 0.985 |
| 1071.492 | COL1A1 | 660 | 670 | EQGVPGDLGAP | EQGVP[113]GDLGAP[113] | EQGVP(1.0000)GDLGAP(1.0000) | 20.90 | 14.15 | 0.973 |
| 1072.455 | COL1A1 | 197 | 207 | GPQGFQGPPGE | GPQ[129]GFQ[129]GPPGE | | 22.25 | 15.16 | 0.987 |
| 1074.465 | COL1A1 | 776 | 787 | GAPGDKGESGPS | GAP[113]GDKGESGPS | GAP(0.9858)GDKGESGP(0.0142)S | 20.87 | 13.65 | 0.991 |
| 1076.553 | COL1A1 | 872 | 883 | GATGFPGAAGRV | GATGFP[113]GAAGRV | GATGFP(1.0000)GAAGRV | 19.91 | 15.80 | 1.000 |
| 1078.477 | COL1A1 | 1189 | 1199 | PGPPSAGFDFS | PGPPSAGFDFS | | 18.31 | 9.94 | 0.997 |
| 1082.623 | COL1A1 | 954 | 964 | IAGQRGVVGLP | IAGQRGVVGLP[113] | IAGQRGVVGLP(1.0000) | 21.83 | 16.05 | 0.977 |
| 1083.610 | COL1A1 | 954 | 964 | IAGQRGVVGLP | IAGQ[129]RGVVGLP[113] | IAGQRGVVGLP(1.0000) | 23.71 | 13.38 | 0.999 |
| 1084.500 | COL1A1 | 986 | 997 | GPSGASGERGPP | GPSGASGERGPP[113] | GP(0.0057)SGASGERGP(0.0080)P(0.9863) | 22.17 | 22.17 | 0.999 |
| 1088.447 | COL1A1 | 197 | 207 | GPQGFQGPPGE | GPQ[129]GFQ[129]GPP[113]GE | GP(0.0053)QGFQGP(0.0071)P(0.9876)GE | 21.21 | 17.39 | 0.939 |
| 1094.477 | COL1A1 | 1189 | 1199 | PGPPSAGFDFS | P[113]GPPSAGFDFS | P(0.9897)GP(0.0052)P(0.0051)SAGFDFS | 20.64 | 15.94 | 1.000 |
| 1094.512 | COL1A1 | 269 | 280 | GFSGLDGAKGDA | GFSGLDGAKGDA | | 21.65 | 7.48 | 1.000 |
| 1097.517 | COL1A1 | 977 | 988 | GPSGEPGKQGPS | GPSGEPGKQGPS | | 22.27 | 22.27 | 1.000 |
| 1098.505 | COL1A1 | 977 | 988 | GPSGEPGKQGPS | GPSGEPGKQ[129]GPS | | 19.88 | 19.88 | 0.938 |
| 1098.592 | COL1A1 | 251 | 262 | GARGLPGTAGLP | GARGLP[113]GTAGLP[113] | GARGLP(1.0000)GTAGLP(1.0000) | 20.58 | 11.40 | 0.998 |
| 1102.563 | COL1A1 | 404 | 415 | GIAGAPGFPGAR | GIAGAP[113]GFP[113]GAR | GIAGAP(1.0000)GFP(1.0000)GAR | 19.93 | 9.48 | 1.000 |
| 1106.525 | COL1A1 | 491 | 502 | GSRGFPGADGVA | GSRGFP[113]GADGVA | GSRGFP(1.0000)GADGVA | 19.79 | 10.63 | 1.000 |
| 1110.514 | COL1A1 | 1007 | 1018 | GPPGESGREGAP | GPPGESGREGAP | | 20.38 | 12.77 | 0.982 |
| 1112.513 | COL1A1 | 659 | 670 | GEQGVPGDLGAP | GEQGVPGDLGAP[113] | GEQGVP(0.0165)GDLGAP(0.9835) | 23.21 | 14.58 | 0.985 |
| 1112.547 | COL1A1 | 1148 | 1159 | GAPGKDGLNGLP | GAPGKDGLN[115]GLP[113] | GAP(0.0107)GKDGLNGLP(0.9893) | 23.61 | 13.15 | 0.999 |
| 1113.492 | COL1A1 | 659 | 670 | GEQGVPGDLGAP | GEQ[129]GVPGDLGAP[113] | GEQGVP(0.0106)GDLGAP(0.9894) | 23.71 | 14.34 | 0.998 |
| 1113.514 | COL1A1 | 977 | 988 | GPSGEPGKQGPS | GPSGEP[113]GKQGPS | GP(0.0055)SGEP(0.9841)GKQGP(0.0104)S | 24.06 | 24.06 | 0.972 |
| 1114.501 | COL1A1 | 977 | 988 | GPSGEPGKQGPS | GPSGEP[113]GKQ[129]GPS | GP(0.0051)SGEP(0.9874)GKQGP(0.0075)S | 21.20 | 21.20 | 0.981 |
| 1115.525 | COL1A1 | 271 | 283 | SGLDGAKGDAGPA | SGLDGAKGDAGPA | | 26.44 | 17.24 | 1.000 |
| 1115.540 | COL1A1 | 1031 | 1042 | GAKGDRGETGPA | GAKGDRGETGPA | | 19.88 | 9.57 | 0.990 |
| 1116.527 | COL1A1 | 635 | 646 | GPAGSPGFQGLP | GPAGSP[113]GFQGLP[113] | GP(0.0232)AGSP(0.9884)GFQGLP(0.9884) | 19.06 | 14.27 | 0.979 |
| 1117.498 | COL1A1 | 635 | 646 | GPAGSPGFQGLP | GPAGSP[113]GFQ[129]GLP[113] | GP(0.0208)AGSP(0.9896)GFQGLP(0.9896) | 21.06 | 17.67 | 0.996 |
| 1118.473 | COL1A1 | 1019 | 1030 | GAEGSPGRDGSP | GAEGSP[113]GRDGSP[113] | GAEGSP(1.0000)GRDGSP(1.0000) | 18.80 | 12.15 | 0.984 |
| 1122.557 | COL1A1 | 914 | 925 | GPAGRPGEVGPP | GPAGRP[113]GEVGPP[113] | GP(0.0116)AGRP(0.9881)GEVGP(0.0122)P(0.9881) | 21.37 | 21.37 | 0.922 |
| 1125.525 | COL1A1 | 626 | 637 | GPAGERGEQGPA | GPAGERGEQGPA | | 22.35 | 22.35 | 0.957 |
| 1126.507 | COL1A1 | 1007 | 1018 | GPPGESGREGAP | GPPGESGREGAP[113] | GP(0.0063)P(0.0076)GESGREGAP(0.9861) | 21.48 | 13.18 | 0.967 |
| 1128.496 | COL1A1 | 1190 | 1200 | GPPSAGFDFSF | GPPSAGFDFSF | | 25.93 | 11.64 | 1.000 |
| 1128.512 | COL1A1 | 659 | 670 | GEQGVPGDLGAP | GEQGVP[113]GDLGAP[113] | GEQGVP(1.0000)GDLGAP(1.0000) | 24.32 | 13.01 | 0.997 |
| 1128.524 | COL1A1 | 151 | 162 | GPPGLGGNFAPQ | GPP[113]GLGGN[115]FAPQ | GP(0.0062)P(0.9879)GLGGNFAP(0.0059)Q | 20.95 | 18.69 | 0.989 |
| 1128.552 | COL1A1 | 1148 | 1159 | GAPGKDGLNGLP | GAP[113]GKDGLN[115]GLP[113] | GAP(1.0000)GKDGLNGLP(1.0000) | 19.77 | 16.88 | 0.926 |
| 1131.536 | COL1A1 | 1061 | 1072 | GKSGDRGETGPA | GKSGDRGETGPA | | 21.20 | 7.42 | 1.000 |
| 1139.656 | COL1A1 | 954 | 965 | IAGQRGVVGLPG | IAGQRGVVGLP[113]G | IAGQRGVVGLP(1.0000)G | 21.49 | 10.78 | 0.998 |
| 1140.636 | COL1A1 | 953 | 964 | GIAGQRGVVGLP | GIAGQ[129]RGVVGLP[113] | GIAGQRGVVGLP(1.0000) | 20.38 | 15.87 | 0.966 |
| 1142.507 | COL1A1 | 1007 | 1018 | GPPGESGREGAP | GPP[113]GESGREGAP[113] | GP(0.0198)P(0.9901)GESGREGAP(0.9901) | 20.04 | 17.43 | 0.980 |
| 1142.544 | COL1A1 | 524 | 535 | GEAGRPGEAGLP | GEAGRP[113]GEAGLP[113] | GEAGRP(1.0000)GEAGLP(1.0000) | 18.32 | 12.86 | 0.945 |
| 1145.520 | COL1A1 | 1187 | 1198 | GPPGPPSAGFDF | GPPGPPSAGFDF | | 24.00 | 17.58 | 0.995 |
| 1154.507 | COL1A1 | 797 | 808 | GAPGDRGEPGPP | GAP[113]GDRGEP[113]GPP[113] | GAP(0.9880)GDRGEP(0.9880)GP(0.0359)P(0.9880) | 20.81 | 20.81 | 0.961 |
| 1154.561 | COL1A1 | 179 | 190 | GPMGPSGPRGLP | GPM[147]GPSGPRGLP[113] | GP(0.0113)M(0.9864)GP(0.0072)SGP(0.0086)RGLP(0.9866) | 20.08 | 18.22 | 0.984 |
| 1154.595 | COL1A1 | 740 | 751 | GPKGDRGDAGPK | GPKGDRGDAGPK | | 19.12 | 15.32 | 0.986 |
| 1155.526 | COL1A1 | 650 | 661 | GPPGEAGKPGEQ | GPP[113]GEAGKP[113]GEQ | GP(0.0205)P(0.9897)GEAGKP(0.9897)GEQ | 23.64 | 20.90 | 0.945 |
| 1156.549 | COL1A1 | 607 | 619 | VGPAGKDGEAGAQ | VGPAGKDGEAGAQ | | 22.49 | 17.68 | 0.967 |
| 1161.512 | COL1A1 | 1187 | 1198 | GPPGPPSAGFDF | GPP[113]GPPSAGFDF | GP(0.0058)P(0.9829)GP(0.0069)P(0.0043)SAGFDF | 32.45 | 30.28 | 1.000 |
| 1163.564 | COL1A1 | 572 | 583 | GARGQAGVMGFP | GARGQAGVMGFP[113] | GARGQAGVM(0.0107)GFP(0.9893) | 20.86 | 11.96 | 0.998 |
| 1164.550 | COL1A1 | 572 | 583 | GARGQAGVMGFP | GARGQ[129]AGVMGFP[113] | GARGQAGVM(0.0107)GFP(0.9893) | 21.07 | 12.49 | 0.998 |
| 1172.575 | COL1A1 | 752 | 764 | GADGSPGKDGVRG | GADGSPGKDGVRG | | 18.76 | 10.35 | 1.000 |
| 1175.544 | COL1A1 | 725 | 736 | GLQGMPGERGAA | GLQGM[147]P[113]GERGAA | GLQGM(1.0000)P(1.0000)GERGAA | 18.86 | 11.15 | 0.965 |
| 1177.508 | COL1A1 | 1187 | 1198 | GPPGPPSAGFDF | GPP[113]GPP[113]SAGFDF | GP(0.0127)P(0.9861)GP(0.0152)P(0.9860)SAGFDF | 24.60 | 22.48 | 0.999 |
| 1179.560 | COL1A1 | 572 | 583 | GARGQAGVMGFP | GARGQAGVM[147]GFP[113] | GARGQAGVM(1.0000)GFP(1.0000) | 22.67 | 12.36 | 0.999 |
| 1179.573 | COL1A1 | 365 | 376 | GPQGVRGEPGPP | GPQGVRGEP[113]GPP[113] | GP(0.0135)QGVRGEP(0.9876)GP(0.0113)P(0.9876) | 23.68 | 23.68 | 0.988 |
| 1180.531 | COL1A1 | 572 | 583 | GARGQAGVMGFP | GARGQ[129]AGVM[147]GFP[113] | GARGQAGVM(1.0000)GFP(1.0000) | 22.83 | 11.28 | 0.972 |
| 1186.528 | COL1A1 | 362 | 373 | GSEGPQGVRGEP | GSEGPQ[129]GVRGEP[113] | GSEGP(0.0193)QGVRGEP(0.9807) | 18.62 | 13.56 | 1.000 |
| 1188.557 | COL1A1 | 752 | 764 | GADGSPGKDGVRG | GADGSP[113]GKDGVRG | GADGSP(1.0000)GKDGVRG | 18.05 | 4.15 | 1.000 |
| 1189.596 | COL1A1 | 1198 | 1207 | FSFLPQPPQE | FSFLPQPPQE | | 20.79 | 12.22 | 0.974 |
| 1196.562 | COL1A1 | 625 | 637 | AGPAGERGEQGPA | AGPAGERGEQGPA | | 20.34 | 20.34 | 0.988 |
| 1205.559 | COL1A1 | 485 | 496 | GERGGPGSRGFP | GERGGP[113]GSRGFP[113] | GERGGP(1.0000)GSRGFP(1.0000) | 20.76 | 10.03 | 1.000 |
| 1212.548 | COL1A1 | 650 | 662 | GPPGEAGKPGEQG | GPP[113]GEAGKP[113]GEQG | GP(0.0210)P(0.9895)GEAGKP(0.9895)GEQG | 21.95 | 19.21 | 0.995 |
| 1215.614 | COL1A1 | 403 | 415 | PGIAGAPGFPGAR | P[113]GIAGAP[113]GFP[113]GAR | P(1.0000)GIAGAP(1.0000)GFP(1.0000)GAR | 28.74 | 16.62 | 1.000 |
| 1223.596 | COL1A1 | 913 | 925 | TGPAGRPGEVGPP | TGPAGRP[113]GEVGP[113]P | TGP(0.0101)AGRP(0.7601)GEVGP(0.6149)P(0.6149) | 22.03 | 22.03 | 0.983 |
| 1227.591 | COL1A1 | 606 | 619 | AVGPAGKDGEAGAQ | AVGPAGKDGEAGAQ | | 23.44 | 11.50 | 1.000 |
| 1229.584 | COL1A1 | 680 | 691 | GFPGERGVQGPP | GFP[113]GERGVQGPP[113] | GFP(0.9161)GERGVQGP(0.1697)P(0.9142) | 24.24 | 24.24 | 0.991 |
| 1230.582 | COL1A1 | 680 | 691 | GFPGERGVQGPP | GFP[113]GERGVQ[129]GP[113]P | GFP(0.7197)GERGVQGP(0.6402)P(0.6402) | 21.18 | 21.18 | 0.988 |
| 1232.547 | COL1A1 | 1187 | 1199 | GPPGPPSAGFDFS | GPPGPPSAGFDFS | | 27.11 | 11.83 | 0.968 |
| 1241.609 | COL1A1 | 395 | 409 | GAKGANGAPGIAGAP | GAKGAN[115]GAP[113]GIAGAP[113] | GAKGANGAP(1.0000)GIAGAP(1.0000) | 19.09 | 12.20 | 0.987 |
| 1242.576 | COL1A1 | 317 | 331 | GPAGARGNDGATGAA | GPAGARGNDGATGAA | | 20.61 | 12.22 | 0.997 |

| | | | | | | | | | |
|---|---|---|---|---|---|---|---|---|---|
| 1243.560 | COL1A1 | 317 | 331 | GPAGARGNDGATGAA | GPAGARGN[115]DGATGAA | | 18.47 | 11.43 | 0.982 |
| 1248.542 | COL1A1 | 1187 | 1199 | GPPGPPSAGFDFS | GPPGPP[113]SAGFDFS | GP(0.0079)P(0.0134)GP(0.3492)P(0.6294)SAGFDFS | 20.31 | 20.29 | 0.977 |
| 1251.650 | COL1A1 | 175 | 187 | ISVPGPMGPSGPR | ISVPGPMGPSGPR | | 18.83 | 9.23 | 0.995 |
| 1262.586 | COL1A1 | 270 | 283 | FSGLDGAKGDAGPA | FSGLDGAKGDAGPA | | 43.16 | 18.37 | 1.000 |
| 1264.539 | COL1A1 | 1187 | 1199 | GPPGPPSAGFDFS | GPP[113]GP[113]PSAGFDFS | GP(0.0134)P(0.9826)GP(0.9823)P(0.0217)SAGFDFS | 19.41 | 18.37 | 1.000 |
| 1267.582 | COL1A1 | 929 | 943 | GPAGEKGSPGADGPA | GPAGEKGSPGADGPA | | 29.11 | 12.24 | 1.000 |
| 1267.649 | COL1A1 | 175 | 187 | ISVPGPMGPSGPR | ISVPGPM[147]GPSGPR | ISVP(0.0028)GP(0.0049)M(0.9860)GP(0.0032)SGP(0.0031)R | 19.87 | 17.53 | 0.995 |
| 1267.716 | COL1A1 | 954 | 966 | IAGQRGVVGLPGQ | IAGQRGVVGLP[113]GQ | IAGQRGVVGLP(1.0000)GQ | 24.39 | 12.30 | 0.998 |
| 1272.533 | COL1A1 | 201 | 213 | FQGPPGEPGEPGA | FQ[129]GPPGEP[113]GEP[113]GA | FQGP(0.0120)P(0.0122)GEP(0.9879)GEP(0.9879)GA | 20.35 | 14.76 | 0.942 |
| 1278.595 | COL1A1 | 270 | 283 | FSGLDGAKGDAGPA | FSGLDGAKGDAGP[113]A | FSGLDGAKGDAGP(1.0000)A | 23.23 | 17.63 | 0.942 |
| 1280.533 | COL1A1 | 1187 | 1199 | GPPGPPSAGFDFS | GP[113]P[113]GPP[113]SAGFDFS | GP(0.9813)P(0.9813)GP(0.0561)P(0.9812)SAGFDFS | 25.32 | 21.53 | 1.000 |
| 1283.568 | COL1A1 | 932 | 946 | GEKGSPGADGPAGAP | GEKGSPGADGP[113]AGAP | GEKGSP(0.0080)GADGP(0.9864)AGAP(0.0056) | 20.05 | 15.79 | 0.991 |
| 1283.582 | COL1A1 | 773 | 787 | GPAGAPGDKGESGPS | GPAGAPGDKGESGPS | | 32.67 | 18.00 | 0.998 |
| 1284.607 | COL1A1 | 605 | 619 | GAVGPAGKDGEAGAQ | GAVGPAGKDGEAGAQ | | 32.77 | 13.40 | 1.000 |
| 1285.595 | COL1A1 | 605 | 619 | GAVGPAGKDGEAGAQ | GAVGPAGKDGEAGAQ[129] | | 23.49 | 15.19 | 0.961 |
| 1286.553 | COL1A1 | 701 | 715 | GAPGNDGAKGDAGAP | GAP[113]GNDGAKGDAGAP[113] | GAP(1.0000)GNDGAKGDAGAP(1.0000) | 18.39 | 11.69 | 0.998 |
| 1286.644 | COL1A1 | 405 | 418 | IAGAPGFPGARGPS | IAGAP[113]GFP[113]GARGPS | IAGAP(0.9826)GFP(0.9825)GARGP(0.0350)S | 18.82 | 18.82 | 0.977 |
| 1287.598 | COL1A1 | 680 | 692 | GFPGERGVQGPPG | GFP[113]GERGVQ[129]GPP[113]G | GFP(0.9682)GERGVQGP(0.0641)P(0.9678)G | 22.46 | 22.46 | 0.999 |
| 1299.575 | COL1A1 | 773 | 787 | GPAGAPGDKGESGPS | GPAGAP[113]GDKGESGPS | GP(0.0053)AGAP(0.9895)GDKGESGP(0.0052)S | 30.00 | 16.85 | 0.993 |
| 1299.575 | COL1A1 | 929 | 943 | GPAGEKGSPGADGPA | GP[113]AGEKGSP[113]GADGPA | GP(0.9852)AGEKGSP(0.9858)GADGP(0.0290)A | 21.02 | 12.88 | 0.980 |
| 1302.587 | COL1A1 | 398 | 412 | GANGAPGIAGAPGFP | GAN[115]GAP[113]GIAGAP[113]GFP[113] | GANGAP(1.0000)GIAGAP(1.0000)GFP(1.0000) | 26.39 | 14.87 | 1.000 |
| 1304.628 | COL1A1 | 677 | 688 | GERGFPGERGVQ | GERGFP[113]GERGVQ | GERGFP(1.0000)GERGVQ | 20.44 | 10.72 | 0.993 |
| 1311.652 | COL1A1 | 1145 | 1159 | GSAGAPGKDGLNGLP | GSAGAPGKDGLN[115]GLP | | 22.42 | 17.68 | 0.961 |
| 1313.551 | COL1A1 | 1121 | 1135 | GPPGSPGEQGPSGAS | GPP[113]GSP[113]GEQGPSGAS | GP(0.0322)P(0.9759)GSP(0.9764)GEQGP(0.0154)SGAS | 24.74 | 24.74 | 0.984 |
| 1313.597 | COL1A1 | 653 | 666 | GEAGKPGEQGVPGD | GEAGKP[113]GEQGVPGD | GEAGKP(0.9875)GEQGVP(0.0125)GD | 19.33 | 12.48 | 0.985 |
| 1315.572 | COL1A1 | 773 | 787 | GPAGAPGDKGESGPS | GPAGAP[113]GDKGESGP[113]S | GP(0.1205)AGAP(0.9403)GDKGESGP(0.9392)S | 18.21 | 10.59 | 0.936 |
| 1319.610 | COL1A1 | 269 | 283 | GFSGLDGAKGDAGPA | GFSGLDGAKGDAGPA | | 40.45 | 23.25 | 1.000 |
| 1322.639 | COL1A1 | 1196 | 1206 | FDFSFLPQPPQ | FDFSFLPQPPQ | | 19.53 | 12.89 | 0.958 |
| 1326.666 | COL1A1 | 1145 | 1159 | GSAGAPGKDGLNGLP | GSAGAPGKDGLNGLP[113] | GSAGAP(0.0189)GKDGLNGLP(0.9811) | 29.12 | 13.86 | 1.000 |
| 1327.640 | COL1A1 | 1145 | 1159 | GSAGAPGKDGLNGLP | GSAGAPGKDGLN[115]GLP[113] | GSAGAP(0.0103)GKDGLNGLP(0.9897) | 18.14 | 10.28 | 0.991 |
| 1333.609 | COL1A1 | 488 | 502 | GGPGSRGFPGADGVA | GGP[113]GSRGFP[113]GADGVA | GGP(1.0000)GSRGFP(1.0000)GADGVA | 19.71 | 12.18 | 0.997 |
| 1335.610 | COL1A1 | 269 | 283 | GFSGLDGAKGDAGPA | GFSGLDGAKGDAGP[113]A | GFSGLDGAKGDAGP(1.0000)A | 27.24 | 13.43 | 1.000 |
| 1342.623 | COL1A1 | 522 | 535 | SPGEAGRPGEAGLP | SP[113]GEAGRP[113]GEAGLP[113] | SP(1.0000)GEAGRP(1.0000)GEAGLP(1.0000) | 18.71 | 10.03 | 0.997 |
| 1342.651 | COL1A1 | 344 | 358 | GFPGAVGAKGEAGPQ | GFPGAVGAKGEAGPQ | | 28.20 | 14.26 | 0.998 |
| 1342.656 | COL1A1 | 1145 | 1159 | GSAGAPGKDGLNGLP | GSAGAP[113]GKDGLNGLP[113] | GSAGAP(1.0000)GKDGLNGLP(1.0000) | 18.29 | 12.75 | 0.937 |
| 1343.621 | COL1A1 | 1145 | 1159 | GSAGAPGKDGLNGLP | GSAGAP[113]GKDGLN[115]GLP[113] | GSAGAP(1.0000)GKDGLNGLP(1.0000) | 24.90 | 13.38 | 1.000 |
| 1343.656 | COL1A1 | 404 | 418 | GIAGAPGFPGARGPS | GIAGAP[113]GFP[113]GARGPS | GIAGAP(0.9632)GFP(0.9627)GARGP(0.0741)S | 28.40 | 22.55 | 0.994 |
| 1343.672 | COL1A1 | 401 | 415 | GAPGIAGAPGFPGAR | GAP[113]GIAGAP[113]GFP[113]GAR | GAP(1.0000)GIAGAP(1.0000)GFP(1.0000)GAR | 35.77 | 21.66 | 1.000 |
| 1350.633 | COL1A1 | 623 | 637 | GPAGPAGERGEQGPA | GPAGPAGERGEQGPA | | 18.09 | 18.09 | 0.959 |
| 1352.589 | COL1A1 | 203 | 217 | GPPGEPGEPGASGPM | GPPGEPGEPGASGP[113]M | GP(0.0033)P(0.0028)GEP(0.0029)GEP(0.0030)GASGP(0.4940)M(0.4940) | 30.09 | 30.09 | 1.000 |
| 1355.608 | COL1A1 | 542 | 556 | GSPGSPGPDGKTGPP | GSP[113]GSP[113]GPDGKTGPP[113] | GSP(0.9856)GSP(0.9855)GP(0.0239)DGKTGP(0.0194)P(0.9856) | 18.82 | 18.82 | 1.000 |
| 1358.577 | COL1A1 | 201 | 214 | FQGPPGEPGEPGAS | FQGPPGEP[113]GEP[113]GAS | FQGP(0.0099)P(0.0099)GEP(0.9901)GEP(0.9901)GAS | 30.04 | 20.84 | 1.000 |
| 1358.668 | COL1A1 | 344 | 358 | GFPGAVGAKGEAGPQ | GFPGAVGAKGEAGP[113]Q | GFP(0.0792)GAVGAKGEAGP(0.9208)Q | 19.96 | 11.71 | 0.998 |
| 1359.637 | COL1A1 | 404 | 418 | GIAGAPGFPGARGPS | GIAGAP[113]GFP[113]GARGP[113]S | GIAGAP(1.0000)GFP(1.0000)GARGP(1.0000)S | 28.48 | 11.76 | 1.000 |
| 1367.651 | COL1A1 | 1004 | 1018 | GLAGPPGESGREGAP | GLAGPPGESGREGAP[113] | GLAGP(0.0071)P(0.0087)GESGREGAP(0.9841) | 30.34 | 21.77 | 1.000 |
| 1368.568 | COL1A1 | 203 | 217 | GPPGEPGEPGASGPM | GPPGEP[113]GEPGASGP[113]M | GP(0.0052)P(0.0047)GEP(0.8117)GEP(0.1022)GASGP(0.5381)M(0.5381) | 29.53 | 29.53 | 0.994 |
| 1368.691 | COL1A1 | 587 | 601 | GAAGEPGKAGERGVP | GAAGEP[113]GKAGERGVP | GAAGEP(0.7636)GKAGERGVP(0.2364) | 18.51 | 12.11 | 0.997 |
| 1371.566 | COL1A1 | 197 | 210 | GPQGFQGPPGEPGE | GPQ[129]GFQ[129]GPPGEP[113]GE | GP(0.0040)QGFQGP(0.0062)P(0.0062)GEP(0.9837)GE | 18.92 | 18.27 | 0.990 |
| 1374.661 | COL1A1 | 344 | 358 | GFPGAVGAKGEAGPQ | GFP[113]GAVGAKGEAGP[113]Q | GFP(1.0000)GAVGAKGEAGP(1.0000)Q | 18.15 | 12.87 | 0.954 |
| 1381.703 | COL1A1 | 248 | 262 | GPQGARGLPGTAGLP | GPQ[129]GARGLP[113]GTAGLP[113] | GP(0.0222)QGARGLP(0.9889)GTAGLP(0.9889) | 21.23 | 13.71 | 0.982 |
| 1383.648 | COL1A1 | 1004 | 1018 | GLAGPPGESGREGAP | GLAGPP[113]GESGREGAP[113] | GLAGP(0.0314)P(0.9843)GESGREGAP(0.9843) | 23.78 | 19.42 | 0.992 |
| 1384.558 | COL1A1 | 203 | 217 | GPPGEPGEPGASGPM | GPPGEP[113]GEP[113]GASGP[113]M | GP(0.0101)P(0.0101)GEP(0.8273)GEP(0.8273)GASGP(0.6626)M(0.6626) | 27.79 | 27.79 | 0.980 |
| 1384.670 | COL1A1 | 587 | 601 | GAAGEPGKAGERGVP | GAAGEP[113]GKAGERGVP[113] | GAAGEP(1.0000)GKAGERGVP(1.0000) | 20.17 | 10.29 | 0.975 |
| 1385.675 | COL1A1 | 726 | 739 | LQGMPGERGAAGLP | LQGMP[113]GERGAAGLP[113] | LQGM(0.0235)P(0.9883)GERGAAGLP(0.9883) | 20.80 | 18.29 | 0.997 |
| 1386.691 | COL1A1 | 752 | 766 | GADGSPGKDGVRGLT | GADGSPGKDGVRGLT | | 21.06 | 8.51 | 1.000 |
| 1394.696 | COL1A1 | 656 | 670 | GKPGEQGVPGDLGAP | GKPGEQGVPGDLGAP[113] | GKP(0.0100)GEQGVP(0.0090)GDLGAP(0.9810) | 21.01 | 15.00 | 0.959 |
| 1397.706 | COL1A1 | 899 | 913 | GPAGKEGGKGPRGET | GPAGKEGGKGPRGET | | 20.06 | 11.89 | 0.989 |
| 1398.603 | COL1A1 | 284 | 298 | GPKGEPGSPGENGAP | GPKGEP[113]GSP[113]GENGAP[113] | GP(0.0321)KGEP(0.9893)GSP(0.9893)GENGAP(0.9893) | 19.69 | 13.98 | 0.985 |
| 1399.584 | COL1A1 | 284 | 298 | GPKGEPGSPGENGAP | GPKGEP[113]GSP[113]GEN[115]GAP[113] | GP(0.0322)KGEP(0.9893)GSP(0.9893)GENGAP(0.9893) | 25.40 | 19.13 | 0.981 |
| 1399.627 | COL1A1 | 521 | 535 | GSPGEAGRPGEAGLP | GSP[113]GEAGRP[113]GEAGLP[113] | GSP(1.0000)GEAGRP(1.0000)GEAGLP(1.0000) | 24.24 | 13.67 | 1.000 |
| 1400.551 | COL1A1 | 203 | 217 | GPPGEPGEPGASGPM | GP[113]P[113]GEP[113]GEP[113]GASGPM | GP(0.6402)P(0.6402)GEP(0.7197)GEP(0.7197)GASGP(0.6402)M(0.6402) | 25.98 | 25.98 | 0.934 |
| 1401.669 | COL1A1 | 726 | 739 | LQGMPGERGAAGLP | LQGM[147]P[113]GERGAAGLP[113] | LQGM(1.0000)P(1.0000)GERGAAGLP(1.0000) | 21.67 | 13.50 | 0.960 |
| 1402.684 | COL1A1 | 752 | 766 | GADGSPGKDGVRGLT | GADGSP[113]GKDGVRGLT | GADGSP(1.0000)GKDGVRGLT | 18.25 | 8.12 | 1.000 |
| 1409.655 | COL1A1 | 911 | 925 | GETGPAGRPGEVGPP | GETGPAGRP[113]GEVGPP[113] | GETGP(0.0104)AGRP(0.9872)GEVGP(0.0152)P(0.9872) | 22.43 | 22.43 | 0.938 |
| 1410.684 | COL1A1 | 656 | 670 | GKPGEQGVPGDLGAP | GKP[113]GEQGVPGDLGAP[113] | GKP(0.9879)GEQGVP(0.0242)GDLGAP(0.9879) | 19.77 | 13.86 | 0.969 |
| 1411.669 | COL1A1 | 656 | 670 | GKPGEQGVPGDLGAP | GKP[113]GEQ[129]GVPGDLGAP[113] | GKP(0.9867)GEQGVP(0.0267)GDLGAP(0.9865) | 18.82 | 12.83 | 0.977 |
| 1412.623 | COL1A1 | 224 | 238 | GPPGKNGDDGEAGKP | GPP[113]GKN[115]GDDGEAGKP | GP(0.0051)P(0.9860)GKNGDDGEAGKP(0.0089) | 29.10 | 25.84 | 1.000 |
| 1414.605 | COL1A1 | 284 | 298 | GPKGEPGSPGENGAP | GP[113]KGEP[113]GSP[113]GENGAP[113] | GP(1.0000)KGEP(1.0000)GSP(1.0000)GENGAP(1.0000) | 21.71 | 11.24 | 0.997 |
| 1415.584 | COL1A1 | 284 | 298 | GPKGEPGSPGENGAP | GP[113]KGEP[113]GSP[113]GEN[115]GAP[113] | GP(1.0000)KGEP(1.0000)GSP(1.0000)GENGAP(1.0000) | 26.64 | 10.96 | 1.000 |
| 1415.610 | COL1A1 | 200 | 214 | GFQGPPGEPGEPGAS | GFQGPPGEP[113]GEP[113]GAS | GFQGP(0.0127)P(0.0129)GEP(0.9872)GEP(0.9872)GAS | 20.23 | 13.44 | 1.000 |
| 1421.779 | COL1A1 | 950 | 964 | GPQGIAGQRGVVGLP | GPQGIAGQRGVVGLP[113] | GP(0.0103)QGIAGQRGVVGLP(0.9897) | 46.36 | 24.00 | 1.000 |
| 1422.765 | COL1A1 | 950 | 964 | GPQGIAGQRGVVGLP | GPQ[129]GIAGQRGVVGLP[113] | GP(0.0132)QGIAGQRGVVGLP(0.9868) | 19.81 | 13.63 | 0.996 |
| 1423.753 | COL1A1 | 950 | 964 | GPQGIAGQRGVVGLP | GPQ[129]GIAGQ[129]RGVVGLP[113] | GP(0.0161)QGIAGQRGVVGLP(0.9839) | 25.87 | 11.79 | 1.000 |
| 1424.656 | COL1A1 | 607 | 622 | VGPAGKDGEAGAQGPP | VGPAGKDGEAGAQ[129]GP[113]P | VGP(0.0042)AGKDGEAGAQGP(0.4979)P(0.4979) | 23.94 | 23.94 | 0.939 |
| 1424.670 | COL1A1 | 650 | 664 | GPPGEAGKPGEQGVP | GPP[113]GEAGKP[113]GEQGVP[113] | GP(0.1147)P(0.9615)GEAGKP(0.9619)GEQGVP(0.9619) | 18.47 | 16.73 | 0.986 |
| 1424.793 | COL1A1 | 954 | 967 | IAGQRGVVGLPGQR | IAGQRGVVGLP[113]GQ[129]R | IAGQRGVVGLP(1.0000)GQR | 21.52 | 9.46 | 1.000 |
| 1426.672 | COL1A1 | 656 | 670 | GKPGEQGVPGDLGAP | GKP[113]GEQGVP[113]GDLGAP[113] | GKP(1.0000)GEQGVP(1.0000)GDLGAP(1.0000) | 36.08 | 19.13 | 0.979 |
| 1427.663 | COL1A1 | 656 | 670 | GKPGEQGVPGDLGAP | GKP[113]GEQ[129]GVP[113]GDLGAP[113] | GKP(1.0000)GEQGVP(1.0000)GDLGAP(1.0000) | 19.42 | 14.46 | 0.949 |
| 1428.622 | COL1A1 | 224 | 238 | GPPGKNGDDGEAGKP | GPP[113]GKN[115]GDDGEAGKP[113] | GP(0.0222)P(0.9889)GKNGDDGEAGKP(0.9889) | 22.57 | 19.15 | 1.000 |
| 1430.685 | COL1A1 | 569 | 583 | GPPGARGQAGVMGFP | GPP[113]GARGQAGVMGFP[113] | GP(0.0325)P(0.9717)GARGQAGVM(0.0232)GFP(0.9725) | 18.62 | 16.84 | 0.994 |
| 1431.646 | COL1A1 | 632 | 646 | GEQGPAGSPGFQGLP | GEQGPAGSP[113]GFQ[129]GLP[113] | GEQGP(0.0335)AGSP(0.9832)GFQGLP(0.9833) | 22.90 | 18.67 | 1.000 |
| 1432.631 | COL1A1 | 632 | 646 | GEQGPAGSPGFQGLP | GEQ[129]GPAGSP[113]GFQ[129]GLP[113] | GEQGP(0.0408)AGSP(0.9795)GFQGLP(0.9797) | 18.31 | 14.59 | 1.000 |
| 1442.701 | COL1A1 | 725 | 739 | GLQGMPGERGAAGLP | GLQGM[147]PGERGAAGLP[113] | GLQGM(0.9850)P(0.0300)GERGAAGLP(0.9850) | 18.22 | 13.88 | 0.923 |

| | | | | | | | | | |
|---|---|---|---|---|---|---|---|---|---|
| 1443.684 | COL1A1 | 725 | 739 | GLQGMPGERGAAGLP | GLQ[129]GMP[113]GERGAAGLP[113] | GLQGM(0.0234)P(0.9883)GERGAAGLP(0.9883) | 20.64 | 15.69 | 0.986 |
| 1446.671 | COL1A1 | 569 | 583 | GPPGARGQAGVMGFP | GPP[113]GARGQAGVM[147]GFP[113] | GP(0.0662)P(0.9779)GARGQAGVM(0.9779)GFP(0.9780) | 19.88 | 17.93 | 0.981 |
| 1447.676 | COL1A1 | 569 | 583 | GPPGARGQAGVMGFP | GPP[113]GARGQ[129]AGVM[147]GFP[113] | GP(0.0727)P(0.9756)GARGQAGVM(0.9758)GFP(0.9758) | 18.63 | 16.69 | 0.993 |
| 1454.707 | COL1A1 | 749 | 764 | GPKGADGSPGKDGVRG | GPKGADGSPGKDGVRG | | 26.13 | 11.19 | 1.000 |
| 1458.685 | COL1A1 | 725 | 739 | GLQGMPGERGAAGLP | GLQGM[147]P[113]GERGAAGLP[113] | GLQGM(1.0000)P(1.0000)GERGAAGLP(1.0000) | 43.01 | 17.59 | 1.000 |
| 1459.678 | COL1A1 | 725 | 739 | GLQGMPGERGAAGLP | GLQ[129]GM[147]P[113]GERGAAGLP[113] | GLQGM(1.0000)P(1.0000)GERGAAGLP(1.0000) | 22.78 | 13.48 | 1.000 |
| 1465.688 | COL1A1 | 650 | 665 | GPPGEAGKPGEQGVPG | GPP[113]GEAGKP[113]GEQGVPG | GP(0.0143)P(0.9874)GEAGKP(0.9876)GEQGVP(0.0107)G | 19.61 | 17.77 | 0.978 |
| 1466.686 | COL1A1 | 1190 | 1203 | GPPSAGFDFSFLPQ | GPPSAGFDFSFLPQ | | 20.17 | 13.90 | 0.955 |
| 1468.692 | COL1A1 | 656 | 671 | GKPGEQGVPGDLGAPG | GKP[113]GEQ[129]GVPGDLGAP[113]G | GKP(0.9899)GEQGVP(0.0206)GDLGAP(0.9895)G | 20.93 | 15.35 | 0.960 |
| 1470.724 | COL1A1 | 749 | 764 | GPKGADGSPGKDGVRG | GPKGADGSP[113]GKDGVRG | GP(0.0114)KGADGSP(0.9886)GKDGVRG | 25.59 | 12.59 | 1.000 |
| 1475.714 | COL1A1 | 268 | 283 | RGFSGLDGAKGDAGPA | RGFSGLDGAKGDAGPA | | 26.45 | 11.49 | 1.000 |
| 1481.694 | COL1A1 | 650 | 665 | GPPGEAGKPGEQGVPG | GPP[113]GEAGKP[113]GEQGVP[113]G | GP(0.0351)P(0.9883)GEAGKP(0.9883)GEQGVP(0.9883)G | 18.68 | 16.84 | 0.968 |
| 1482.708 | COL1A1 | 655 | 670 | AGKPGEQGVPGDLGAP | AGKP[113]GEQ[129]GVPGDLGAP[113] | AGKP(0.9884)GEQGVP(0.0232)GDLGAP(0.9884) | 19.87 | 15.44 | 0.978 |
| 1498.703 | COL1A1 | 655 | 670 | AGKPGEQGVPGDLGAP | AGKP[113]GEQ[129]GVP[113]GDLGAP[113] | AGKP(1.0000)GEQGVP(1.0000)GDLGAP(1.0000) | 18.25 | 11.99 | 0.984 |
| 1529.627 | COL1A1 | 698 | 715 | GANGAPGNDGAKGDAGAP | GAN[115]GAP[113]GNDGAKGDAGAP[113] | GANGAP(1.0000)GNDGAKGDAGAP(1.0000) | 27.22 | 14.44 | 1.000 |
| 1535.731 | COL1A1 | 605 | 622 | GAVGPAGKDGEAGAQGPP | GAVGPAGKDGEAGAQGPP | | 25.58 | 14.11 | 0.971 |
| 1536.730 | COL1A1 | 605 | 622 | GAVGPAGKDGEAGAQGPP | GAVGPAGKDGEAGAQ[129]GPP | | 18.78 | 13.85 | 0.975 |
| 1538.665 | COL1A1 | 1121 | 1138 | GPPGSPGEQGPSGASGPA | GP[113]PGSP[113]GEQGPSGASGPA | GP(0.5874)P(0.5874)GSP(0.8113)GEQGP(0.0065)SGASGP(0.0074)A | 30.35 | 30.35 | 1.000 |
| 1550.799 | COL1A1 | 175 | 190 | ISVPGPMGPSGPRGLP | ISVPGPM[147]GPSGPRGLP[113] | ISVP(0.0100)GP(0.1433)M(0.8944)GP(0.0137)SGP(0.0139)RGLP(0.9246) | 21.36 | 21.33 | 1.000 |
| 1551.724 | COL1A1 | 602 | 619 | GPPGAVGPAGKDGEAGAQ | GPP[113]GAVGPAGKDGEAGAQ | GP(0.0100)P(0.9840)GAVGP(0.0060)AGKDGEAGAQ | 21.93 | 20.03 | 0.965 |
| 1552.708 | COL1A1 | 605 | 622 | GAVGPAGKDGEAGAQGPP | GAVGPAGKDGEAGAQ[129]GPP[113] | GAVGP(0.0074)AGKDGEAGAQGP(0.0084)P(0.9842) | 32.46 | 32.46 | 0.997 |
| 1552.818 | COL1A1 | 851 | 868 | GPIGNVGAPGAKGARGSA | GPIGNVGAP[113]GAKGARGSA | GP(0.0138)IGNVGAP(0.9862)GAKGARGSA | 27.01 | 11.34 | 1.000 |
| 1553.721 | COL1A1 | 167 | 181 | YDEKSTGGISVPGPM | YDEKSTGGISVPGP[113]M | YDEKSTGGISVP(0.0056)GP(0.4972)M(0.4972) | 26.59 | 26.59 | 1.000 |
| 1557.757 | COL1A1 | 395 | 412 | GAKGANGAPGIAGAPGFP | GAKGANGAP[113]GIAGAP[113]GFP[113] | GAKGANGAP(1.0000)GIAGAP(1.0000)GFP(1.0000) | 18.12 | 9.71 | 0.993 |
| 1558.719 | COL1A1 | 1091 | 1105 | GPRGDKGETGEQGDR | GPRGDKGETGEQGDR | | 19.05 | 8.71 | 0.999 |
| 1558.739 | COL1A1 | 395 | 412 | GAKGANGAPGIAGAPGFP | GAKGAN[115]GAP[113]GIAGAP[113]GFP[113] | GAKGANGAP(1.0000)GIAGAP(1.0000)GFP(1.0000) | 24.68 | 10.81 | 1.000 |
| 1562.774 | COL1A1 | 1142 | 1159 | GPPGSAGAPGKDGLNGLP | GPPGSAGAPGKDGLN[115]GLP | | 22.73 | 11.98 | 0.998 |
| 1567.724 | COL1A1 | 824 | 841 | GAKGEPGDAGAKGDAGPP | GAKGEPGDAGAKGDAGP[113]P | GAKGEP(0.0044)GDAGAKGDAGP(0.4978)P(0.4978) | 21.09 | 21.09 | 0.977 |
| 1569.725 | COL1A1 | 540 | 556 | LTGSPGSPGPDGKTGPP | LTGSP[113]GSP[113]GPDGKTGPP[113] | LTGSP(0.9620)GSP(0.9605)GP(0.0961)DGKTGP(0.0194)P(0.9620) | 30.77 | 30.77 | 0.998 |
| 1572.741 | COL1A1 | 677 | 691 | GERGFPGERGVQGPP | GERGFP[113]GERGVQ[129]GPP[113] | GERGFP(0.9864)GERGVQGP(0.0273)P(0.9863) | 20.41 | 18.45 | 0.951 |
| 1574.710 | COL1A1 | 1091 | 1105 | GPRGDKGETGEQGDR | GP[113]RGDKGETGEQGDR | GP(1.0000)RGDKGETGEQGDR | 21.04 | 12.12 | 0.998 |
| 1578.770 | COL1A1 | 1142 | 1159 | GPPGSAGAPGKDGLNGLP | GPPGSAGAPGKDGLN[115]GLP[113] | GP(0.0074)P(0.0077)GSAGAP(0.0130)GKDGLNGLP(0.9718) | 24.33 | 16.57 | 0.941 |
| 1580.708 | COL1A1 | 650 | 666 | GPPGEAGKPGEQGVPGD | GPP[113]GEAGKP[113]GEQGVPGD | GP(0.0144)P(0.9866)GEAGKP(0.9867)GEQGVP(0.0123)GD | 22.54 | 20.55 | 1.000 |
| 1581.699 | COL1A1 | 650 | 666 | GPPGEAGKPGEQGVPGD | GPP[113]GEAGKP[113]GEQ[129]GVPGD | GP(0.0221)P(0.9823)GEAGKP(0.9826)GEQGVP(0.0130)GD | 22.08 | 20.10 | 1.000 |
| 1583.719 | COL1A1 | 824 | 841 | GAKGEPGDAGAKGDAGPP | GAKGEP[113]GDAGAKGDAGPP[113] | GAKGEP(0.9556)GDAGAKGDAGP(0.0896)P(0.9548) | 21.51 | 21.51 | 0.960 |
| 1584.766 | COL1A1 | 401 | 418 | GAPGIAGAPGFPGARGPS | GAP[113]GIAGAP[113]GFP[113]GARGPS | GAP(0.9484)GIAGAP(0.9484)GFP(0.9478)GARGP(0.1553)S | 22.73 | 20.76 | 0.990 |
| 1586.744 | COL1A1 | 398 | 415 | GANGAPGIAGAPGFPGAR | GAN[115]GAP[113]GIAGAP[113]GFP[113]GAR | GANGAP(1.0000)GIAGAP(1.0000)GFP(1.0000)GAR | 30.09 | 17.40 | 1.000 |
| 1588.794 | COL1A1 | 674 | 688 | GARGERGFPGERGVQ | GARGERGFP[113]GERGVQ | GARGERGFP(1.0000)GERGVQ | 18.24 | 11.69 | 0.949 |
| 1593.776 | COL1A1 | 1142 | 1159 | GPPGSAGAPGKDGLNGLP | GPP[113]GSAGAPGKDGLNGLP[113] | GP(0.0394)P(0.9730)GSAGAP(0.0136)GKDGLNGLP(0.9740) | 35.86 | 33.69 | 1.000 |
| 1594.765 | COL1A1 | 1142 | 1159 | GPPGSAGAPGKDGLNGLP | GPP[113]GSAGAPGKDGLN[115]GLP[113] | GP(0.0138)P(0.9843)GSAGAP(0.0178)GKDGLNGLP(0.9841) | 45.88 | 43.38 | 1.000 |
| 1596.707 | COL1A1 | 650 | 666 | GPPGEAGKPGEQGVPGD | GP[113]PGEAGKP[113]GEQGVP[113]GD | GP(0.7216)P(0.7216)GEAGKP(0.7784)GEQGVP(0.7784)GD | 24.07 | 24.07 | 1.000 |
| 1600.761 | COL1A1 | 401 | 418 | GAPGIAGAPGFPGARGPS | GAP[113]GIAGAP[113]GFP[113]GARGP[113]S | GAP(1.0000)GIAGAP(1.0000)GFP(1.0000)GARGP(1.0000)S | 24.07 | 12.84 | 0.996 |
| 1609.768 | COL1A1 | 1142 | 1159 | GPPGSAGAPGKDGLNGLP | GPP[113]GSAGAP[113]GKDGLNGLP[113] | GP(0.0371)P(0.9876)GSAGAP(0.9876)GKDGLNGLP(0.9876) | 19.85 | 17.96 | 0.999 |
| 1609.781 | COL1A1 | 1055 | 1072 | GPVGPAGKSGDRGETGPA | GPVGPAGKSGDRGETGPA | | 20.09 | 12.66 | 1.000 |
| 1609.791 | COL1A1 | 341 | 358 | GPPGFPGAVGAKGEAGPQ | GPP[113]GFPGAVGAKGEAGPQ | GP(0.0069)P(0.9840)GFP(0.0039)GAVGAKGEAGP(0.0052)Q | 22.34 | 20.47 | 0.980 |
| 1610.762 | COL1A1 | 1142 | 1159 | GPPGSAGAPGKDGLNGLP | GPP[113]GSAGAP[113]GKDGLN[115]GLP[113] | GP(0.0524)P(0.9825)GSAGAP(0.9826)GKDGLNGLP(0.9826) | 20.05 | 20.05 | 0.995 |
| 1612.672 | COL1A1 | 197 | 213 | GPQGFQGPPGEPGEPGA | GPQ[129]GFQ[129]GPPGEP[113]GEP[113]GA | GP(0.0109)QGFQGP(0.0092)P(0.0097)GEP(0.9852)GEP(0.9851)GA | 32.35 | 23.63 | 0.997 |
| 1615.750 | COL1A1 | 1091 | 1106 | GPRGDKGETGEQGDRG | GPRGDKGETGEQGDRG | | 25.67 | 8.26 | 1.000 |
| 1622.820 | COL1A1 | 899 | 916 | GPAGKEGGKGPRGETGPA | GPAGKEGGKGPRGETGPA | | 19.44 | 10.68 | 1.000 |
| 1625.783 | COL1A1 | 341 | 358 | GPPGFPGAVGAKGEAGPQ | GPP[113]GFP[113]GAVGAKGEAGPQ | GP(0.0194)P(0.9786)GFP(0.9785)GAVGAKGEAGP(0.0235)Q | 36.40 | 36.40 | 1.000 |
| 1628.661 | COL1A1 | 197 | 213 | GPQGFQGPPGEPGEPGA | GPQ[129]GFQ[129]GPP[113]GEP[113]GEP[113]GA | GP(0.0163)QGFQGP(0.0214)P(0.9874)GEP(0.9875)GEP(0.9875)GA | 22.90 | 21.01 | 0.992 |
| 1631.729 | COL1A1 | 1091 | 1106 | GPRGDKGETGEQGDRG | GP[113]RGDKGETGEQGDRG | GP(1.0000)RGDKGETGEQGDRG | 29.89 | 10.39 | 1.000 |
| 1634.771 | COL1A1 | 1001 | 1018 | GPPGLAGPPGESGREGAP | GPPGLAGPP[113]GESGREGAP[113] | GP(0.0070)P(0.0070)GLAGP(0.0087)P(0.9890)GESGREGAP(0.9883) | 23.92 | 19.92 | 0.995 |
| 1641.783 | COL1A1 | 341 | 358 | GPPGFPGAVGAKGEAGPQ | GPP[113]GFP[113]GAVGAKGEAGP[113]Q | GP(0.0538)P(0.9824)GFP(0.9825)GAVGAKGEAGP(0.9814)Q | 18.60 | 17.12 | 0.998 |
| 1643.695 | COL1A1 | 201 | 217 | FQGPPGEPGEPGASGPM | FQGPPGEP[113]GEP[113]GASGPM | FQGP(0.0059)P(0.0060)GEP(0.9878)GEP(0.9877)GASGP(0.0063)M(0.0063) | 32.02 | 22.99 | 1.000 |
| 1650.845 | COL1A1 | 584 | 601 | GPKGAAGEPGKAGERGVP | GPKGAAGEP[113]GKAGERGVP | GP(0.0091)KGAAGEP(0.9825)GKAGERGVP(0.0084) | 26.56 | 14.75 | 1.000 |
| 1651.792 | COL1A1 | 653 | 670 | GEAGKPGEQGVPGDLGAP | GEAGKPGEQGVPGDLGAP[113] | GEAGKP(0.0058)GEQGVP(0.0056)GDLGAP(0.9886) | 22.92 | 15.81 | 0.954 |
| 1656.811 | COL1A1 | 440 | 457 | GSKGDTGAKGEPGPVGVQ | GSKGDTGAKGEP[113]GPVGVQ | GSKGDTGAKGEP(0.9890)GP(0.0110)VGVQ | 24.09 | 18.53 | 1.000 |
| 1659.688 | COL1A1 | 201 | 217 | FQGPPGEPGEPGASGPM | FQGPPGEP[113]GEP[113]GASGP[113]M | FQGP(0.0101)P(0.0101)GEP(0.8241)GEP(0.8226)GASGP(0.6666)M(0.6666) | 19.97 | 19.97 | 0.982 |
| 1660.675 | COL1A1 | 201 | 217 | FQGPPGEPGEPGASGPM | FQ[129]GPPGEP[113]GEP[113]GASGP[113]M | FQGP(0.0095)P(0.0095)GEP(0.8292)GEP(0.8292)GASGP(0.6613)M(0.6613) | 31.43 | 31.43 | 1.000 |
| 1666.835 | COL1A1 | 584 | 601 | GPKGAAGEPGKAGERGVP | GPKGAAGEP[113]GKAGERGVP[113] | GP(0.0359)KGAAGEP(0.9820)GKAGERGVP(0.9821) | 27.40 | 17.07 | 1.000 |
| 1667.777 | COL1A1 | 653 | 670 | GEAGKPGEQGVPGDLGAP | GEAGKPGEQGVP[113]GDLGAP[113] | GEAGKP(0.0368)GEQGVP(0.9816)GDLGAP(0.9816) | 25.02 | 14.59 | 0.986 |
| 1668.762 | COL1A1 | 653 | 670 | GEAGKPGEQGVPGDLGAP | GEAGKP[113]GEQ[129]GVPGDLGAP[113] | GEAGKP(0.9800)GEQGVP(0.0400)GDLGAP(0.9800) | 23.61 | 17.91 | 1.000 |
| 1668.848 | COL1A1 | 749 | 766 | GPKGADGSPGKDGVRGLT | GPKGADGSPGKDGVRGLT | | 18.65 | 12.16 | 1.000 |
| 1669.779 | COL1A1 | 266 | 283 | GHRGFSGLDGAKGDAGPA | GHRGFSGLDGAKGDAGPA | | 19.71 | 11.46 | 1.000 |
| 1675.671 | COL1A1 | 201 | 217 | FQGPPGEPGEPGASGPM | FQGPP[113]GEP[113]GEP[113]GASGPM[147] | FQGP(0.0882)P(0.8731)GEP(0.8738)GEP(0.8738)GASGP(0.4279)M(0.8632) | 34.16 | 34.16 | 0.997 |
| 1676.677 | COL1A1 | 201 | 217 | FQGPPGEPGEPGASGPM | FQ[129]GP[113]P[113]GEP[113]GEP[113]GASGPM | FQGP(0.6414)P(0.6414)GEP(0.7172)GEP(0.7171)GASGP(0.6414)M(0.6414) | 20.28 | 20.28 | 0.999 |
| 1676.860 | COL1A1 | 1200 | 1214 | FLPQPPQEKAHDGGR | FLPQPPQEKAHDGGR | | 19.49 | 10.34 | 1.000 |
| 1681.799 | COL1A1 | 518 | 535 | GPKGSPGEAGRPGEAGLP | GPKGSP[113]GEAGRP[113]GEAGLP[113] | GP(0.0377)KGSP(0.9874)GEAGRP(0.9874)GEAGLP(0.9874) | 21.26 | 15.67 | 0.983 |
| 1682.828 | COL1A1 | 584 | 601 | GPKGAAGEPGKAGERGVP | GP[113]KGAAGEP[113]GKAGERGVP[113] | GP(1.0000)KGAAGEP(1.0000)GKAGERGVP(1.0000) | 29.71 | 16.62 | 1.000 |
| 1683.767 | COL1A1 | 653 | 670 | GEAGKPGEQGVPGDLGAP | GEAGKP[113]GEQGVP[113]GDLGAP[113] | GEAGKP(1.0000)GEQGVP(1.0000)GDLGAP(1.0000) | 25.92 | 14.15 | 0.992 |
| 1684.759 | COL1A1 | 653 | 670 | GEAGKPGEQGVPGDLGAP | GEAGKP[113]GEQ[129]GVP[113]GDLGAP[113] | GEAGKP(1.0000)GEQGVP(1.0000)GDLGAP(1.0000) | 28.13 | 14.89 | 1.000 |
| 1684.846 | COL1A1 | 749 | 766 | GPKGADGSPGKDGVRGLT | GPKGADGSP[113]GKDGVRGLT | GP(0.0297)KGADGSP(0.9703)GKDGVRGLT | 25.71 | 12.94 | 1.000 |
| 1697.726 | COL1A1 | 197 | 214 | GPQGFQGPPGEPGEPGAS | GPQGFQGPPGEP[113]GEP[113]GAS | GP(0.0100)QGFQGP(0.0071)P(0.0071)GEP(0.9879)GEP(0.9878)GAS | 22.26 | 14.40 | 0.989 |
| 1698.708 | COL1A1 | 197 | 214 | GPQGFQGPPGEPGEPGAS | GPQGFQ[129]GPPGEP[113]GEP[113]GAS | GP(0.0111)QGFQGP(0.0078)P(0.0080)GEP(0.9865)GEP(0.9865)GAS | 31.49 | 25.30 | 0.999 |
| 1699.702 | COL1A1 | 197 | 214 | GPQGFQGPPGEPGEPGAS | GPQ[129]GFQ[129]GPPGEP[113]GEP[113]GAS | GP(0.0089)QGFQGP(0.0071)P(0.0071)GEP(0.9884)GEP(0.9885)GAS | 25.75 | 17.05 | 1.000 |
| 1699.799 | COL1A1 | 722 | 739 | GAPGLQGMPGERGAAGLP | GAP[113]GLQGM[147]P[113]GERGAAGLP[113] | GAP(1.0000)GLQGM(1.0000)P(1.0000)GERGAAGLP(1.0000) | 24.04 | 13.88 | 0.999 |
| 1700.790 | COL1A1 | 722 | 739 | GAPGLQGMPGERGAAGLP | GAP[113]GLQ[129]GM[147]P[113]GERGAAGLP[113] | GAP(1.0000)GLQGM(1.0000)P(1.0000)GERGAAGLP(1.0000) | 22.51 | 11.38 | 1.000 |
| 1700.797 | COL1A1 | 1025 | 1042 | GRDGSPGAKGDRGETGPA | GRDGSP[113]GAKGDRGETGPA | GRDGSP(0.9868)GAKGDRGETGP(0.0132)A | 19.44 | 7.74 | 1.000 |
| 1713.734 | COL1A1 | 197 | 214 | GPQGFQGPPGEPGEPGAS | GPQGFQGPP[113]GEP[113]GEP[113]GAS | GP(0.0275)QGFQGP(0.4284)P(0.8363)GEP(0.8539)GEP(0.8539)GAS | 22.11 | 20.27 | 0.997 |
| 1716.711 | COL1A1 | 200 | 217 | GFQGPPGEPGEPGASGPM | GFQGPPGEP[113]GEP[113]GASGP[113]M | GFQGP(0.0091)P(0.0091)GEP(0.8275)GEP(0.8275)GASGP(0.6634)M(0.6634) | 36.31 | 36.31 | 1.000 |
| 1719.833 | COL1A1 | 908 | 925 | GPRGETGPAGRPGEVGPP | GPRGETGPAGRP[113]GEVGPP[113] | GP(0.0142)RGETGP(0.0190)AGRP(0.9732)GEVGP(0.0204)P(0.9731) | 21.63 | 21.63 | 0.969 |
| 1725.770 | COL1A1 | 653 | 671 | GEAGKPGEQGVPGDLGAPG | GEAGKP[113]GEQ[129]GVPGDLGAP[113]G | GEAGKP(0.9801)GEQGVP(0.0400)GDLGAP(0.9799)G | 22.02 | 11.63 | 1.000 |

### Supplemental Table 3. Col1a1 peptide sequences from normal breast by LC-MS/MS.

| | | | | | | | | | |
|---|---|---|---|---|---|---|---|---|---|
| 1731.710 | COL1A1 | 284 | 301 | GPKGEPGSPGENGAPGQM | GPKGEP[113]GSP[113]GEN[115]GAP[113]GQM[147] | GP(0.0635)KGEP(0.9841)GSP(0.9841)GENGAP(0.9841)GQM(0.9841) | 19.33 | 15.49 | 0.947 |
| 1738.846 | COL1A1 | 461 | 478 | GPAGEEGKRGARGEPGPT | GPAGEEGKRGARGEP[113]GPT | GP(0.0061)AGEEGKRGARGEP(0.9885)GP(0.0054)T | 18.10 | 14.02 | 0.998 |
| 1742.834 | COL1A1 | 1202 | 1216 | PQPPQEKAHDGGRYY | PQPPQEKAHDGGRYY | | 20.23 | 11.26 | 1.000 |
| 1762.846 | COL1A1 | 359 | 376 | GPRGSEGPQGVRGEPGPP | GPRGSEGPQGVRGEP[113]GPP[113] | GP(0.0070)RGSEGP(0.0070)QGVRGEP(0.9890)GP(0.0080)P(0.9890) | 18.66 | 18.66 | 1.000 |
| 1768.931 | COL1A1 | 960 | 976 | VVGLPGQRGERGFPGLP | VVGLP[113]GQ[129]RGERGFPGLP[113] | VVGLP(0.9807)GQRGERGFP(0.0385)GLP(0.9809) | 18.91 | 15.64 | 0.987 |
| 1774.781 | COL1A1 | 629 | 646 | GERGEQGPAGSPGFQGLP | GERGEQ[129]GPAGSP[113]GFQ[129]GLP[113] | GERGEQGP(0.0295)AGSP(0.9853)GFQGLP(0.9853) | 21.35 | 16.61 | 1.000 |
| 1788.837 | COL1A1 | 1190 | 1206 | GPPSAGFDFSFLPQPPQ | GPPSAGFDFSFLPQPPQ | | 21.37 | 11.43 | 0.942 |
| 1802.857 | COL1A1 | 602 | 622 | GPPGAVGPAGKDGEAGAQGPP | GPPGAVGPAGKDGEAGAQGPP[113] | GP(0.0028)P(0.0028)GAVGP(0.0025)AGKDGEAGAQGP(0.0038)P(0.9880) | 20.22 | 20.22 | 0.990 |
| 1818.847 | COL1A1 | 602 | 622 | GPPGAVGPAGKDGEAGAQGPP | GP[113]P[113]GAVGPAGKDGEAGAQGPP | GP(0.4987)P(0.4987)GAVGP(0.0052)AGKDGEAGAQGP(0.4987)P(0.4987) | 18.11 | 18.11 | 0.979 |
| 1819.945 | COL1A1 | 851 | 871 | GPIGNVGAPGAKGARGSAGPP | GPIGNVGAP[113]GAKGARGSAGP[113]P | GP(0.0109)IGNVGAP(0.7639)GAKGARGSAGP(0.6126)P(0.6126) | 19.57 | 19.57 | 0.981 |
| 1827.859 | COL1A1 | 398 | 418 | GANGAPGIAGAPGFPGARGPS | GAN[115]GAP[113]GIAGAP[113]GFP[113]GARGPS | GANGAP(0.9088)GIAGAP(0.9087)GFP(0.9072)GARGP(0.2753)S | 24.55 | 24.55 | 0.999 |
| 1839.807 | COL1A1 | 695 | 715 | GPRGANGAPGNDGAKGDAGAP | GPRGAN[115]GAP[113]GNDGAKGDAGAP[113] | GP(0.0398)RGANGAP(0.9801)GNDGAKGDAGAP(0.9801) | 27.31 | 24.09 | 1.000 |
| 1840.783 | COL1A1 | 695 | 715 | GPRGANGAPGNDGAKGDAGAP | GPRGAN[115]GAP[113]GN[115]DGAKGDAGAP[113] | GP(0.0288)RGANGAP(0.9856)GNDGAKGDAGAP(0.9856) | 20.20 | 12.38 | 0.998 |
| 1841.893 | COL1A1 | 395 | 415 | GAKGANGAPGIAGAPGFPGAR | GAKGANGAP[113]GIAGAP[113]GFP[113]GAR | GAKGANGAP(1.0000)GIAGAP(1.0000)GFP(1.0000)GAR | 23.65 | 10.58 | 1.000 |
| 1842.889 | COL1A1 | 395 | 415 | GAKGANGAPGIAGAPGFPGAR | GAKGAN[115]GAP[113]GIAGAP[113]GFP[113]GAR | GAKGANGAP(1.0000)GIAGAP(1.0000)GFP(1.0000)GAR | 33.31 | 20.48 | 1.000 |
| 1876.930 | COL1A1 | 1145 | 1165 | GSAGAPGKDGLNGLPGPIGPP | GSAGAPGKDGLNGLP[113]GPIGP[113]P[113] | GSAGAP(0.0169)GKDGLNGLP(0.9668)GP(0.0856)IGP(0.9654)P(0.9654) | 30.16 | 24.10 | 0.982 |
| 1877.879 | COL1A1 | 1145 | 1165 | GSAGAPGKDGLNGLPGPIGPP | GSAGAPGKDGLN[115]GLP[113]GPIGP[113]P[113] | GSAGAP(0.0425)GKDGLNGLP(0.9628)GP(0.0676)IGP(0.9638)P(0.9633) | 18.97 | 17.12 | 0.998 |
| 1882.886 | COL1A1 | 536 | 556 | GAKGLTGSPGSPGPDGKTGPP | GAKGLTGSP[113]GSP[113]GPDGKTGPP[113] | GAKGLTGSP(0.9824)GSP(0.9824)GP(0.0228)DGKTGP(0.0300)P(0.9822) | 28.03 | 28.03 | 1.000 |
| 1899.995 | COL1A1 | 956 | 973 | GQRGVVGLPGQRGERGFP | GQRGVVGLP[113]GQ[129]RGERGFP[113] | GQRGVVGLP(1.0000)GQRGERGFP(1.0000) | 19.44 | 12.14 | 1.000 |
| 1917.847 | COL1A1 | 431 | 451 | GNSGEPGAPGSKGDTGAKGEP | GNSGEP[113]GAP[113]GSKGDTGAKGEP[113] | GNSGEP(1.0000)GAP(1.0000)GSKGDTGAKGEP(1.0000) | 20.62 | 11.29 | 0.998 |
| 1918.018 | COL1A1 | 944 | 964 | GAPGTPGPQGIAGQRGVVGLP | GAP[113]GTPGPQGIAGQRGVVGLP[113] | GAP(0.9888)GTP(0.0106)GP(0.0119)QGIAGQRGVVGLP(0.9888) | 22.83 | 18.25 | 1.000 |
| 1918.903 | COL1A1 | 650 | 670 | GPPGEAGKPGEQGVPGDLGAP | GP[113]PGEAGKPGEQGVPGDLGAP[113] | GP(0.5882)P(0.5882)GEAGKP(0.0055)GEQGVP(0.0084)GDLGAP(0.8097) | 24.93 | 24.93 | 0.981 |
| 1921.887 | COL1A1 | 977 | 997 | GPSGEPGKQGPSGASGERGPP | GPSGEPGKQGPSGASGERGPP[113] | GP(0.0038)SGEP(0.0065)GKQGP(0.0049)SGASGERGP(0.0068)P(0.9780) | 25.49 | 25.49 | 1.000 |
| 1923.918 | COL1A1 | 440 | 460 | GSKGDTGAKGEPGPVGVQGPP | GSKGDTGAKGEP[113]GPVGVQGPP[113] | GSKGDTGAKGEP(0.9681)GP(0.0376)VGVQGP(0.0258)P(0.9686) | 21.85 | 21.85 | 0.995 |
| 1934.881 | COL1A1 | 650 | 670 | GPPGEAGKPGEQGVPGDLGAP | GPP[113]GEAGKP[113]GEQGVPGDLGAP[113] | GP(0.0200)P(0.9874)GEAGKP(0.9874)GEQGVP(0.0177)GDLGAP(0.9875) | 50.97 | 50.97 | 1.000 |
| 1935.849 | COL1A1 | 650 | 670 | GPPGEAGKPGEQGVPGDLGAP | GPP[113]GEAGKP[113]GEQ[129]GVP[113]GDLGAP | GP(0.1042)P(0.8323)GEAGKP(0.8342)GEQGVP(0.8342)GDLGAP(0.3950) | 19.84 | 19.84 | 0.999 |
| 1937.877 | COL1A1 | 977 | 997 | GPSGEPGKQGPSGASGERGPP | GPSGEP[113]GKQGPSGASGERGP[113]P | GP(0.0063)SGEP(0.8035)GKQGP(0.0123)SGASGERGP(0.5890)P(0.5890) | 19.99 | 19.99 | 1.000 |
| 1939.916 | COL1A1 | 440 | 460 | GSKGDTGAKGEPGPVGVQGPP | GSKGDTGAKGEP[113]GP[113]VGVQGP[113]P | GSKGDTGAKGEP(0.7798)GP(0.7798)VGVQGP(0.7202)P(0.7202) | 22.44 | 22.44 | 0.977 |
| 1942.896 | COL1A1 | 482 | 502 | GPPGERGGPGSRGFPGADGVA | GPP[113]GERGGP[113]GSRGFP[113]GADGVA | GP(0.0311)P(0.9896)GERGGP(0.9896)GSRGFP(0.9896)GADGVA | 22.50 | 20.31 | 0.996 |
| 1950.856 | COL1A1 | 650 | 670 | GPPGEAGKPGEQGVPGDLGAP | GPP[113]GEAGKP[113]GEQGVP[113]GDLGAP[113] | GP(0.0510)P(0.9872)GEAGKP(0.9873)GEQGVP(0.9872)GDLGAP(0.9872) | 47.82 | 47.82 | 1.000 |
| 1951.882 | COL1A1 | 650 | 670 | GPPGEAGKPGEQGVPGDLGAP | GP[113]PGEAGKP[113]GEQ[129]GVP[113]GDLGAP[113] | GP(0.7706)P(0.7706)GEAGKP(0.8196)GEQGVP(0.8196)GDLGAP(0.8196) | 18.97 | 18.97 | 0.976 |
| 1982.844 | COL1A1 | 197 | 217 | GPQGFQGPPGEPGEPGASGPM | GPQGFQGPPGEP[113]GEP[113]GASGPM | GP(0.0045)QGFQGP(0.0044)P(0.0042)GEP(0.9872)GEP(0.9867)GASGP(0.0065)M(0.0065) | 30.34 | 21.92 | 0.998 |
| 1998.847 | COL1A1 | 197 | 217 | GPQGFQGPPGEPGEPGASGPM | GPQGFQGPPGEP[113]GEP[113]GASGPM[147] | GP(0.0080)QGFQGP(0.0078)P(0.0078)GEP(0.9876)GEP(0.9876)GASGP(0.0140)M(0.9872) | 63.88 | 63.88 | 1.000 |
| 1999.833 | COL1A1 | 197 | 217 | GPQGFQGPPGEPGEPGASGPM | GPQGFQ[129]GPPGEP[113]GEP[113]GASGP[113]M | GP(0.0122)QGFQGP(0.0075)P(0.0075)GEP(0.8470)GEP(0.8468)GASGP(0.6395)M(0.6395) | 18.10 | 18.10 | 0.965 |
| 1999.901 | COL1A1 | 626 | 646 | GPAGERGEQGPAGSPGFQGLP | GPAGERGEQ[129]GPAGSP[113]GFQ[129]GLP[113] | GP(0.0277)AGERGEQGP(0.0304)AGSP(0.9707)GFQGLP(0.9712) | 18.89 | 14.16 | 1.000 |
| 2000.802 | COL1A1 | 197 | 217 | GPQGFQGPPGEPGEPGASGPM | GPQ[129]GFQ[129]GPPGEP[113]GEP[113]GASGP[113]M | GP(0.0085)QGFQGP(0.0092)P(0.0093)GEP(0.8443)GEP(0.8444)GASGP(0.6421)M(0.6421) | 24.36 | 24.36 | 1.000 |
| 2002.966 | COL1A1 | 1200 | 1216 | FLPQPPQEKAHDGGRYY | FLPQPPQEKAHDGGRYY | | 20.01 | 11.18 | 1.000 |
| 2003.967 | COL1A1 | 1200 | 1216 | FLPQPPQEKAHDGGRYY | FLPQPPQ[129]EKAHDGGRYY | | 20.94 | 20.94 | 1.000 |
| 2014.838 | COL1A1 | 197 | 217 | GPQGFQGPPGEPGEPGASGPM | GPQGFQGPP[113]GEP[113]GEP[113]GASGP[113]M | GP(0.0121)QGFQGP(0.0198)P(0.8489)GEP(0.8491)GEP(0.8491)GASGP(0.7105)M(0.7105) | 27.04 | 27.04 | 0.952 |
| 2015.821 | COL1A1 | 197 | 217 | GPQGFQGPPGEPGEPGASGPM | GPQGFQ[129]GP[113]PGEP[113]GEP[113]GASGP[113]M | GP(0.0139)QGFQGP(0.8432)P(0.0291)GEP(0.8436)GEP(0.8435)GASGP(0.7133)M(0.7133) | 40.35 | 40.35 | 1.000 |
| 2030.839 | COL1A1 | 197 | 217 | GPQGFQGPPGEPGEPGASGPM | GPQGFQGP[113]P[113]GEP[113]GEP[113]GASGP[113]M | GP(0.0335)QGFQGP(0.8534)P(0.8534)GEP(0.8534)GEP(0.8535)GASGP(0.7763)M(0.7763) | 23.89 | 18.28 | 0.999 |
| 2041.877 | COL1A1 | 284 | 304 | GPKGEPGSPGENGAPGQMGPR | GPKGEP[113]GSP[113]GEN[115]GAP[113]GQM[147]GPR | GP(0.0302)KGEP(0.8318)GSP(0.8319)GENGAP(0.8181)GQM(0.7440)GP(0.7440)R | 21.40 | 21.40 | 0.989 |
| 2051.915 | COL1A1 | 386 | 409 | GNPGADGQPGAKGANGAPGIAGAP | GNP[113]GADGQ[129]PGAKGAN[115]GAP[113]GIAGAP[113] | GNP(0.9896)GADGQP(0.0312)GAKGANGAP(0.9895)GIAGAP(0.9896) | 19.05 | 15.14 | 0.941 |
| 2079.956 | COL1A1 | 227 | 247 | GKNGDDGEAGKPGRPGERGPP | GKNGDDGEAGKPGRP[113]GERGP[113]P | GKNGDDGEAGKP(0.0108)GRP(0.7642)GERGP(0.6125)P(0.6125) | 23.96 | 23.96 | 1.000 |
| 2080.954 | COL1A1 | 227 | 247 | GKNGDDGEAGKPGRPGERGPP | GKN[115]GDDGEAGKPGRP[113]GERGP[113]P | GKNGDDGEAGKP(0.0104)GRP(0.7649)GERGP(0.6123)P(0.6123) | 29.97 | 29.97 | 1.000 |
| 2083.012 | COL1A1 | 395 | 418 | GAKGANGAPGIAGAPGFPGARGPS | GAKGANGAP[113]GIAGAP[113]GFP[113]GARGPS | GAKGANGAP(0.9366)GIAGAP(0.9366)GFP(0.9357)GARGP(0.1911)S | 29.19 | 29.19 | 1.000 |
| 2083.966 | COL1A1 | 395 | 418 | GAKGANGAPGIAGAPGFPGARGPS | GAKGAN[115]GAP[113]GIAGAP[113]GFP[113]GARGPS | GAKGANGAP(0.9397)GIAGAP(0.9393)GFP(0.9389)GARGP(0.1821)S | 23.64 | 22.20 | 1.000 |
| 2084.098 | COL1A1 | 954 | 973 | IAGQRGVVGLPGQRGERGFP | IAGQRGVVGLP[113]GQ[129]RGERGFP[113] | IAGQRGVVGLP(1.0000)GQRGERGFP(1.0000) | 19.22 | 12.28 | 0.996 |
| 2085.087 | COL1A1 | 954 | 973 | IAGQRGVVGLPGQRGERGFP | IAGQ[129]RGVVGLP[113]GQ[129]RGERGFP[113] | IAGQRGVVGLP(1.0000)GQRGERGFP(1.0000) | 23.31 | 16.75 | 1.000 |
| 2093.917 | COL1A1 | 818 | 841 | GADGQPGAKGEPGDAGAKGDAGPP | GADGQ[129]PGAKGEPGDAGAKGDAGP[113]P | GADGQP(0.0043)GAKGEP(0.0032)GDAGAKGDAGP(0.4963)P(0.4963) | 31.30 | 31.30 | 1.000 |
| 2096.936 | COL1A1 | 227 | 247 | GKNGDDGEAGKPGRPGERGPP | GKN[115]GDDGEAGKP[113]GRP[113]GERGP[113]P | GKNGDDGEAGKP(0.7784)GRP(0.7783)GERGP(0.7216)P(0.7216) | 27.24 | 27.24 | 1.000 |
| 2099.972 | COL1A1 | 395 | 418 | GAKGANGAPGIAGAPGFPGARGPS | GAKGAN[115]GAP[113]GIAGAP[113]GFP[113]GARGP[113]S | GAKGANGAP(1.0000)GIAGAP(1.0000)GFP(1.0000)GARGP(1.0000)S | 22.51 | 12.50 | 1.000 |
| 2101.964 | COL1A1 | 162 | 181 | QLSYGYDEKSTGGISVPGPM | QLSYGYDEKSTGGISVPGP[113]M | QLSYGYDEKSTGGISVP(0.0048)GP(0.4976)M(0.4976) | 19.94 | 19.94 | 1.000 |
| 2108.927 | COL1A1 | 818 | 841 | GADGQPGAKGEPGDAGAKGDAGPP | GADGQPGAKGEP[113]GDAGAKGDAGPP[113] | GADGQP(0.0169)GAKGEP(0.9851)GDAGAKGDAGP(0.0127)P(0.9853) | 19.76 | 19.76 | 1.000 |
| 2109.905 | COL1A1 | 818 | 841 | GADGQPGAKGEPGDAGAKGDAGPP | GADGQ[129]PGAKGEP[113]GDAGAKGDAGP[113]P | GADGQP(0.0102)GAKGEP(0.7643)GDAGAKGDAGP(0.6128)P(0.6128) | 23.21 | 23.21 | 1.000 |
| 2124.934 | COL1A1 | 818 | 841 | GADGQPGAKGEPGDAGAKGDAGPP | GADGQP[113]GAKGEP[113]GDAGAKGDAGP[113]P | GADGQP(0.7798)GAKGEP(0.7798)GDAGAKGDAGP(0.7202)P(0.7202) | 18.75 | 18.75 | 0.991 |
| 2125.892 | COL1A1 | 818 | 841 | GADGQPGAKGEPGDAGAKGDAGPP | GADGQ[129]P[113]GAKGEP[113]GDAGAKGDAGPP[113] | GADGQP(0.9869)GAKGEP(0.9869)GDAGAKGDAGP(0.0391)P(0.9871) | 33.28 | 33.28 | 1.000 |
| 2200.002 | COL1A1 | 428 | 451 | GPKGNSGEPGAPGSKGDTGAKGEP | GPKGNSGEP[113]GAP[113]GSKGDTGAKGEP[113] | GP(0.0338)KGNSGEP(0.9887)GAP(0.9887)GSKGDTGAKGEP(0.9887) | 20.20 | 14.65 | 0.987 |
| 2215.000 | COL1A1 | 1019 | 1042 | GAEGSPGRDGSPGAKGDRGETGPA | GAEGSP[113]GRDGSP[113]GAKGDRGETGPA | GAEGSP(0.9216)GRDGSP(0.9211)GAKGDRGETGP(0.1573)A | 19.42 | 11.41 | 0.985 |
| 2225.057 | COL1A1 | 482 | 505 | GPPGERGGPGSRGFPGADGVAGPK | GPP[113]GERGGP[113]GSRGFP[113]GADGVAGPK | GP(0.0284)P(0.9829)GERGGP(0.9829)GSRGFP(0.9829)GADGVAGP(0.0229)K | 21.39 | 17.66 | 0.988 |
| 2256.067 | COL1A1 | 743 | 766 | GDRGDAGPKGADGSPGKDGVRGLT | GDRGDAGPKGADGSP[113]GKDGVRGLT | GDRGDAGP(0.0124)KGADGSP(0.9876)GKDGVRGLT | 31.84 | 16.40 | 1.000 |
| 2264.126 | COL1A1 | 1091 | 1112 | GPRGDKGETGEQGDRGIKGHRG | GPRGDKGETGEQGDRGIKGHRG | | 24.44 | 12.66 | 1.000 |
| 2308.124 | COL1A1 | 740 | 764 | GPKGDRGDAGPKGADGSPGKDGVRG | GPKGDRGDAGPKGADGSPGKDGVRG | | 23.43 | 11.06 | 0.987 |
| 2324.113 | COL1A1 | 740 | 764 | GPKGDRGDAGPKGADGSPGKDGVRG | GPKGDRGDAGPKGADGSP[113]GKDGVRG | GP(0.0179)KGDRGDAGP(0.0137)KGADGSP(0.9684)GKDGVRG | 24.09 | 14.80 | 1.000 |
| 2333.112 | COL1A1 | 1049 | 1075 | GAPGAPGPVGPAGKSGDRGETGPAGPA | GAP[113]GAP[113]GPVGPAGKSGDRGETGP[113]AGPA | GAP(0.9585)GAP(0.9581)GP(0.0227)VGP(0.0306)AGKSGDRGETGP(0.9558)AGP(0.0743)A | 24.37 | 24.37 | 0.994 |
| 2347.086 | COL1A1 | 224 | 247 | GPPGKNGDDGEAGKPGRPGERGPP | GPP[113]GKNGDDGEAGKPGRP[113]GERGP[113]P | GP(0.0455)P(0.8114)GKNGDDGEAGKP(0.0158)GRP(0.8125)GERGP(0.6574)P(0.6574) | 27.59 | 27.59 | 1.000 |
| 2348.067 | COL1A1 | 224 | 247 | GPPGKNGDDGEAGKPGRPGERGPP | GPP[113]GKN[115]GDDGEAGKP[113]GRP[113]GERGPP | GP(0.0077)P(0.7118)GKNGDDGEAGKP(0.5701)GRP(0.5701)GERGP(0.5701)P(0.5701) | 20.61 | 20.61 | 1.000 |
| 2363.090 | COL1A1 | 224 | 247 | GPPGKNGDDGEAGKPGRPGERGPP | GPP[113]GKNGDDGEAGKP[113]GRPGERGP[113]P[113] | GP(0.0877)P(0.8190)GKNGDDGEAGKP(0.7274)GRP(0.7274)GERGP(0.8192)P(0.8192) | 21.07 | 21.07 | 1.000 |
| 2364.065 | COL1A1 | 224 | 247 | GPPGKNGDDGEAGKPGRPGERGPP | GPP[113]GKN[115]GDDGEAGKP[113]GRP[113]GERGPP[113] | GP(0.1320)P(0.8760)GKNGDDGEAGKP(0.8691)GRP(0.8691)GERGP(0.3767)P(0.8771) | 20.28 | 20.28 | 0.967 |
| 2412.158 | COL1A1 | 359 | 385 | GPRGSEGPQGVRGEPGPPGPAGAAGPA | GPRGSEGPQGVRGEP[113]GPP[113]GPAGAAGPA | GP(0.0071)RGSEGP(0.0055)QGVRGEP(0.9605)GP(0.0400)P(0.9744)GP(0.0079)AGAAGP(0.0046)A | 28.24 | 26.27 | 0.973 |
| 2522.254 | COL1A1 | 740 | 766 | GPKGDRGDAGPKGADGSPGKDGVRGLT | GPKGDRGDAGPKGADGSPGKDGVRGLT | | 27.39 | 15.23 | 1.000 |
| 2538.247 | COL1A1 | 740 | 766 | GPKGDRGDAGPKGADGSPGKDGVRGLT | GPKGDRGDAGPKGADGSP[113]GKDGVRGLT | GP(0.0078)KGDRGDAGP(0.0068)KGADGSP(0.9854)GKDGVRGLT | 22.99 | 12.18 | 1.000 |
| 2554.253 | COL1A1 | 740 | 766 | GPKGDRGDAGPKGADGSPGKDGVRGLT | GP[113]KGDRGDAGPKGADGSP[113]GKDGVRGLT | GP(0.9743)KGDRGDAGP(0.0512)KGADGSP(0.9745)GKDGVRGLT | 31.99 | 23.37 | 1.000 |
| 2653.235 | COL1A1 | 386 | 415 | GNPGADGQPGAKGANGAPGIAGAPGFP[illegible] | GNP[113]GADGQ[129]PGAKGANGAP[113]GIAGAP[113]GFP[113]G[illegible] | GNP(0.9650)GADGQP(0.1425)GAKGANGAP(0.9628)GIAGAP(0.9648)GFP(0.9648)GAR | 18.76 | 18.71 | 0.993 |
| 2805.419 | COL1A1 | 740 | 769 | GPKGDRGDAGPKGADGSPGKDGVRGLT[illegible] | GPKGDRGDAGPKGADGSP[113]GKDGVRGLTGPI | GP(0.0083)KGDRGDAGP(0.0035)KGADGSP(0.9824)GKDGVRGLTGP(0.0058)I | 33.64 | 24.58 | 1.000 |

### Supplemental Table 3. Col1a2 peptide sequences from normal breast by LC-MS/MS.

| M+H | Gene | Domain Start | Domain End | Peptide | Modified Peptide [113] means HYP | Sequence with HYP probability PM:15.9949 | Hyperscore | Nextscore | Peptide Prophet Probability |
|---|---|---|---|---|---|---|---|---|---|
| 659.334 | COL1A2 | 629 | 636 | AVGTAGPS | AVGTAGPS | | 18.94 | 15.02 | 0.943 |
| 711.293 | COL1A2 | 1102 | 1109 | GVSGGGYD | GVSGGGYD | | 19.38 | 9.84 | 1.000 |
| 730.405 | COL1A2 | 625 | 633 | GVVGAVGTA | GVVGAVGTA | | 19.95 | 11.34 | 0.984 |
| 737.417 | COL1A2 | 896 | 903 | PLGIAGPP | PLGIAGPP[113] | P(0.0062)LGIAGP(0.0061)P(0.9877) | 19.79 | 17.86 | 0.969 |
| 741.375 | COL1A2 | 833 | 840 | EVGAVGPP | EVGAVGP[113]P | EVGAVGP(0.5000)P(0.5000) | 18.52 | 18.52 | 0.938 |
| 755.392 | COL1A2 | 344 | 351 | LVGEPGPA | LVGEP[113]GPA | LVGEP(0.9888)GP(0.0112)A | 19.30 | 15.45 | 0.945 |
| 755.402 | COL1A2 | 952 | 960 | GNIGPVGAA | GNIGPVGAA | | 19.71 | 12.92 | 0.975 |
| 756.387 | COL1A2 | 952 | 960 | GNIGPVGAA | GN[115]IGPVGAA | | 20.49 | 14.97 | 0.945 |
| 772.383 | COL1A2 | 886 | 894 | GVAGAVGEP | GVAGAVGEP[113] | GVAGAVGEP(1.0000) | 20.89 | 16.59 | 0.934 |
| 774.357 | COL1A2 | 910 | 918 | GAVGSPGVN | GAVGSP[113]GVN[115] | GAVGSP(1.0000)GVN | 21.65 | 12.55 | 0.968 |
| 784.382 | COL1A2 | 799 | 807 | GPSGISGPP | GPSGISGPP[113] | GP(0.0052)SGISGP(0.0056)P(0.9892) | 21.72 | 19.90 | 0.956 |
| 784.396 | COL1A2 | 86 | 94 | VGLGPGPMG | VGLGPGPMG | | 22.69 | 12.05 | 1.000 |
| 785.374 | COL1A2 | 454 | 462 | GSPGNIGPA | GSP[113]GNIGPA | GSP(0.9891)GNIGP(0.0109)A | 18.68 | 11.15 | 0.977 |
| 785.440 | COL1A2 | 1028 | 1035 | LQGLPGIA | LQ[129]GLP[113]GIA | LQGLP(1.0000)GIA | 18.68 | 14.79 | 0.954 |
| 787.369 | COL1A2 | 721 | 729 | GPNGFAGPA | GPNGFAGPA | | 19.58 | 10.35 | 0.996 |
| 788.354 | COL1A2 | 721 | 729 | GPNGFAGPA | GPN[115]GFAGPA | | 21.30 | 11.87 | 0.989 |
| 794.439 | COL1A2 | 895 | 903 | GPLGIAGPP | GPLGIAGPP[113] | GP(0.0053)LGIAGP(0.0059)P(0.9889) | 19.64 | 17.71 | 0.963 |
| 800.394 | COL1A2 | 86 | 94 | VGLGPGPMG | VGLGPGPM[147]G | VGLGP(0.0057)GP(0.0157)M(0.9786)G | 19.00 | 17.04 | 1.000 |
| 810.471 | COL1A2 | 868 | 876 | GAPGILGLP | GAPGILGLP[113] | GAP(0.0104)GILGLP(0.9896) | 21.77 | 16.98 | 0.938 |
| 812.412 | COL1A2 | 343 | 351 | GLVGEPGPA | GLVGEP[113]GPA | GLVGEP(0.9881)GP(0.0119)A | 21.16 | 15.86 | 0.944 |
| 815.419 | COL1A2 | 627 | 636 | VGAVGTAGPS | VGAVGTAGPS | | 20.53 | 12.02 | 0.988 |
| 824.439 | COL1A2 | 427 | 435 | GPAGVRGPN | GPAGVRGPN | | 19.00 | 9.83 | 0.997 |
| 825.437 | COL1A2 | 954 | 963 | IGPVGAAGAP | IGPVGAAGAP[113] | IGP(0.0104)VGAAGAP(0.9896) | 18.62 | 11.14 | 0.946 |
| 826.434 | COL1A2 | 267 | 276 | IGAVGNAGPA | IGAVGNAGPA | | 19.90 | 12.20 | 0.974 |
| 826.469 | COL1A2 | 868 | 876 | GAPGILGLP | GAP[113]GILGLP[113] | GAP(1.0000)GILGLP(1.0000) | 19.36 | 16.56 | 0.950 |
| 827.420 | COL1A2 | 267 | 276 | IGAVGNAGPA | IGAVGN[115]AGPA | | 21.65 | 12.50 | 0.997 |
| 827.426 | COL1A2 | 562 | 570 | GPAGEVGKP | GPAGEVGKP[113] | GP(0.0120)AGEVGKP(0.9880) | 19.23 | 12.26 | 0.941 |
| 830.379 | COL1A2 | 748 | 756 | GENGVVGPT | GEN[115]GVVGPT | | 19.66 | 12.97 | 0.958 |
| 840.355 | COL1A2 | 784 | 792 | GMTGFPGAA | GM[147]TGFP[113]GAA | GM(1.0000)TGFP(1.0000)GAA | 19.40 | 11.14 | 1.000 |
| 840.445 | COL1A2 | 466 | 474 | GPVGLPGID | GPVGLP[113]GID | GP(0.0104)VGLP(0.9896)GID | 18.43 | 16.78 | 0.946 |
| 841.420 | COL1A2 | 85 | 94 | GVGLGPGPMG | GVGLGPGPMG | | 20.66 | 9.57 | 1.000 |
| 843.453 | COL1A2 | 624 | 633 | PGVVGAVGTA | P[113]GVVGAVGTA | P(1.0000)GVVGAVGTA | 18.02 | 10.26 | 0.999 |
| 844.439 | COL1A2 | 283 | 291 | GEVGLPGLS | GEVGLP[113]GLS | GEVGLP(1.0000)GLS | 20.43 | 15.46 | 0.978 |
| 848.401 | COL1A2 | 1081 | 1089 | GPQGHQGPA | GPQGHQGPA | | 19.44 | 12.47 | 0.999 |
| 849.395 | COL1A2 | 409 | 417 | GADGRAGVM | GADGRAGVM[147] | GADGRAGVM(1.0000) | 19.21 | 10.56 | 0.991 |
| 857.415 | COL1A2 | 85 | 94 | GVGLGPGPMG | GVGLGPGPM[147]G | GVGLGP(0.0050)GP(0.0116)M(0.9834)G | 22.92 | 20.83 | 0.998 |
| 858.355 | COL1A2 | 1102 | 1110 | GVSGGGYDF | GVSGGGYDF | | 19.66 | 13.21 | 0.960 |
| 858.406 | COL1A2 | 109 | 117 | GPQGFQGPA | GPQGFQGPA | | 21.82 | 13.88 | 0.978 |
| 863.385 | COL1A2 | 1099 | 1108 | GPPGVSGGGY | GPP[113]GVSGGGY | GP(0.0120)P(0.9880)GVSGGGY | 21.88 | 19.46 | 0.977 |
| 868.446 | COL1A2 | 951 | 960 | PGNIGPVGAA | P[113]GNIGPVGAA | P(0.9832)GNIGP(0.0168)VGAA | 23.64 | 13.32 | 0.967 |
| 869.471 | COL1A2 | 463 | 471 | GKEGPVGLP | GKEGPVGLP[113] | GKEGP(0.0113)VGLP(0.9887) | 18.13 | 14.57 | 0.925 |
| 874.429 | COL1A2 | 166 | 174 | GFPGTPGLP | GFP[113]GTPGLP[113] | GFP(0.9901)GTP(0.0197)GLP(0.9901) | 23.10 | 13.81 | 0.998 |
| 883.453 | COL1A2 | 862 | 871 | GPQGLLGAPG | GPQ[129]GLLGAP[113]G | GP(0.0193)QGLLGAP(0.9807)G | 18.13 | 16.12 | 0.958 |
| 884.446 | COL1A2 | 562 | 571 | GPAGEVGKPG | GPAGEVGKP[113]G | GP(0.0104)AGEVGKP(0.9896)G | 19.07 | 12.03 | 0.987 |
| 884.494 | COL1A2 | 811 | 819 | GPAGKEGLR | GPAGKEGLR | | 18.16 | 10.57 | 0.997 |
| 885.445 | COL1A2 | 575 | 582 | LHGEFGLP | LHGEFGLP[113] | LHGEFGLP(1.0000) | 19.72 | 11.56 | 0.991 |
| 888.430 | COL1A2 | 88 | 96 | LGPGPMGLM | LGPGPMGLM[147] | LGP(0.0039)GP(0.0039)M(0.0040)GLM(0.9882) | 18.92 | 10.79 | 1.000 |
| 890.419 | COL1A2 | 949 | 957 | GYPGNIGPV | GYP[113]GN[115]IGPV | GYP(0.9817)GNIGP(0.0183)V | 18.53 | 14.98 | 0.999 |
| 890.427 | COL1A2 | 166 | 174 | GFPGTPGLP | GFP[113]GTP[113]GLP[113] | GFP(1.0000)GTP(1.0000)GLP(1.0000) | 18.28 | 14.60 | 0.954 |
| 897.466 | COL1A2 | 466 | 475 | GPVGLPGIDG | GPVGLP[113]GIDG | GP(0.0120)VGLP(0.9880)GIDG | 18.53 | 16.49 | 0.968 |
| 899.441 | COL1A2 | 831 | 840 | TGEVGAVGPP | TGEVGAVGPP[113] | TGEVGAVGP(0.0132)P(0.9868) | 22.07 | 20.05 | 0.973 |
| 901.442 | COL1A2 | 550 | 558 | GPPGFQGLP | GPP[113]GFQGLP[113] | GP(0.0213)P(0.9894)GFQGLP(0.9894) | 20.40 | 18.37 | 0.959 |
| 902.423 | COL1A2 | 550 | 558 | GPPGFQGLP | GPP[113]GFQ[129]GLP[113] | GP(0.0266)P(0.9867)GFQGLP(0.9867) | 19.87 | 17.87 | 0.965 |
| 904.426 | COL1A2 | 88 | 96 | LGPGPMGLM | LGPGPM[147]GLM[147] | LGP(0.0104)GP(0.0202)M(0.9845)GLM(0.9849) | 20.27 | 14.82 | 0.994 |
| 909.442 | COL1A2 | 1068 | 1077 | TGHPGTVGPA | TGHP[113]GTVGPA | TGHP(0.9891)GTVGP(0.0109)A | 18.12 | 12.12 | 0.973 |
| 911.515 | COL1A2 | 285 | 294 | VGLPGLSGPV | VGLP[113]GLSGPV | VGLP(0.9889)GLSGP(0.0111)V | 24.28 | 14.06 | 1.000 |
| 912.472 | COL1A2 | 311 | 321 | AAGLPGVAGAP | AAGLP[113]GVAGAP[113] | AAGLP(1.0000)GVAGAP(1.0000) | 19.41 | 12.60 | 0.975 |
| 914.488 | COL1A2 | 626 | 636 | VVGAVGTAGPS | VVGAVGTAGPS | | 35.01 | 15.94 | 1.000 |
| 915.376 | COL1A2 | 1102 | 1111 | GVSGGGYDFG | GVSGGGYDFG | | 20.56 | 12.74 | 0.997 |
| 924.511 | COL1A2 | 459 | 468 | IGPAGKEGPV | IGPAGKEGPV | | 19.69 | 12.35 | 0.974 |
| 925.500 | COL1A2 | 891 | 900 | VGEPGPLGIA | VGEP[113]GPLGIA | VGEP(0.9891)GP(0.0109)LGIA | 18.81 | 14.58 | 0.951 |
| 939.546 | COL1A2 | 867 | 876 | LGAPGILGLP | LGAP[113]GILGLP[113] | LGAP(1.0000)GILGLP(1.0000) | 23.06 | 15.97 | 0.964 |
| 942.468 | COL1A2 | 574 | 582 | GLHGEFGLP | GLHGEFGLP[113] | GLHGEFGLP(1.0000) | 18.07 | 9.46 | 0.998 |
| 947.447 | COL1A2 | 949 | 958 | GYPGNIGPVG | GYP[113]GN[115]IGPVG | GYP(0.9793)GNIGP(0.0207)VG | 20.72 | 14.54 | 0.999 |
| 956.468 | COL1A2 | 632 | 642 | TAGPSGPSGLP | TAGPSGPSGLP[113] | TAGP(0.0054)SGP(0.0050)SGLP(0.9896) | 20.72 | 15.09 | 0.951 |
| 959.447 | COL1A2 | 550 | 559 | GPPGFQGLPG | GPP[113]GFQ[129]GLP[113]G | GP(0.0282)P(0.9859)GFQGLP(0.9859)G | 18.14 | 16.24 | 0.965 |
| 969.497 | COL1A2 | 310 | 321 | GAAGLPGVAGAP | GAAGLP[113]GVAGAP[113] | GAAGLP(1.0000)GVAGAP(1.0000) | 20.47 | 17.36 | 0.971 |
| 971.511 | COL1A2 | 625 | 636 | GVVGAVGTAGPS | GVVGAVGTAGPS | | 32.50 | 19.07 | 1.000 |
| 972.502 | COL1A2 | 623 | 633 | EPGVVGAVGTA | EP[113]GVVGAVGTA | EP(1.0000)GVVGAVGTA | 23.12 | 10.23 | 1.000 |

| | | | | | | | | | |
|---|---|---|---|---|---|---|---|---|---|
| 974.457 | COL1A2 | 69 | 79 | GPPGLGGNFAA | GPP[113]GLGGN[115]FAA | GP(0.0119)P(0.9881)GLGGNFAA | 18.03 | 16.11 | 0.990 |
| 978.408 | COL1A2 | 1099 | 1109 | GPPGVSGGGYD | GPP[113]GVSGGGYD | GP(0.0105)P(0.9895)GVSGGGYD | 26.45 | 24.17 | 1.000 |
| 980.512 | COL1A2 | 952 | 963 | GNIGPVGAAGAP | GNIGPVGAAGAP | | 19.98 | 11.55 | 0.994 |
| 983.374 | COL1A2 | 1110 | 1117 | FGYDGDFY | FGYDGDFY | | 20.52 | 10.18 | 0.999 |
| 986.469 | COL1A2 | 721 | 732 | GPNGFAGPAGAA | GPNGFAGPAGAA | | 18.02 | 8.08 | 0.998 |
| 987.443 | COL1A2 | 721 | 732 | GPNGFAGPAGAA | GPN[115]GFAGPAGAA | | 30.56 | 18.79 | 1.000 |
| 987.504 | COL1A2 | 625 | 636 | GVVGAVGTAGPS | GVVGAVGTAGP[113]S | GVVGAVGTAGP(1.0000)S | 19.05 | 13.47 | 0.992 |
| 996.504 | COL1A2 | 952 | 963 | GNIGPVGAAGAP | GNIGPVGAAGAP[113] | GNIGP(0.0100)VGAAGAP(0.9900) | 24.21 | 12.04 | 1.000 |
| 996.535 | COL1A2 | 890 | 900 | AVGEPGPLGIA | AVGEP[113]GPLGIA | AVGEP(0.9775)GP(0.0225)LGIA | 18.68 | 17.38 | 0.990 |
| 996.571 | COL1A2 | 867 | 877 | LGAPGILGLPG | LGAP[113]GILGLP[113]G | LGAP(1.0000)GILGLP(1.0000)G | 22.68 | 14.42 | 0.988 |
| 997.476 | COL1A2 | 427 | 437 | GPAGVRGPNGD | GPAGVRGPN[115]GD | | 18.59 | 10.25 | 0.985 |
| 997.492 | COL1A2 | 952 | 963 | GNIGPVGAAGAP | GN[115]IGPVGAAGAP[113] | GNIGP(0.0117)VGAAGAP(0.9883) | 21.14 | 14.48 | 0.974 |
| 999.468 | COL1A2 | 235 | 246 | GSDGSVGPVGPA | GSDGSVGPVGPA | | 21.49 | 13.10 | 0.977 |
| 999.517 | COL1A2 | 826 | 836 | GPVGRTGEVGA | GPVGRTGEVGA | | 20.42 | 12.44 | 0.945 |
| 1012.501 | COL1A2 | 265 | 276 | GEIGAVGNAGPA | GEIGAVGNAGPA | | 24.14 | 15.49 | 0.997 |
| 1013.484 | COL1A2 | 265 | 276 | GEIGAVGNAGPA | GEIGAVGN[115]AGPA | | 27.72 | 21.09 | 1.000 |
| 1015.464 | COL1A2 | 910 | 921 | GAVGSPGVNGAP | GAVGSP[113]GVN[115]GAP[113] | GAVGSP(1.0000)GVNGAP(1.0000) | 22.19 | 15.69 | 0.988 |
| 1021.423 | COL1A2 | 1103 | 1112 | VSGGGYDFGY | VSGGGYDFGY | | 18.55 | 11.34 | 0.996 |
| 1029.515 | COL1A2 | 622 | 633 | GEPGVVGAVGTA | GEP[113]GVVGAVGTA | GEP(1.0000)GVVGAVGTA | 24.38 | 10.63 | 1.000 |
| 1032.467 | COL1A2 | 688 | 699 | GATGDRGEAGAA | GATGDRGEAGAA | | 21.69 | 14.66 | 0.966 |
| 1032.472 | COL1A2 | 438 | 447 | AGRPGEPGLM | AGRP[113]GEP[113]GLM[147] | AGRP(1.0000)GEP(1.0000)GLM(1.0000) | 22.90 | 11.14 | 0.998 |
| 1036.635 | COL1A2 | 866 | 876 | LLGAPGILGLP | LLGAPGILGLP[113] | LLGAP(0.0105)GILGLP(0.9895) | 22.75 | 10.19 | 0.983 |
| 1039.539 | COL1A2 | 886 | 897 | GVAGAVGEPGPL | GVAGAVGEP[113]GPL | GVAGAVGEP(0.9885)GP(0.0115)L | 21.50 | 18.34 | 0.976 |
| 1040.474 | COL1A2 | 772 | 783 | GPAGSRGDGGPP | GPAGSRGDGGPP[113] | GP(0.0054)AGSRGDGGP(0.0060)P(0.9886) | 18.86 | 17.22 | 0.986 |
| 1041.571 | COL1A2 | 827 | 837 | PVGRTGEVGAV | PVGRTGEVGAV | | 18.94 | 11.65 | 0.974 |
| 1052.625 | COL1A2 | 866 | 876 | LLGAPGILGLP | LLGAP[113]GILGLP[113] | LLGAP(1.0000)GILGLP(1.0000) | 27.58 | 14.21 | 0.977 |
| 1053.551 | COL1A2 | 889 | 900 | GAVGEPGPLGIA | GAVGEP[113]GPLGIA | GAVGEP(0.9888)GP(0.0112)LGIA | 25.79 | 18.11 | 1.000 |
| 1055.531 | COL1A2 | 883 | 894 | GLPGVAGAVGEP | GLP[113]GVAGAVGEP[113] | GLP(1.0000)GVAGAVGEP(1.0000) | 27.54 | 17.77 | 1.000 |
| 1058.520 | COL1A2 | 232 | 243 | GARGSDGSVGPV | GARGSDGSVGPV | | 25.69 | 12.01 | 0.998 |
| 1060.520 | COL1A2 | 86 | 96 | VGLGPGPMGLM | VGLGPGPM[147]GLM[147] | VGLGP(0.0133)GP(0.0356)M(0.9752)GLM(0.9759) | 25.92 | 19.89 | 1.000 |
| 1068.500 | COL1A2 | 427 | 438 | GPAGVRGPNGDA | GPAGVRGPN[115]GDA | | 19.24 | 12.60 | 0.966 |
| 1068.528 | COL1A2 | 451 | 462 | GLPGSPGNIGPA | GLP[113]GSP[113]GNIGPA | GLP(0.9876)GSP(0.9876)GNIGP(0.0248)A | 18.37 | 12.63 | 0.963 |
| 1069.565 | COL1A2 | 262 | 273 | GPKGEIGAVGNA | GPKGEIGAVGNA | | 19.95 | 12.32 | 0.983 |
| 1072.547 | COL1A2 | 949 | 960 | GYPGNIGPVGAA | GYPGNIGPVGAA | | 20.75 | 13.83 | 0.992 |
| 1075.492 | COL1A2 | 1034 | 1044 | IAGHHGDQGAP | IAGHHGDQGAP[113] | IAGHHGDQGAP(1.0000) | 18.13 | 11.23 | 0.994 |
| 1078.434 | COL1A2 | 1102 | 1112 | GVSGGGYDFGY | GVSGGGYDFGY | | 21.81 | 10.66 | 1.000 |
| 1080.571 | COL1A2 | 886 | 898 | GVAGAVGEPGPLG | GVAGAVGEPGPLG | | 23.99 | 20.06 | 0.980 |
| 1085.524 | COL1A2 | 247 | 258 | GPIGSAGPPGFP | GPIGSAGPP[113]GFP[113] | GP(0.0118)IGSAGP(0.0127)P(0.9877)GFP(0.9878) | 21.09 | 17.39 | 1.000 |
| 1088.533 | COL1A2 | 949 | 960 | GYPGNIGPVGAA | GYP[113]GNIGPVGAA | GYP(0.9896)GNIGP(0.0104)VGAA | 28.67 | 14.31 | 1.000 |
| 1089.513 | COL1A2 | 949 | 960 | GYPGNIGPVGAA | GYP[113]GN[115]IGPVGAA | GYP(0.9886)GNIGP(0.0114)VGAA | 22.81 | 14.46 | 0.964 |
| 1096.558 | COL1A2 | 886 | 898 | GVAGAVGEPGPLG | GVAGAVGEP[113]GPLG | GVAGAVGEP(0.9830)GP(0.0170)LG | 28.23 | 20.29 | 0.999 |
| 1096.577 | COL1A2 | 340 | 351 | GARGLVGEPGPA | GARGLVGEP[113]GPA | GARGLVGEP(0.9896)GP(0.0104)A | 22.22 | 16.52 | 0.994 |
| 1097.577 | COL1A2 | 283 | 294 | GEVGLPGLSGPV | GEVGLP[113]GLSGPV | GEVGLP(0.9863)GLSGP(0.0137)V | 20.21 | 11.39 | 0.966 |
| 1097.599 | COL1A2 | 281 | 291 | PRGEVGLPGLS | PRGEVGLP[113]GLS | P(0.0112)RGEVGLP(0.9888)GLS | 19.07 | 10.69 | 0.958 |
| 1098.571 | COL1A2 | 826 | 837 | GPVGRTGEVGAV | GPVGRTGEVGAV | | 18.01 | 10.28 | 0.967 |
| 1099.536 | COL1A2 | 454 | 465 | GSPGNIGPAGKE | GSP[113]GNIGPAGKE | GSP(0.9886)GNIGP(0.0114)AGKE | 18.94 | 10.14 | 1.000 |
| 1100.520 | COL1A2 | 454 | 465 | GSPGNIGPAGKE | GSP[113]GN[115]IGPAGKE | GSP(0.9891)GNIGP(0.0109)AGKE | 18.31 | 11.04 | 0.997 |
| 1101.519 | COL1A2 | 247 | 258 | GPIGSAGPPGFP | GPIGSAGP[113]P[113]GFP[113] | GP(0.0344)IGSAGP(0.9885)P(0.9886)GFP(0.9885) | 20.75 | 12.12 | 1.000 |
| 1101.569 | COL1A2 | 158 | 168 | VVGPQGARGFP | VVGPQ[129]GARGFP[113] | VVGP(0.0119)QGARGFP(0.9881) | 18.00 | 10.19 | 0.999 |
| 1109.606 | COL1A2 | 1072 | 1083 | GTVGPAGIRGPQ | GTVGPAGIRGPQ | | 22.06 | 10.09 | 1.000 |
| 1109.652 | COL1A2 | 865 | 876 | GLLGAPGILGLP | GLLGAP[113]GILGLP[113] | GLLGAP(1.0000)GILGLP(1.0000) | 27.51 | 13.70 | 1.000 |
| 1110.561 | COL1A2 | 575 | 585 | LHGEFGLPGPA | LHGEFGLP[113]GPA | LHGEFGLP(0.9891)GP(0.0109)A | 20.75 | 15.87 | 0.961 |
| 1112.532 | COL1A2 | 709 | 720 | GSPGERGEVGPA | GSPGERGEVGPA | | 20.41 | 13.18 | 0.990 |
| 1112.555 | COL1A2 | 745 | 756 | GPKGENGVVGPT | GPKGEN[115]GVVGPT | | 19.56 | 15.14 | 0.920 |
| 1112.570 | COL1A2 | 829 | 840 | GRTGEVGAVGPP | GRTGEVGAVGPP[113] | GRTGEVGAVGP(0.0131)P(0.9869) | 20.27 | 20.27 | 0.952 |
| 1116.505 | COL1A2 | 409 | 420 | GADGRAGVMGPP | GADGRAGVM[147]GPP[113] | GADGRAGVM(0.9857)GP(0.0286)P(0.9856) | 22.89 | 22.89 | 0.959 |
| 1117.475 | COL1A2 | 113 | 123 | FQGPAGEPGEP | FQGPAGEP[113]GEP[113] | FQGP(0.0195)AGEP(0.9902)GEP(0.9902) | 19.17 | 11.85 | 0.981 |
| 1117.532 | COL1A2 | 85 | 96 | GVGLGPGPMGLM | GVGLGPGP[113]MGLM[147] | GVGLGP(0.0144)GP(0.6113)M(0.6113)GLM(0.7631) | 18.10 | 14.55 | 0.975 |
| 1123.558 | COL1A2 | 856 | 867 | GPPGTPGPQGLL | GPP[113]GTP[113]GPQ[129]GLL | GP(0.0117)P(0.9887)GTP(0.9887)GP(0.0109)QGLL | 21.58 | 19.55 | 0.995 |
| 1125.474 | COL1A2 | 1099 | 1110 | GPPGVSGGGYDF | GPP[113]GVSGGGYDF | GP(0.0106)P(0.9894)GVSGGGYDF | 21.64 | 19.61 | 1.000 |
| 1127.572 | COL1A2 | 745 | 756 | GPKGENGVVGPT | GP[113]KGENGVVGPT | GP(0.9843)KGENGVVGP(0.0157)T | 21.57 | 15.72 | 0.976 |
| 1128.524 | COL1A2 | 709 | 720 | GSPGERGEVGPA | GSP[113]GERGEVGPA | GSP(0.9888)GERGEVGP(0.0112)A | 19.21 | 11.86 | 0.978 |
| 1128.544 | COL1A2 | 745 | 756 | GPKGENGVVGPT | GP[113]KGEN[115]GVVGPT | GP(0.9864)KGENGVVGP(0.0136)T | 20.40 | 11.51 | 0.990 |
| 1132.507 | COL1A2 | 1033 | 1044 | GIAGHHGDQGAP | GIAGHHGDQGAP[113] | GIAGHHGDQGAP(1.0000) | 22.05 | 11.66 | 0.996 |
| 1136.445 | COL1A2 | 1103 | 1113 | VSGGGYDFGYD | VSGGGYDFGYD | | 22.40 | 10.45 | 1.000 |
| 1139.483 | COL1A2 | 1110 | 1118 | FGYDGDFYR | FGYDGDFYR | | 18.29 | 10.03 | 0.974 |
| 1142.549 | COL1A2 | 550 | 561 | GPPGFQGLPGPS | GPP[113]GFQGLP[113]GPS | GP(0.1566)P(0.8685)GFQGLP(0.8721)GP(0.1028)S | 22.04 | 21.92 | 0.999 |
| 1150.617 | COL1A2 | 466 | 477 | GPVGLPGIDGRP | GPVGLP[113]GIDGRP | GP(0.0052)VGLP(0.9779)GIDGRP(0.0169) | 29.36 | 14.42 | 0.998 |
| 1154.522 | COL1A2 | 784 | 795 | GMTGFPGAAGRT | GM[147]TGFP[113]GAAGRT | GM(1.0000)TGFP(1.0000)GAAGRT | 19.70 | 9.35 | 1.000 |
| 1154.590 | COL1A2 | 280 | 291 | GPRGEVGLPGLS | GPRGEVGLP[113]GLS | GP(0.0350)RGEVGLP(0.9650)GLS | 21.03 | 11.52 | 1.000 |
| 1154.603 | COL1A2 | 463 | 474 | GKEGPVGLPGID | GKEGPVGLP[113]GID | GKEGP(0.0110)VGLP(0.9890)GID | 27.21 | 16.04 | 0.989 |
| 1166.607 | COL1A2 | 466 | 477 | GPVGLPGIDGRP | GPVGLP[113]GIDGRP[113] | GP(0.0212)VGLP(0.9894)GIDGRP(0.9894) | 29.59 | 13.39 | 0.988 |

| | | | | | | | | | |
|---|---|---|---|---|---|---|---|---|---|
| 1169.584 | COL1A2 | 562 | 573 | GPAGEVGKPGER | GPAGEVGKP[113]GER | GP(0.0109)AGEVGKP(0.9891)GER | 18.81 | 10.02 | 0.946 |
| 1174.590 | COL1A2 | 163 | 174 | GARGFPGTPGLP | GARGFP[113]GTP[113]GLP[113] | GARGFP(1.0000)GTP(1.0000)GLP(1.0000) | 19.56 | 13.90 | 0.973 |
| 1174.596 | COL1A2 | 1078 | 1089 | GIRGPQGHQGPA | GIRGPQGHQGPA | | 26.76 | 10.68 | 1.000 |
| 1182.478 | COL1A2 | 1099 | 1111 | GPPGVSGGGYDFG | GPP[113]GVSGGGYDFG | GP(0.0105)P(0.9895)GVSGGGYDFG | 39.16 | 36.56 | 1.000 |
| 1193.464 | COL1A2 | 1102 | 1113 | GVSGGGYDFGYD | GVSGGGYDFGYD | | 32.81 | 15.51 | 1.000 |
| 1198.506 | COL1A2 | 1099 | 1111 | GPPGVSGGGYDFG | GP[113]P[113]GVSGGGYDFG | GP(1.0000)P(1.0000)GVSGGGYDFG | 18.70 | 8.04 | 1.000 |
| 1203.568 | COL1A2 | 484 | 495 | GARGEPGNIGFP | GARGEP[113]GNIGFP[113] | GARGEP(1.0000)GNIGFP(1.0000) | 19.76 | 11.75 | 0.946 |
| 1204.520 | COL1A2 | 436 | 447 | GDAGRPGEPGLM | GDAGRP[113]GEP[113]GLM[147] | GDAGRP(1.0000)GEP(1.0000)GLM(1.0000) | 21.23 | 11.31 | 0.988 |
| 1204.558 | COL1A2 | 484 | 495 | GARGEPGNIGFP | GARGEP[113]GN[115]IGFP[113] | GARGEP(1.0000)GNIGFP(1.0000) | 18.40 | 12.72 | 0.973 |
| 1207.670 | COL1A2 | 459 | 471 | IGPAGKEGPVGLP | IGPAGKEGPVGLP[113] | IGP(0.0097)AGKEGP(0.0081)VGLP(0.9822) | 27.41 | 21.97 | 0.996 |
| 1210.516 | COL1A2 | 1110 | 1119 | FGYDGDFYRA | FGYDGDFYRA | | 19.12 | 9.81 | 0.999 |
| 1211.623 | COL1A2 | 463 | 475 | GKEGPVGLPGIDG | GKEGPVGLP[113]GIDG | GKEGP(0.0106)VGLP(0.9894)GIDG | 25.56 | 13.77 | 0.999 |
| 1220.649 | COL1A2 | 219 | 231 | PGERGRVGAPGPA | PGERGRVGAPGPA | | 19.61 | 14.94 | 0.926 |
| 1222.657 | COL1A2 | 1071 | 1083 | PGTVGPAGIRGPQ | P[113]GTVGPAGIRGPQ | P(0.9892)GTVGP(0.0055)AGIRGP(0.0054)Q | 21.35 | 13.44 | 0.994 |
| 1223.654 | COL1A2 | 887 | 900 | VAGAVGEPGPLGIA | VAGAVGEP[113]GPLGIA | VAGAVGEP(0.9528)GP(0.0472)LGIA | 26.13 | 22.26 | 0.956 |
| 1226.606 | COL1A2 | 562 | 574 | GPAGEVGKPGERG | GPAGEVGKP[113]GERG | GP(0.0103)AGEVGKP(0.9897)GERG | 20.70 | 12.84 | 0.991 |
| 1235.655 | COL1A2 | 1068 | 1080 | TGHPGTVGPAGIR | TGHP[113]GTVGPAGIR | TGHP(0.9886)GTVGP(0.0114)AGIR | 19.95 | 9.80 | 1.000 |
| 1237.667 | COL1A2 | 178 | 189 | GIRGHNGLDGLK | GIRGHN[115]GLDGLK | | 22.72 | 14.50 | 1.000 |
| 1250.483 | COL1A2 | 1102 | 1114 | GVSGGGYDFGYDG | GVSGGGYDFGYDG | | 30.21 | 10.73 | 1.000 |
| 1257.569 | COL1A2 | 685 | 699 | GPAGATGDRGEAGAA | GPAGATGDRGEAGAA | | 36.58 | 13.08 | 1.000 |
| 1263.655 | COL1A2 | 890 | 903 | AVGEPGPLGIAGPP | AVGEP[113]GPLGIAGPP[113] | AVGEP(0.6118)GP(0.6118)LGIAGP(0.0093)P(0.7671) | 19.14 | 19.14 | 0.986 |
| 1270.621 | COL1A2 | 622 | 636 | GEPGVVGAVGTAGPS | GEP[113]GVVGAVGTAGPS | GEP(0.9896)GVVGAVGTAGP(0.0104)S | 27.92 | 15.75 | 1.000 |
| 1272.609 | COL1A2 | 298 | 312 | GNPGANGLTGAKGAA | GNP[113]GAN[115]GLTGAKGAA | GNP(1.0000)GANGLTGAKGAA | 19.90 | 11.82 | 1.000 |
| 1278.590 | COL1A2 | 842 | 855 | FAGEKGPSGEAGTA | FAGEKGPSGEAGTA | | 35.71 | 11.33 | 1.000 |
| 1280.679 | COL1A2 | 886 | 900 | GVAGAVGEPGPLGIA | GVAGAVGEP[113]GPLGIA | GVAGAVGEP(0.9870)GP(0.0130)LGIA | 29.73 | 23.51 | 1.000 |
| 1283.620 | COL1A2 | 232 | 246 | GARGSDGSVGPVGPA | GARGSDGSVGPVGPA | | 26.19 | 13.93 | 0.993 |
| 1284.630 | COL1A2 | 571 | 582 | GERGLHGEFGLP | GERGLHGEFGLP[113] | GERGLHGEFGLP(1.0000) | 18.27 | 10.82 | 0.976 |
| 1294.587 | COL1A2 | 842 | 855 | FAGEKGPSGEAGTA | FAGEKGP[113]SGEAGTA | FAGEKGP(1.0000)SGEAGTA | 24.87 | 12.51 | 1.000 |
| 1294.667 | COL1A2 | 262 | 276 | GPKGEIGAVGNAGPA | GPKGEIGAVGNAGPA | | 20.02 | 13.53 | 0.987 |
| 1295.647 | COL1A2 | 262 | 276 | GPKGEIGAVGNAGPA | GPKGEIGAVGN[115]AGPA | | 34.89 | 21.74 | 1.000 |
| 1296.566 | COL1A2 | 116 | 129 | PAGEPGEPGQTGPA | PAGEP[113]GEP[113]GQTGPA | P(0.0281)AGEP(0.9802)GEP(0.9809)GQTGP(0.0109)A | 22.72 | 20.36 | 0.991 |
| 1310.622 | COL1A2 | 247 | 261 | GPIGSAGPPGFPGAP | GPIGSAGP[113]PGFPGAP[113] | GP(0.0162)IGSAGP(0.5833)P(0.5833)GFP(0.0062)GAP(0.8109) | 18.87 | 18.87 | 0.992 |
| 1310.669 | COL1A2 | 262 | 276 | GPKGEIGAVGNAGPA | GP[113]KGEIGAVGNAGPA | GP(0.9897)KGEIGAVGNAGP(0.0103)A | 35.47 | 24.11 | 1.000 |
| 1311.648 | COL1A2 | 262 | 276 | GPKGEIGAVGNAGPA | GP[113]KGEIGAVGN[115]AGPA | GP(0.9886)KGEIGAVGNAGP(0.0114)A | 34.60 | 23.60 | 0.999 |
| 1320.679 | COL1A2 | 889 | 903 | GAVGEPGPLGIAGPP | GAVGEP[113]GPLGIAGPP[113] | GAVGEP(0.9631)GP(0.0150)LGIAGP(0.0609)P(0.9610) | 28.11 | 25.89 | 0.999 |
| 1325.678 | COL1A2 | 337 | 351 | GATGARGLVGEPGPA | GATGARGLVGEP[113]GPA | GATGARGLVGEP(0.9885)GP(0.0115)A | 24.47 | 18.78 | 0.997 |
| 1326.628 | COL1A2 | 247 | 261 | GPIGSAGPPGFPGAP | GPIGSAGPP[113]GFP[113]GAP[113] | GP(0.0175)IGSAGP(0.0332)P(0.9829)GFP(0.9832)GAP(0.9832) | 19.96 | 19.96 | 0.990 |
| 1328.687 | COL1A2 | 619 | 633 | GNKGEPGVVGAVGTA | GNKGEP[113]GVVGAVGTA | GNKGEP(1.0000)GVVGAVGTA | 18.38 | 9.16 | 0.998 |
| 1329.636 | COL1A2 | 949 | 963 | GYPGNIGPVGAAGAP | GYP[113]GNIGPVGAAGAP[113] | GYP(0.9866)GNIGP(0.0268)VGAAGAP(0.9866) | 27.85 | 18.91 | 1.000 |
| 1330.620 | COL1A2 | 949 | 963 | GYPGNIGPVGAAGAP | GYP[113]GN[115]IGPVGAAGAP[113] | GYP(0.9891)GNIGP(0.0219)VGAAGAP(0.9890) | 23.36 | 14.70 | 0.992 |
| 1335.604 | COL1A2 | 841 | 855 | GFAGEKGPSGEAGTA | GFAGEKGPSGEAGTA | | 45.78 | 22.55 | 1.000 |
| 1339.692 | COL1A2 | 562 | 575 | GPAGEVGKPGERGL | GPAGEVGKP[113]GERGL | GP(0.0103)AGEVGKP(0.9897)GERGL | 18.23 | 11.70 | 1.000 |
| 1345.564 | COL1A2 | 1099 | 1112 | GPPGVSGGGYDFGY | GP[113]PGVSGGGYDFGY | GP(0.5000)P(0.5000)GVSGGGYDFGY | 26.21 | 26.21 | 1.000 |
| 1351.610 | COL1A2 | 841 | 855 | GFAGEKGPSGEAGTA | GFAGEKGP[113]SGEAGTA | GFAGEKGP(1.0000)SGEAGTA | 25.16 | 15.31 | 0.999 |
| 1351.694 | COL1A2 | 637 | 651 | GPSGLPGERGAAGIP | GPSGLPGERGAAGIP[113] | GP(0.0063)SGLP(0.0073)GERGAAGIP(0.9864) | 29.31 | 19.86 | 0.994 |
| 1352.679 | COL1A2 | 454 | 468 | GSPGNIGPAGKEGPV | GSP[113]GNIGPAGKEGPV | GSP(0.9887)GNIGP(0.0056)AGKEGP(0.0057)V | 28.86 | 16.09 | 1.000 |
| 1353.580 | COL1A2 | 115 | 129 | GPAGEPGEPGQTGPA | GPAGEP[113]GEP[113]GQTGPA | GP(0.0669)AGEP(0.9601)GEP(0.9625)GQTGP(0.0105)A | 30.06 | 29.77 | 0.999 |
| 1353.641 | COL1A2 | 454 | 468 | GSPGNIGPAGKEGPV | GSP[113]GN[115]IGPAGKEGPV | GSP(0.9868)GNIGP(0.0063)AGKEGP(0.0069)V | 23.46 | 17.91 | 1.000 |
| 1354.563 | COL1A2 | 115 | 129 | GPAGEPGEPGQTGPA | GPAGEP[113]GEP[113]GQ[129]TGPA | GP(0.0107)AGEP(0.9880)GEP(0.9879)GQTGP(0.0133)A | 25.91 | 21.86 | 0.969 |
| 1365.693 | COL1A2 | 826 | 840 | GPVGRTGEVGAVGPP | GPVGRTGEVGAVGPP[113] | GP(0.0053)VGRTGEVGAVGP(0.0057)P(0.9890) | 45.20 | 45.20 | 1.000 |
| 1367.694 | COL1A2 | 637 | 651 | GPSGLPGERGAAGIP | GPSGLP[113]GERGAAGIP[113] | GP(0.0268)SGLP(0.9866)GERGAAGIP(0.9865) | 22.81 | 17.72 | 0.980 |
| 1370.618 | COL1A2 | 115 | 129 | GPAGEPGEPGQTGPA | GP[113]AGEP[113]GEPGQTGP[113]A | GP(0.7641)AGEP(0.7621)GEP(0.6819)GQTGP(0.7919)A | 19.36 | 19.18 | 0.983 |
| 1372.683 | COL1A2 | 158 | 171 | VVGPQGARGFPGTP | VVGPQ[129]GARGFP[113]GTP[113] | VVGP(0.0406)QGARGFP(0.9796)GTP(0.9797) | 18.47 | 10.75 | 0.997 |
| 1378.731 | COL1A2 | 457 | 471 | GNIGPAGKEGPVGLP | GNIGPAGKEGPVGLP[113] | GNIGP(0.0117)AGKEGP(0.0055)VGLP(0.9828) | 30.46 | 25.53 | 1.000 |
| 1379.711 | COL1A2 | 883 | 898 | GLPGVAGAVGEPGPLG | GLP[113]GVAGAVGEPGP[113]LG | GLP(0.7895)GVAGAVGEP(0.4335)GP(0.7770)LG | 31.81 | 29.46 | 0.993 |
| 1379.711 | COL1A2 | 460 | 474 | GPAGKEGPVGLPGID | GPAGKEGPVGLP[113]GID | GP(0.0057)AGKEGP(0.0055)VGLP(0.9888)GID | 19.60 | 16.33 | 0.952 |
| 1381.707 | COL1A2 | 826 | 840 | GPVGRTGEVGAVGPP | GPVGRTGEVGAVGP[113]P[113] | GP(0.0331)VGRTGEVGAVGP(0.9835)P(0.9835) | 27.29 | 17.93 | 0.992 |
| 1384.693 | COL1A2 | 661 | 674 | GLRGEIGNPGRDGA | GLRGEIGNP[113]GRDGA | GLRGEIGNP(1.0000)GRDGA | 19.86 | 11.87 | 1.000 |
| 1384.708 | COL1A2 | 300 | 315 | PGANGLTGAKGAAGLP | P[113]GAN[115]GLTGAKGAAGLP[113] | P(1.0000)GANGLTGAKGAAGLP(1.0000) | 27.72 | 11.49 | 1.000 |
| 1391.782 | COL1A2 | 862 | 876 | GPQGLLGAPGILGLP | GPQGLLGAP[113]GILGLP[113] | GP(0.0316)QGLLGAP(0.9842)GILGLP(0.9842) | 23.27 | 12.26 | 0.987 |
| 1397.699 | COL1A2 | 880 | 894 | GERGLPGVAGAVGEP | GERGLP[113]GVAGAVGEP[113] | GERGLP(1.0000)GVAGAVGEP(1.0000) | 21.60 | 17.98 | 0.933 |
| 1398.685 | COL1A2 | 823 | 837 | GDQGPVGRTGEVGAV | GDQGPVGRTGEVGAV | | 34.61 | 13.37 | 1.000 |
| 1398.703 | COL1A2 | 621 | 636 | KGEPGVVGAVGTAGPS | KGEP[113]GVVGAVGTAGPS | KGEP(0.9894)GVVGAVGTAGP(0.0106)S | 31.22 | 11.90 | 1.000 |
| 1399.623 | COL1A2 | 196 | 210 | GVKGEPGAPGENGTP | GVKGEP[113]GAP[113]GEN[115]GTP | GVKGEP(0.9835)GAP(0.9834)GENGTP(0.0330) | 23.89 | 14.66 | 0.982 |
| 1403.599 | COL1A2 | 113 | 126 | FQGPAGEPGEPGQT | FQGPAGEP[113]GEP[113]GQT | FQGP(0.0215)AGEP(0.9893)GEP(0.9893)GQT | 26.33 | 18.37 | 1.000 |
| 1407.737 | COL1A2 | 280 | 294 | GPRGEVGLPGLSGPV | GPRGEVGLP[113]GLSGPV | GP(0.0059)RGEVGLP(0.9885)GLSGP(0.0056)V | 27.46 | 14.34 | 1.000 |
| 1408.672 | COL1A2 | 535 | 549 | GPQGVQGGKGEQGPP | GPQGVQGGKGEQGPP[113] | GP(0.0060)QGVQGGKGEQGP(0.0063)P(0.9876) | 22.51 | 22.51 | 1.000 |
| 1409.655 | COL1A2 | 535 | 549 | GPQGVQGGKGEQGPP | GPQGVQ[129]GGKGEQGPP[113] | GP(0.0142)QGVQGGKGEQGP(0.0064)P(0.9794) | 24.96 | 24.96 | 0.957 |
| 1415.613 | COL1A2 | 196 | 210 | GVKGEPGAPGENGTP | GVKGEP[113]GAP[113]GEN[115]GTP[113] | GVKGEP(1.0000)GAP(1.0000)GENGTP(1.0000) | 23.45 | 9.97 | 0.986 |
| 1416.723 | COL1A2 | 1069 | 1083 | GHPGTVGPAGIRGPQ | GHP[113]GTVGPAGIRGPQ | GHP(0.9871)GTVGP(0.0068)AGIRGP(0.0061)Q | 27.36 | 18.13 | 1.000 |
| 1417.716 | COL1A2 | 1069 | 1083 | GHPGTVGPAGIRGPQ | GHP[113]GTVGPAGIRGPQ[129] | GHP(0.9893)GTVGP(0.0053)AGIRGP(0.0054)Q | 19.28 | 11.83 | 0.999 |
| 1421.685 | COL1A2 | 970 | 984 | GPAGKHGNRGETGPS | GPAGKHGNRGETGPS | | 23.46 | 12.61 | 1.000 |

### Supplemental Table 3. Col1a1 peptide sequences from normal breast by LC-MS/MS.

| | | | | | | | | | |
|---|---|---|---|---|---|---|---|---|---|
| 1422.700 | COL1A2 | 1060 | 1074 | GPAGKDGRTGHPGTV | GPAGKDGRTGHP[113]GTV | GP(0.0157)AGKDGRTGHP(0.9843)GTV | 22.71 | 11.47 | 1.000 |
| 1422.734 | COL1A2 | 826 | 841 | GPVGRTGEVGAVGPPG | GPVGRTGEVGAVGPP[113]G | GP(0.0057)VGRTGEVGAVGP(0.0115)P(0.9828)G | 27.49 | 27.49 | 0.992 |
| 1425.681 | COL1A2 | 616 | 631 | GPDGNKGEPGVVGAVG | GPDGNKGEP[113]GVVGAVG | GP(0.0388)DGNKGEP(0.9612)GVVGAVG | 19.71 | 15.81 | 0.975 |
| 1428.685 | COL1A2 | 481 | 495 | GPAGARGEPGNIGFP | GPAGARGEP[113]GNIGFP[113] | GP(0.0277)AGARGEP(0.9861)GNIGFP(0.9861) | 18.32 | 12.41 | 0.971 |
| 1431.678 | COL1A2 | 946 | 960 | GERGYPGNIGPVGAA | GERGYP[113]GN[115]IGPVGAA | GERGYP(0.9703)GNIGP(0.0297)VGAA | 27.83 | 15.74 | 0.996 |
| 1432.689 | COL1A2 | 403 | 417 | GSRGLPGADGRAGVM | GSRGLP[113]GADGRAGVM[147] | GSRGLP(1.0000)GADGRAGVM(1.0000) | 24.05 | 11.75 | 1.000 |
| 1435.748 | COL1A2 | 457 | 472 | GNIGPAGKEGPVGLPG | GNIGPAGKEGPVGLP[113]G | GNIGP(0.0053)AGKEGP(0.0052)VGLP(0.9895)G | 25.01 | 16.14 | 0.996 |
| 1437.630 | COL1A2 | 784 | 798 | GMTGFPGAAGRTGPP | GM[147]TGFP[113]GAAGRTGP[113]P[113] | GM(1.0000)TGFP(1.0000)GAAGRTGP(1.0000)P(1.0000) | 28.03 | 10.42 | 1.000 |
| 1441.593 | COL1A2 | 922 | 936 | GEAGRDGNPGNDGPP | GEAGRDGNP[113]GNDGPP[113] | GEAGRDGNP(0.9883)GNDGP(0.0234)P(0.9883) | 18.40 | 18.40 | 0.976 |
| 1443.694 | COL1A2 | 618 | 633 | DGNKGEPGVVGAVGTA | DGNKGEP[113]GVVGAVGTA | DGNKGEP(1.0000)GVVGAVGTA | 22.19 | 11.86 | 1.000 |
| 1457.630 | COL1A2 | 433 | 447 | GPNGDAGRPGEPGLM | GPN[115]GDAGRP[113]GEP[113]GLM | GP(0.0115)NGDAGRP(0.9808)GEP(0.9802)GLM(0.0275) | 30.18 | 26.05 | 1.000 |
| 1460.589 | COL1A2 | 1099 | 1113 | GPPGVSGGGYDFGYD | GPP[113]GVSGGGYDFGYD | GP(0.0202)P(0.9798)GVSGGGYDFGYD | 18.18 | 16.30 | 0.996 |
| 1464.747 | COL1A2 | 463 | 477 | GKEGPVGLPGIDGRP | GKEGPVGLP[113]GIDGRP | GKEGP(0.0052)VGLP(0.9895)GIDGRP(0.0052) | 23.94 | 14.20 | 1.000 |
| 1465.660 | COL1A2 | 499 | 514 | GPTGDPGKNGDKGHAG | GPTGDPGKN[115]GDKGHAG | | 19.35 | 12.59 | 0.960 |
| 1472.637 | COL1A2 | 433 | 447 | GPNGDAGRPGEPGLM | GPNGDAGRP[113]GEP[113]GLM[147] | GP(0.0432)NGDAGRP(0.9856)GEP(0.9856)GLM(0.9856) | 20.20 | 7.08 | 1.000 |
| 1473.618 | COL1A2 | 433 | 447 | GPNGDAGRPGEPGLM | GPN[115]GDAGRP[113]GEP[113]GLM[147] | GP(0.0327)NGDAGRP(0.9891)GEP(0.9891)GLM(0.9891) | 28.92 | 11.74 | 1.000 |
| 1476.747 | COL1A2 | 562 | 576 | GPAGEVGKPGERGLH | GPAGEVGKP[113]GERGLH | GP(0.0123)AGEVGKP(0.9877)GERGLH | 18.34 | 10.54 | 0.996 |
| 1480.757 | COL1A2 | 463 | 477 | GKEGPVGLPGIDGRP | GKEGPVGLP[113]GIDGRP[113] | GKEGP(0.0224)VGLP(0.9888)GIDGRP(0.9888) | 25.12 | 13.83 | 1.000 |
| 1481.669 | COL1A2 | 499 | 514 | GPTGDPGKNGDKGHAG | GPTGDP[113]GKN[115]GDKGHAG | GP(0.0100)TGDP(0.9900)GKNGDKGHAG | 19.40 | 10.51 | 0.999 |
| 1482.686 | COL1A2 | 685 | 702 | GPAGATGDRGEAGAAGPA | GPAGATGDRGEAGAAGPA | | 32.71 | 12.15 | 1.000 |
| 1490.786 | COL1A2 | 887 | 903 | VAGAVGEPGPLGIAGPP | VAGAVGEP[113]GPLGIAGPP[113] | VAGAVGEP(0.9692)GP(0.0219)LGIAGP(0.0404)P(0.9685) | 28.50 | 28.50 | 0.997 |
| 1492.787 | COL1A2 | 459 | 474 | IGPAGKEGPVGLPGID | IGPAGKEGPVGLP[113]GID | IGP(0.0057)AGKEGP(0.0054)VGLP(0.9889)GID | 20.85 | 12.02 | 0.987 |
| 1498.677 | COL1A2 | 685 | 702 | GPAGATGDRGEAGAAGPA | GPAGATGDRGEAGAAGP[113]A | GP(0.0105)AGATGDRGEAGAAGP(0.9895)A | 31.29 | 13.24 | 1.000 |
| 1501.777 | COL1A2 | 1068 | 1083 | TGHPGTVGPAGIRGPQ | TGHPGTVGPAGIRGPQ | | 27.39 | 9.51 | 1.000 |
| 1508.736 | COL1A2 | 229 | 246 | GPAGARGSDGSVGPVGPA | GPAGARGSDGSVGPVGPA | | 28.30 | 12.96 | 1.000 |
| 1511.803 | COL1A2 | 1004 | 1017 | IRGDKGEPGEKGPR | IRGDKGEP[113]GEKGPR | IRGDKGEP(0.9829)GEKGP(0.0171)R | 20.76 | 14.84 | 1.000 |
| 1517.615 | COL1A2 | 1099 | 1114 | GPPGVSGGGYDFGYDG | GP[113]PGVSGGGYDFGYDG | GP(0.5000)P(0.5000)GVSGGGYDFGYDG | 35.81 | 35.81 | 1.000 |
| 1517.757 | COL1A2 | 1068 | 1083 | TGHPGTVGPAGIRGPQ | TGHP[113]GTVGPAGIRGPQ | TGHP(0.9861)GTVGP(0.0075)AGIRGP(0.0064)Q | 22.07 | 15.38 | 1.000 |
| 1518.747 | COL1A2 | 1068 | 1083 | TGHPGTVGPAGIRGPQ | TGHP[113]GTVGPAGIRGPQ[129] | TGHP(0.9891)GTVGP(0.0052)AGIRGP(0.0057)Q | 21.86 | 13.14 | 1.000 |
| 1524.805 | COL1A2 | 661 | 675 | GLRGEIGNPGRDGAR | GLRGEIGNPGRDGAR | | 21.13 | 12.29 | 0.994 |
| 1527.724 | COL1A2 | 1034 | 1050 | IAGHHGDQGAPGSVGPA | IAGHHGDQGAPGSVGPA | | 19.03 | 12.33 | 0.955 |
| 1533.775 | COL1A2 | 562 | 577 | GPAGEVGKPGERGLHG | GPAGEVGKP[113]GERGLHG | GP(0.0103)AGEVGKP(0.9897)GERGLHG | 19.99 | 12.32 | 0.997 |
| 1538.760 | COL1A2 | 820 | 835 | GPRGDQGPVGRTGEVG | GPRGDQGPVGRTGEVG | | 21.58 | 11.99 | 1.000 |
| 1539.750 | COL1A2 | 298 | 315 | GNPGANGLTGAKGAAGLP | GNPGAN[115]GLTGAKGAAGLP[113] | GNP(0.0189)GANGLTGAKGAAGLP(0.9811) | 24.50 | 17.97 | 1.000 |
| 1540.756 | COL1A2 | 617 | 633 | PDGNKGEPGVVGAVGTA | PDGNKGEP[113]GVVGAVGTA | P(0.0142)DGNKGEP(0.9858)GVVGAVGTA | 19.39 | 15.25 | 0.993 |
| 1540.794 | COL1A2 | 661 | 675 | GLRGEIGNPGRDGAR | GLRGEIGNP[113]GRDGAR | GLRGEIGNP(1.0000)GRDGAR | 18.28 | 8.44 | 1.000 |
| 1541.776 | COL1A2 | 661 | 675 | GLRGEIGNPGRDGAR | GLRGEIGN[115]P[113]GRDGAR | GLRGEIGNP(1.0000)GRDGAR | 19.33 | 14.46 | 1.000 |
| 1543.718 | COL1A2 | 1034 | 1050 | IAGHHGDQGAPGSVGPA | IAGHHGDQGAP[113]GSVGPA | IAGHHGDQGAP(0.9892)GSVGP(0.0108)A | 21.90 | 13.17 | 1.000 |
| 1545.710 | COL1A2 | 842 | 858 | FAGEKGPSGEAGTAGPP | FAGEKGPSGEAGTAGPP[113] | FAGEKGP(0.0058)SGEAGTAGP(0.0062)P(0.9880) | 45.77 | 45.77 | 1.000 |
| 1547.786 | COL1A2 | 886 | 903 | GVAGAVGEPGPLGIAGPP | GVAGAVGEP[113]GPLGIAGP[113]P | GVAGAVGEP(0.7372)GP(0.0663)LGIAGP(0.5982)P(0.5982) | 41.35 | 41.35 | 1.000 |
| 1552.807 | COL1A2 | 331 | 348 | GPVGAAGATGARGLVGEP | GPVGAAGATGARGLVGEP[113] | GP(0.0124)VGAAGATGARGLVGEP(0.9876) | 43.71 | 13.85 | 1.000 |
| 1555.766 | COL1A2 | 298 | 315 | GNPGANGLTGAKGAAGLP | GNP[113]GAN[115]GLTGAKGAAGLP[113] | GNP(1.0000)GANGLTGAKGAAGLP(1.0000) | 26.86 | 18.05 | 1.000 |
| 1556.724 | COL1A2 | 298 | 315 | GNPGANGLTGAKGAAGLP | GN[115]P[113]GAN[115]GLTGAKGAAGLP[113] | GNP(1.0000)GANGLTGAKGAAGLP(1.0000) | 22.99 | 13.20 | 1.000 |
| 1563.816 | COL1A2 | 883 | 900 | GLPGVAGAVGEPGPLGIA | GLP[113]GVAGAVGEP[113]GPLGIA | GLP(0.9714)GVAGAVGEP(0.9711)GP(0.0575)LGIA | 32.07 | 27.90 | 1.000 |
| 1569.777 | COL1A2 | 619 | 636 | GNKGEPGVVGAVGTAGPS | GNKGEP[113]GVVGAVGTAGPS | GNKGEP(0.9892)GVVGAVGTAGP(0.0108)S | 35.31 | 13.40 | 1.000 |
| 1581.779 | COL1A2 | 616 | 633 | GPDGNKGEPGVVGAVGTA | GPDGNKGEPGVVGAVGTA | | 30.21 | 11.53 | 1.000 |
| 1582.766 | COL1A2 | 616 | 633 | GPDGNKGEPGVVGAVGTA | GPDGN[115]KGEPGVVGAVGTA | | 19.44 | 10.48 | 0.993 |
| 1584.745 | COL1A2 | 1033 | 1050 | GIAGHHGDQGAPGSVGPA | GIAGHHGDQGAPGSVGPA | | 18.30 | 12.64 | 0.954 |
| 1597.752 | COL1A2 | 616 | 633 | GPDGNKGEPGVVGAVGTA | GPDGNKGEP[113]GVVGAVGTA | GP(0.0106)DGNKGEP(0.9894)GVVGAVGTA | 49.74 | 28.14 | 1.000 |
| 1598.759 | COL1A2 | 616 | 633 | GPDGNKGEPGVVGAVGTA | GPDGN[115]KGEP[113]GVVGAVGTA | GP(0.0103)DGNKGEP(0.9897)GVVGAVGTA | 47.84 | 31.64 | 1.000 |
| 1599.730 | COL1A2 | 910 | 927 | GAVGSPGVNGAPGEAGRD | GAVGSP[113]GVNGAP[113]GEAGRD | GAVGSP(1.0000)GVNGAP(1.0000)GEAGRD | 25.58 | 9.66 | 1.000 |
| 1600.699 | COL1A2 | 910 | 927 | GAVGSPGVNGAPGEAGRD | GAVGSP[113]GVN[115]GAP[113]GEAGRD | GAVGSP(1.0000)GVNGAP(1.0000)GEAGRD | 23.72 | 10.04 | 1.000 |
| 1600.733 | COL1A2 | 1033 | 1050 | GIAGHHGDQGAPGSVGPA | GIAGHHGDQGAP[113]GSVGPA | GIAGHHGDQGAP(0.9883)GSVGP(0.0117)A | 21.78 | 13.37 | 0.997 |
| 1602.723 | COL1A2 | 841 | 858 | GFAGEKGPSGEAGTAGPP | GFAGEKGPSGEAGTAGPP[113] | GFAGEKGP(0.0057)SGEAGTAGP(0.0062)P(0.9881) | 66.98 | 66.98 | 1.000 |
| 1602.735 | COL1A2 | 349 | 366 | GPAGSKGESGNKGEPGSA | GPAGSKGESGNKGEP[113]GSA | GP(0.0133)AGSKGESGNKGEP(0.9867)GSA | 21.61 | 12.75 | 1.000 |
| 1609.794 | COL1A2 | 820 | 836 | GPRGDQGPVGRTGEVGA | GPRGDQGPVGRTGEVGA | | 27.27 | 10.75 | 1.000 |
| 1610.782 | COL1A2 | 820 | 836 | GPRGDQGPVGRTGEVGA | GPRGDQ[129]GPVGRTGEVGA | | 19.27 | 11.23 | 0.993 |
| 1613.767 | COL1A2 | 616 | 633 | GPDGNKGEPGVVGAVGTA | GP[113]DGNKGEP[113]GVVGAVGTA | GP(1.0000)DGNKGEP(1.0000)GVVGAVGTA | 33.34 | 12.18 | 1.000 |
| 1614.757 | COL1A2 | 616 | 633 | GPDGNKGEPGVVGAVGTA | GP[113]DGN[115]KGEP[113]GVVGAVGTA | GP(1.0000)DGNKGEP(1.0000)GVVGAVGTA | 21.51 | 12.08 | 0.997 |
| 1618.635 | COL1A2 | 1103 | 1117 | VSGGGYDFGYDGDFY | VSGGGYDFGYDGDFY | | 33.04 | 12.49 | 1.000 |
| 1618.713 | COL1A2 | 841 | 858 | GFAGEKGPSGEAGTAGPP | GFAGEKGPSGEAGTAGP[113]P[113] | GFAGEKGP(0.0255)SGEAGTAGP(0.9872)P(0.9872) | 37.54 | 15.30 | 1.000 |
| 1628.698 | COL1A2 | 113 | 129 | FQGPAGEPGEPGQTGPA | FQGPAGEP[113]GEP[113]GQTGPA | FQGP(0.0118)AGEP(0.9836)GEP(0.9833)GQTGP(0.0213)A | 31.81 | 20.90 | 1.000 |
| 1629.693 | COL1A2 | 113 | 129 | FQGPAGEPGEPGQTGPA | FQGPAGEP[113]GEP[113]GQ[129]TGPA | FQGP(0.0110)AGEP(0.9863)GEP(0.9861)GQTGP(0.0167)A | 23.07 | 19.50 | 1.000 |
| 1631.814 | COL1A2 | 1060 | 1077 | GPAGKDGRTGHPGTVGPA | GPAGKDGRTGHPGTVGPA | | 20.22 | 9.84 | 1.000 |
| 1635.827 | COL1A2 | 454 | 471 | GSPGNIGPAGKEGPVGLP | GSP[113]GNIGPAGKEGPVGLP[113] | GSP(0.9865)GNIGP(0.0126)AGKEGP(0.0146)VGLP(0.9863) | 30.47 | 28.25 | 1.000 |
| 1636.815 | COL1A2 | 454 | 471 | GSPGNIGPAGKEGPVGLP | GSP[113]GN[115]IGPAGKEGPVGLP[113] | GSP(0.9818)GNIGP(0.0185)AGKEGP(0.0182)VGLP(0.9815) | 18.18 | 18.18 | 0.972 |
| 1640.724 | COL1A2 | 193 | 210 | GAPGVKGEPGAPGENGTP | GAP[113]GVKGEP[113]GAP[113]GEN[115]GTP | GAP(0.9885)GVKGEP(0.9885)GAP(0.9885)GENGTP(0.0344) | 22.38 | 15.31 | 1.000 |
| 1640.733 | COL1A2 | 712 | 729 | GERGEVGPAGPNGFAGPA | GERGEVGPAGPN[115]GFAGPA | | 24.73 | 11.97 | 1.000 |
| 1645.728 | COL1A2 | 113 | 129 | FQGPAGEPGEPGQTGPA | FQGP[113]AGEP[113]GEPGQTGP[113]A | FQGP(0.7368)AGEP(0.7673)GEP(0.7140)GQTGP(0.7819)A | 25.22 | 21.73 | 0.982 |
| 1646.728 | COL1A2 | 113 | 129 | FQGPAGEPGEPGQTGPA | FQGP[113]AGEP[113]GEPGQ[129]TGP[113]A | FQGP(0.7597)AGEP(0.7702)GEP(0.6721)GQTGP(0.7979)A | 21.10 | 20.66 | 0.955 |
| 1647.810 | COL1A2 | 1060 | 1077 | GPAGKDGRTGHPGTVGPA | GPAGKDGRTGHP[113]GTVGPA | GP(0.0078)AGKDGRTGHP(0.9861)GTVGP(0.0060)A | 21.77 | 14.53 | 0.977 |
| 1649.778 | COL1A2 | 499 | 516 | GPTGDPGKNGDKGHAGLA | GPTGDPGKN[115]GDKGHAGLA | | 20.50 | 9.60 | 1.000 |

### Supplemental Table 3. Col1a1 peptide sequences from normal breast by LC-MS/MS.

| | | | | | | | | | |
|---|---|---|---|---|---|---|---|---|---|
| 1656.728 | COL1A2 | 193 | 210 | GAPGVKGEPGAPGENGTP | GAP[113]GVKGEP[113]GAP[113]GEN[115]GTP[113] | GAP(1.0000)GVKGEP(1.0000)GAP(1.0000)GENGTP(1.0000) | 24.81 | 16.27 | 1.000 |
| 1656.753 | COL1A2 | 373 | 390 | GPSGEEGKRGPNGEAGSA | GPSGEEGKRGPNGEAGSA | | 23.55 | 12.30 | 1.000 |
| 1657.734 | COL1A2 | 373 | 390 | GPSGEEGKRGPNGEAGSA | GPSGEEGKRGPN[115]GEAGSA | | 20.22 | 11.68 | 0.994 |
| 1659.746 | COL1A2 | 352 | 369 | GSKGESGNKGEPGSAGPQ | GSKGESGNKGEP[113]GSAGPQ | GSKGESGNKGEP(0.9895)GSAGP(0.0105)Q | 24.33 | 15.53 | 0.991 |
| 1660.733 | COL1A2 | 352 | 369 | GSKGESGNKGEPGSAGPQ | GSKGESGN[115]KGEP[113]GSAGPQ | GSKGESGNKGEP(0.9898)GSAGP(0.0102)Q | 23.49 | 14.97 | 0.986 |
| 1662.815 | COL1A2 | 562 | 578 | GPAGEVGKPGERGLHGE | GPAGEVGKP[113]GERGLHGE | GP(0.0136)AGEVGKP(0.9864)GERGLHGE | 23.70 | 11.81 | 1.000 |
| 1664.810 | COL1A2 | 499 | 516 | GPTGDPGKNGDKGHAGLA | GPTGDP[113]GKNGDKGHAGLA | GP(0.0342)TGDP(0.9658)GKNGDKGHAGLA | 19.08 | 11.22 | 0.999 |
| 1665.774 | COL1A2 | 499 | 516 | GPTGDPGKNGDKGHAGLA | GPTGDP[113]GKN[115]GDKGHAGLA | GP(0.0142)TGDP(0.9858)GKNGDKGHAGLA | 19.31 | 12.87 | 0.989 |
| 1665.802 | COL1A2 | 586 | 603 | GPRGERGPPGESGAAGPT | GPRGERGP[113]PGESGAAGPT | GP(0.0042)RGERGP(0.9690)P(0.0234)GESGAAGP(0.0035)T | 19.82 | 19.82 | 0.966 |
| 1665.812 | COL1A2 | 823 | 840 | GDQGPVGRTGEVGAVGPP | GDQGPVGRTGEVGAVGPP[113] | GDQGP(0.0061)VGRTGEVGAVGP(0.0193)P(0.9746) | 41.94 | 41.94 | 1.000 |
| 1672.778 | COL1A2 | 946 | 963 | GERGYPGNIGPVGAAGAP | GERGYP[113]GN[115]IGPVGAAGAP[113] | GERGYP(0.9782)GNIGP(0.0436)VGAAGAP(0.9782) | 25.25 | 15.40 | 0.983 |
| 1674.819 | COL1A2 | 967 | 984 | GPVGPAGKHGNRGETGPS | GPVGPAGKHGNRGETGPS | | 20.04 | 12.26 | 0.974 |
| 1674.877 | COL1A2 | 280 | 297 | GPRGEVGLPGLSGPVGPP | GPRGEVGLP[113]GLSGPVGPP[113] | GP(0.0077)RGEVGLP(0.9877)GLSGP(0.0070)VGP(0.0101)P(0.9876) | 31.51 | 31.51 | 1.000 |
| 1675.644 | COL1A2 | 1102 | 1117 | GVSGGGYDFGYDGDFY | GVSGGGYDFGYDGDFY | | 22.92 | 10.70 | 1.000 |
| 1675.810 | COL1A2 | 967 | 984 | GPVGPAGKHGNRGETGPS | GPVGPAGKHGN[115]RGETGPS | | 20.70 | 10.77 | 1.000 |
| 1682.697 | COL1A2 | 919 | 936 | GAPGEAGRDGNPGNDGPP | GAP[113]GEAGRDGNP[113]GNDGPP[113] | GAP(0.9779)GEAGRDGNP(0.9779)GNDGP(0.0666)P(0.9777) | 18.71 | 18.71 | 1.000 |
| 1684.788 | COL1A2 | 618 | 636 | DGNKGEPGVVGAVGTAGPS | DGNKGEP[113]GVVGAVGTAGPS | DGNKGEP(0.9886)GVVGAVGTAGP(0.0114)S | 31.12 | 14.46 | 1.000 |
| 1685.732 | COL1A2 | 113 | 130 | FQGPAGEPGEPGQTGPAG | FQGPAGEP[113]GEP[113]GQTGPAG | FQGP(0.0133)AGEP(0.9749)GEP(0.9739)GQTGP(0.0378)AG | 33.99 | 24.85 | 1.000 |
| 1686.706 | COL1A2 | 113 | 130 | FQGPAGEPGEPGQTGPAG | FQGPAGEP[113]GEP[113]GQ[129]TGPAG | FQGP(0.0114)AGEP(0.9850)GEP(0.9847)GQTGP(0.0189)AG | 28.57 | 19.98 | 0.980 |
| 1692.836 | COL1A2 | 454 | 472 | GSPGNIGPAGKEGPVGLPG | GSP[113]GNIGPAGKEGPVGLP[113]G | GSP(0.9869)GNIGP(0.0146)AGKEGP(0.0115)VGLP(0.9869)G | 21.06 | 17.46 | 1.000 |
| 1693.832 | COL1A2 | 454 | 472 | GSPGNIGPAGKEGPVGLPG | GSP[113]GN[115]IGPAGKEGPVGLP[113]G | GSP(0.9826)GNIGP(0.0227)AGKEGP(0.0117)VGLP(0.9830)G | 19.73 | 15.63 | 0.943 |
| 1698.827 | COL1A2 | 643 | 660 | GERGAAGIPGGKGEKGEP | GERGAAGIP[113]GGKGEKGEP[113] | GERGAAGIP(1.0000)GGKGEKGEP(1.0000) | 19.40 | 11.49 | 0.993 |
| 1704.832 | COL1A2 | 1060 | 1078 | GPAGKDGRTGHPGTVGPAG | GPAGKDGRTGHP[113]GTVGPAG | GP(0.0122)AGKDGRTGHP(0.9814)GTVGP(0.0064)AG | 22.97 | 17.31 | 0.981 |
| 1707.803 | COL1A2 | 685 | 705 | GPAGATGDRGEAGAAGPAGPA | GPAGATGDRGEAGAAGPAGPA | | 20.88 | 14.05 | 0.983 |
| 1708.853 | COL1A2 | 820 | 837 | GPRGDQGPVGRTGEVGAV | GPRGDQGPVGRTGEVGAV | | 23.91 | 12.08 | 1.000 |
| 1709.848 | COL1A2 | 820 | 837 | GPRGDQGPVGRTGEVGAV | GPRGDQ[129]GPVGRTGEVGAV | | 24.30 | 12.54 | 1.000 |
| 1712.798 | COL1A2 | 431 | 447 | VRGPNGDAGRPGEPGLM | VRGPN[115]GDAGRP[113]GEPGLM[147] | VRGP(0.0130)NGDAGRP(0.9883)GEP(0.0102)GLM(0.9884) | 19.88 | 15.41 | 0.982 |
| 1724.882 | COL1A2 | 662 | 678 | LRGEIGNPGRDGARGAP | LRGEIGNP[113]GRDGARGAP[113] | LRGEIGNP(1.0000)GRDGARGAP(1.0000) | 20.33 | 11.34 | 0.994 |
| 1728.782 | COL1A2 | 431 | 447 | VRGPNGDAGRPGEPGLM | VRGPN[115]GDAGRP[113]GEP[113]GLM[147] | VRGP(0.0316)NGDAGRP(0.9895)GEP(0.9895)GLM(0.9895) | 20.41 | 11.69 | 0.970 |
| 1742.756 | COL1A2 | 109 | 126 | GPQGFQGPAGEPGEPGQT | GPQGFQGPAGEP[113]GEP[113]GQT | GP(0.0247)QGFQGP(0.0130)AGEP(0.9812)GEP(0.9811)GQT | 24.89 | 16.44 | 1.000 |
| 1743.737 | COL1A2 | 109 | 126 | GPQGFQGPAGEPGEPGQT | GPQGFQ[129]GPAGEP[113]GEP[113]GQT | GP(0.0283)QGFQGP(0.0140)AGEP(0.9808)GEP(0.9768)GQT | 28.16 | 24.28 | 1.000 |
| 1750.804 | COL1A2 | 427 | 445 | GPAGVRGPNGDAGRPGEPG | GPAGVRGPN[115]GDAGRP[113]GEP[113]G | GP(0.0190)AGVRGP(0.0440)NGDAGRP(0.9682)GEP(0.9688)G | 19.63 | 10.08 | 0.987 |
| 1753.847 | COL1A2 | 943 | 960 | GHKGERGYPGNIGPVGAA | GHKGERGYP[113]GN[115]IGPVGAA | GHKGERGYP(0.9894)GNIGP(0.0106)VGAA | 21.69 | 15.58 | 0.966 |
| 1769.811 | COL1A2 | 430 | 447 | GVRGPNGDAGRPGEPGLM | GVRGPN[115]GDAGRP[113]GEP[113]GLM | GVRGP(0.0106)NGDAGRP(0.9895)GEP(0.9896)GLM(0.0103) | 22.59 | 15.83 | 0.997 |
| 1771.769 | COL1A2 | 910 | 929 | GAVGSPGVNGAPGEAGRDGN | GAVGSP[113]GVN[115]GAP[113]GEAGRDGN | GAVGSP(1.0000)GVNGAP(1.0000)GEAGRDGN | 21.13 | 11.10 | 1.000 |
| 1772.740 | COL1A2 | 910 | 929 | GAVGSPGVNGAPGEAGRDGN | GAVGSP[113]GVN[115]GAP[113]GEAGRDGN[115] | GAVGSP(1.0000)GVNGAP(1.0000)GEAGRDGN | 19.88 | 12.75 | 0.989 |
| 1774.735 | COL1A2 | 1103 | 1118 | VSGGGYDFGYDGDFYR | VSGGGYDFGYDGDFYR | | 23.96 | 9.49 | 1.000 |
| 1777.910 | COL1A2 | 331 | 351 | GPVGAAGATGARGLVGEPGPA | GPVGAAGATGARGLVGEP[113]GPA | GP(0.0071)VGAAGATGARGLVGEP(0.9512)GP(0.0417)A | 35.50 | 35.50 | 1.000 |
| 1781.881 | COL1A2 | 661 | 678 | GLRGEIGNPGRDGARGAP | GLRGEIGNP[113]GRDGARGAP[113] | GLRGEIGNP(1.0000)GRDGARGAP(1.0000) | 23.03 | 18.77 | 1.000 |
| 1784.830 | COL1A2 | 430 | 447 | GVRGPNGDAGRPGEPGLM | GVRGPNGDAGRP[113]GEP[113]GLM[147] | GVRGP(0.0298)NGDAGRP(0.9901)GEP(0.9901)GLM(0.9900) | 19.66 | 12.14 | 0.996 |
| 1784.872 | COL1A2 | 661 | 678 | GLRGEIGNPGRDGARGAP | GLRGEIGN[115]P[113]GRDGARGAP[113] | GLRGEIGNP(1.0000)GRDGARGAP(1.0000) | 21.47 | 15.72 | 1.000 |
| 1785.808 | COL1A2 | 430 | 447 | GVRGPNGDAGRPGEPGLM | GVRGPN[115]GDAGRP[113]GEP[113]GLM[147] | GVRGP(0.0399)NGDAGRP(0.9867)GEP(0.9867)GLM(0.9867) | 20.54 | 12.81 | 0.999 |
| 1809.883 | COL1A2 | 562 | 579 | GPAGEVGKPGERGLHGEF | GPAGEVGKP[113]GERGLHGEF | GP(0.0132)AGEVGKP(0.9868)GERGLHGEF | 37.82 | 18.46 | 1.000 |
| 1816.821 | COL1A2 | 842 | 861 | FAGEKGPSGEAGTAGPPGTP | FAGEKGPSGEAGTAGPP[113]GTP[113] | FAGEKGP(0.0188)SGEAGTAGP(0.0181)P(0.9814)GTP(0.9817) | 18.15 | 18.15 | 0.942 |
| 1818.940 | COL1A2 | 459 | 477 | IGPAGKEGPVGLPGIDGRP | IGPAGKEGPVGLP[113]GIDGRP[113] | IGP(0.0247)AGKEGP(0.0118)VGLP(0.9818)GIDGRP(0.9818) | 23.48 | 13.83 | 0.966 |
| 1822.892 | COL1A2 | 616 | 636 | GPDGNKGEPGVVGAVGTAGPS | GPDGNKGEPGVVGAVGTAGPS | | 33.39 | 14.10 | 1.000 |
| 1830.953 | COL1A2 | 883 | 903 | GLPGVAGAVGEPGPLGIAGPP | GLP[113]GVAGAVGEP[113]GPLGIAGP[113]P | GLP(0.8001)GVAGAVGEP(0.8006)GP(0.0159)LGIAGP(0.6917)P(0.6917) | 48.41 | 48.41 | 1.000 |
| 1831.744 | COL1A2 | 1102 | 1118 | GVSGGGYDFGYDGDFYR | GVSGGGYDFGYDGDFYR | | 24.58 | 11.90 | 1.000 |
| 1838.852 | COL1A2 | 616 | 636 | GPDGNKGEPGVVGAVGTAGPS | GPDGNKGEP[113]GVVGAVGTAGPS | GP(0.0057)DGNKGEP(0.9882)GVVGAVGTAGP(0.0061)S | 54.93 | 34.46 | 1.000 |
| 1839.870 | COL1A2 | 616 | 636 | GPDGNKGEPGVVGAVGTAGPS | GPDGN[115]KGEP[113]GVVGAVGTAGPS | GP(0.0050)DGNKGEP(0.9901)GVVGAVGTAGP(0.0049)S | 42.74 | 32.46 | 1.000 |
| 1845.769 | COL1A2 | 1103 | 1119 | VSGGGYDFGYDGDFYRA | VSGGGYDFGYDGDFYRA | | 31.74 | 8.49 | 1.000 |
| 1850.946 | COL1A2 | 1000 | 1017 | GPQGIRGDKGEPGEKGPR | GPQGIRGDKGEP[113]GEKGPR | GP(0.0087)QGIRGDKGEP(0.9746)GEKGP(0.0167)R | 22.11 | 15.50 | 1.000 |
| 1851.921 | COL1A2 | 565 | 582 | GEVGKPGERGLHGEFGLP | GEVGKPGERGLHGEFGLP[113] | GEVGKP(0.0109)GERGLHGEFGLP(0.9891) | 19.76 | 13.31 | 0.938 |
| 1854.872 | COL1A2 | 616 | 636 | GPDGNKGEPGVVGAVGTAGPS | GP[113]DGNKGEP[113]GVVGAVGTAGPS | GP(0.9901)DGNKGEP(0.9901)GVVGAVGTAGP(0.0198)S | 24.19 | 10.80 | 1.000 |
| 1855.859 | COL1A2 | 616 | 636 | GPDGNKGEPGVVGAVGTAGPS | GPDGN[115]KGEP[113]GVVGAVGTAGP[113]S | GP(0.0470)DGNKGEP(0.9764)GVVGAVGTAGP(0.9766)S | 24.16 | 17.31 | 1.000 |
| 1856.839 | COL1A2 | 429 | 447 | AGVRGPNGDAGRPGEPGLM | AGVRGPN[115]GDAGRP[113]GEP[113]GLM[147] | AGVRGP(0.0355)NGDAGRP(0.9882)GEP(0.9882)GLM(0.9882) | 19.67 | 11.19 | 1.000 |
| 1858.898 | COL1A2 | 950 | 970 | YPGNIGPVGAAGAPGPHGPVG | YPGN[115]IGPVGAAGAP[113]GPHGPVG | YP(0.0025)GNIGP(0.0025)VGAAGAP(0.9892)GP(0.0029)HGP(0.0029)VG | 18.52 | 16.87 | 0.939 |
| 1866.941 | COL1A2 | 1000 | 1017 | GPQGIRGDKGEPGEKGPR | GP[113]QGIRGDKGEP[113]GEKGPR | GP(0.9888)QGIRGDKGEP(0.9905)GEKGP(0.0206)R | 21.63 | 13.15 | 1.000 |
| 1867.929 | COL1A2 | 565 | 582 | GEVGKPGERGLHGEFGLP | GEVGKP[113]GERGLHGEFGLP[113] | GEVGKP(1.0000)GERGLHGEFGLP(1.0000) | 30.27 | 16.62 | 1.000 |
| 1867.934 | COL1A2 | 1000 | 1017 | GPQGIRGDKGEPGEKGPR | GP[113]Q[129]GIRGDKGEP[113]GEKGPR | GP(0.9883)QGIRGDKGEP(0.9907)GEKGP(0.0210)R | 20.80 | 19.28 | 0.977 |
| 1868.825 | COL1A2 | 910 | 930 | GAVGSPGVNGAPGEAGRDGNP | GAVGSP[113]GVN[115]GAP[113]GEAGRDGNP | GAVGSP(0.9306)GVNGAP(0.9293)GEAGRDGNP(0.1401) | 22.23 | 19.98 | 0.999 |
| 1869.854 | COL1A2 | 838 | 858 | GPPGFAGEKGPSGEAGTAGPP | GPP[113]GFAGEKGPSGEAGTAGP[113]P | GP(0.0108)P(0.8094)GFAGEKGP(0.0066)SGEAGTAGP(0.5866)P(0.5866) | 26.77 | 26.77 | 1.000 |
| 1873.847 | COL1A2 | 841 | 861 | GFAGEKGPSGEAGTAGPPGTP | GFAGEKGPSGEAGTAGPP[113]GTP[113] | GFAGEKGP(0.0113)SGEAGTAGP(0.0120)P(0.9883)GTP(0.9884) | 35.55 | 33.28 | 1.000 |
| 1883.851 | COL1A2 | 910 | 930 | GAVGSPGVNGAPGEAGRDGNP | GAVGSP[113]GVNGAP[113]GEAGRDGNP[113] | GAVGSP(1.0000)GVNGAP(1.0000)GEAGRDGNP(1.0000) | 21.03 | 8.55 | 1.000 |
| 1884.809 | COL1A2 | 910 | 930 | GAVGSPGVNGAPGEAGRDGNP | GAVGSP[113]GVN[115]GAP[113]GEAGRDGNP[113] | GAVGSP(1.0000)GVNGAP(1.0000)GEAGRDGNP(1.0000) | 30.28 | 15.40 | 1.000 |
| 1884.853 | COL1A2 | 349 | 369 | GPAGSKGESGNKGEPGSAGPQ | GPAGSKGESGNKGEP[113]GSAGPQ | GP(0.0058)AGSKGESGNKGEP(0.9870)GSAGP(0.0072)Q | 27.91 | 21.19 | 1.000 |
| 1889.839 | COL1A2 | 841 | 861 | GFAGEKGPSGEAGTAGPPGTP | GFAGEKGP[113]SGEAGTAGPP[113]GTP[113] | GFAGEKGP(0.9854)SGEAGTAGP(0.0440)P(0.9854)GTP(0.9852) | 22.10 | 19.84 | 0.987 |
| 1902.791 | COL1A2 | 1102 | 1119 | GVSGGGYDFGYDGDFYRA | GVSGGGYDFGYDGDFYRA | | 22.85 | 9.59 | 1.000 |
| 1912.871 | COL1A2 | 113 | 132 | FQGPAGEPGEPGQTGPAGAR | FQGPAGEP[113]GEP[113]GQTGPAGAR | FQGP(0.0099)AGEP(0.9894)GEP(0.9894)GQTGP(0.0113)AGAR | 22.49 | 13.47 | 0.997 |
| 1913.848 | COL1A2 | 113 | 132 | FQGPAGEPGEPGQTGPAGAR | FQGPAGEP[113]GEP[113]GQ[129]TGPAGAR | FQGP(0.0100)AGEP(0.9885)GEP(0.9884)GQTGP(0.0131)AGAR | 26.45 | 19.43 | 0.976 |
| 1924.857 | COL1A2 | 373 | 393 | GPSGEEGKRGPNGEAGSAGPP | GPSGEEGKRGPN[115]GEAGSAGPP[113] | GP(0.0034)SGEEGKRGP(0.0034)NGEAGSAGP(0.0044)P(0.9888) | 19.37 | 19.37 | 0.940 |
| 1933.937 | COL1A2 | 499 | 519 | GPTGDPGKNGDKGHAGLAGAR | GPTGDPGKN[115]GDKGHAGLAGAR | | 28.35 | 20.74 | 1.000 |

### Supplemental Table 3. Col1a1 peptide sequences from normal breast by LC-MS/MS.

| | | | | | | | | | |
|---|---|---|---|---|---|---|---|---|---|
| 1937.822 | COL1A2 | 916 | 936 | GVNGAPGEAGRDGNPGNDGPP | GVN[115]GAP[113]GEAGRDGNPGNDGPP[113] | GVNGAP(0.9836)GEAGRDGNP(0.0135)GNDGP(0.0195)P(0.9834) | 19.37 | 19.37 | 1.000 |
| 1938.963 | COL1A2 | 564 | 582 | AGEVGKPGERGLHGEFGLP | AGEVGKP[113]GERGLHGEFGLP[113] | AGEVGKP(1.0000)GERGLHGEFGLP(1.0000) | 22.68 | 15.04 | 0.975 |
| 1942.768 | COL1A2 | 1099 | 1117 | GPPGVSGGGYDFGYDGDFY | GP[113]PGVSGGGYDFGYDGDFY | GP(0.5000)P(0.5000)GVSGGGYDFGYDGDFY | 39.51 | 39.51 | 1.000 |
| 1949.936 | COL1A2 | 499 | 519 | GPTGDPGKNGDKGHAGLAGAR | GPTGDP[113]GKN[115]GDKGHAGLAGAR | GP(0.0102)TGDP(0.9898)GKNGDKGHAGLAGAR | 28.19 | 22.84 | 1.000 |
| 1952.826 | COL1A2 | 916 | 936 | GVNGAPGEAGRDGNPGNDGPP | GVNGAP[113]GEAGRDGNP[113]GNDGPP[113] | GVNGAP(0.9840)GEAGRDGNP(0.9840)GNDGP(0.0480)P(0.9840) | 18.59 | 18.59 | 1.000 |
| 1953.801 | COL1A2 | 916 | 936 | GVNGAPGEAGRDGNPGNDGPP | GVN[115]GAP[113]GEAGRDGNP[113]GNDGP[113]P | GVNGAP(0.7802)GEAGRDGNP(0.7801)GNDGP(0.7198)P(0.7198) | 18.62 | 18.62 | 0.979 |
| 1959.807 | COL1A2 | 1099 | 1117 | GPPGVSGGGYDFGYDGDFY | GP[113]P[113]GVSGGGYDFGYDGDFY | GP(1.0000)P(1.0000)GVSGGGYDFGYDGDFY | 21.47 | 10.82 | 1.000 |
| 1967.856 | COL1A2 | 109 | 129 | GPQGFQGPAGEPGEPGQTGPA | GPQGFQGPAGEP[113]GEP[113]GQTGPA | GP(0.0131)QGFQGP(0.0071)AGEP(0.9612)GEP(0.9557)GQTGP(0.0628)A | 22.68 | 22.65 | 0.963 |
| 1968.837 | COL1A2 | 109 | 129 | GPQGFQGPAGEPGEPGQTGPA | GPQGFQGPAGEP[113]GEP[113]GQ[129]TGPA | GP(0.0068)QGFQGP(0.0068)AGEP(0.9291)GEP(0.9128)GQTGP(0.1445)A | 31.11 | 26.71 | 1.000 |
| 1969.835 | COL1A2 | 109 | 129 | GPQGFQGPAGEPGEPGQTGPA | GPQGFQ[129]GPAGEP[113]GEP[113]GQ[129]TGPA | GP(0.0068)QGFQGP(0.0068)AGEP(0.9864)GEP(0.9858)GQTGP(0.0141)A | 29.07 | 29.07 | 1.000 |
| 1974.005 | COL1A2 | 457 | 477 | GNIGPAGKEGPVGLPGIDGRP | GNIGPAGKEGPVGLP[113]GIDGRP | GNIGP(0.0042)AGKEGP(0.0040)VGLP(0.9825)GIDGRP(0.0093) | 35.33 | 30.31 | 1.000 |
| 1974.031 | COL1A2 | 1060 | 1080 | GPAGKDGRTGHPGTVGPAGIR | GPAGKDGRTGHP[113]GTVGPAGIR | GP(0.0054)AGKDGRTGHP(0.9895)GTVGP(0.0051)AGIR | 27.46 | 16.72 | 1.000 |
| 1975.989 | COL1A2 | 820 | 840 | GPRGDQGPVGRTGEVGAVGPP | GPRGDQGPVGRTGEVGAVGPP[113] | GP(0.0036)RGDQGP(0.0034)VGRTGEVGAVGP(0.1073)P(0.8858) | 31.21 | 31.21 | 1.000 |
| 1976.977 | COL1A2 | 820 | 840 | GPRGDQGPVGRTGEVGAVGPP | GPRGDQ[129]GPVGRTGEVGAVGP[113]P | GP(0.0100)RGDQGP(0.0033)VGRTGEVGAVGP(0.9783)P(0.0084) | 24.17 | 24.17 | 0.989 |
| 1983.850 | COL1A2 | 109 | 129 | GPQGFQGPAGEPGEPGQTGPA | GPQGFQGP[113]AGEP[113]GEP[113]GQTGPA | GP(0.0645)QGFQGP(0.9721)AGEP(0.9729)GEP(0.9729)GQTGP(0.0176)A | 33.01 | 26.35 | 1.000 |
| 1990.001 | COL1A2 | 457 | 477 | GNIGPAGKEGPVGLPGIDGRP | GNIGPAGKEGPVGLP[113]GIDGRP[113] | GNIGP(0.0116)AGKEGP(0.0116)VGLP(0.9884)GIDGRP(0.9884) | 22.53 | 13.57 | 1.000 |
| 1993.942 | COL1A2 | 427 | 447 | GPAGVRGPNGDAGRPGEPGLM | GPAGVRGPNGDAGRP[113]GEP[113]GLM | GP(0.0086)AGVRGP(0.0100)NGDAGRP(0.9869)GEP(0.9871)GLM(0.0074) | 27.68 | 22.46 | 1.000 |
| 1993.972 | COL1A2 | 943 | 963 | GHKGERGYPGNIGPVGAAGAP | GHKGERGYP[113]GNIGPVGAAGAP[113] | GHKGERGYP(0.9894)GNIGP(0.0211)VGAAGAP(0.9895) | 22.01 | 14.65 | 1.000 |
| 1994.913 | COL1A2 | 427 | 447 | GPAGVRGPNGDAGRPGEPGLM | GPAGVRGPN[115]GDAGRP[113]GEP[113]GLM | GP(0.0072)AGVRGP(0.0073)NGDAGRP(0.9892)GEP(0.9892)GLM(0.0072) | 44.41 | 36.74 | 1.000 |
| 1994.966 | COL1A2 | 943 | 963 | GHKGERGYPGNIGPVGAAGAP | GHKGERGYP[113]GN[115]IGPVGAAGAP[113] | GHKGERGYP(0.9746)GNIGP(0.0505)VGAAGAP(0.9749) | 19.64 | 14.60 | 0.998 |
| 2009.931 | COL1A2 | 427 | 447 | GPAGVRGPNGDAGRPGEPGLM | GPAGVRGPNGDAGRP[113]GEP[113]GLM[147] | GP(0.0325)AGVRGP(0.0164)NGDAGRP(0.9837)GEP(0.9837)GLM(0.9837) | 18.61 | 10.85 | 0.976 |
| 2010.914 | COL1A2 | 427 | 447 | GPAGVRGPNGDAGRPGEPGLM | GPAGVRGPN[115]GDAGRP[113]GEP[113]GLM[147] | GP(0.0181)AGVRGP(0.0191)NGDAGRP(0.9876)GEP(0.9876)GLM(0.9876) | 23.48 | 15.64 | 0.965 |
| 2023.990 | COL1A2 | 298 | 321 | GNPGANGLTGAKGAAGLPGVAGAP | GNP[113]GAN[115]GLTGAKGAAGLP[113]GVAGAP[113] | GNP(1.0000)GANGLTGAKGAAGLP(1.0000)GVAGAP(1.0000) | 18.62 | 18.62 | 0.942 |
| 2026.912 | COL1A2 | 427 | 447 | GPAGVRGPNGDAGRPGEPGLM | GP[113]AGVRGPN[115]GDAGRP[113]GEP[113]GLM[147] | GP(0.8410)AGVRGP(0.6147)NGDAGRP(0.8481)GEP(0.8481)GLM(0.8481) | 18.42 | 16.69 | 0.990 |
| 2031.057 | COL1A2 | 1063 | 1083 | GKDGRTGHPGTVGPAGIRGPQ | GKDGRTGHP[113]GTVGPAGIRGPQ | GKDGRTGHP(0.9872)GTVGP(0.0069)AGIRGP(0.0058)Q | 27.83 | 17.60 | 1.000 |
| 2033.012 | COL1A2 | 820 | 841 | GPRGDQGPVGRTGEVGAVGPPG | GPRGDQGPVGRTGEVGAVGPP[113]G | GP(0.0036)RGDQGP(0.0036)VGRTGEVGAVGP(0.0765)P(0.9163)G | 23.27 | 23.27 | 0.958 |
| 2034.009 | COL1A2 | 820 | 841 | GPRGDQGPVGRTGEVGAVGPPG | GPRGDQ[129]GPVGRTGEVGAVGP[113]PG | GP(0.0034)RGDQGP(0.0033)VGRTGEVGAVGP(0.4967)P(0.4967)G | 27.16 | 27.16 | 1.000 |
| 2067.052 | COL1A2 | 649 | 669 | GIPGGKGEKGEPGLRGEIGNP | GIP[113]GGKGEKGEP[113]GLRGEIGNP[113] | GIP(1.0000)GGKGEKGEP(1.0000)GLRGEIGNP(1.0000) | 24.05 | 12.11 | 1.000 |
| 2077.038 | COL1A2 | 562 | 582 | GPAGEVGKPGERGLHGEFGLP | GPAGEVGKPGERGLHGEFGLP[113] | GP(0.0060)AGEVGKP(0.0059)GERGLHGEFGLP(0.9881) | 20.48 | 14.92 | 0.945 |
| 2093.011 | COL1A2 | 562 | 582 | GPAGEVGKPGERGLHGEFGLP | GPAGEVGKP[113]GERGLHGEFGLP[113] | GP(0.0199)AGEVGKP(0.9900)GERGLHGEFGLP(0.9901) | 31.25 | 15.91 | 1.000 |
| 2098.877 | COL1A2 | 1099 | 1118 | GPPGVSGGGYDFGYDGDFYR | GP[113]PGVSGGGYDFGYDGDFYR | GP(0.5000)P(0.5000)GVSGGGYDFGYDGDFYR | 18.36 | 18.36 | 0.998 |
| 2100.006 | COL1A2 | 950 | 973 | YPGNIGPVGAAGAPGPHGPVGPAG | YPGN[115]IGPVGAAGAP[113]GPHGPVGP[113]AG | YP(0.0046)GNIGP(0.0047)VGAAGAP(0.8140)GP(0.5192)HGP(0.0886)VGP(0.5689)AG | 27.07 | 27.07 | 1.000 |
| 2109.030 | COL1A2 | 562 | 582 | GPAGEVGKPGERGLHGEFGLP | GP[113]AGEVGKP[113]GERGLHGEFGLP[113] | GP(1.0000)AGEVGKP(1.0000)GERGLHGEFGLP(1.0000) | 20.58 | 14.24 | 0.978 |
| 2154.959 | COL1A2 | 113 | 135 | FQGPAGEPGEPGQTGPAGARGPA | FQGPAGEP[113]GEP[113]GQ[129]TGPAGARGP[113]A | FQGP(0.0220)AGEP(0.9860)GEP(0.9859)GQTGP(0.0202)AGARGP(0.9859)A | 18.69 | 14.52 | 0.979 |
| 2169.916 | COL1A2 | 1099 | 1119 | GPPGVSGGGYDFGYDGDFYRA | GPP[113]GVSGGGYDFGYDGDFYRA | GP(0.0135)P(0.9865)GVSGGGYDFGYDGDFYRA | 26.90 | 26.90 | 1.000 |
| 2170.914 | COL1A2 | 910 | 933 | GAVGSPGVNGAPGEAGRDGNPGND | GAVGSP[113]GVN[115]GAP[113]GEAGRDGNP[113]GND | GAVGSP(1.0000)GVNGAP(1.0000)GEAGRDGNP(1.0000)GND | 24.18 | 10.63 | 1.000 |
| 2175.057 | COL1A2 | 499 | 522 | GPTGDPGKNGDKGHAGLAGARGAP | GPTGDPGKN[115]GDKGHAGLAGARGAP[113] | GP(0.0064)TGDP(0.0069)GKNGDKGHAGLAGARGAP(0.9867) | 21.05 | 12.50 | 1.000 |
| 2180.087 | COL1A2 | 561 | 582 | SGPAGEVGKPGERGLHGEFGLP | SGPAGEVGKP[113]GERGLHGEFGLP[113] | SGP(0.0474)AGEVGKP(0.9762)GERGLHGEFGLP(0.9763) | 18.85 | 13.92 | 0.926 |
| 2191.035 | COL1A2 | 499 | 522 | GPTGDPGKNGDKGHAGLAGARGAP | GPTGDP[113]GKN[115]GDKGHAGLAGARGAP[113] | GP(0.0350)TGDP(0.9826)GKNGDKGHAGLAGARGAP(0.9825) | 19.19 | 14.55 | 0.995 |
| 2207.065 | COL1A2 | 706 | 729 | GPRGSPGERGEVGPAGPNGFAGPA | GPRGSPGERGEVGP[113]AGPNGFAGPA | GP(0.2615)RGSP(0.2489)GERGEVGP(0.4729)AGP(0.0123)NGFAGP(0.0045)A | 33.56 | 31.44 | 1.000 |
| 2207.113 | COL1A2 | 637 | 660 | GPSGLPGERGAAGIPGGKGEKGEP | GPSGLPGERGAAGIP[113]GGKGEKGEP[113] | GP(0.0159)SGLP(0.0148)GERGAAGIP(0.9848)GGKGEKGEP(0.9845) | 26.19 | 17.77 | 0.998 |
| 2223.104 | COL1A2 | 637 | 660 | GPSGLPGERGAAGIPGGKGEKGEP | GPSGLP[113]GERGAAGIP[113]GGKGEKGEP[113] | GP(0.1832)SGLP(0.9385)GERGAAGIP(0.9392)GGKGEKGEP(0.9391) | 22.66 | 18.12 | 0.962 |
| 2225.996 | COL1A2 | 424 | 447 | GASGPAGVRGPNGDAGRPGEPGLM | GASGPAGVRGPN[115]GDAGRP[113]GEP[113]GLM[147] | GASGP(0.0197)AGVRGP(0.0238)NGDAGRP(0.9854)GEP(0.9855)GLM(0.9855) | 27.22 | 19.04 | 1.000 |
| 2231.131 | COL1A2 | 454 | 477 | GSPGNIGPAGKEGPVGLPGIDGRP | GSP[113]GNIGPAGKEGPVGLP[113]GIDGRP | GSP(0.9834)GNIGP(0.0136)AGKEGP(0.0088)VGLP(0.9831)GIDGRP(0.0111) | 25.89 | 21.62 | 0.989 |
| 2232.110 | COL1A2 | 454 | 477 | GSPGNIGPAGKEGPVGLPGIDGRP | GSP[113]GN[115]IGPAGKEGPVGLP[113]GIDGRP | GSP(0.9798)GNIGP(0.0080)AGKEGP(0.0083)VGLP(0.9782)GIDGRP(0.0258) | 29.99 | 23.74 | 1.000 |
| 2247.112 | COL1A2 | 454 | 477 | GSPGNIGPAGKEGPVGLPGIDGRP | GSP[113]GNIGPAGKEGPVGLP[113]GIDGRP[113] | GSP(0.9881)GNIGP(0.0190)AGKEGP(0.0168)VGLP(0.9881)GIDGRP(0.9881) | 28.87 | 19.52 | 1.000 |
| 2248.082 | COL1A2 | 454 | 477 | GSPGNIGPAGKEGPVGLPGIDGRP | GSP[113]GN[115]IGPAGKEGPVGLP[113]GIDGRP[113] | GSP(0.9886)GNIGP(0.0170)AGKEGP(0.0172)VGLP(0.9886)GIDGRP(0.9886) | 24.69 | 17.92 | 1.000 |
| 2256.104 | COL1A2 | 1060 | 1083 | GPAGKDGRTGHPGTVGPAGIRGPQ | GPAGKDGRTGHP[113]GTVGPAGIRGPQ | GP(0.0050)AGKDGRTGHP(0.9829)GTVGP(0.0065)AGIRGP(0.0057)Q | 26.21 | 20.30 | 1.000 |
| 2257.130 | COL1A2 | 1060 | 1083 | GPAGKDGRTGHPGTVGPAGIRGPQ | GPAGKDGRTGHP[113]GTVGPAGIRGPQ[129] | GP(0.0127)AGKDGRTGHP(0.9799)GTVGP(0.0038)AGIRGP(0.0036)Q | 20.08 | 15.15 | 1.000 |
| 2264.130 | COL1A2 | 637 | 661 | GPSGLPGERGAAGIPGGKGEKGEPG | GPSGLPGERGAAGIP[113]GGKGEKGEP[113]G | GP(0.0742)SGLP(0.0196)GERGAAGIP(0.9559)GGKGEKGEP(0.9503)G | 23.82 | 18.86 | 1.000 |
| 2266.133 | COL1A2 | 646 | 669 | GAAGIPGGKGEKGEPGLRGEIGNP | GAAGIP[113]GGKGEKGEP[113]GLRGEIGNP[113] | GAAGIP(1.0000)GGKGEKGEP(1.0000)GLRGEIGNP(1.0000) | 23.07 | 11.32 | 1.000 |
| 2266.154 | COL1A2 | 651 | 673 | PGGKGEKGEPGLRGEIGNPGRDG | P[113]GGKGEKGEP[113]GLRGEIGNPGRDG | P(0.9887)GGKGEKGEP(0.9887)GLRGEIGNP(0.0227)GRDG | 27.60 | 15.64 | 1.000 |
| 2280.125 | COL1A2 | 637 | 661 | GPSGLPGERGAAGIPGGKGEKGEPG | GPSGLP[113]GERGAAGIP[113]GGKGEKGEP[113]G | GP(0.0371)SGLP(0.9877)GERGAAGIP(0.9876)GGKGEKGEP(0.9876)G | 23.30 | 16.31 | 0.994 |
| 2318.160 | COL1A2 | 562 | 585 | GPAGEVGKPGERGLHGEFGLPGPA | GPAGEVGKP[113]GERGLHGEFGLP[113]GPA | GP(0.0281)AGEVGKP(0.9778)GERGLHGEFGLP(0.9783)GP(0.0159)A | 28.19 | 25.63 | 1.000 |
| 2355.173 | COL1A2 | 139 | 162 | GKAGEDGHPGKPGRPGERGVVGPQ | GKAGEDGHPGKPGRP[113]GERGVVGPQ | GKAGEDGHP(0.0038)GKP(0.0050)GRP(0.9840)GERGVVGP(0.0072)Q | 24.18 | 24.18 | 1.000 |
| 2371.172 | COL1A2 | 139 | 162 | GKAGEDGHPGKPGRPGERGVVGPQ | GKAGEDGHP[113]GKPGRP[113]GERGVVGPQ | GKAGEDGHP(0.7631)GKP(0.5979)GRP(0.6255)GERGVVGP(0.0135)Q | 30.10 | 28.14 | 1.000 |
| 2372.159 | COL1A2 | 139 | 162 | GKAGEDGHPGKPGRPGERGVVGPQ | GKAGEDGHP[113]GKPGRP[113]GERGVVGPQ[129] | GKAGEDGHP(0.9753)GKP(0.0370)GRP(0.9744)GERGVVGP(0.0132)Q | 23.68 | 23.68 | 1.000 |
| 2381.098 | COL1A2 | 937 | 960 | GRDGQPGHKGERGYPGNIGPVGAA | GRDGQ[129]P[113]GHKGERGYP[113]GN[115]IGPVGAA | GRDGQP(0.9876)GHKGERGYP(0.9875)GNIGP(0.0249)VGAA | 19.50 | 12.65 | 0.998 |
| 2387.168 | COL1A2 | 139 | 162 | GKAGEDGHPGKPGRPGERGVVGPQ | GKAGEDGHP[113]GKP[113]GRP[113]GERGVVGPQ | GKAGEDGHP(0.9339)GKP(0.9333)GRP(0.9331)GERGVVGP(0.1997)Q | 30.23 | 22.99 | 1.000 |
| 2388.169 | COL1A2 | 139 | 162 | GKAGEDGHPGKPGRPGERGVVGPQ | GKAGEDGHP[113]GKP[113]GRP[113]GERGVVGPQ[129] | GKAGEDGHP(0.8960)GKP(0.8944)GRP(0.8944)GERGVVGP(0.3153)Q | 18.80 | 17.00 | 0.983 |
| 2396.107 | COL1A2 | 344 | 369 | LVGEPGPAGSKGESGNKGEPGSAGPQ | LVGEP[113]GPAGSKGESGNKGEP[113]GSAGPQ | LVGEP(0.7826)GP(0.3862)AGSKGESGNKGEP(0.8220)GSAGP(0.0092)Q | 31.76 | 27.44 | 1.000 |
| 2396.198 | COL1A2 | 652 | 675 | GGKGEKGEPGLRGEIGNPGRDGAR | GGKGEKGEP[113]GLRGEIGNP[113]GRDGAR | GGKGEKGEP(1.0000)GLRGEIGNP(1.0000)GRDGAR | 29.04 | 15.23 | 1.000 |
| 2397.088 | COL1A2 | 344 | 369 | LVGEPGPAGSKGESGNKGEPGSAGPQ | LVGEP[113]GP[113]AGSKGESGN[115]KGEPGSAGPQ | LVGEP(0.7667)GP(0.7275)AGSKGESGNKGEP(0.4838)GSAGP(0.0220)Q | 20.00 | 20.00 | 0.983 |
| 2412.111 | COL1A2 | 344 | 369 | LVGEPGPAGSKGESGNKGEPGSAGPQ | LVGEP[113]GP[113]AGSKGESGNKGEP[113]GSAGPQ | LVGEP(0.9737)GP(0.9737)AGSKGESGNKGEP(0.9742)GSAGP(0.0784)Q | 18.62 | 12.52 | 0.947 |
| 2421.025 | COL1A2 | 910 | 936 | GAVGSPGVNGAPGEAGRDGNPGNDGPP | GAVGSP[113]GVNGAP[113]GEAGRDGNPGNDGP[113]P | GAVGSP(0.8016)GVNGAP(0.8017)GEAGRDGNP(0.0243)GNDGP(0.6861)P(0.6861) | 26.78 | 26.78 | 1.000 |
| 2422.033 | COL1A2 | 910 | 936 | GAVGSPGVNGAPGEAGRDGNPGNDGPP | GAVGSP[113]GVN[115]GAP[113]GEAGRDGNPGNDGPP[113] | GAVGSP(0.9874)GVNGAP(0.9874)GEAGRDGNP(0.0197)GNDGP(0.0182)P(0.9873) | 53.62 | 53.62 | 1.000 |
| 2423.033 | COL1A2 | 910 | 936 | GAVGSPGVNGAPGEAGRDGNPGNDGPP | GAVGSP[113]GVN[115]GAP[113]GEAGRDGNPGN[115]DGP[113]P | GAVGSP(0.8049)GVNGAP(0.8049)GEAGRDGNP(0.0139)GNDGP(0.6881)P(0.6881) | 38.64 | 38.64 | 1.000 |
| 2437.041 | COL1A2 | 910 | 936 | GAVGSPGVNGAPGEAGRDGNPGNDGPP | GAVGSP[113]GVNGAP[113]GEAGRDGNP[113]GNDGPP[113] | GAVGSP(0.9827)GVNGAP(0.9827)GEAGRDGNP(0.9827)GNDGP(0.0692)P(0.9827) | 25.93 | 25.93 | 1.000 |
| 2438.015 | COL1A2 | 910 | 936 | GAVGSPGVNGAPGEAGRDGNPGNDGPP | GAVGSP[113]GVN[115]GAPGEAGRDGNP[113]GNDGP[113]P[113] | GAVGSP(0.8385)GVNGAP(0.6731)GEAGRDGNP(0.8351)GNDGP(0.8267)P(0.8267) | 24.14 | 24.14 | 1.000 |
| 2453.148 | COL1A2 | 343 | 369 | GLVGEPGPAGSKGESGNKGEPGSAGPQ | GLVGEP[113]GPAGSKGESGNKGEP[113]GSAGPQ | GLVGEP(0.9109)GP(0.1587)AGSKGESGNKGEP(0.9187)GSAGP(0.0118)Q | 30.26 | 27.93 | 1.000 |
| 2454.119 | COL1A2 | 343 | 369 | GLVGEPGPAGSKGESGNKGEPGSAGPQ | GLVGEP[113]GPAGSKGESGN[115]KGEP[113]GSAGPQ | GLVGEP(0.9770)GP(0.0338)AGSKGESGNKGEP(0.9772)GSAGP(0.0120)Q | 29.34 | 27.21 | 1.000 |
| 2459.093 | COL1A2 | 772 | 798 | GPAGSRGDGGPPGMTGFPGAAGRTGPP | GPAGSRGDGGPP[113]GM[147]TGFP[113]GAAGRTGP[113]P[113] | GP(0.1304)AGSRGDGGP(0.3347)P(0.9049)GM(0.9049)TGFP(0.9085)GAAGRTGP(0.9084)P(0.908 | 18.69 | 18.69 | 0.994 |

| | | | | | | | | | |
|---|---|---|---|---|---|---|---|---|---|
| 2469.140 | COL1A2 | 343 | 369 | GLVGEPGPAGSKGESGNKGEPGSAGPQ | GLVGEP[113]GP[113]AGSKGESGNKGEP[113]GSAGPQ | GLVGEP(0.9878)GP(0.9878)AGSKGESGNKGEP(0.9881)GSAGP(0.0362)Q | 19.30 | 12.20 | 0.976 |
| 2473.265 | COL1A2 | 877 | 903 | GSRGERGLPGVAGAVGEPGPLGIAGPP | GSRGERGLP[113]GVAGAVGEP[113]GPLGIAGP[113]P | GSRGERGLP(0.7900)GVAGAVGEP(0.7889)GP(0.0697)LGIAGP(0.6757)P(0.6757) | 28.22 | 28.22 | 1.000 |
| 2477.105 | COL1A2 | 109 | 135 | GPQGFQGPAGEPGEPGQTGPAGARGPA | GPQGFQGPAGEP[113]GEP[113]GQTGPAGARGPA | GP(0.0187)QGFQGP(0.0113)AGEP(0.9769)GEP(0.9776)GQTGP(0.0066)AGARGP(0.0088)A | 31.37 | 29.22 | 1.000 |
| 2493.105 | COL1A2 | 109 | 135 | GPQGFQGPAGEPGEPGQTGPAGARGPA | GPQGFQGPAGEP[113]GEP[113]GQTGPAGARGP[113]A | GP(0.0209)QGFQGP(0.0142)AGEP(0.9808)GEP(0.9802)GQTGP(0.0235)AGARGP(0.9804)A | 30.82 | 26.48 | 1.000 |
| 2494.112 | COL1A2 | 109 | 135 | GPQGFQGPAGEPGEPGQTGPAGARGPA | GPQGFQGPAGEP[113]GEP[113]GQ[129]TGPAGARGP[113]A | GP(0.0151)QGFQGP(0.0127)AGEP(0.9810)GEP(0.9808)GQTGP(0.0301)AGARGP(0.9803)A | 22.40 | 16.95 | 0.971 |
| 2509.194 | COL1A2 | 421 | 447 | GSRGASGPAGVRGPNGDAGRPGEPGLM | GSRGASGPAGVRGPNGDAGRP[113]GEP[113]GLM | GSRGASGP(0.3010)AGVRGP(0.1787)NGDAGRP(0.7447)GEP(0.7664)GLM(0.0092) | 26.63 | 22.19 | 1.000 |
| 2525.181 | COL1A2 | 421 | 447 | GSRGASGPAGVRGPNGDAGRPGEPGLM | GSRGASGPAGVRGPNGDAGRP[113]GEP[113]GLM[147] | GSRGASGP(0.6596)AGVRGP(0.1260)NGDAGRP(0.6659)GEP(0.7735)GLM(0.7750) | 19.86 | 19.07 | 0.995 |
| 2592.305 | COL1A2 | 643 | 669 | GERGAAGIPGGKGEKGEPGLRGEIGNP | GERGAAGIP[113]GGKGEKGEPGLRGEIGNP[113] | GERGAAGIP(0.9865)GGKGEKGEP(0.0271)GLRGEIGNP(0.9865) | 26.93 | 21.17 | 1.000 |
| 2608.278 | COL1A2 | 643 | 669 | GERGAAGIPGGKGEKGEPGLRGEIGNP | GERGAAGIP[113]GGKGEKGEP[113]GLRGEIGNP[113] | GERGAAGIP(1.0000)GGKGEKGEP(1.0000)GLRGEIGNP(1.0000) | 26.86 | 14.79 | 1.000 |
| 2622.320 | COL1A2 | 136 | 162 | GPPGKAGEDGHPGKPGRPGERGVVGPQ | GP[113]PGKAGEDGHPGKPGRP[113]GERGVVGPQ | GP(0.8867)P(0.0093)GKAGEDGHP(0.0050)GKP(0.0067)GRP(0.8006)GERGVVGP(0.2917)Q | 20.92 | 19.40 | 1.000 |
| 2638.296 | COL1A2 | 136 | 162 | GPPGKAGEDGHPGKPGRPGERGVVGPQ | GPP[113]GKAGEDGHP[113]GKPGRP[113]GERGVVGPQ | GP(0.0513)P(0.8400)GKAGEDGHP(0.8412)GKP(0.0131)GRP(0.7842)GERGVVGP(0.4702)Q | 24.43 | 24.43 | 1.000 |
| 2654.298 | COL1A2 | 136 | 162 | GPPGKAGEDGHPGKPGRPGERGVVGPQ | GPP[113]GKAGEDGHP[113]GKP[113]GRP[113]GERGVVGPQ | GP(0.0225)P(0.9843)GKAGEDGHP(0.9843)GKP(0.9842)GRP(0.9841)GERGVVGP(0.0406)Q | 18.06 | 18.06 | 1.000 |
| 2728.440 | COL1A2 | 168 | 195 | PGTPGLPGFKGIRGHNGLDGLKGQPGAP | P[113]GTPGLPGFKGIRGHNGLDGLKGQ[129]P[113]GAP | P(0.8045)GTP(0.0146)GLP(0.0146)GFKGIRGHNGLDGLKGQP(0.5832)GAP(0.5832) | 20.17 | 20.17 | 0.983 |
| 2803.401 | COL1A2 | 1062 | 1091 | AGKDGRTGHPGTVGPAGIRGPQGHQGPAGP | AGKDGRTGHPGTVGP[113]AGIRGPQGHQGPAGP | AGKDGRTGHP(0.0122)GTVGP(0.9301)AGIRGP(0.0192)QGHQGP(0.0214)AGP(0.0171) | 18.28 | 17.81 | 0.972 |
| 2878.476 | COL1A2 | 646 | 675 | GAAGIPGGKGEKGEPGLRGEIGNPGRDGAR | GAAGIP[113]GGKGEKGEP[113]GLRGEIGNP[113]GRDGAR | GAAGIP(1.0000)GGKGEKGEP(1.0000)GLRGEIGNP(1.0000)GRDGAR | 24.17 | 11.15 | 1.000 |

| M+H | Gene | Domain Start | Domain End | Peptide | Modified Peptide [113] means HYP | Sequence with HYP probability PM:15.9949 | Hyperscore | Nextscore | Peptide Prophet Probability |
|---|---|---|---|---|---|---|---|---|---|
| 741.452 | COL3A1 | 946 | 953 | PLGIAGIT | PLGIAGIT | | 18.07 | 13.03 | 0.984 |
| 743.370 | COL3A1 | 702 | 710 | GPEGGKGAA | GPEGGKGAA | | 19.36 | 12.34 | 0.992 |
| 762.340 | COL3A1 | 867 | 875 | GGPGAAGFP | GGP[113]GAAGFP[113] | GGP(1.0000)GAAGFP(1.0000) | 19.14 | 13.16 | 0.966 |
| 774.344 | COL3A1 | 1215 | 1221 | GFAPYYG | GFAPYYG | | 18.41 | 11.17 | 0.999 |
| 788.377 | COL3A1 | 1002 | 1010 | GLAGTAGEP | GLAGTAGEP[113] | GLAGTAGEP(1.0000) | 19.37 | 12.39 | 0.987 |
| 798.467 | COL3A1 | 945 | 953 | GPLGIAGIT | GPLGIAGIT | | 21.90 | 16.36 | 0.960 |
| 806.438 | COL3A1 | 164 | 172 | VAVGGLAGY | VAVGGLAGY | | 19.21 | 11.33 | 1.000 |
| 815.428 | COL3A1 | 639 | 647 | GLQGLPGTG | GLQGLP[113]GTG | GLQGLP(1.0000)GTG | 18.05 | 14.96 | 0.964 |
| 830.396 | COL3A1 | 1125 | 1133 | GQQGAIGSP | GQQGAIGSP[113] | GQQGAIGSP(1.0000) | 21.06 | 14.63 | 1.000 |
| 832.381 | COL3A1 | 336 | 344 | GPPGTAGFP | GPP[113]GTAGFP[113] | GP(0.0309)P(0.9845)GTAGFP(0.9846) | 22.18 | 21.72 | 0.993 |
| 844.421 | COL3A1 | 252 | 260 | GPAGIPGFP | GPAGIP[113]GFP[113] | GP(0.0208)AGIP(0.9896)GFP(0.9896) | 20.63 | 20.63 | 0.922 |
| 845.400 | COL3A1 | 1002 | 1011 | GLAGTAGEPG | GLAGTAGEP[113]G | GLAGTAGEP(1.0000)G | 19.48 | 10.80 | 0.993 |
| 855.489 | COL3A1 | 945 | 954 | GPLGIAGITG | GPLGIAGITG | | 22.78 | 15.51 | 0.956 |
| 858.436 | COL3A1 | 972 | 980 | GPQGVKGES | GPQGVKGES | | 18.74 | 10.03 | 0.997 |
| 891.384 | COL3A1 | 195 | 203 | GSPGYQGPP | GSP[113]GYQGPP[113] | GSP(0.9879)GYQGP(0.0223)P(0.9898) | 19.25 | 17.43 | 0.994 |
| 914.384 | COL3A1 | 644 | 653 | PGTGGPPGEN | P[113]GTGGPP[113]GEN | P(0.9886)GTGGP(0.0228)P(0.9886)GEN | 21.08 | 17.29 | 0.996 |
| 919.467 | COL3A1 | 164 | 173 | VAVGGLAGYP | VAVGGLAGYP[113] | VAVGGLAGYP(1.0000) | 24.34 | 11.63 | 1.000 |
| 921.412 | COL3A1 | 573 | 581 | GQPGVMGFP | GQPGVM[147]GFP[113] | GQP(0.0493)GVM(0.9752)GFP(0.9755) | 18.50 | 13.65 | 1.000 |
| 923.446 | COL3A1 | 840 | 851 | GVAGPPGGSGPA | GVAGPPGGSGPA | | 19.78 | 10.04 | 0.983 |
| 926.524 | COL3A1 | 945 | 955 | GPLGIAGITGA | GPLGIAGITGA | | 25.32 | 19.37 | 1.000 |
| 937.401 | COL3A1 | 573 | 581 | GQPGVMGFP | GQP[113]GVM[147]GFP[113] | GQP(1.0000)GVM(1.0000)GFP(1.0000) | 22.22 | 11.69 | 1.000 |
| 976.506 | COL3A1 | 163 | 173 | GVAVGGLAGYP | GVAVGGLAGYP[113] | GVAVGGLAGYP(1.0000) | 23.02 | 17.83 | 0.983 |
| 985.579 | COL3A1 | 947 | 957 | LGIAGITGARG | LGIAGITGARG | | 21.82 | 14.55 | 0.998 |
| 999.591 | COL3A1 | 949 | 959 | IAGITGARGLA | IAGITGARGLA | | 19.73 | 10.11 | 0.981 |
| 1019.437 | COL3A1 | 864 | 875 | GSPGGPGAAGFP | GSP[113]GGP[113]GAAGFP[113] | GSP(1.0000)GGP(1.0000)GAAGFP(1.0000) | 22.02 | 14.34 | 0.987 |
| 1025.518 | COL3A1 | 640 | 650 | LQGLPGTGGPP | LQGLP[113]GTGGPP[113] | LQGLP(0.9884)GTGGP(0.0232)P(0.9884) | 19.31 | 17.23 | 0.995 |
| 1025.610 | COL3A1 | 946 | 956 | PLGIAGITGAR | PLGIAGITGAR | | 25.85 | 11.85 | 0.998 |
| 1027.499 | COL3A1 | 780 | 791 | GEGGAPGLPGIA | GEGGAP[113]GLP[113]GIA | GEGGAP(1.0000)GLP(1.0000)GIA | 19.12 | 11.64 | 0.971 |
| 1042.502 | COL3A1 | 636 | 646 | GPQGLQGLPGT | GPQ[129]GLQ[129]GLP[113]GT | GP(0.0115)QGLQGLP(0.9885)GT | 20.38 | 11.81 | 1.000 |
| 1055.516 | COL3A1 | 1122 | 1133 | GPAGQQGAIGSP | GPAGQQGAIGSP[113] | GP(0.0113)AGQQGAIGSP(0.9887) | 18.12 | 11.84 | 0.991 |
| 1056.491 | COL3A1 | 1122 | 1133 | GPAGQQGAIGSP | GPAGQ[129]QGAIGSP[113] | GP(0.0111)AGQQGAIGSP(0.9889) | 18.04 | 16.02 | 0.994 |
| 1056.617 | COL3A1 | 948 | 959 | GIAGITGARGLA | GIAGITGARGLA | | 18.56 | 9.59 | 0.996 |
| 1059.505 | COL3A1 | 1003 | 1013 | LAGTAGEPGRD | LAGTAGEP[113]GRD | LAGTAGEP(1.0000)GRD | 20.11 | 9.86 | 1.000 |
| 1063.537 | COL3A1 | 162 | 173 | SGVAVGGLAGYP | SGVAVGGLAGYP[113] | SGVAVGGLAGYP(1.0000) | 29.31 | 13.55 | 1.000 |
| 1070.471 | COL3A1 | 624 | 635 | GPGGDKGDTGPP | GPGGDKGDTGPP[113] | GP(0.0056)GGDKGDTGP(0.0064)P(0.9880) | 18.66 | 18.66 | 0.979 |
| 1073.486 | COL3A1 | 336 | 347 | GPPGTAGFPGSP | GP[113]PGTAGFPGSP[113] | GP(0.6149)P(0.6149)GTAGFP(0.0099)GSP(0.7602) | 21.88 | 21.88 | 1.000 |
| 1082.531 | COL3A1 | 639 | 650 | GLQGLPGTGGPP | GLQGLP[113]GTGGPP[113] | GLQGLP(0.9891)GTGGP(0.0218)P(0.9891) | 30.12 | 27.94 | 1.000 |
| 1082.617 | COL3A1 | 945 | 956 | GPLGIAGITGAR | GPLGIAGITGAR | | 32.53 | 17.65 | 1.000 |
| 1083.522 | COL3A1 | 639 | 650 | GLQGLPGTGGPP | GLQ[129]GLP[113]GTGGP[113]P | GLQGLP(0.7197)GTGGP(0.6401)P(0.6401) | 18.33 | 18.33 | 0.963 |
| 1084.486 | COL3A1 | 894 | 905 | GPSGSPGKDGPP | GPSGSP[113]GKDGP[113]P | GP(0.0087)SGSP(0.7649)GKDGP(0.6132)P(0.6132) | 24.71 | 24.71 | 0.961 |
| 1085.600 | COL3A1 | 250 | 260 | IKGPAGIPGFP | IKGPAGIP[113]GFP[113] | IKGP(0.0198)AGIP(0.9901)GFP(0.9901) | 18.57 | 12.65 | 0.988 |
| 1088.502 | COL3A1 | 717 | 728 | GAAGTPGLQGMP | GAAGTP[113]GLQGMP[113] | GAAGTP(0.9885)GLQGM(0.0228)P(0.9887) | 18.81 | 16.58 | 0.993 |
| 1089.471 | COL3A1 | 336 | 347 | GPPGTAGFPGSP | GPP[113]GTAGFP[113]GSP[113] | GP(0.0397)P(0.9868)GTAGFP(0.9868)GSP(0.9868) | 22.10 | 20.09 | 0.976 |
| 1097.555 | COL3A1 | 636 | 647 | GPQGLQGLPGTG | GPQGLQGLP[113]GTG | GP(0.0103)QGLQGLP(0.9897)GTG | 21.69 | 9.13 | 1.000 |
| 1098.535 | COL3A1 | 636 | 647 | GPQGLQGLPGTG | GPQGLQ[129]GLP[113]GTG | GP(0.0112)QGLQGLP(0.9888)GTG | 20.50 | 13.14 | 1.000 |
| 1101.468 | COL3A1 | 468 | 479 | GSPGEPGANGLP | GSP[113]GEP[113]GAN[115]GLP[113] | GSP(1.0000)GEP(1.0000)GANGLP(1.0000) | 19.37 | 12.41 | 0.996 |
| 1101.587 | COL3A1 | 250 | 260 | IKGPAGIPGFP | IKGP[113]AGIP[113]GFP[113] | IKGP(1.0000)AGIP(1.0000)GFP(1.0000) | 18.11 | 11.82 | 0.974 |
| 1106.521 | COL3A1 | 1146 | 1157 | GPPGKDGTSGHP | GPPGKDGTSGHP | | 18.57 | 12.13 | 0.965 |
| 1109.563 | COL3A1 | 495 | 506 | GPNGIPGEKGPA | GPNGIP[113]GEKGPA | GP(0.0050)NGIP(0.9865)GEKGP(0.0085)A | 19.22 | 12.62 | 0.985 |
| 1110.540 | COL3A1 | 495 | 506 | GPNGIPGEKGPA | GPN[115]GIP[113]GEKGPA | GP(0.0056)NGIP(0.9888)GEKGP(0.0056)A | 20.14 | 13.38 | 0.965 |
| 1116.525 | COL3A1 | 1002 | 1013 | GLAGTAGEPGRD | GLAGTAGEP[113]GRD | GLAGTAGEP(1.0000)GRD | 24.38 | 12.73 | 0.995 |
| 1117.520 | COL3A1 | 1059 | 1070 | GKSGDRGESGPA | GKSGDRGESGPA | | 18.63 | 11.75 | 0.959 |
| 1125.566 | COL3A1 | 750 | 761 | GADGVPGKDGPR | GADGVPGKDGPR | | 21.60 | 12.59 | 0.999 |
| 1127.541 | COL3A1 | 225 | 236 | GPAGKDGESGRP | GPAGKDGESGRP | | 19.80 | 11.22 | 1.000 |
| 1128.520 | COL3A1 | 984 | 995 | GANGLSGERGPP | GAN[115]GLSGERGPP[113] | GANGLSGERGP(0.0489)P(0.9511) | 25.69 | 23.84 | 0.997 |
| 1138.556 | COL3A1 | 1149 | 1160 | GKDGTSGHPGPI | GKDGTSGHP[113]GPI | GKDGTSGHP(0.9841)GP(0.0159)I | 18.74 | 15.59 | 0.990 |
| 1139.636 | COL3A1 | 945 | 957 | GPLGIAGITGARG | GPLGIAGITGARG | | 32.60 | 8.57 | 1.000 |
| 1142.618 | COL3A1 | 249 | 260 | GIKGPAGIPGFP | GIKGPAGIP[113]GFP[113] | GIKGP(0.0204)AGIP(0.9898)GFP(0.9898) | 20.63 | 9.50 | 1.000 |
| 1143.545 | COL3A1 | 225 | 236 | GPAGKDGESGRP | GPAGKDGESGRP[113] | GP(0.0100)AGKDGESGRP(0.9900) | 18.38 | 9.57 | 0.998 |
| 1154.537 | COL3A1 | 606 | 617 | GPPGKNGETGPQ | GPP[113]GKNGETGPQ | GP(0.0051)P(0.9899)GKNGETGP(0.0051)Q | 20.48 | 17.29 | 0.968 |
| 1155.524 | COL3A1 | 606 | 617 | GPPGKNGETGPQ | GPP[113]GKN[115]GETGPQ | GP(0.0058)P(0.9887)GKNGETGP(0.0055)Q | 24.20 | 23.00 | 0.993 |
| 1156.588 | COL3A1 | 972 | 983 | GPQGVKGESGKP | GPQGVKGESGKP[113] | GP(0.0103)QGVKGESGKP(0.9897) | 24.61 | 9.97 | 1.000 |
| 1158.614 | COL3A1 | 249 | 260 | GIKGPAGIPGFP | GIKGP[113]AGIP[113]GFP[113] | GIKGP(1.0000)AGIP(1.0000)GFP(1.0000) | 18.07 | 12.86 | 0.980 |
| 1161.571 | COL3A1 | 480 | 491 | GAAGERGAPGFR | GAAGERGAP[113]GFR | GAAGERGAP(1.0000)GFR | 25.22 | 8.34 | 1.000 |
| 1162.512 | COL3A1 | 726 | 737 | GMPGERGGLGSP | GM[147]P[113]GERGGLGSP[113] | GM(1.0000)P(1.0000)GERGGLGSP(1.0000) | 18.22 | 9.17 | 0.998 |
| 1169.703 | COL3A1 | 947 | 959 | LGIAGITGARGLA | LGIAGITGARGLA | | 20.84 | 9.72 | 0.999 |
| 1172.510 | COL3A1 | 795 | 806 | GSPGERGETGPP | GSP[113]GERGETGPP[113] | GSP(0.9874)GERGETGP(0.0253)P(0.9874) | 18.76 | 18.76 | 0.947 |
| 1173.544 | COL3A1 | 1002 | 1014 | GLAGTAGEPGRDG | GLAGTAGEP[113]GRDG | GLAGTAGEP(1.0000)GRDG | 21.27 | 10.74 | 0.992 |
| 1184.602 | COL3A1 | 855 | 866 | GPQGVKGERGSP | GPQGVKGERGSP[113] | GP(0.0100)QGVKGERGSP(0.9900) | 18.45 | 6.76 | 1.000 |
| 1231.548 | COL3A1 | 1003 | 1015 | LAGTAGEPGRDGN | LAGTAGEP[113]GRDGN[115] | LAGTAGEP(1.0000)GRDGN | 22.99 | 10.61 | 1.000 |
| 1232.573 | COL3A1 | 570 | 581 | GPRGQPGVMGFP | GPRGQ[129]PGVM[147]GFP[113] | GP(0.0119)RGQP(0.0119)GVM(0.9881)GFP(0.9881) | 18.19 | 13.97 | 0.983 |
| 1248.567 | COL3A1 | 570 | 581 | GPRGQPGVMGFP | GPRGQ[129]P[113]GVM[147]GFP[113] | GP(0.0367)RGQP(0.9877)GVM(0.9878)GFP(0.9878) | 21.43 | 14.99 | 0.998 |
| 1252.732 | COL3A1 | 945 | 958 | GPLGIAGITGARGL | GPLGIAGITGARGL | | 21.27 | 9.83 | 1.000 |
| 1266.752 | COL3A1 | 946 | 959 | PLGIAGITGARGLA | PLGIAGITGARGLA | | 28.85 | 13.45 | 1.000 |
| 1287.586 | COL3A1 | 1002 | 1015 | GLAGTAGEPGRDGN | GLAGTAGEP[113]GRDGN | GLAGTAGEP(1.0000)GRDGN | 19.35 | 10.02 | 0.993 |
| 1288.573 | COL3A1 | 1002 | 1015 | GLAGTAGEPGRDGN | GLAGTAGEP[113]GRDGN[115] | GLAGTAGEP(1.0000)GRDGN | 25.28 | 11.94 | 1.000 |
| 1295.582 | COL3A1 | 771 | 785 | GPAGQPGDKGEGGAP | GPAGQ[129]PGDKGEGGAP | | 22.17 | 15.21 | 0.937 |
| 1311.658 | COL3A1 | 777 | 791 | GDKGEGGAPGLPGIA | GDKGEGGAPGLP[113]GIA | GDKGEGGAP(0.0143)GLP(0.9857)GIA | 22.95 | 18.09 | 0.996 |
| 1323.756 | COL3A1 | 945 | 959 | GPLGIAGITGARGLA | GPLGIAGITGARGLA | | 42.31 | 20.20 | 1.000 |
| 1325.588 | COL3A1 | 621 | 635 | GPTGPGGDKGDTGPP | GPTGPGGDKGDTGPP[113] | GP(0.0052)TGP(0.0053)GGDKGDTGP(0.0036)P(0.9859) | 22.42 | 22.42 | 0.996 |

| | | | | | | | | | |
|---|---|---|---|---|---|---|---|---|---|
| 1325.618 | COL3A1 | 640 | 653 | LQGLPGTGGPPGEN | LQGLP[113]GTGGPP[113]GEN | LQGLP(0.9127)GTGGP(0.1765)P(0.9107)GEN | 18.59 | 18.50 | 0.953 |
| 1327.568 | COL3A1 | 540 | 554 | GSPGGPGSDGKPGPP | GSP[113]GGP[113]GSDGKP[113]GP[113]P | GSP(0.8205)GGP(0.8204)GSDGKP(0.8161)GP(0.7715)P(0.7715) | 21.32 | 21.32 | 0.987 |
| 1327.636 | COL3A1 | 393 | 407 | GGKGEMGPAGIPGAP | GGKGEMGPAGIP[113]GAP[113] | GGKGEM(0.0132)GP(0.0137)AGIP(0.9865)GAP(0.9865) | 24.68 | 17.34 | 0.979 |
| 1327.652 | COL3A1 | 777 | 791 | GDKGEGGAPGLPGIA | GDKGEGGAP[113]GLP[113]GIA | GDKGEGGAP(1.0000)GLP(1.0000)GIA | 21.62 | 14.63 | 0.989 |
| 1329.663 | COL3A1 | 481 | 494 | AAGERGAPGFRGPA | AAGERGAP[113]GFRGPA | AAGERGAP(0.9761)GFRGP(0.0239)A | 18.20 | 13.87 | 1.000 |
| 1336.646 | COL3A1 | 747 | 761 | GGPGADGVPGKDGPR | GGPGADGVPGKDGPR | | 24.92 | 8.06 | 1.000 |
| 1341.599 | COL3A1 | 519 | 533 | GAAGEPGRDGVPGGP | GAAGEP[113]GRDGVP[113]GGP[113] | GAAGEP(1.0000)GRDGVP(1.0000)GGP(1.0000) | 19.94 | 13.16 | 0.973 |
| 1342.618 | COL3A1 | 915 | 929 | GSPGVSGPKGDAGQP | GSP[113]GVSGPKGDAGQP[113] | GSP(0.9895)GVSGP(0.0211)KGDAGQP(0.9895) | 23.28 | 13.72 | 1.000 |
| 1343.605 | COL3A1 | 1003 | 1016 | LAGTAGEPGRDGNP | LAGTAGEP[113]GRDGNP[113] | LAGTAGEP(1.0000)GRDGNP(1.0000) | 28.66 | 13.17 | 1.000 |
| 1343.628 | COL3A1 | 393 | 407 | GGKGEMGPAGIPGAP | GGKGEM[147]GPAGIP[113]GAP[113] | GGKGEM(0.9789)GP(0.0630)AGIP(0.9790)GAP(0.9790) | 22.52 | 17.08 | 0.993 |
| 1352.640 | COL3A1 | 747 | 761 | GGPGADGVPGKDGPR | GGP[113]GADGVPGKDGPR | GGP(0.9884)GADGVP(0.0059)GKDGP(0.0057)R | 20.60 | 11.10 | 1.000 |
| 1354.630 | COL3A1 | 672 | 686 | GGKGDAGAPGERGPP | GGKGDAGAP[113]GERGP[113]P | GGKGDAGAP(0.7195)GERGP(0.6403)P(0.6403) | 19.44 | 19.44 | 0.950 |
| 1358.622 | COL3A1 | 429 | 443 | GGAGEPGKNGAKGEP | GGAGEP[113]GKN[115]GAKGEP[113] | GGAGEP(1.0000)GKNGAKGEP(1.0000) | 18.20 | 13.66 | 0.971 |
| 1361.590 | COL3A1 | 387 | 401 | GINGSPGGKGEMGPA | GIN[115]GSP[113]GGKGEM[147]GPA | GINGSP(0.9890)GGKGEM(0.9890)GP(0.0220)A | 24.40 | 18.79 | 0.985 |
| 1361.601 | COL3A1 | 861 | 875 | GERGSPGGPGAAGFP | GERGSP[113]GGP[113]GAAGFP[113] | GERGSP(1.0000)GGP(1.0000)GAAGFP(1.0000) | 23.42 | 15.01 | 0.995 |
| 1364.674 | COL3A1 | 636 | 650 | GPQGLQGLPGTGGPP | GPQGLQGLP[113]GTGGP[113]P | GP(0.0099)QGLQGLP(0.7648)GTGGP(0.6126)P(0.6126) | 23.96 | 21.80 | 1.000 |
| 1365.664 | COL3A1 | 636 | 650 | GPQGLQGLPGTGGPP | GPQGLQ[129]GLP[113]GTGGPP[113] | GP(0.0129)QGLQGLP(0.9720)GTGGP(0.0444)P(0.9706) | 27.16 | 25.05 | 0.996 |
| 1371.613 | COL3A1 | 714 | 728 | GPPGAAGTPGLQGMP | GP[113]PGAAGTP[113]GLQGM[147]P[113] | GP(0.8600)P(0.5465)GAAGTP(0.8646)GLQGM(0.8645)P(0.8645) | 19.83 | 19.83 | 0.994 |
| 1382.643 | COL3A1 | 639 | 653 | GLQGLPGTGGPPGEN | GLQGLP[113]GTGGPP[113]GEN | GLQGLP(0.9843)GTGGP(0.0315)P(0.9842)GEN | 24.02 | 20.15 | 1.000 |
| 1384.635 | COL3A1 | 1002 | 1016 | GLAGTAGEPGRDGNP | GLAGTAGEP[113]GRDGNP | GLAGTAGEP(0.9887)GRDGNP(0.0113) | 24.31 | 15.97 | 0.999 |
| 1385.645 | COL3A1 | 283 | 296 | LKGENGLPGENGAP | LKGEN[115]GLP[113]GENGAP[113] | LKGENGLP(1.0000)GENGAP(1.0000) | 19.07 | 12.00 | 0.980 |
| 1386.627 | COL3A1 | 283 | 296 | LKGENGLPGENGAP | LKGEN[115]GLP[113]GEN[115]GAP[113] | LKGENGLP(1.0000)GENGAP(1.0000) | 20.42 | 12.64 | 0.991 |
| 1386.681 | COL3A1 | 480 | 494 | GAAGERGAPGFRGPA | GAAGERGAP[113]GFRGPA | GAAGERGAP(0.9853)GFRGP(0.0147)A | 18.22 | 15.72 | 0.990 |
| 1400.630 | COL3A1 | 1002 | 1016 | GLAGTAGEPGRDGNP | GLAGTAGEP[113]GRDGNP[113] | GLAGTAGEP(1.0000)GRDGNP(1.0000) | 37.77 | 13.29 | 1.000 |
| 1401.611 | COL3A1 | 465 | 479 | GKDGSPGEPGANGLP | GKDGSP[113]GEP[113]GAN[115]GLP[113] | GKDGSP(1.0000)GEP(1.0000)GANGLP(1.0000) | 19.19 | 12.06 | 0.991 |
| 1403.601 | COL3A1 | 384 | 398 | GPPGINGSPGGKGEM | GPP[113]GIN[115]GSP[113]GGKGEM[147] | GP(0.0325)P(0.9892)GINGSP(0.9892)GGKGEM(0.9892) | 20.98 | 18.70 | 0.964 |
| 1403.658 | COL3A1 | 724 | 737 | LQGMPGERGGLGSP | LQGM[147]P[113]GERGGLGSP[113] | LQGM(1.0000)P(1.0000)GERGGLGSP(1.0000) | 19.86 | 13.51 | 0.977 |
| 1409.740 | COL3A1 | 246 | 260 | GPPGIKGPAGIPGFP | GPP[113]GIKGPAGIP[113]GFP[113] | GP(0.3884)P(0.8539)GIKGP(0.0193)AGIP(0.8692)GFP(0.8692) | 21.82 | 20.65 | 0.983 |
| 1410.689 | COL3A1 | 981 | 995 | GKPGANGLSGERGPP | GKPGAN[115]GLSGERGP[113]P | GKP(0.0041)GANGLSGERGP(0.4979)P(0.4979) | 19.93 | 19.93 | 0.969 |
| 1413.663 | COL3A1 | 921 | 935 | GPKGDAGQPGEKGSP | GPKGDAGQP[113]GEKGSP[113] | GP(0.0194)KGDAGQP(0.9903)GEKGSP(0.9903) | 19.80 | 11.68 | 0.994 |
| 1425.735 | COL3A1 | 246 | 260 | GPPGIKGPAGIPGFP | GP[113]P[113]GIKGPAGIP[113]GFP[113] | GP(0.8174)P(0.8244)GIKGP(0.6961)AGIP(0.8310)GFP(0.8310) | 24.61 | 22.91 | 1.000 |
| 1426.679 | COL3A1 | 282 | 296 | GLKGENGLPGENGAP | GLKGENGLPGEN[115]GAP[113] | GLKGENGLP(0.0197)GENGAP(0.9803) | 19.81 | 13.08 | 0.989 |
| 1426.680 | COL3A1 | 981 | 995 | GKPGANGLSGERGPP | GKP[113]GAN[115]GLSGERGP[113]P | GKP(0.7186)GANGLSGERGP(0.6407)P(0.6407) | 21.45 | 21.45 | 1.000 |
| 1427.664 | COL3A1 | 282 | 296 | GLKGENGLPGENGAP | GLKGEN[115]GLPGEN[115]GAP[113] | GLKGENGLP(0.0195)GENGAP(0.9805) | 27.86 | 15.15 | 1.000 |
| 1441.660 | COL3A1 | 1002 | 1017 | GLAGTAGEPGRDGNPG | GLAGTAGEP[113]GRDGNPG | GLAGTAGEP(0.9890)GRDGNP(0.0110)G | 24.53 | 20.40 | 1.000 |
| 1441.690 | COL3A1 | 282 | 296 | GLKGENGLPGENGAP | GLKGENGLP[113]GENGAP[113] | GLKGENGLP(1.0000)GENGAP(1.0000) | 21.29 | 13.36 | 0.999 |
| 1442.670 | COL3A1 | 282 | 296 | GLKGENGLPGENGAP | GLKGENGLP[113]GEN[115]GAP[113] | GLKGENGLP(1.0000)GENGAP(1.0000) | 18.61 | 10.63 | 0.993 |
| 1443.654 | COL3A1 | 282 | 296 | GLKGENGLPGENGAP | GLKGEN[115]GLP[113]GEN[115]GAP[113] | GLKGENGLP(1.0000)GENGAP(1.0000) | 18.07 | 10.73 | 0.999 |
| 1444.678 | COL3A1 | 723 | 737 | GLQGMPGERGGLGSP | GLQGM[147]PGERGGLGSP[113] | GLQGM(0.9790)P(0.0422)GERGGLGSP(0.9788) | 26.72 | 21.31 | 0.966 |
| 1446.601 | COL3A1 | 195 | 209 | GSPGYQGPPGEPGQA | GSP[113]GYQGPP[113]GEP[113]GQA | GSP(0.9461)GYQGP(0.1617)P(0.9458)GEP(0.9465)GQA | 26.46 | 24.77 | 0.987 |
| 1455.727 | COL3A1 | 972 | 987 | GPQGVKGESGKPGANG | GPQGVKGESGKP[113]GANG | GP(0.0468)QGVKGESGKP(0.9532)GANG | 21.46 | 12.40 | 1.000 |
| 1457.650 | COL3A1 | 1002 | 1017 | GLAGTAGEPGRDGNPG | GLAGTAGEP[113]GRDGNP[113]G | GLAGTAGEP(1.0000)GRDGNP(1.0000)G | 26.62 | 10.73 | 1.000 |
| 1457.696 | COL3A1 | 1089 | 1103 | GPRGDKGETGERGAA | GPRGDKGETGERGAA | | 18.79 | 10.64 | 0.997 |
| 1460.666 | COL3A1 | 723 | 737 | GLQGMPGERGGLGSP | GLQGM[147]P[113]GERGGLGSP[113] | GLQGM(1.0000)P(1.0000)GERGGLGSP(1.0000) | 30.99 | 14.09 | 1.000 |
| 1461.659 | COL3A1 | 723 | 737 | GLQGMPGERGGLGSP | GLQ[129]GM[147]P[113]GERGGLGSP[113] | GLQGM(1.0000)P(1.0000)GERGGLGSP(1.0000) | 32.37 | 15.38 | 1.000 |
| 1462.671 | COL3A1 | 366 | 380 | GQRGEPGPQGHAGAQ | GQRGEP[113]GPQGHAGAQ | GQRGEP(0.9883)GP(0.0117)QGHAGAQ | 19.79 | 15.99 | 0.977 |
| 1473.706 | COL3A1 | 1089 | 1103 | GPRGDKGETGERGAA | GP[113]RGDKGETGERGAA | GP(1.0000)RGDKGETGERGAA | 18.89 | 12.56 | 1.000 |
| 1488.656 | COL3A1 | 525 | 539 | GRDGVPGGPGMRGMP | GRDGVP[113]GGP[113]GMRGMP[113] | GRDGVP(0.9867)GGP(0.9866)GM(0.0204)RGM(0.0197)P(0.9867) | 22.49 | 20.45 | 1.000 |
| 1504.651 | COL3A1 | 525 | 539 | GRDGVPGGPGMRGMP | GRDGVP[113]GGP[113]GM[147]RGMP[113] | GRDGVP(0.9708)GGP(0.9707)GM(0.9706)RGM(0.1172)P(0.9708) | 21.72 | 19.88 | 0.995 |
| 1508.660 | COL3A1 | 267 | 281 | GFDGRNGEKGETGAP | GFDGRN[115]GEKGETGAP[113] | GFDGRNGEKGETGAP(1.0000) | 21.56 | 12.95 | 1.000 |
| 1548.877 | COL3A1 | 942 | 959 | GAPGPLGIAGITGARGLA | GAPGPLGIAGITGARGLA | | 29.46 | 11.44 | 1.000 |
| 1564.869 | COL3A1 | 942 | 959 | GAPGPLGIAGITGARGLA | GAP[113]GPLGIAGITGARGLA | GAP(0.9886)GP(0.0114)LGIAGITGARGLA | 30.86 | 25.91 | 1.000 |
| 1578.724 | COL3A1 | 828 | 845 | GAPGEKGEGGPPGVAGPP | GAP[113]GEKGEGGPP[113]GVAGP[113]P | GAP(0.7869)GEKGEGGP(0.0637)P(0.7913)GVAGP(0.6791)P(0.6791) | 25.13 | 25.13 | 0.969 |
| 1588.770 | COL3A1 | 859 | 875 | VKGERGSPGGPGAAGFP | VKGERGSP[113]GGP[113]GAAGFP[113] | VKGERGSP(1.0000)GGP(1.0000)GAAGFP(1.0000) | 18.36 | 13.32 | 0.972 |
| 1590.894 | COL3A1 | 945 | 962 | GPLGIAGITGARGLAGPP | GPLGIAGITGARGLAGPP[113] | GP(0.0057)LGIAGITGARGLAGP(0.0183)P(0.9760) | 44.01 | 44.01 | 1.000 |
| 1593.782 | COL3A1 | 774 | 791 | GQPGDKGEGGAPGLPGIA | GQPGDKGEGGAPGLP[113]GIA | GQP(0.0078)GDKGEGGAP(0.0066)GLP(0.9856)GIA | 19.11 | 12.87 | 0.984 |
| 1595.729 | COL3A1 | 669 | 686 | GAPGGKGDAGAPGERGPP | GAP[113]GGKGDAGAP[113]GERGP[113]P | GAP(0.7800)GGKGDAGAP(0.7800)GERGP(0.7200)P(0.7200) | 25.40 | 25.40 | 1.000 |
| 1595.764 | COL3A1 | 1053 | 1070 | GPVGPAGKSGDRGESGPA | GPVGPAGKSGDRGESGPA | | 23.94 | 14.97 | 0.999 |
| 1602.687 | COL3A1 | 1003 | 1019 | LAGTAGEPGRDGNPGSD | LAGTAGEP[113]GRDGNP[113]GSD | LAGTAGEP(1.0000)GRDGNP(1.0000)GSD | 29.46 | 8.34 | 1.000 |
| 1609.774 | COL3A1 | 774 | 791 | GQPGDKGEGGAPGLPGIA | GQP[113]GDKGEGGAPGLP[113]GIA | GQP(0.9888)GDKGEGGAP(0.0222)GLP(0.9889)GIA | 21.48 | 15.01 | 0.992 |
| 1612.718 | COL3A1 | 384 | 401 | GPPGINGSPGGKGEMGPA | GPP[113]GIN[115]GSP[113]GGKGEMGPA | GP(0.0142)P(0.9754)GINGSP(0.9757)GGKGEM(0.0106)GP(0.0242)A | 18.98 | 18.98 | 0.941 |
| 1625.775 | COL3A1 | 774 | 791 | GQPGDKGEGGAPGLPGIA | GQP[113]GDKGEGGAP[113]GLP[113]GIA | GQP(1.0000)GDKGEGGAP(1.0000)GLP(1.0000)GIA | 33.58 | 14.72 | 1.000 |
| 1628.713 | COL3A1 | 384 | 401 | GPPGINGSPGGKGEMGPA | GPP[113]GIN[115]GSP[113]GGKGEM[147]GPA | GP(0.0173)P(0.9884)GINGSP(0.9879)GGKGEM(0.9884)GP(0.0180)A | 19.42 | 17.73 | 0.982 |
| 1637.822 | COL3A1 | 777 | 794 | GDKGEGGAPGLPGIAGPR | GDKGEGGAP[113]GLP[113]GIAGPR | GDKGEGGAP(0.9892)GLP(0.9892)GIAGP(0.0216)R | 30.84 | 20.90 | 1.000 |
| 1643.719 | COL3A1 | 1002 | 1019 | GLAGTAGEPGRDGNPGSD | GLAGTAGEP[113]GRDGNPGSD | GLAGTAGEP(0.9894)GRDGNP(0.0106)GSD | 30.07 | 15.95 | 1.000 |
| 1645.787 | COL3A1 | 858 | 875 | GVKGERGSPGGPGAAGFP | GVKGERGSP[113]GGP[113]GAAGFP[113] | GVKGERGSP(1.0000)GGP(1.0000)GAAGFP(1.0000) | 18.79 | 11.86 | 0.976 |
| 1651.779 | COL3A1 | 516 | 533 | GPRGAAGEPGRDGVPGGP | GPRGAAGEP[113]GRDGVP[113]GGP[113] | GP(0.0334)RGAAGEP(0.9889)GRDGVP(0.9889)GGP(0.9889) | 20.01 | 13.05 | 0.995 |
| 1659.705 | COL3A1 | 1002 | 1019 | GLAGTAGEPGRDGNPGSD | GLAGTAGEP[113]GRDGNP[113]GSD | GLAGTAGEP(1.0000)GRDGNP(1.0000)GSD | 34.45 | 8.58 | 1.000 |
| 1660.691 | COL3A1 | 1002 | 1019 | GLAGTAGEPGRDGNPGSD | GLAGTAGEP[113]GRDGN[115]P[113]GSD | GLAGTAGEP(1.0000)GRDGNP(1.0000)GSD | 21.32 | 7.89 | 1.000 |
| 1664.779 | COL3A1 | 636 | 653 | GPQGLQGLPGTGGPPGEN | GPQGLQGLP[113]GTGGPP[113]GEN | GP(0.0130)QGLQGLP(0.9828)GTGGP(0.0219)P(0.9824)GEN | 38.10 | 33.49 | 1.000 |
| 1665.774 | COL3A1 | 636 | 653 | GPQGLQGLPGTGGPPGEN | GPQ[129]GLQGLP[113]GTGGPP[113]GEN | GP(0.0178)QGLQGLP(0.9738)GTGGP(0.0356)P(0.9729)GEN | 24.24 | 24.24 | 1.000 |
| 1666.730 | COL3A1 | 642 | 659 | GLPGTGGPPGENGKPGEP | GLP[113]GTGGPP[113]GEN[115]GKPGEP[113] | GLP(0.9300)GTGGP(0.1621)P(0.9265)GENGKP(0.0519)GEP(0.9293) | 21.60 | 21.60 | 1.000 |
| 1667.750 | COL3A1 | 771 | 789 | GPAGQPGDKGEGGAPGLPG | GPAGQ[129]P[113]GDKGEGGAP[113]GLP[113]G | GP(0.0483)AGQP(0.9839)GDKGEGGAP(0.9839)GLP(0.9839)G | 18.09 | 12.89 | 0.992 |
| 1668.791 | COL3A1 | 822 | 839 | GGKGERGAPGEKGEGGPP | GGKGERGAP[113]GEKGEGGPP[113] | GGKGERGAP(0.9888)GEKGEGGP(0.0224)P(0.9888) | 18.90 | 18.90 | 0.990 |
| 1669.769 | COL3A1 | 921 | 938 | GPKGDAGQPGEKGSPGAQ | GPKGDAGQP[113]GEKGSP[113]GAQ | GP(0.0209)KGDAGQP(0.9895)GEKGSP(0.9895)GAQ | 27.77 | 14.37 | 1.000 |
| 1681.747 | COL3A1 | 642 | 659 | GLPGTGGPPGENGKPGEP | GLP[113]GTGGPP[113]GENGKP[113]GEP[113] | GLP(0.9883)GTGGP(0.0470)P(0.9883)GENGKP(0.9882)GEP(0.9882) | 23.42 | 19.27 | 0.959 |
| 1682.740 | COL3A1 | 642 | 659 | GLPGTGGPPGENGKPGEP | GLP[113]GTGGPP[113]GEN[115]GKP[113]GEP[113] | GLP(0.9890)GTGGP(0.0439)P(0.9890)GENGKP(0.9890)GEP(0.9890) | 25.18 | 20.88 | 1.000 |
| 1683.771 | COL3A1 | 999 | 1016 | GLPGLAGTAGEPGRDGNP | GLP[113]GLAGTAGEP[113]GRDGNP[113] | GLP(1.0000)GLAGTAGEP(1.0000)GRDGNP(1.0000) | 22.10 | 12.84 | 0.993 |
| 1684.814 | COL3A1 | 426 | 443 | GLRGGAGEPGKNGAKGEP | GLRGGAGEP[113]GKN[115]GAKGEP[113] | GLRGGAGEP(1.0000)GKNGAKGEP(1.0000) | 30.77 | 16.57 | 1.000 |
| 1685.753 | COL3A1 | 519 | 536 | GAAGEPGRDGVPGGPGMR | GAAGEP[113]GRDGVP[113]GGP[113]GMR | GAAGEP(0.7802)GRDGVP(0.7801)GGP(0.7199)GM(0.7199)R | 21.86 | 21.86 | 0.964 |
| 1687.707 | COL3A1 | 195 | 212 | GSPGYQGPPGEPGQAGPS | GSP[113]GYQGPP[113]GEP[113]GQAGPS | GSP(0.9891)GYQGP(0.0163)P(0.9893)GEP(0.9894)GQAGP(0.0159)S | 41.28 | 36.54 | 1.000 |
| 1688.686 | COL3A1 | 195 | 212 | GSPGYQGPPGEPGQAGPS | GSP[113]GYQGPP[113]GEP[113]GQ[129]AGPS | GSP(0.9875)GYQGP(0.0195)P(0.9877)GEP(0.9878)GQAGP(0.0174)S | 21.15 | 19.45 | 0.996 |
| 1692.790 | COL3A1 | 495 | 512 | GPNGIPGEKGPAGERGAP | GPNGIP[113]GEKGPAGERGAP[113] | GP(0.0081)NGIP(0.7895)GEKGP(0.4610)AGERGAP(0.7414) | 19.07 | 18.65 | 1.000 |

### Supplemental Table 3. Col3a1 peptide sequences from normal breast by LC-MS/MS.

| | | | | | | | | | |
|---|---|---|---|---|---|---|---|---|---|
| 1693.800 | COL3A1 | 495 | 512 | GPNGIPGEKGPAGERGAP | GPN[115]GIP[113]GEKGPAGERGAP[113] | GP(0.0105)NGIP(0.9890)GEKGP(0.0116)AGERGAP(0.9889) | 30.02 | 23.65 | 1.000 |
| 1699.780 | COL3A1 | 978 | 995 | GESGKPGANGLSGERGPP | GESGKP[113]GAN[115]GLSGERGP[113]P | GESGKP(0.7196)GANGLSGERGP(0.6402)P(0.6402) | 18.12 | 18.12 | 0.996 |
| 1701.752 | COL3A1 | 519 | 536 | GAAGEPGRDGVPGGPGMR | GAAGEP[113]GRDGVP[113]GGP[113]GM[147]R | GAAGEP(1.0000)GRDGVP(1.0000)GGP(1.0000)GM(1.0000)R | 20.21 | 11.79 | 0.998 |
| 1703.703 | COL3A1 | 192 | 209 | GSPGSPGYQGPPGEPGQA | GSP[113]GSP[113]GYQGPP[113]GEP[113]GQA | GSP(0.9870)GSP(0.9871)GYQGP(0.0518)P(0.9871)GEP(0.9871)GQA | 28.61 | 26.81 | 1.000 |
| 1709.809 | COL3A1 | 495 | 512 | GPNGIPGEKGPAGERGAP | GPN[115]GIP[113]GEKGP[113]AGERGAP[113] | GP(0.4293)NGIP(0.8593)GEKGP(0.8554)AGERGAP(0.8560) | 22.08 | 13.44 | 0.996 |
| 1716.732 | COL3A1 | 1002 | 1020 | GLAGTAGEPGRDGNPGSDG | GLAGTAGEP[113]GRDGNP[113]GSDG | GLAGTAGEP(1.0000)GRDGNP(1.0000)GSDG | 18.53 | 9.53 | 1.000 |
| 1731.799 | COL3A1 | 720 | 737 | GTPGLQGMPGERGGLGSP | GTP[113]GLQGM[147]P[113]GERGGLGSP[113] | GTP(1.0000)GLQGM(1.0000)P(1.0000)GERGGLGSP(1.0000) | 19.54 | 12.47 | 0.993 |
| 1744.762 | COL3A1 | 282 | 299 | GLKGENGLPGENGAPGPM | GLKGEN[115]GLP[113]GEN[115]GAP[113]GP[113]M | GLKGENGLP(0.9627)GENGAP(0.9626)GP(0.9623)M(0.1124) | 23.52 | 23.52 | 0.998 |
| 1750.865 | COL3A1 | 776 | 794 | PGDKGEGGAPGLPGIAGPR | P[113]GDKGEGGAP[113]GLP[113]GIAGPR | P(0.9715)GDKGEGGAP(0.9716)GLP(0.9713)GIAGP(0.0855)R | 18.97 | 13.50 | 1.000 |
| 1834.899 | COL3A1 | 771 | 791 | GPAGQPGDKGEGGAPGLPGIA | GPAGQP[113]GDKGEGGAPGLP[113]GIA | GP(0.0166)AGQP(0.9814)GDKGEGGAP(0.0209)GLP(0.9812)GIA | 18.15 | 14.49 | 0.998 |
| 1835.885 | COL3A1 | 771 | 791 | GPAGQPGDKGEGGAPGLPGIA | GPAGQ[129]P[113]GDKGEGGAPGLP[113]GIA | GP(0.0115)AGQP(0.9878)GDKGEGGAP(0.0129)GLP(0.9878)GIA | 20.60 | 16.77 | 0.983 |
| 1850.881 | COL3A1 | 771 | 791 | GPAGQPGDKGEGGAPGLPGIA | GPAGQP[113]GDKGEGGAP[113]GLP[113]GIA | GP(0.1961)AGQP(0.9340)GDKGEGGAP(0.9349)GLP(0.9349)GIA | 19.79 | 18.63 | 0.986 |
| 1851.862 | COL3A1 | 771 | 791 | GPAGQPGDKGEGGAPGLPGIA | GPAGQ[129]P[113]GDKGEGGAP[113]GLP[113]GIA | GP(0.0453)AGQP(0.9849)GDKGEGGAP(0.9849)GLP(0.9849)GIA | 20.13 | 16.31 | 0.963 |
| 1853.864 | COL3A1 | 1003 | 1022 | LAGTAGEPGRDGNPGSDGLP | LAGTAGEP[113]GRDGNPGSDGLP | LAGTAGEP(0.9854)GRDGNP(0.0081)GSDGLP(0.0065) | 19.21 | 12.76 | 0.974 |
| 1869.854 | COL3A1 | 1003 | 1022 | LAGTAGEPGRDGNPGSDGLP | LAGTAGEP[113]GRDGNPGSDGLP[113] | LAGTAGEP(0.9879)GRDGNP(0.0242)GSDGLP(0.9879) | 20.23 | 16.16 | 0.995 |
| 1870.844 | COL3A1 | 1003 | 1022 | LAGTAGEPGRDGNPGSDGLP | LAGTAGEP[113]GRDGN[115]PGSDGLP[113] | LAGTAGEP(0.9723)GRDGNP(0.0556)GSDGLP(0.9720) | 20.33 | 16.23 | 1.000 |
| 1885.833 | COL3A1 | 1003 | 1022 | LAGTAGEPGRDGNPGSDGLP | LAGTAGEP[113]GRDGNP[113]GSDGLP[113] | LAGTAGEP(1.0000)GRDGNP(1.0000)GSDGLP(1.0000) | 20.55 | 11.52 | 0.960 |
| 1885.860 | COL3A1 | 387 | 407 | GINGSPGGKGEMGPAGIPGAP | GIN[115]GSP[113]GGKGEM[147]GPAGIP[113]GAP[113] | GINGSP(0.8811)GGKGEM(0.8783)GP(0.4779)AGIP(0.8814)GAP(0.8814) | 22.62 | 19.16 | 0.997 |
| 1886.848 | COL3A1 | 1003 | 1022 | LAGTAGEPGRDGNPGSDGLP | LAGTAGEP[113]GRDGN[115]P[113]GSDGLP[113] | LAGTAGEP(1.0000)GRDGNP(1.0000)GSDGLP(1.0000) | 19.12 | 12.89 | 0.986 |
| 1907.890 | COL3A1 | 640 | 659 | LQGLPGTGGPPGENGKPGEP | LQGLP[113]GTGGPP[113]GEN[115]GKPGEP[113] | LQGLP(0.9773)GTGGP(0.0344)P(0.9771)GENGKP(0.0341)GEP(0.9771) | 20.81 | 20.81 | 0.976 |
| 1907.906 | COL3A1 | 738 | 758 | GPKGDKGEPGGPGADGVPGKD | GPKGDKGEP[113]GGPGADGVPGKD | GP(0.0034)KGDKGEP(0.9887)GGP(0.0040)GADGVP(0.0038)GKD | 21.55 | 17.45 | 1.000 |
| 1910.882 | COL3A1 | 1002 | 1022 | GLAGTAGEPGRDGNPGSDGLP | GLAGTAGEP[113]GRDGNPGSDGLP | GLAGTAGEP(0.9859)GRDGNP(0.0082)GSDGLP(0.0059) | 20.92 | 12.82 | 0.976 |
| 1911.873 | COL3A1 | 384 | 404 | GPPGINGSPGGKGEMGPAGIP | GPP[113]GIN[115]GSP[113]GGKGEM[147]GPAGIP[113] | GP(0.0265)P(0.9876)GINGSP(0.9876)GGKGEM(0.9876)GP(0.0231)AGIP(0.9876) | 26.24 | 26.24 | 1.000 |
| 1913.873 | COL3A1 | 915 | 935 | GSPGVSGPKGDAGQPGEKGSP | GSP[113]GVSGPKGDAGQP[113]GEKGSP[113] | GSP(0.9666)GVSGP(0.0996)KGDAGQP(0.9669)GEKGSP(0.9669) | 24.35 | 18.59 | 1.000 |
| 1914.864 | COL3A1 | 915 | 935 | GSPGVSGPKGDAGQPGEKGSP | GSP[113]GVSGPKGDAGQ[129]P[113]GEKGSP[113] | GSP(0.9898)GVSGP(0.0306)KGDAGQP(0.9898)GEKGSP(0.9898) | 20.59 | 16.45 | 1.000 |
| 1914.892 | COL3A1 | 717 | 737 | GAAGTPGLQGMPGERGGLGSP | GAAGTPGLQGM[147]P[113]GERGGLGSP[113] | GAAGTP(0.0561)GLQGM(0.9812)P(0.9813)GERGGLGSP(0.9813) | 20.69 | 12.69 | 0.992 |
| 1919.918 | COL3A1 | 741 | 761 | GDKGEPGGPGADGVPGKDGPR | GDKGEPGGPGADGVPGKDGPR | | 18.76 | 10.64 | 1.000 |
| 1919.932 | COL3A1 | 774 | 794 | GQPGDKGEGGAPGLPGIAGPR | GQPGDKGEGGAP[113]GLP[113]GIAGPR | GQP(0.0115)GDKGEGGAP(0.9881)GLP(0.9880)GIAGP(0.0124)R | 21.83 | 14.80 | 1.000 |
| 1922.901 | COL3A1 | 640 | 659 | LQGLPGTGGPPGENGKPGEP | LQGLP[113]GTGGP[113]PGENGKP[113]GEP[113] | LQGLP(0.8194)GTGGP(0.7710)P(0.7710)GENGKP(0.8193)GEP(0.8194) | 22.83 | 22.83 | 1.000 |
| 1923.885 | COL3A1 | 640 | 659 | LQGLPGTGGPPGENGKPGEP | LQGLP[113]GTGGP[113]PGEN[115]GKP[113]GEP[113] | LQGLP(0.8192)GTGGP(0.7715)P(0.7715)GENGKP(0.8190)GEP(0.8188) | 21.68 | 21.68 | 1.000 |
| 1926.874 | COL3A1 | 1002 | 1022 | GLAGTAGEPGRDGNPGSDGLP | GLAGTAGEP[113]GRDGNPGSDGLP[113] | GLAGTAGEP(0.9857)GRDGNP(0.0286)GSDGLP(0.9858) | 18.03 | 14.48 | 0.985 |
| 1926.974 | COL3A1 | 976 | 995 | VKGESGKPGANGLSGERGPP | VKGESGKP[113]GAN[115]GLSGERGP[113]P | VKGESGKP(0.7208)GANGLSGERGP(0.6396)P(0.6396) | 20.90 | 20.90 | 0.999 |
| 1927.858 | COL3A1 | 1002 | 1022 | GLAGTAGEPGRDGNPGSDGLP | GLAGTAGEP[113]GRDGN[115]PGSDGLP[113] | GLAGTAGEP(0.9902)GRDGNP(0.0197)GSDGLP(0.9901) | 34.04 | 23.33 | 1.000 |
| 1927.917 | COL3A1 | 855 | 875 | GPQGVKGERGSPGGPGAAGFP | GPQGVKGERGSP[113]GGP[113]GAAGFP[113] | GP(0.5330)QGVKGERGSP(0.8174)GGP(0.8240)GAAGFP(0.8256) | 21.03 | 13.74 | 0.996 |
| 1928.908 | COL3A1 | 855 | 875 | GPQGVKGERGSPGGPGAAGFP | GPQ[129]GVKGERGSP[113]GGP[113]GAAGFP[113] | GP(0.2181)QGVKGERGSP(0.9269)GGP(0.9270)GAAGFP(0.9280) | 18.11 | 12.89 | 0.976 |
| 1930.885 | COL3A1 | 717 | 737 | GAAGTPGLQGMPGERGGLGSP | GAAGTP[113]GLQGM[147]P[113]GERGGLGSP[113] | GAAGTP(1.0000)GLQGM(1.0000)P(1.0000)GERGGLGSP(1.0000) | 18.95 | 10.63 | 0.999 |
| 1935.905 | COL3A1 | 741 | 761 | GDKGEPGGPGADGVPGKDGPR | GDKGEP[113]GGPGADGVPGKDGPR | GDKGEP(0.9868)GGP(0.0042)GADGVP(0.0046)GKDGP(0.0045)R | 20.19 | 15.93 | 1.000 |
| 1935.928 | COL3A1 | 774 | 794 | GQPGDKGEGGAPGLPGIAGPR | GQP[113]GDKGEGGAP[113]GLP[113]GIAGPR | GQP(0.9711)GDKGEGGAP(0.9711)GLP(0.9709)GIAGP(0.0869)R | 22.95 | 20.98 | 1.000 |
| 1942.832 | COL3A1 | 1002 | 1022 | GLAGTAGEPGRDGNPGSDGLP | GLAGTAGEP[113]GRDGNP[113]GSDGLP[113] | GLAGTAGEP(1.0000)GRDGNP(1.0000)GSDGLP(1.0000) | 62.38 | 12.85 | 1.000 |
| 1942.859 | COL3A1 | 459 | 479 | GAKGEDGKDGSPGEPGANGLP | GAKGEDGKDGSPGEP[113]GAN[115]GLP[113] | GAKGEDGKDGSP(0.1961)GEP(0.9007)GANGLP(0.9032) | 19.09 | 15.40 | 0.997 |
| 1943.851 | COL3A1 | 1002 | 1022 | GLAGTAGEPGRDGNPGSDGLP | GLAGTAGEP[113]GRDGN[115]P[113]GSDGLP[113] | GLAGTAGEP(1.0000)GRDGNP(1.0000)GSDGLP(1.0000) | 21.92 | 14.38 | 0.989 |
| 1943.919 | COL3A1 | 855 | 875 | GPQGVKGERGSPGGPGAAGFP | GP[113]QGVKGERGSP[113]GGP[113]GAAGFP[113] | GP(1.0000)QGVKGERGSP(1.0000)GGP(1.0000)GAAGFP(1.0000) | 21.46 | 12.57 | 0.996 |
| 1944.797 | COL3A1 | 192 | 212 | GSPGSPGYQGPPGEPGQAGPS | GSP[113]GSP[113]GYQGPP[113]GEP[113]GQAGPS | GSP(0.9831)GSP(0.9831)GYQGP(0.0444)P(0.9829)GEP(0.9831)GQAGP(0.0233)S | 31.94 | 29.84 | 1.000 |
| 1945.791 | COL3A1 | 192 | 212 | GSPGSPGYQGPPGEPGQAGPS | GSP[113]GSP[113]GYQGPP[113]GEP[113]GQ[129]AGPS | GSP(0.9502)GSP(0.9501)GYQGP(0.1784)P(0.9483)GEP(0.9504)GQAGP(0.0226)S | 35.40 | 33.03 | 0.990 |
| 1950.877 | COL3A1 | 648 | 668 | GPPGENGKPGEPGPKGDAGAP | GPP[113]GEN[115]GKP[113]GEPGP[113]KGDAGAP[113] | GP(0.0201)P(0.9798)GENGKP(0.9768)GEP(0.0633)GP(0.9797)KGDAGAP(0.9802) | 19.52 | 17.88 | 0.988 |
| 1951.912 | COL3A1 | 741 | 761 | GDKGEPGGPGADGVPGKDGPR | GDKGEP[113]GGP[113]GADGVPGKDGPR | GDKGEP(0.9883)GGP(0.9883)GADGVP(0.0108)GKDGP(0.0126)R | 25.34 | 15.14 | 1.000 |
| 1958.849 | COL3A1 | 459 | 479 | GAKGEDGKDGSPGEPGANGLP | GAKGEDGKDGSP[113]GEP[113]GAN[115]GLP[113] | GAKGEDGKDGSP(1.0000)GEP(1.0000)GANGLP(1.0000) | 19.38 | 10.27 | 0.995 |
| 1964.914 | COL3A1 | 639 | 659 | GLQGLPGTGGPPGENGKPGEP | GLQGLP[113]GTGGPP[113]GEN[115]GKP[113]GEP | GLQGLP(0.9528)GTGGP(0.0439)P(0.9525)GENGKP(0.9513)GEP(0.0995) | 23.87 | 23.87 | 0.998 |
| 1970.869 | COL3A1 | 1002 | 1023 | GLAGTAGEPGRDGNPGSDGLPG | GLAGTAGEP[113]GRDGNPGSDGLPG | GLAGTAGEP(0.9884)GRDGNP(0.0058)GSDGLP(0.0058)G | 20.05 | 19.87 | 1.000 |
| 1979.920 | COL3A1 | 639 | 659 | GLQGLPGTGGPPGENGKPGEP | GLQGLP[113]GTGGPP[113]GENGKP[113]GEP[113] | GLQGLP(0.9814)GTGGP(0.0744)P(0.9813)GENGKP(0.9814)GEP(0.9814) | 23.28 | 21.18 | 0.992 |
| 1980.890 | COL3A1 | 639 | 659 | GLQGLPGTGGPPGENGKPGEP | GLQGLP[113]GTGGPP[113]GEN[115]GKP[113]GEP[113] | GLQGLP(0.9856)GTGGP(0.0567)P(0.9859)GENGKP(0.9859)GEP(0.9859) | 29.98 | 25.99 | 1.000 |
| 1983.959 | COL3A1 | 975 | 995 | GVKGESGKPGANGLSGERGPP | GVKGESGKP[113]GAN[115]GLSGERGPP[113] | GVKGESGKP(0.9756)GANGLSGERGP(0.0490)P(0.9754) | 22.40 | 22.40 | 1.000 |
| 1986.861 | COL3A1 | 519 | 539 | GAAGEPGRDGVPGGPGMRGMP | GAAGEP[113]GRDGVP[113]GGP[113]GMRGMP[113] | GAAGEP(0.8197)GRDGVP(0.8182)GGP(0.8052)GM(0.4861)RGM(0.2497)P(0.8211) | 21.31 | 21.31 | 0.999 |
| 1998.975 | COL3A1 | 972 | 992 | GPQGVKGESGKPGANGLSGER | GPQGVKGESGKP[113]GAN[115]GLSGER | GP(0.0122)QGVKGESGKP(0.9878)GANGLSGER | 18.99 | 9.12 | 1.000 |
| 2002.844 | COL3A1 | 519 | 539 | GAAGEPGRDGVPGGPGMRGMP | GAAGEP[113]GRDGVP[113]GGP[113]GMRGM[147]P[113] | GAAGEP(0.8468)GRDGVP(0.8468)GGP(0.8065)GM(0.8065)RGM(0.8468)P(0.8466) | 21.73 | 20.08 | 1.000 |
| 2079.034 | COL3A1 | 225 | 245 | GPAGKDGESGRPGRPGERGLP | GPAGKDGESGRPGRP[113]GERGLP[113] | GP(0.0229)AGKDGESGRP(0.0127)GRP(0.9822)GERGLP(0.9822) | 18.92 | 16.83 | 1.000 |
| 2095.030 | COL3A1 | 225 | 245 | GPAGKDGESGRPGRPGERGLP | GPAGKDGESGRP[113]GRP[113]GERGLP[113] | GP(0.0516)AGKDGESGRP(0.9828)GRP(0.9828)GERGLP(0.9828) | 19.00 | 9.78 | 1.000 |
| 2113.056 | COL3A1 | 1092 | 1112 | GDKGETGERGAAGIKGHRGFP | GDKGETGERGAAGIKGHRGFP[113] | GDKGETGERGAAGIKGHRGFP(1.0000) | 18.50 | 11.25 | 1.000 |
| 2129.052 | COL3A1 | 1048 | 1070 | HPGPPGPVGPAGKSGDRGESGPA | HP[113]GP[113]P[113]GPVGPAGKSGDRGESGPA | HP(0.9840)GP(0.9840)P(0.9840)GP(0.0119)VGP(0.0159)AGKSGDRGESGP(0.0202)A | 23.00 | 18.25 | 0.981 |
| 2155.999 | COL3A1 | 1002 | 1024 | GLAGTAGEPGRDGNPGSDGLPGR | GLAGTAGEP[113]GRDGNP[113]GSDGLP[113]GR | GLAGTAGEP(1.0000)GRDGNP(1.0000)GSDGLP(1.0000)GR | 19.50 | 15.07 | 1.000 |
| 2161.041 | COL3A1 | 771 | 794 | GPAGQPGDKGEGGAPGLPGIAGPR | GPAGQP[113]GDKGEGGAP[113]GLP[113]GIAGPR | GP(0.0187)AGQP(0.9594)GDKGEGGAP(0.9594)GLP(0.9577)GIAGP(0.1048)R | 28.60 | 26.38 | 1.000 |
| 2165.949 | COL3A1 | 645 | 668 | GTGGPPGENGKPGEPGPKGDAGAP | GTGGPP[113]GEN[115]GKP[113]GEP[113]GPKGDAGAP[113] | GTGGP(0.0558)P(0.9656)GENGKP(0.9635)GEP(0.9653)GP(0.0839)KGDAGAP(0.9659) | 21.57 | 19.73 | 1.000 |
| 2169.975 | COL3A1 | 915 | 938 | GSPGVSGPKGDAGQPGEKGSPGAQ | GSP[113]GVSGPKGDAGQP[113]GEKGSP[113]GAQ | GSP(0.9894)GVSGP(0.0318)KGDAGQP(0.9894)GEKGSP(0.9894)GAQ | 24.81 | 20.43 | 1.000 |
| 2202.077 | COL3A1 | 738 | 761 | GPKGDKGEPGGPGADGVPGKDGPR | GPKGDKGEPGGPGADGVPGKDGPR | | 27.03 | 13.92 | 1.000 |
| 2210.028 | COL3A1 | 999 | 1022 | GLPGLAGTAGEPGRDGNPGSDGLP | GLP[113]GLAGTAGEP[113]GRDGNPGSDGLP[113] | GLP(0.9893)GLAGTAGEP(0.9893)GRDGNP(0.0322)GSDGLP(0.9893) | 21.32 | 21.32 | 1.000 |
| 2218.058 | COL3A1 | 738 | 761 | GPKGDKGEPGGPGADGVPGKDGPR | GPKGDKGEP[113]GGPGADGVPGKDGPR | GP(0.0031)KGDKGEP(0.9871)GGP(0.0039)GADGVP(0.0030)GKDGP(0.0030)R | 25.52 | 21.05 | 1.000 |
| 2226.024 | COL3A1 | 999 | 1022 | GLPGLAGTAGEPGRDGNPGSDGLP | GLP[113]GLAGTAGEP[113]GRDGNP[113]GSDGLP[113] | GLP(1.0000)GLAGTAGEP(1.0000)GRDGNP(1.0000)GSDGLP(1.0000) | 21.19 | 8.83 | 0.998 |
| 2228.001 | COL3A1 | 456 | 479 | GVPGAKGEDGKDGSPGEPGANGLP | GVP[113]GAKGEDGKDGSP[113]GEP[113]GAN[115]GLP[113] | GVP(1.0000)GAKGEDGKDGSP(1.0000)GEP(1.0000)GANGLP(1.0000) | 20.66 | 12.24 | 0.995 |
| 2234.062 | COL3A1 | 738 | 761 | GPKGDKGEPGGPGADGVPGKDGPR | GP[113]KGDKGEP[113]GGPGADGVPGKDGPR | GP(0.9863)KGDKGEP(0.9890)GGP(0.0080)GADGVP(0.0085)GKDGP(0.0082)R | 29.67 | 27.94 | 1.000 |
| 2250.032 | COL3A1 | 738 | 761 | GPKGDKGEPGGPGADGVPGKDGPR | GP[113]KGDKGEP[113]GGP[113]GADGVPGKDGPR | GP(0.9875)KGDKGEP(0.9878)GGP(0.9879)GADGVP(0.0187)GKDGP(0.0181)R | 24.62 | 16.27 | 0.985 |
| 2250.094 | COL3A1 | 972 | 995 | GPQGVKGESGKPGANGLSGERGPP | GPQGVKGESGKPGAN[115]GLSGERGP[113]P | GP(0.0036)QGVKGESGKP(0.0038)GANGLSGERGP(0.4963)P(0.4963) | 19.78 | 19.78 | 1.000 |
| 2266.091 | COL3A1 | 972 | 995 | GPQGVKGESGKPGANGLSGERGPP | GPQGVKGESGKP[113]GAN[115]GLSGERGP[113]P | GP(0.0406)QGVKGESGKP(0.7518)GANGLSGERGP(0.6038)P(0.6038) | 40.20 | 40.20 | 1.000 |
| 2267.087 | COL3A1 | 972 | 995 | GPQGVKGESGKPGANGLSGERGPP | GPQ[129]GVKGESGKP[113]GAN[115]GLSGERGP[113]P | GP(0.0639)QGVKGESGKP(0.7431)GANGLSGERGP(0.5965)P(0.5965) | 22.29 | 22.29 | 1.000 |
| 2271.031 | COL3A1 | 1002 | 1025 | GLAGTAGEPGRDGNPGSDGLPGRD | GLAGTAGEP[113]GRDGNP[113]GSDGLP[113]GRD | GLAGTAGEP(1.0000)GRDGNP(1.0000)GSDGLP(1.0000)GRD | 27.32 | 13.72 | 1.000 |
| 2281.101 | COL3A1 | 972 | 995 | GPQGVKGESGKPGANGLSGERGPP | GP[113]QGVKGESGKP[113]GANGLSGERGP[113]P | GP(0.7795)QGVKGESGKP(0.7797)GANGLSGERGP(0.7204)P(0.7204) | 20.09 | 20.09 | 1.000 |
| 2282.086 | COL3A1 | 972 | 995 | GPQGVKGESGKPGANGLSGERGPP | GP[113]QGVKGESGKP[113]GAN[115]GLSGERGP[113]P | GP(0.7796)QGVKGESGKP(0.7801)GANGLSGERGP(0.7201)P(0.7201) | 25.25 | 25.25 | 1.000 |
| 2297.044 | COL3A1 | 516 | 539 | GPRGAAGEPGRDGVPGGPGMRGMP | GPRGAAGEP[113]GRDGVP[113]GGP[113]GMRGMP[113] | GP(0.0161)RGAAGEP(0.9764)GRDGVP(0.9765)GGP(0.9763)GM(0.0579)RGM(0.0213)P(0.9754) | 23.45 | 21.31 | 0.981 |
| 2313.032 | COL3A1 | 516 | 539 | GPRGAAGEPGRDGVPGGPGMRGMP | GPRGAAGEP[113]GRDGVP[113]GGP[113]GMRGM[147]P[113] | GP(0.0886)RGAAGEP(0.9041)GRDGVP(0.9041)GGP(0.9045)GM(0.3952)RGM(0.8994)P(0.9042) | 27.82 | 25.81 | 1.000 |
| 2394.111 | COL3A1 | 417 | 443 | GPAGANGAPGLRGGAGEPGKNGAKGEP | GPAGAN[115]GAP[113]GLRGGAGEP[113]GKN[115]GAKGEP[113] | GP(0.0506)AGANGAP(0.9832)GLRGGAGEP(0.9831)GKNGAKGEP(0.9831) | 24.06 | 18.39 | 1.000 |
| 2394.175 | COL3A1 | 417 | 443 | GPAGANGAPGLRGGAGEPGKNGAKGEP | GPAGANGAP[113]GLRGGAGEP[113]GKN[115]GAKGEP[113] | GP(0.0915)AGANGAP(0.9696)GLRGGAGEP(0.9695)GKNGAKGEP(0.9694) | 19.67 | 19.06 | 1.000 |
| 2433.117 | COL3A1 | 642 | 668 | GLPGTGGPPGENGKPGEPGPKGDAGAP | GLP[113]GTGGPP[113]GEN[115]GKPGEP[113]GPKGDAGAP[113] | GLP(0.8798)GTGGP(0.0206)P(0.8797)GENGKP(0.4654)GEP(0.8534)GP(0.0231)KGDAGAP(0.8780) | 27.11 | 27.11 | 1.000 |
| 2437.101 | COL3A1 | 915 | 941 | GSPGVSGPKGDAGQPGEKGSPGAQGPP | GSP[113]GVSGPKGDAGQP[113]GEKGSP[113]GAQGPP[113] | GSP(0.7390)GVSGP(0.7390)KGDAGQP(0.8329)GEKGSP(0.8330)GAQGP(0.0233)P(0.8329) | 29.37 | 29.37 | 1.000 |
| 2438.090 | COL3A1 | 915 | 941 | GSPGVSGPKGDAGQPGEKGSPGAQGPP | GSP[113]GVSGPKGDAGQP[113]GEKGSP[113]GAQ[129]GPP[113] | GSP(0.7364)GVSGP(0.7364)KGDAGQP(0.8279)GEKGSP(0.8279)GAQGP(0.0439)P(0.8276) | 20.74 | 20.74 | 0.961 |

### Supplemental Table 3. Col3a1 peptide sequences from normal breast by LC-MS/MS.

| | | | | | | | | | |
|---|---|---|---|---|---|---|---|---|---|
| 2439.111 | COL3A1 | 1003 | 1028 | LAGTAGEPGRDGNPGSDGLPGRDGSP | LAGTAGEP[113]GRDGNPGSDGLPGRDGSP[113] | LAGTAGEP(0.7463)GRDGNP(0.2564)GSDGLP(0.2503)GRDGSP(0.7470) | 19.15 | 18.70 | 0.969 |
| 2455.091 | COL3A1 | 1003 | 1028 | LAGTAGEPGRDGNPGSDGLPGRDGSP | LAGTAGEP[113]GRDGNPGSDGLP[113]GRDGSP[113] | LAGTAGEP(0.9964)GRDGNP(0.0528)GSDGLP(0.9746)GRDGSP(0.9762) | 24.79 | 20.63 | 1.000 |
| 2471.091 | COL3A1 | 1003 | 1028 | LAGTAGEPGRDGNPGSDGLPGRDGSP | LAGTAGEP[113]GRDGNP[113]GSDGLP[113]GRDGSP[113] | LAGTAGEP(1.0000)GRDGNP(1.0000)GSDGLP(1.0000)GRDGSP(1.0000) | 25.91 | 8.00 | 1.000 |
| 2472.085 | COL3A1 | 1003 | 1028 | LAGTAGEPGRDGNPGSDGLPGRDGSP | LAGTAGEP[113]GRDGN[115]P[113]GSDGLP[113]GRDGSP[113] | LAGTAGEP(1.0000)GRDGNP(1.0000)GSDGLP(1.0000)GRDGSP(1.0000) | 23.49 | 12.04 | 1.000 |
| 2496.122 | COL3A1 | 1002 | 1028 | GLAGTAGEPGRDGNPGSDGLPGRDGSF | GLAGTAGEP[113]GRDGNPGSDGLP[113]GRDGSP | GLAGTAGEP(0.8450)GRDGNP(0.0243)GSDGLP(0.8202)GRDGSP(0.3105) | 26.97 | 24.76 | 1.000 |
| 2499.142 | COL3A1 | 459 | 485 | GAKGEDGKDGSPGEPGANGLPGAAGEF | GAKGEDGKDGSP[113]GEP[113]GANGLP[113]GAAGER | GAKGEDGKDGSP(1.0000)GEP(1.0000)GANGLP(1.0000)GAAGER | 20.25 | 10.76 | 1.000 |
| 2500.089 | COL3A1 | 459 | 485 | GAKGEDGKDGSPGEPGANGLPGAAGEF | GAKGEDGKDGSP[113]GEP[113]GAN[115]GLP[113]GAAGER | GAKGEDGKDGSP(1.0000)GEP(1.0000)GANGLP(1.0000)GAAGER | 24.43 | 17.86 | 1.000 |
| 2512.080 | COL3A1 | 1002 | 1028 | GLAGTAGEPGRDGNPGSDGLPGRDGSF | GLAGTAGEP[113]GRDGNP[113]GSDGLPGRDGSP[113] | GLAGTAGEP(0.8089)GRDGNP(0.8087)GSDGLP(0.5852)GRDGSP(0.7972) | 31.48 | 31.07 | 1.000 |
| 2513.116 | COL3A1 | 1002 | 1028 | GLAGTAGEPGRDGNPGSDGLPGRDGSF | GLAGTAGEP[113]GRDGN[115]PGSDGLP[113]GRDGSP[113] | GLAGTAGEP(0.9865)GRDGNP(0.0406)GSDGLP(0.9865)GRDGSP(0.9865) | 20.24 | 16.35 | 1.000 |
| 2528.062 | COL3A1 | 1002 | 1028 | GLAGTAGEPGRDGNPGSDGLPGRDGSF | GLAGTAGEP[113]GRDGNP[113]GSDGLP[113]GRDGSP[113] | GLAGTAGEP(1.0000)GRDGNP(1.0000)GSDGLP(1.0000)GRDGSP(1.0000) | 84.76 | 23.53 | 1.000 |
| 2529.109 | COL3A1 | 1002 | 1028 | GLAGTAGEPGRDGNPGSDGLPGRDGSF | GLAGTAGEP[113]GRDGN[115]P[113]GSDGLP[113]GRDGSP[113] | GLAGTAGEP(1.0000)GRDGNP(1.0000)GSDGLP(1.0000)GRDGSP(1.0000) | 26.73 | 21.93 | 1.000 |
| 2674.254 | COL3A1 | 640 | 668 | LQGLPGTGGPPGENGKPGEPGPKGDAG | LQGLP[113]GTGGP[113]P[113]GEN[115]GKPGEPGPKGDAGAP[1 | LQGLP(0.6943)GTGGP(0.5852)P(0.5852)GENGKP(0.3058)GEP(0.5852)GP(0.5852)KGDAGAP(0.6593) | 21.23 | 21.23 | 1.000 |
| 2725.222 | COL3A1 | 459 | 488 | GAKGEDGKDGSPGEPGANGLPGAAGEF | GAKGEDGKDGSPGEP[113]GAN[115]GLP[113]GAAGERGAP[113] | GAKGEDGKDGSP(0.0425)GEP(0.9858)GANGLP(0.9859)GAAGERGAP(0.9858) | 26.19 | 20.88 | 1.000 |
| 2730.294 | COL3A1 | 639 | 668 | GLQGLPGTGGPPGENGKPGEPGPKGDA | GLQGLP[113]GTGGPPGENGKP[113]GEP[113]GP[113]KGDAGAP | GLQGLP(0.8166)GTGGP(0.1275)P(0.1555)GENGKP(0.8129)GEP(0.8129)GP(0.7797)KGDAGAP(0.4948) | 18.14 | 18.14 | 0.997 |
| 2731.269 | COL3A1 | 639 | 668 | GLQGLPGTGGPPGENGKPGEPGPKGDA | GLQGLP[113]GTGGP[113]PGEN[115]GKPGEP[113]GPKGDAGAP[ | GLQGLP(0.7511)GTGGP(0.6307)P(0.6307)GENGKP(0.5249)GEP(0.6946)GP(0.0182)KGDAGAP(0.7497) | 25.90 | 25.90 | 1.000 |
| 2740.201 | COL3A1 | 459 | 488 | GAKGEDGKDGSPGEPGANGLPGAAGEF | GAKGEDGKDGSP[113]GEP[113]GANGLP[113]GAAGERGAP[113] | GAKGEDGKDGSP(1.0000)GEP(1.0000)GANGLP(1.0000)GAAGERGAP(1.0000) | 30.69 | 11.71 | 1.000 |
| 2741.202 | COL3A1 | 459 | 488 | GAKGEDGKDGSPGEPGANGLPGAAGEF | GAKGEDGKDGSP[113]GEP[113]GAN[115]GLP[113]GAAGERGAP[ | GAKGEDGKDGSP(1.0000)GEP(1.0000)GANGLP(1.0000)GAAGERGAP(1.0000) | 22.84 | 20.14 | 1.000 |
